# Deep evolutionary origin of nematode SL2 *trans*-splicing revealed by genome-wide analysis of the *Trichinella spiralis* transcriptome

**DOI:** 10.1101/642082

**Authors:** Marius Wenzel, Christopher Johnston, Berndt Müller, Jonathan Pettitt, Bernadette Connolly

**Affiliations:** School of Medicine, Medical Sciences and Nutrition, University of Aberdeen, Institute of Medical Sciences, Foresterhill, Aberdeen, AB25 2ZD, UK; Centre of Genome-Enabled Biology and Medicine, University of Aberdeen, 23 St Machar Drive, Aberdeen AB24 3RY, UK

**Author notes:** Corresponding Author: Jonathan Pettitt, School of Medicine, Medical Sciences and Nutrition, University of Aberdeen, Institute of Medical Sciences, Foresterhill, Aberdeen, AB25 2ZD, UK.

**Keywords:** spliced leader *trans*-splicing, polycistronic RNA processing, eukaryotic operons, RNA splicing, nematode genome evolution

## Abstract

Spliced leader *trans*-splicing is intimately associated with the presence of eukaryotic operons, allowing the processing of polycistronic RNAs into individual mRNAs. Most of our understanding of spliced leader *trans*-splicing as it relates to operon gene expression comes from studies in *C. elegans*. In this organism, two distinct spliced leader *trans*-splicing events are recognised: SL1, which is used to replace the 5’ ends of pre-mRNAs that have a nascent monomethyl guanosine cap; and SL2, which provides the 5’ end to uncapped pre-mRNAs derived from polycistronic RNAs. Limited data on operons and spliced leader *trans*-splicing in other nematodes suggested that SL2-type *trans*-splicing is a relatively recent innovation, associated with increased efficiency of polycistronic processing, and confined to only one of the five major nematode clades, Clade V. We have conducted the first transcriptome-wide analysis of spliced leader *trans*-splicing in a nematode species, *Trichinella spiralis*, which belongs to a clade distantly related to Clade V. Our work identifies a set of *T. spiralis* SL2-type spliced leaders that are specifically used to process polycistronic RNAs, the first examples of specialised spliced leaders that have been found outside of Clade V. These *T. spiralis* spliced leader RNAs possess a perfectly conserved stem-loop motif previously shown to be essential for polycistronic RNA processing in *C. elegans*. We show that this motif is found in specific sets of spliced leader RNAs broadly distributed across the nematode phylum. This work substantially revises our understanding of the evolution of nematode spliced leader *trans*-splicing, showing that the machinery for SL2 *trans*-splicing evolved much earlier during nematode evolution than was previously appreciated, and has been conserved throughout the radiation of the nematode phylum.

## INTRODUCTION

The organisation of multiple genes into a single transcriptional control unit, termed an operon, is a common, though sparsely distributed, gene expression strategy in eukaryotes (Blumenthal 2004; Douris et al. 2010; Danks et al. 2015). Although the general adaptive significance of this mode of gene organisation remains uncertain (Zaslaver et al. 2011; Danks et al. 2015), a common feature in all cases is the presence of spliced leader *trans*-splicing. This derived version of *cis*-splicing allows the addition of short, ‘spliced leader’ RNAs to the 5’ ends of pre-mRNAs via an intermolecular splicing event (Lasda and Blumenthal 2011).

Spliced leader *trans*-splicing is an essential element in the generation of mRNAs derived from genes situated downstream of the first gene in an operon since the spliced leader RNA provides the 5’ cap for such mRNAs, allowing the *trans*-spliced mRNAs to be recognised by the translation machinery. Addition of the spliced leader is also thought to prevent termination of transcription following polyadenylation of the upstream mRNA (Evans et al. 2001; Lasda et al. 2010). Thus, spliced leader *trans*-splicing is an essential prerequisite for the evolution of eukaryotic operons.

Spliced leader *trans*-splicing and operon organisation are best understood in the nematode *C. elegans* (Blumenthal et al. 2015; Blumenthal 2012). At least 84% of *C. elegans* genes encode mRNAs that are spliced leader *trans*-spliced, with approximately 9% of these arising from downstream genes in operons (Allen et al. 2011; Tourasse et al. 2017). There are two functionally distinct types of spliced leaders in *C. elegans*. The first to be discovered, SL1, is acquired by mRNAs through a *trans*-splicing reaction that removes the 5’ UTR, also known as the ‘outron’, of capped pre-mRNAs derived from monocistronic genes and the first genes in operons. SL1 *trans*-splicing serves to sanitise the 5’ ends of mRNAs, and thus impacts their translational efficiency (Yang et al. 2017). The other spliced leader type, SL2, is added to mRNAs encoded by downstream operonic genes, which are otherwise uncapped and thus cannot be translated without SL2 *trans*-splicing (Spieth et al. 1993).

The two spliced leader RNA types have differentiated biochemical interactions (Evans et al. 2001; MacMorris et al. 2007). SL2 RNAs have a motif present in the third stem-loop that is not present in SL1 RNA (Evans and Blumenthal 2000). The sequence composition of this stem-loop is essential for the specificity of SL2 *trans*-splicing to downstream operonic gene pre-mRNAs, as well as its interaction with Cleavage Stimulation Factor (CSTF) 2 (Evans et al. 2001), the factor involved in coordinating polyadenylation of the upstream pre-mRNA with the spliced leader *trans*-splicing of the downstream pre-mRNA.

*C. elegans* is a member of Clade V, one of five major clades into which the nematodes are divided. Other species within this clade also use SL2-type *trans*-splicing to resolve polycistronic RNA into mRNAs (Lee and Sommer 2003; Evans et al. 1997). In contrast, other nematode clades appear to lack SL2-type *trans*-splicing, using the same set of spliced leaders for the processing of all pre-RNAs, regardless of their origin as monocistronic or polycistronic RNAs (Guiliano and Blaxter 2006; Ghedin et al. 2007). This led to the view that SL2 *trans*-splicing is a relatively recent innovation, associated with a more efficient processing of polycistronic RNAs (Blumenthal 2012) that evolved only in Clade V nematodes. However, the patterns of spliced leader RNA usage in the other major nematode clades have not been investigated in the detail needed to properly interrogate this hypothesis.

*Trichinella spiralis* is a vertebrate parasitic nematode that belongs to the Dorylaimia, one of three major nematode classes. *C. elegans* and all other nematodes in which polycistronic processing has been studied are located in the class Chromadoria. Thus, the lineages that led to *T. spiralis* and *C. elegans* separated early on during the radiation of the nematode phylum, so studies of polycistronic processing in *T. spiralis* allows us to investigate the extent to which this process is conserved across the nematode phylum.

The genome of *T. spiralis* encodes at least 15 distinct spliced leader RNAs (Tsp-SLs). The spliced leader sequences show a high degree of sequence polymorphism, and no sequence similarity with *C. elegans* SL1 or SL2 (Pettitt et al. 2008). The sequence diversity of these SL RNAs might simply be a consequence of unconstrained sequence variation but could also reflect functional differences. Here we describe the results of a transcriptome-wide investigation of spliced leader usage in the muscle larva of *T. spiralis*. We show that this nematode possesses a set of spliced leaders that, like *C. elegans* SL2, are specialised for the processing of pre-mRNAs derived from downstream genes in operons. These spliced leaders share a motif in their third stem-loops that is identical to that found in *C. elegans* SL2, suggesting that their specificity arises from the same conserved CSTF2 interaction. This feature is found in spliced leader RNAs in multiple nematodes lineages, and our data demonstrate for the first time that SL2-type spliced leader *trans*-splicing is broadly distributed throughout the nematode phylum, thus providing a means for the genome-wide discovery of nematode operons that was previously confined to only a small group of nematodes. The remarkable conservation of the SL2 *trans*-splicing machinery has important implications, showing that the evolution of SL2 *trans*-splicing predated the radiation of the nematode phylum, significantly improving our understanding of the evolution of nematode polycistronic RNA processing.

## RESULTS

### Conservation of the *C. elegans* SL2 RNA third stem-loop motif in Clade I nematodes

Analysis of the predicted secondary structures of the fifteen known *T. spiralis* spliced leader RNAs revealed that three, *Tsp*-SL2, 10 and 12, have predicted stem-loops that share an invariant motif (Fig 1A). Strikingly, this motif is identical to that found in *C. elegans* SL2, which is essential for the specialised activity of this family of spliced leaders (Evans and Blumenthal 2000; Evans et al. 2001). Inspection of the predicted structures of spliced leader RNAs from other Clade I nematodes, *Trichinella pseudospiralis, Trichuris muris, Soboliphyme baturini* and *Prionchulus punctatus* as well as Clade IV nematodes, *Bursaphelenchus xylophilus* and *Strongyloides ratti*, showed that this motif is broadly conserved in the predicted third stem-loops of a subset of nematode spliced leader RNAs (Supplemental Fig S1). The striking conservation of this motif suggested to us the possibility of functional conservation between these spliced leader RNAs and *C. elegans* SL2 RNA. To investigate this hypothesis, we identified spliced leader usage in the transcriptome of *T. spiralis* muscle larvae, with the aim of determining whether *Tsp*-SL2, 10 and 12 were indeed functionally distinct, and associated with *trans*-splicing to mRNAs produced by downstream operonic genes.

**Figure 1:**
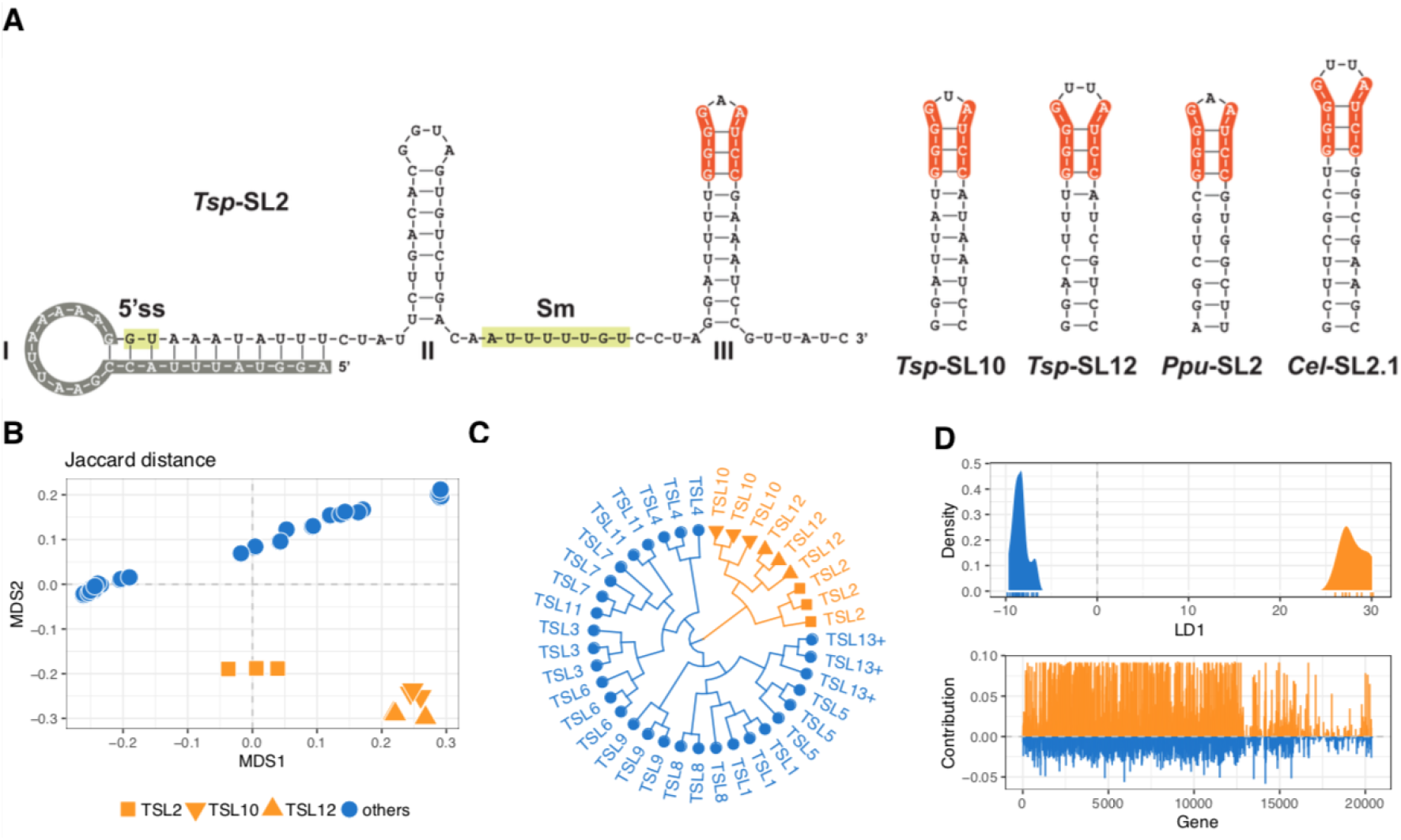
Functional similarities between *trans*-spliced leader RNA sequences *Tsp*-SL2, *Tsp*-SL10 and *Tsp*-SL12. (A) Secondary structure predictions of SL2-type spliced leader RNAs in *T. spiralis* highlight conserved residues (orange shading) of stem-loop III across *Tsp*-SL2, *Tsp*-SL10 and *Tsp*-SL12, as well as homologous SL2-type structures predicted for *Prionchulus punctatus* (*Ppu*-SL2) and *Caenorhabditis elegans* (*Cel*-SL2.1). *Tsp*-SL2 is shown in full length with numbered stem loops (I, II, III) and background shading indicating the spliced leader sequence (grey) as well as the donor splice site (5’ss) and Sm-binding site (green). (B) Multi-dimensional scaling (MDS) plot of gene-based Jaccard distances based on presence/absence of spliced leader reads (*Tsp*-SL1-15; *Tsp*-SL is abbreviated to ‘TSL’ for clarity) from three replicate RNAseq libraries. Data are based on *de novo* gene annotations (BRAKER+TRINITY exon-corrected) and 8-bp *Tsp*-SL read classification stringency (see main text). (C) Hierarchical clustering (Ward’s criterion) of gene-based Jaccard distances. Note that TSL13+ contains *Tsp*-SL13-15, which are not distinguishable at 8bp screening stringency. (D) Linear discriminant analysis of gene-based presence/absence information between read sets containing *Tsp*-SL2, *Tsp*-SL10 or *Tsp*-SL12 versus all other *Tsp*-SLs. The density of the first linear discriminant function is plotted and coloured by group. Below, the contribution of each gene to the group discrimination is plotted and coloured according to the direction along the discriminant function.

### Transcriptome assembly and *Tsp*-SL read screening

We sequenced the transcriptomes of three independent pools of approximately 100,000 *T. spiralis* L1 muscle larvae, generating 201,164,867 high-quality read pairs across all pools (Supplemental Table S1). Of these read pairs, 84.19 % mapped concordantly end-to-end to the *T. spiralis* reference genome. We used these mapped reads to assemble *de novo* gene annotations using four different approaches that yielded an average of 17,574 genes (see Methods). The *T. spiralis* reference annotations contain 16,380 genes composed of 87,853 non-redundant exons (Mitreva et al. 2011); our RNA-Seq reads covered 15,025 of these genes, suggesting that 92% of the reference genes are expressed in *T. spiralis* L1 muscle larvae. However, no more than 49.6% of reference exons and 10.8% of reference transcripts matched our *de novo* annotations, indicating that the reference annotations do not accurately reflect the gene structures suggested by our RNA-Seq data. Since accurate gene structures are crucial for the robustness of *Tsp*-SL screening and operon identification, we carried out all further core analyses on both the reference annotations and our four sets of *de novo* annotations.

In order to identify RNA-Seq reads that were candidates for containing spliced leader sequences, we investigated the 2.85% of read pairs that contained a single unmapped read. Of these unmapped reads, 6.14% (0.18 % of all read pairs) could be unambiguously assigned to one of 15 known *Tsp*-SL types, based on the characteristic 10 bp sequence at the 3’ end of the spliced leader sequence (Supplemental Table S1). The numbers of read pairs per *Tsp*-SL type varied considerably, ranging from 1,290 (*Tsp*-SL9) to 58,838 (*Tsp*-SL11) (Supplemental Table S1).

Since the numbers of recovered *Tsp*-SL reads were rather low because dependence of ≥10 bp matches to identify *Tsp*-SL-containing reads meant that reads with a shorter *Tsp*-SL sequence would be discarded, we undertook a second screen using relaxed stringency that required matches of ≥8 bp from the 3’ end of the spliced leader sequence (8-bp dataset). This stringency could no longer distinguish between *Tsp*-SL13, 14 and 15, but it increased the number of reads identified to 8.93 % of candidate read pairs (0.25 % total read pairs) and the number of read pairs per *Tsp*-SL type from 1,722 (*Tsp*-SL9) to 97,730 (*Tsp*-SL6) (Supplemental Table S1). We carried out all analyses using both datasets with qualitatively similar results, but focus on results from the 8-bp dataset since it contains more data.

The screened read pairs mapped back to the genome at rates of 91.55–98.94 % (properly paired 79.17–96.98%), and the *Tsp*-SL-containing single-end read alignments were unambiguously assigned to single exon annotations at rates of 48.76–73.49% for the reference annotations, and 53.77–87.95% for *de novo* annotations (Supplemental Table S1). Based on the 8 bp match reads, the percentage of *T. spiralis* genes whose mRNAs are spliced leader *trans*-spliced ranges from 16.4% to 22.9%. Since our approach discards *bona-fide* spliced leader-containing reads that have shorter matches than 8 bp, these figures are underestimates. Nevertheless, it is clear that the proportion of genes that generate mRNAs subject to SL *trans*-splicing is considerably smaller in *T. spiralis* compared to the 80 - 90% reported for other nematodes (Wang et al. 2017; Sinha et al. 2014; Allen et al. 2011; Tourasse et al. 2017).

### Genome-wide patterns of *Tsp*-SL *trans*-splicing suggest two functionally distinct spliced leader RNA classes

The presence of the conserved motif found in the third stem-loops of *Tsp*-SL2, 10 and 12 RNAs raised the possibility that they might have distinct patterns of *trans*-splicing. In order to investigate this, we systematically compared the sets of expressed genes associated with each spliced leader type using the *Tsp*-SL read counts and gene annotations generated above. Pairwise distance comparisons between spliced leaders based on read presence/absence across all genes (Jaccard distance) indicated high consistency in the patterns of spliced leader usage between the three replicate sequencing libraries (Figs 1B and C). Strikingly, we found that the fifteen spliced leaders could be grouped into two major clusters, one composed of *Tsp*-SL2, −10 and −12, and the other containing all other spliced leaders (Figs 1B and C). The differentiation signal between the two spliced leader clusters was supported by mRNAs derived from a large number of genes across the whole genome, showing that the two groups are robustly differentiated (Fig 1D). This clustering behaviour indicates that *Tsp*-SL2, −10 and −12 are functionally distinct from the other *T. spiralis* spliced leaders.

The spliced-leader usage patterns were robust across all genome annotation sets and also between read presence/absence information and normalized read counts information. However, the degree of association between *Tsp*-SL2 and *Tsp*-SL10/12 varied such that *Tsp*-SL2 was sometimes not clustered with *Tsp*-SL10/12, particularly when using normalised read counts (Supplemental Fig S2). This suggests that although they share similar activity, *Tsp*-SL2 RNAs may have some distinct properties compared to *Tsp*-SL10 and *Tsp*-SL12 RNAs.

### Three spliced leaders, *Tsp*-SL2, *Tsp*-SL10 and *Tsp*-SL12, define mRNAs derived from downstream genes in *T. spiralis* operons

The striking difference between *Tsp*-SL2, −10 and −12, and the other known *T. spiralis* spliced leaders in terms of both RNA structure and general *trans*-splicing patterns could be explained if these spliced leaders were targeted specifically to mRNAs derived from downstream operonic genes. To investigate this possibility, we manually examined the mapping locations of *Tsp*-SL-containing reads for a set of putative operons in *T. spiralis* defined on the basis of their syntenic conservation in *C. elegans* and other nematodes (Pettitt et al. 2014; Johnston 2018). We curated this dataset, selecting only those pairs of genes where the predicted AUG of the downstream gene is within 1000 bp of the stop codon of the upstream gene, giving us a set of 45 putative operons (Supplemental Table S2). We investigated the correlation between *Tsp*-SL2, −10 and −12 read mapping locations and the upstream or downstream genes predicted from this dataset.

Although in almost all cases the mapping locations of the reads led to considerable revisions of the predicted gene structures involved in each putative operon (Fig 2), we observed a strong correlation between the identity of the *Tsp*-SL added to the mRNA and the position of the corresponding gene in the operon (Supplemental Table S2). All predicted downstream operonic genes produced mRNAs that were *trans*-spliced almost exclusively to *Tsp*-SL2, −10 or −12 (Fig 2; Supplemental Table S2). In contrast, predicted upstream operonic genes, if they were *trans*-spliced, generally received one of the other *T. spiralis* spliced leaders, though in many cases as many as 10% of reads contained *Tsp*-SL2, −10 or −12.

**Figure 2.**
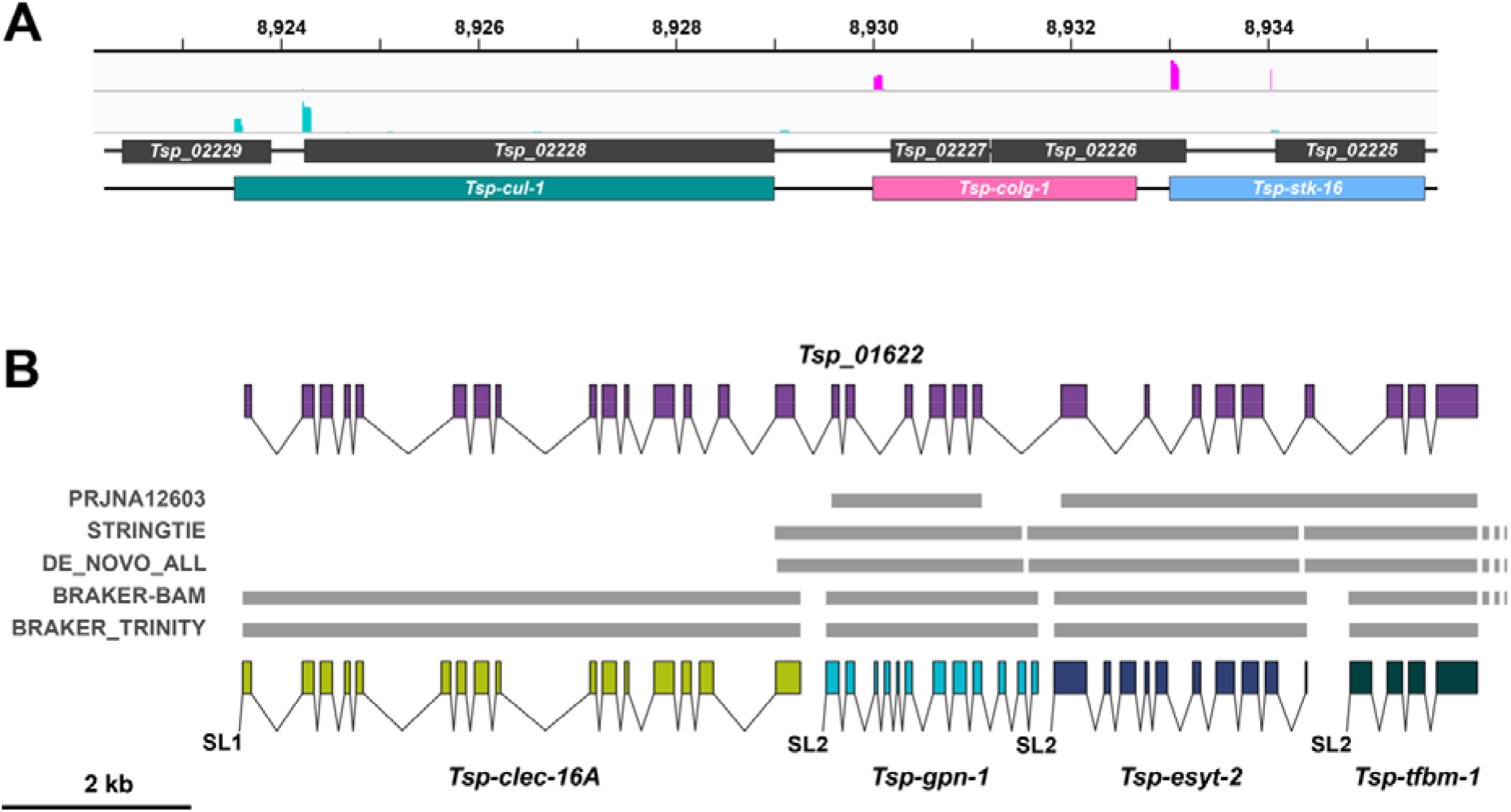
Spliced leader read mapping leads to major revisions of gene annotations. (A) Example of original gene predictions (represented as dark shaded boxes) involving five genes that are shown to be a three gene operon upon mapping of *Tsp*-SL-containing reads (read depth for both tracks is scaled to 80; cyan peaks indicate SL1-type reads; pink, SL2-type reads). Shown below are the revised *T. spiralis* gene predictions, named based on their *C. elegans* and/or human orthologues. Note the exon/intron structure of the genes is not shown. X-axis indicates distance in kilobases along Scaffold GL622787. (B) Benchmarking operon prediction pipelines. The structure of the original gene annotation is shown on the top line. The revised, manually curated gene predictions (named based on *C. elegans* and/or human orthologues) informed by sequence reads and sequence similarity are shown on the bottom line. Grey bars indicate the gene predictions generated using five different approaches (dashed lines indicate predictions that extend beyond the region shown). Exons are indicated by coloured boxes and lines indicate splicing of introns. Spliced leader transsplicing is indicated by a single line, with ‘SL1’ indicating SL1-type *trans*-splicing and ‘SL2’ indicating transsplicing to SL2-type spliced leaders.

These patterns immediately suggest that the *Tsp*-SL RNAs fall into two functional classes. *Tsp*-SL2, −10 and −12 appear to be the only spliced leaders routinely used in the resolution of downstream operon transcripts. This is consistent with the conservation of the motif in the third stem-loop only in these SL RNAs (Fig 1A; Supplemental Fig S1), suggesting that they are likely to be recruited to the intercistronic region between open reading frames in polycistronic RNAs (Evans et al. 2001; Evans and Blumenthal 2000). We thus refer to *Tsp*-SL2, 10 and 12 as SL2-type spliced leaders (Supplemental Table S2). Since all the other *Tsp*-SL RNAs are *trans*-spliced almost exclusively to mRNAs derived from monocistronic and upstream operonic genes, we classify these SL RNAs as SL1-type (Supplemental Table S2).

Genome-wide classification of genes showed that they fell into two categories based on whether they produced transcripts that were *trans*-spliced to either predominantly SL1-type or SL2-type spliced leaders (Fig 3A). However, there initially appeared to be two subcategories of SL1-type *trans*-spliced genes: 1573 genes apparently produced transcripts *trans*-spliced exclusively to SL1-type spliced leaders; while 967 genes encoded transcripts that were *trans*-spliced to SL2-type spliced leaders in up to 10% of cases (Fig 3A). Analysis of read depths for these two datasets revealed that the “SL1-type only” category of genes was mostly an artefact of read depth, such that there were too few reads to detect any SL2-type *trans*-spliced transcripts (Fig 3B). Thus, transcripts derived from monocistronic or upstream genes showed a strong preference for SL1-type spliced leaders, but approximately 10% of these mRNAs were *trans*-spliced to SL2-type spliced leaders (Fig 3C). Strikingly, in this subset of mRNAs, the SL2-type spliced leader used was more frequently *Tsp*-SL2, compared to *Tsp*-SL10 or 12, suggesting some functional distinction between the SL2-type spliced leader RNAs (Fig 3D).

**Figure 3:**
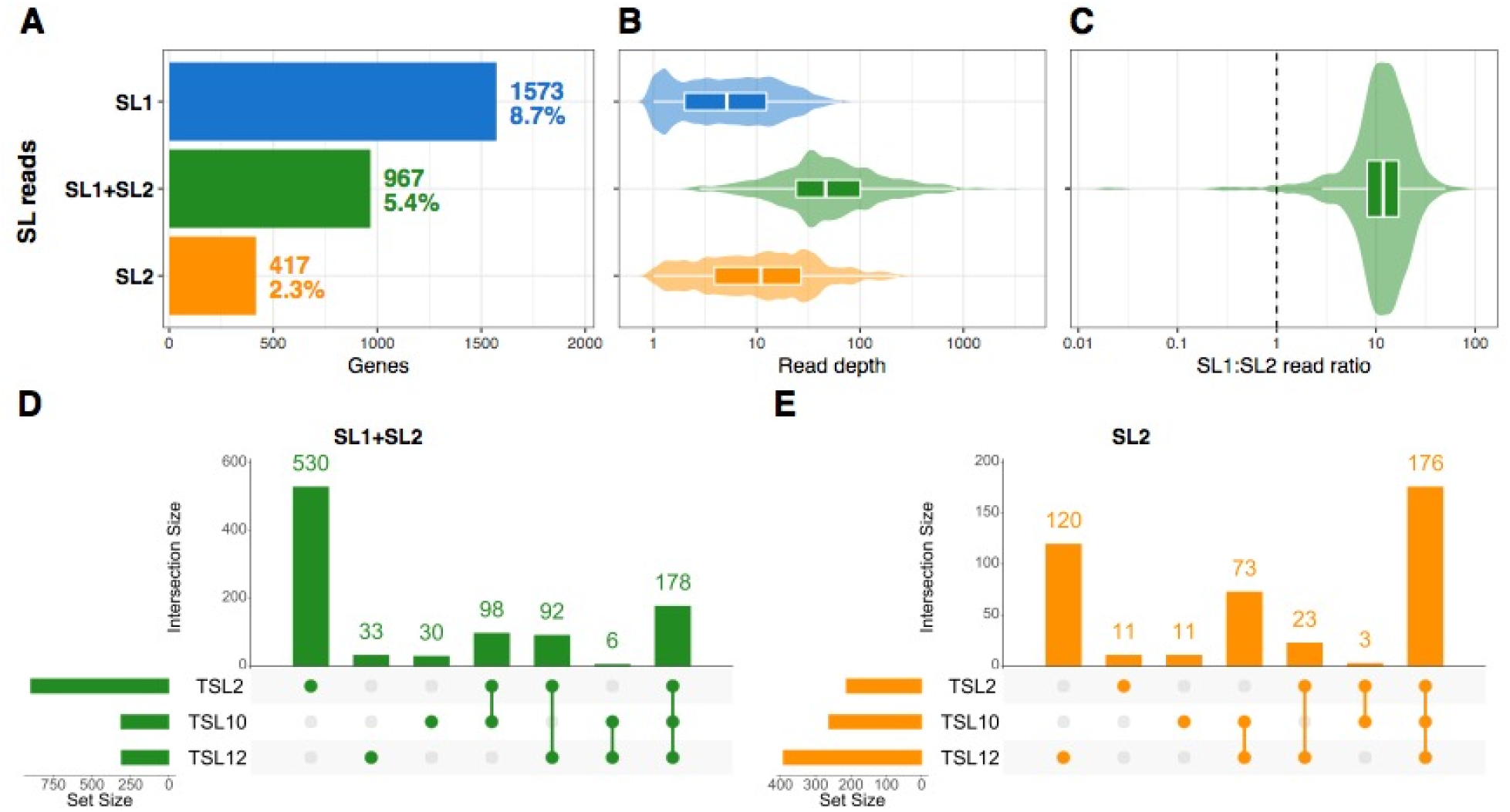
Genome-wide analysis identifies differential usage of *T. spiralis* spliced leader RNAs. Genome-wide identification of *T. spiralis* genes receiving either SL2-type spliced leaders (*Tsp*-SL2, *Tsp*-SL10 and *Tsp*-SL12), SL1-type spliced leaders (all other **Tsp*-SLs)* or a mixture of both types (“SL1+SL2”). A) Numbers and fractions of expressed genes allocated to each category. B) Distributions of spliced-leader read depth in each category. C) Distribution of read-depth ratio between SL1- and SL2-type spliced-leaders in genes receiving both types. (D, E) Intersection plots illustrating the usage patterns of *Tsp*-SL2, *Tsp*-SL10 and *Tsp*-SL12 across genes that receive both SL1-type and SL2-type spliced leaders (D), or SL2-type only (E). Data are based on *de novo* gene annotations (BRAKER+TRINITY exon-corrected) and 8-bp *Tsp*-SL read classification stringency (see main text).

In contrast to SL1-type *trans*-spliced genes, genes which encoded transcripts that were SL2-type *trans*-spliced were only rarely *trans*-spliced to SL1-type spliced leaders. We were able to confirm that this was not an artefact of read depths (Fig 3B), by investigating the top 100 downstream operonic genes ranked on the basis of their SL2-type read coverage (ε 89 reads): the mean read counts for SL1- and SL2-type reads in this dataset were 0.43 and 198.21, respectively. Thus, the low proportion of downstream gene transcripts *trans*-spliced to SL2-type spliced leaders suggests that SL1-type spliced leader RNAs are very poor substrates for the *trans*-splicing of pre-mRNAs from downstream operonic genes.

We also found differences in the frequencies with which the three SL2-type spliced leaders are *trans*-spliced to downstream operonic mRNAs (Fig 3E). *Tsp*-SL10 and 12 appear to be more likely to be added to these mRNAs than *Tsp*-SL2. These patterns confirm that the functional differentiation between the two groups of spliced leaders is not complete and that the three SL2-type spliced leader RNAs may also be functionally subdivided, consistent with the initial results from the multivariate analysis (Figs 1B and D).

### Genome-wide prediction of operons in *T. spiralis*

Eukaryotic operons are challenging to unambiguously identify solely on the basis of polycistronic transcription or gene organisation (Pettitt et al. 2014; Blumenthal 2004). However, the overwhelming correlation between downstream genes in our set of 47 operons and *trans*-splicing of their corresponding mRNAs to SL2-type spliced leaders provides a powerful method to systematically identify the extent of operon organisation in the *T. spiralis* genome.

We inferred operons on the basis that all genes receiving only SL2-type spliced leaders should, by definition, be downstream operonic genes. By this criterion an operon would consists of an uninterrupted series of consecutive SL2-type spliced leader-associated (i.e. downstream operonic) genes on the same DNA strand, together with a single gene immediately upstream of the tract of downstream genes, irrespective of its *Tsp*-SL status.

The utility of this strategy depends entirely on the quality of the genome annotations used to quantify the mapping locations of *Tsp*-SL-containing RNAseq reads. As stated previously, the reference gene structures correlated poorly with the gene structures suggested by our RNAseq data, leading us to assemble four alternative sets of *de novo* annotations. A further complication was noted when quantifying *Tsp*-SL reads at the exon level instead of the gene level. As we observed during the manual curation of our set of 47 candidate operons, as well as our genome-wide analysis, *Tsp*-SL-containing reads frequently mapped to exons that were annotated as being internal, rather than the first exon of the gene prediction. This was a particular issue when dealing with operons that were incorrectly predicted as single genes (Fig 2B). To alleviate this problem, we assumed that each *Tsp*-SL-receiving exon demarcates the beginning of a new gene and corrected the gene annotations accordingly (see Methods). When correcting all sets of annotations in this way, we gained 250 to 1,103 genes per dataset (Supplemental Table S3). Of all 20 datasets (five sets of gene annotations, two *Tsp*-SL screening stringencies, and using exon- or gene-based gene annotations), we found that the BRAKER-Trinity gene set with 8 bp *Tsp*-SL screening stringency and exon-based gene corrections is the most robust set based on its ability to predict our manually predicted operon set, though none of the datasets was perfectly consistent with known operon structures (Fig 2).

On average across all 20 datasets, our strategy identified 308 operons comprising 634 genes (Table 1). Datasets using the reference gene annotations predicted somewhat fewer operons than *de novo* annotations, though not significantly so (Welch’s *t*-test: P=0.129) (Table 1; Supplemental Table S3). *Tsp*-SL screening stringency had no effect on operon prediction (P>0.71) (Table 1; Supplemental Table S3). Correcting gene annotations by splitting genes at internal exons based on distinct peaks of *Tsp*-SL reads considerably increased (P<0.001) the numbers of operons and operonic genes (Table 1). Consistent with the low proportion of SL2-type-only genes (Fig 3A), the number of inferred operonic genes was much smaller than non-operonic (monocistronic) genes (Fig 4A).

**Figure 4:**
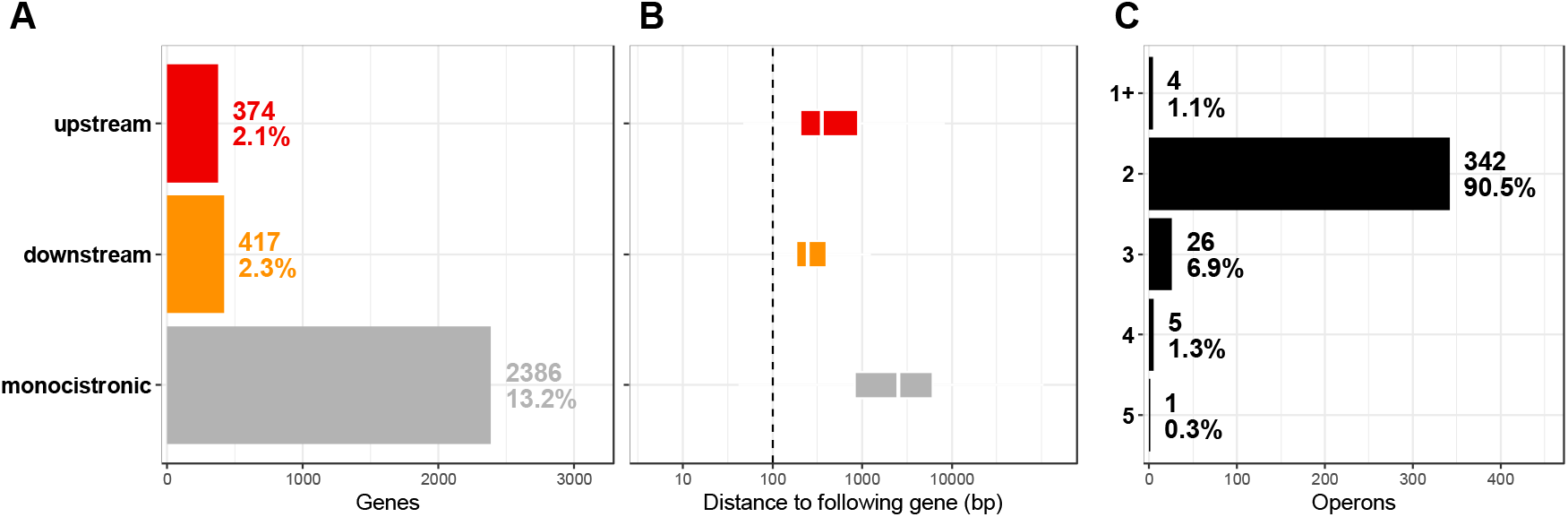
Genome-wide identification of *T. spiralis* operons. (A) Numbers and proportion of expressed genes that are predicted to be upstream or downstream operonic or monocistronic (*trans*-spliced, but not operonic). B) Distributions of physical intergenic distances in the three gene classes (upstream: distance between first and second gene in operon; downstream: distance between downstream genes in same operon; monocistronic: distance to following gene, *trans*-spliced or not). C) Numbers of inferred operons of particular sizes (1+ operons comprise a single SL2-type spliced leader *trans* spliced gene without an upstream gene on the same genomic contig). Data are based on *de novo* gene annotations (BRAKER+TRINITY exon-corrected) and 8-bp *Tsp*-SL read classification stringency (see main text).

**Table 1:**
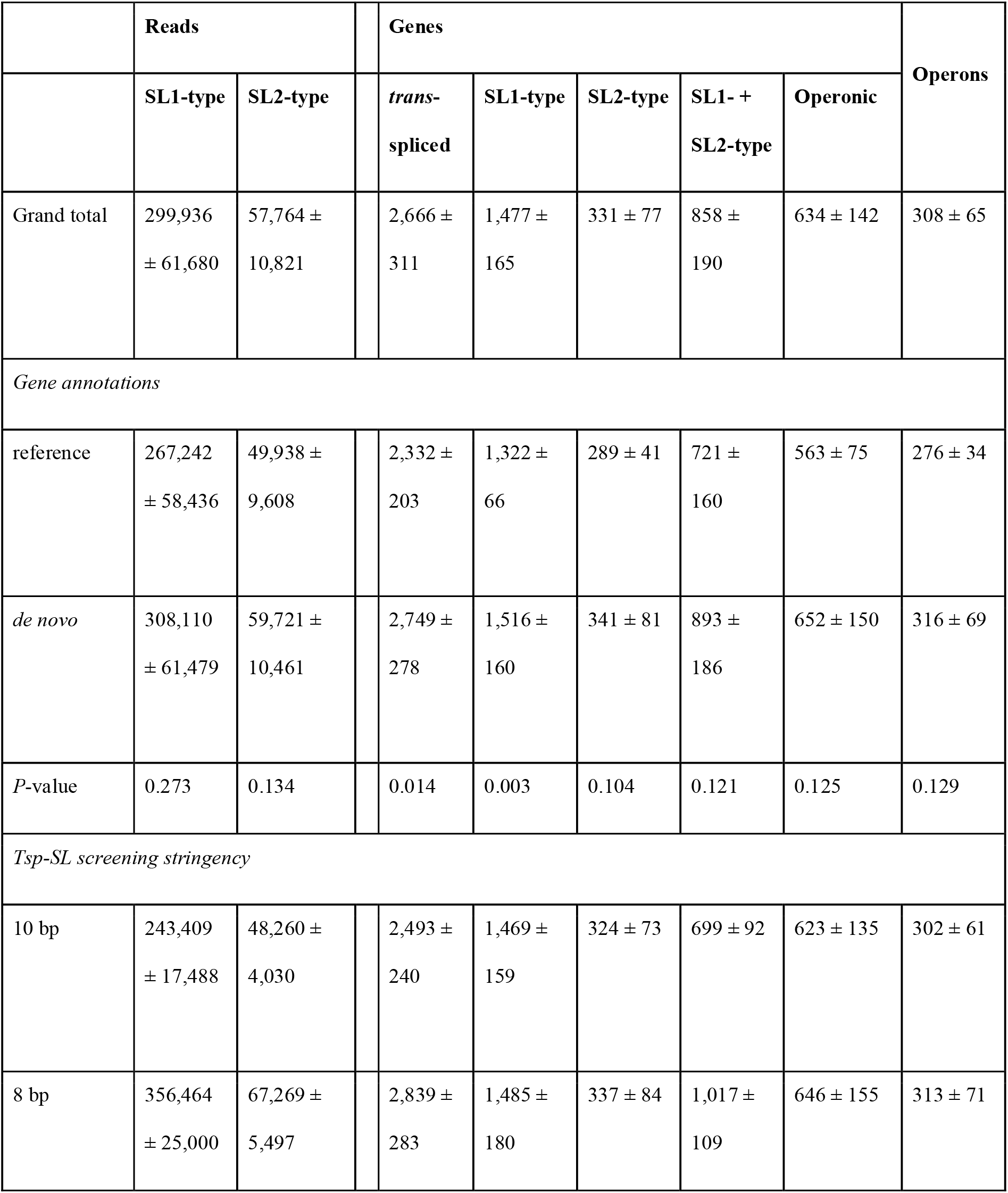
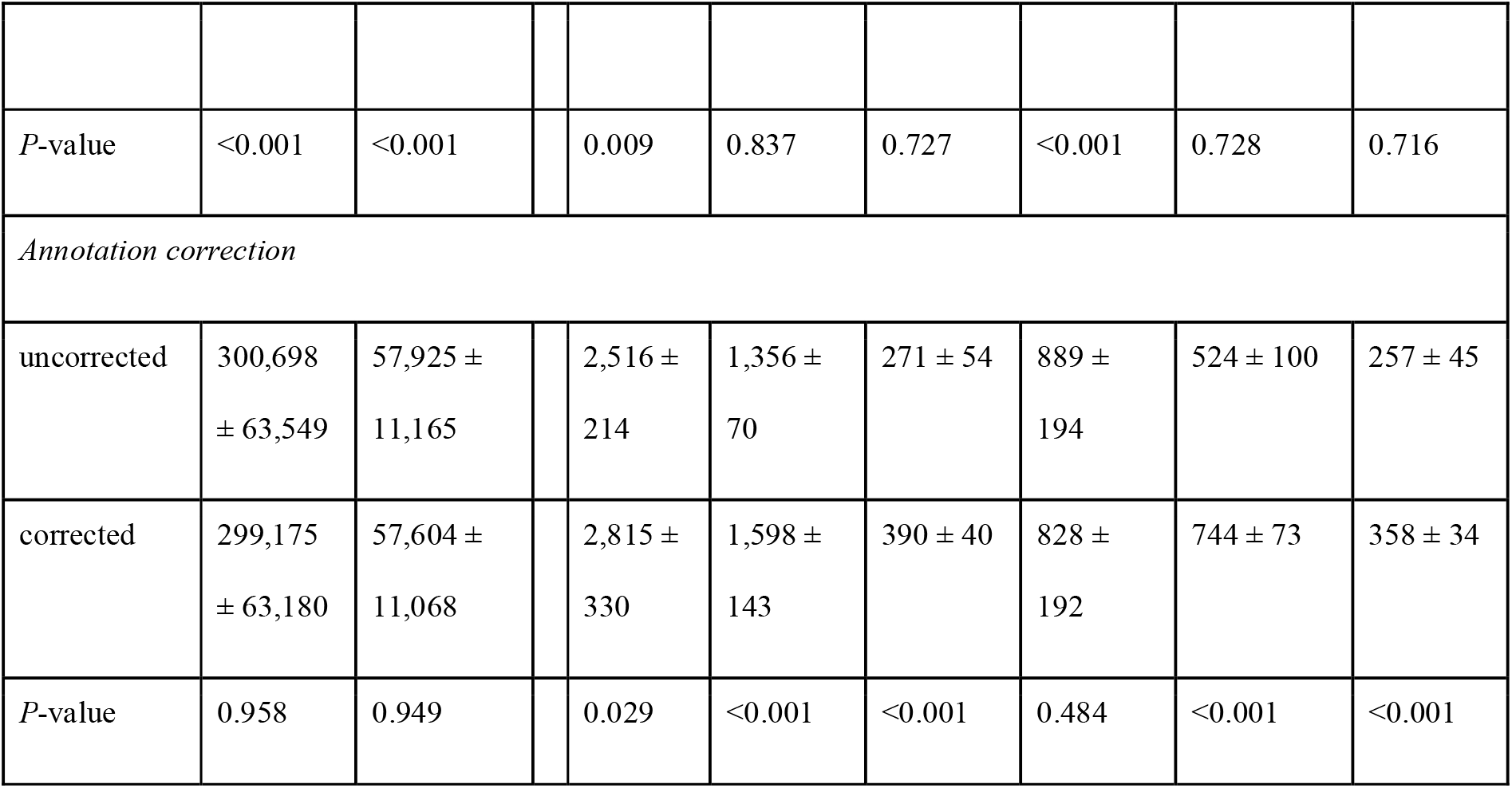
Summary of gene classification by *Tsp*-SL type and prediction of operonic organisation. Numbers of reads, genes and operons (all mean ± SD) are presented across 20 datasets and split by gene-annotation type, *Tsp*-SL-screening stringency or the use of annotation correction via **Tsp*-SL* reads at internal exons. Means were compared between groups using Welch’s *t*-tests, and the *P*-values are presented for each comparison.

A key feature of known nematode operons is that the distance between coding regions, the intercistronic region, is unusually short. For example, in *C. elegans* most intercistronic distances are 50 - 200 nucleotides, and approximately 80% of intercistronic regions are less than, or equal to 500 nucleotides (Sinha et al. 2014). Consistent with this, we found that within *T. spiralis* operons the intercistronic distances were substantially reduced compared to non-operonic (monocistronic) genes, with a median of 246 nucleotides (Fig 4B). In contrast, *Tsp*-SL-associated, monocistronic genes were a median of 2,475 nucleotides apart and showed no difference in intergenic distance compared to non-*Tsp*-SL-receiving genes.

The vast majority of predicted operons consisted of two genes (Fig 4C), and the maximum operon length observed in some datasets was five genes (Supplemental Table S3). We detected up to seven cases per dataset where a strictly SL2-type-receiving gene had no upstream gene (Supplemental Table S3) due to the fragmented nature of the genome assembly.

### Functional characterisation of predicted operonic genes

Previous studies have shown that *C. elegans* operons preferentially contain genes encoding molecules critical to basic eukaryotic cell biology (Blumenthal 2004; Blumenthal and Gleason 2003), and RNAi studies show that operons are enriched for genes associated with observable loss-of-function phenotypes (Kamath et al. 2003). Operonic genes are also more likely to be expressed in the hermaphrodite germline compared to monocistronic genes (Reinke and Cutter 2009).

To investigate whether similar patterns exist for *T. spiralis* operonic genes, we examined the association of operonic genes with defined biological processes. We identified 1,332 *T. spiralis* genes whose functions are predicted to be associated with germline processes, based on orthology to genes expressed in the *C. elegans* germline (see Methods). Of these, 11–27 % (depending on the set of operon predictions used) correspond to genes predicted to reside in operons (Supplemental Table S4). This proportion is significantly larger than the genomic background rate of 3–6 % operonic genes (binomial test: P ≪ 0.001), showing that the enrichment of germline expressed genes in operons is likely a general feature of nematode operons.

Additionally, between 6,758 and 14,444 genes were mapped to 52,055–69,177 UNIPROT proteins, which yielded 2,690–3,210 unique GeneOntology annotations (Supplemental Table S5). Across all datasets, 117 unique terms in the biological process ontology were significantly overrepresented among operonic genes. These were simplified to 39 representative terms following clustering by semantic similarity. All these terms represent essential cellular processes, for example RNA modification and metabolism, cellular component organisation/localisation, protein modification and cellular respiration (Supplemental Table S5). This is consistent with a view that operonic genes are predominantly involved in general cellular metabolism and regulation of gene expression.

## DISCUSSION

Operons and spliced leader *trans*-splicing are conserved features of nematode genomes (Wang et al. 2017; Pettitt et al. 2014; Blumenthal 2012). Spliced leader *trans*-splicing is essential for the processing and expression of mRNAs derived from downstream operonic genes. Previous studies of polycistronic RNA resolution by spliced leader *trans*-splicing, which have focused almost exclusively on *C. elegans*, show that the maturation of pre-mRNAs derived from downstream operonic genes involves the *trans*-splicing of a specialised set of spliced leader RNAs, the SL2 family (Spieth et al. 1993). These SL2 RNAs are recruited to the intercistronic regions of polycistronic RNAs through a mechanism that is distinct from that involved in SL1 *trans*-splicing of the majority of *C. elegans* pre-mRNAs (Graber et al. 2007).

Prior to this study, there was limited information about polycistronic RNA processing in other nematodes, beyond a few species relatively closely related to *C. elegans* (Uyar et al. 2012; Sinha et al. 2014; Evans et al. 1997; Guiliano and Blaxter 2006). These data indicated that SL2-type *trans*-splicing was confined to the Rhabditina (Clade V), the clade containing *C. elegans* (Blaxter 2011); it was thought that in more distantly related species, the same *trans*-splicing mechanism was used to generate operonic and non-operonic mRNAs (Guiliano and Blaxter 2006).

Here we describe the first detailed analysis of spliced leader *trans*-splicing as it relates to processing of operon transcripts in a nematode outside of Clade V (Blaxter 2011). We found that three of the fifteen known *T. spiralis* spliced leaders, *Tsp*-SL2, SL10 and SL12, show a distinct pattern of *trans*-splicing and define a set of target mRNAs that correspond to genes that are downstream in operons. This suggests the existence of a specific mechanism that directs these spliced leaders to the pre-mRNAs derived from these genes. Our work provides definitive evidence that intercistron-specific *trans*-splicing, as exemplified by *C. elegans* SL2 *trans*-splicing, occurs during the pre-mRNA processing of *T. spiralis* downstream operonic genes.

Several lines of evidence indicate that *trans*-splicing of *Tsp*-SL2, 10 and 12 uses the same mechanism as the *C. elegans* SL2 family. Firstly, the motif in stem-loop 3 common to all *C. elegans* SL2 RNAs and which is specifically required for SL2 *trans*-splicing (Evans et al. 2001; Evans and Blumenthal 2000), is strictly conserved in these three *T. spiralis* SL RNAs. Secondly, we can identify features of the *T. spiralis* intercistronic regions that have properties consistent with them being Ur elements (Pettitt et al. 2014), the motifs that are essential for specifically directing SL2 *trans*-splicing to the splice acceptor site of the downstream pre-mRNA (Liu et al. 2003; Huang et al. 2001; Lasda et al. 2010). Moreover, previous studies have shown that at least one of these *T. spiralis* intercistronic regions is recognised by *C. elegans* SL2 *trans*-splicing (Pettitt et al. 2014), further supporting the functional conservation of the process between the two nematodes.

Based on the conservation of the stem-loop 3 motif, it seems likely that SL2-type SL RNAs are conserved throughout the Dorylaimia (Supplementary Fig S1), and functional evidence from previous studies supports this. Limited data for spliced leader usage and operon organisation in *T. muris* shows that the three candidate SL2-type splice leaders are *trans*-spliced to mRNAs expressed by putative downstream operonic genes (Pettitt et al. 2014). More significantly, heterologous expression of one of the *P. punctatus* putative SL2-type spliced leaders, *Ppu*-SL2 (Fig 1), in *C. elegans* results in the specific *trans*-splicing of the spliced leader onto mRNAs derived from downstream operonic genes (Harrison et al. 2010). Thus, *Ppu*-SL2 must interact specifically with the intercistronic regions of *C. elegans* polycistronic RNAs, presumably through the interaction with the polyadenylation machinery mediated by its conserved third stem-loop.

Since the spliced leader *trans*-splicing mechanism for resolving polycistronic RNAs, including the almost invariant stem-loop 3 motif, is conserved between two major nematode classes, this mechanism almost certainly arose prior to, or during early nematode evolution. No information is available regarding operon organisation or spliced leader *trans*-splicing from Enoplia, the earliest diverging nematode lineage (Koutsovoulos 2015). However, we found spliced leader RNAs with the same conserved stem-loop 3 motif in *B. xylophilus* and *S. ratti*; nematodes belonging to Clade IV, members of which have not previously been shown to carry out SL2-type *trans*-splicing. In the case of *B. xylophilus*, these spliced leaders are associated with transcripts derived from downstream operonic genes (data not shown). We were not able to identify stem-loop 3 motifs in any of the spliced leader RNAs from the remaining clade, Clade III. However, credible operons have been identified in a member of this clade, *Brugia malayi* (Ghedin et al. 2007; Guiliano and Blaxter 2006), and putative intercistronic regions have been characterised (Liu et al. 2010), including a potential Ur element, which was shown to be important for the expression of the downstream gene in a transgenic assay (Liu et al. 2010). This work showed that the Ur element behaves similarly to Ur elements in *C. elegans* polycistronic RNAs, and since Ur elements are specifically involved in recruiting SL2 RNAs to downstream pre-mRNAs (Liu et al. 2003; Lasda et al. 2010), this suggests that intercistronic region-specific spliced leader RNAs also exist in *B. malayi*. Thus, all available evidence suggests that the spliced leader *trans*-splicing mechanism specific for downstream pre-mRNAs was present during early nematode evolution and has been preserved in most, possibly all nematode clades.

Since most data on nematode spliced leader *trans*-splicing comes from *C. elegans*, our genome-wide analysis in *T. spiralis* provides the first opportunity to compare this process in two distantly related nematode lineages. The most striking difference between the two nematodes is that the proportion of genes whose mRNAs are subject to *trans*-splicing is much lower in *T. spiralis*. Although our estimate is conservative, it is clear that the actual value cannot be as high as the 80-90% of estimates for *C. elegans*. Similarly, high values have been obtained for *Pristionchus pacificus* and *Ascaris suum*, but these three nematodes belong to the same class as *C. elegans*, whereas *T. spiralis*, as stated above, belongs to a distinct, deeply divergent nematode class. Thus, the differences may simply lie in the different demographic histories of the two lineages.

Although the main finding of our work is that both *C. elegans* and *T. spiralis* employ two distinct types of spliced leader *trans*-splicing, there are differences in the usage of SL1- and SL2-type spliced leaders. Many *C. elegans* SL2 *trans*-splice sites are subject to low levels of SL1 *trans*-splicing, whereas in *T. spiralis*, downstream mRNAs are only very rarely *trans*-spliced to an SL1-type spliced leader. It is likely that this difference is simply due to the relative expression levels of the spliced leader RNAs in the two nematodes; *C. elegans* SL1 RNA is more abundant than SL2 RNA, and so is able to compete with SL2 even though it is poorly recruited to intercistronic regions (Allen et al. 2011).

Conversely, in *T. spiralis*, on average, 10% of outron-containing pre-mRNAs (which are predominantly *trans*-spliced to SL1-type spliced leaders) receive SL2-type spliced leaders. In *C. elegans*, SL2 is almost never recruited to outron-containing pre-mRNAs (Allen et al. 2011). Again, this could be explained by differences in the expression levels of the two types of spliced leader RNAs. It is notable that endogenous *C. elegans* SL2 is *trans*-spliced to SL1 targets in *rrs-1* mutants, which lack the SL1 RNA, and SL2 can rescue the lack of SL1 in these animals when overexpressed (Ferguson et al. 1996). Thus, SL2 RNAs in *C. elegans* are able to interact with outrons, albeit with less efficiency than they interact with their target intercistronic regions. Intriguingly, *Tsp*-SL2 appears to be more effective at *trans*-splicing to outrons than *Tsp*-SL10 and 12. A simple explanation would be that it is the more highly expressed of the SL2-type SL RNAs. However, expression levels cannot explain all the differences between spliced leaders, since we would then expect it to also be the dominant spliced leader *trans*-spliced to downstream pre-mRNAs, whereas we find that *Tsp*-SL12 is the most frequently selected. This suggests that there are subtle functional differences between the three SL2-type SL RNAs for which we cannot currently account.

The identification of distinct spliced leaders associated with downstream genes in operons allowed us to definitively identify operons in *T. spiralis* using the same strategy employed for *C. elegans* and the other closely related nematodes that use SL2 *trans*-splicing. Although the proportion of *T. spiralis* genes that are operonic (about 4%) is lower than in *C. elegans* (15%), the genes that are found in operons show the same general pattern, encoding basic cellular functions, and showing enrichment for germline expression. This supports the hypothesis that operons facilitate recovery from developmental arrest, a life cycle strategy that is common throughout the nematode phylum (Zaslaver et al. 2011). Similarly, the size distribution of operons in *T. spiralis*, in terms of gene number per operon, resembles those for *C. elegans* and *C. briggsae* (Uyar et al. 2012). Taken together the data suggest that the dynamics of operon formation and maintenance are broadly the same across the nematode phylum.

Operons, and the spliced leader *trans*-splicing that facilitates this mode of gene organisation, appear to have independently evolved in multiple eukaryotic lineages (Hastings 2005; Douris et al. 2010; Pettitt et al. 2010). Our data indicates that a mechanism involving spliced leaders specialised for resolving polycistronic RNAs evolved early on during nematode evolution, which clearly involved adaptation of spliced leader function to allow an interaction with the polyadenylation machinery. Although resolution of polycistronic RNA through *trans*-splicing has been described in several other animal groups (Marlétaz et al. 2008; Ganot et al. 2004; Davis and Hodgson 1997; Protasio et al. 2012; Satou et al. 2008), specialised spliced leaders of the kind exemplified by the nematode SL2-type, have not been described. The chaetognath and tunicate operons that have been described in detail resemble the minor class of *C. elegans* operons termed ‘SL1 operons’, in which the 3’ end of the upstream cistron occurs very close to or coincident with the *trans*-splice acceptor site (Williams et al. 1999). It will be important to establish whether interactions between the spliced leader *trans*-splicing and polyadenylation mechanisms are a feature of polycistronic pre-mRNA processing outside of the nematodes.

The discovery that *trans*-splicing of specialised spliced leaders to mRNAs derived from downstream operonic genes is a broadly distributed nematode trait should lead to dramatic improvements to nematode genome annotations. Conservation of the stem-loop 3 motif can be used to identify SL2-type spliced leaders. Provided that the spliced leader sequence can be diagnostically associated with this motif (which is the case for all instances that we have analysed, but *a priori* need not be), then identification of transcripts *trans*spliced to these spliced leaders would provide a definitive means to identify operons and operonic genes, an approach that has previously been confined to *C. elegans* and its close relatives, but which should see broad application in multiple nematode genomes.

In conclusion, the work presented here significantly advances our understanding of the evolution of an essential process for nematode gene expression, showing that the functionally distinct SL1 and SL2-type *trans*-splicing mechanisms were early innovations that were in place prior to the divergence of the major nematode lineages. The use of this information to improve the genome annotation of the *T. spiralis* demonstrates the importance of this approach, which can be used to similarly refine the genome annotations of multiple nematode genomes. Moreover, since we have shown that SL2-*trans*-splicing is more widely distributed in the nematodes than previously understood, this work will have impact beyond the nematodes. Spliced leader *trans*-splicing of operon-derived polycistronic RNAs is a widespread eukaryotic gene expression strategy, that likely evolved multiple times in multiple groups. The interaction of the *trans*-splicing and polyadenylation machinery exemplified by nematode SL2-*trans*-splicing may represent one of only a limited number of viable solutions to the problem of targeting a spliced leader RNA to a downstream operonic pre-mRNA.

## METHODS

### SL RNA secondary structure prediction

Secondary structure prediction of SL RNAs were performed using MFOLD Version 3.6 (Zuker 2003) using the default folding conditions and with the constraint that the putative Sm-binding sites were required to be single stranded.

### RNA-Seq library preparation and sequencing

Three independent sets of five mice were infected with *T. spiralis* muscle larvae. From the resulting infections, approximately 100,000 muscle larvae were isolated from each set. The larvae were pooled for RNA extraction with the PureLink RNA mini kit (Ambion by Life Technologies) following manufacturer’s instructions and including an on-column DNA removal step with PureLink DNase. Three unstranded Illumina TruSeq mRNA V2 libraries were prepared and sequenced on a single lane of an Illumina HiSeq 2000/2500 instrument in 101 bp paired-end mode at The Genome Analysis Centre (TGAC) in Norwich, UK (now The Earlham Institute).

Data quality was assessed using FASTQC 0.11.3 (Andrews and Others 2010), followed by trimming of Illumina adapter sequences and bases with phred quality score below 20 using TRIM_GALORE 0.4.0 (Krueger 2015). Read pairs with at least one read shorter than 30 bp after trimming were discarded.

### *Tsp*-SL read screening

Read pairs that contained one of 15 known *Tsp*-SL sequences (Pettitt et al. 2008b) at the 5’ end of one read were identified using a strategy modified from published pipelines designed for *C. elegans* (Yague-Sanz and Hermand 2018; Tourasse et al. 2017). Read pairs were mapped to the *Trichinella spiralis 3.7.1* reference genome assembly (NCBI accession GCA_000181795.2; BioProject PRJNA12603) using HISAT 2.1.0 (Kim et al. 2015) with enforced end-to-end alignment of reads and concordant mapping of read pairs. Reads containing a spliced-leader tag are expected not to map end-to-end due to the spliced leader overhang, whereas their mate read is expected to map end-to-end (Yague-Sanz and Hermand 2018). Since the RNAseq libraries were unstranded, spliced-leader tags can occur on R1 or R2 reads. Therefore, for ease of downstream processing, candidate read pairs were re-oriented such that all unmapped reads were designated as R1 reads and all mapped reads as R2 reads, using SAMTOOLS 1.6 (Li et al. 2009). Finally, the candidate read pairs were screened for 15 *Tsp*-SL sequences at their 5’ ends using CUTADAPT 1.15 (Martin 2011), enforcing a minimum perfect match of 10 bp in order to ensure unambiguous *Tsp*-SL assignment, and a minimum read length of 20 bp after trimming all matching *Tsp*-SL bases. An alternative screening was undertaken with a minimum match of 8 bp; while this recovered a larger number of *Tsp*-SL reads, *Tsp*-SL13, *Tsp*-SL14 and *Tsp*-SL15 cannot be distinguished at this stringency (Pettitt et al. 2008).

### *De novo* genome re-annotation

The concordant read-pair alignments generated from all three libraries in the first step of the *Tsp*-SL screening above were merged into a single file and used to generate *de novo* gene annotations for the reference genome. Transcript sequences were assembled in TRINITY 2.5.0 (Grabherr et al. 2011), ORFs were extracted using TRANSDECODER 5.3.0 (Haas et al. 2013), and translated protein sequences were clustered at 100% similarity using CD-HIT 4.7 (Fu et al. 2012). This non-redundant set of proteins was then used alongside the RNA-Seq alignments to generate AUGUSTUS-based gene predictions using BRAKER 2.1.0 (Hoff et al. 2015) after soft-masking repetitive sequences with REPEATMASKER 4.0.1 (Chen 2004). For comparison, two further sets of annotations were generated from the RNA-Seq alignments only, using BRAKER or STRINGTIE 1.3.4d (Pertea et al. 2015). A final set was generated by merging these three sets to a non-redundant set of loci using GFFCOMPARE 0.10.6 (https://ccb.jhu.edu/software/stringtie/gffcompare.shtml).

### *Tsp*-SL read quantification

The screened and trimmed *Tsp*-SL read pairs were mapped back to the genome using HISAT2, enforcing end-to-end alignment. The mapped R1 reads were extracted using SAMTOOLS and quantified at the gene level using FEATURECOUNTS 1.6.1 (Liao et al. 2013). All read-count analyses were carried out in R 3.5.1 (R Core team 2013).

Since *Tsp*-SL reads are expected to map to the first exon of the *trans*-spliced gene, incorrect gene annotations can be identified via internal exons that receive *Tsp*-SL reads. We generated a non-redundant set of exon annotations across isoforms using BEDTOOLS MERGE 2.26 (Quinlan and Hall 2010) and re-quantified reads at the exon level. These exon counts were used to modify the corresponding gene annotations such that *Tsp*-SL-receiving exons strictly denote the first exon of a gene. The read counts were added across the three libraries and each gene annotation was then split at internal exons that received a distinct peak of at least four *Tsp*-SL counts (and at least 1 count per library) compared to neighbouring exons.

### Multivariate exploration of genome-wide *trans*-splicing events

The gene-by-*Tsp*-SL read-count matrices were visualised and explored with multivariate methods using the ADEGENET 2.1.1 (Jombart 2008) and DESEQ2 1.22.1 (Love et al. 2014) R packages. The counts were transformed into a presence/absence matrix, where a gene is present if it received at least one read. Jaccard distances among *Tsp*-SLs were estimated from this matrix, projected onto two-dimensional space using metric multidimensional scaling (MDS) and hierarchically clustered using Ward’s criterion. The *Tsp*-SL samples were then assigned to two groups, comprising SL2-type spliced leaders (*Tsp*-SL2, *Tsp*-SL10 and *Tsp*-SL12) versus all other *Tsp*-SLs, and linear discriminant analysis of PCA (DAPC) was carried out to identify the relative contribution of each gene to the multidimensional separation between these two groups (Jombart 2008). The best number of principal components to retain was identified using the cross-validation function within DAPC. The same analyses were also carried out on variance-stabilized normalized read counts using PCA instead of MDS (Love et al. 2014).

### Identification of conserved operons

From the *C. elegans* annotation, WS247 gff3 file hosted by WormBase (available at ftp://ftp.wormbase.org/pub/wormbase/species/c_elegans/gff/), a list of operonic genes was derived by extracting all lines designated as operon in the feature field. The gene IDs were isolated from the last field of the operon gff entries, and the protein fasta data for these genes was extracted from the *C. elegans* WS247 protein fasta file (ftp://ftp.wormbase.org/pub/wormbase/species/c_elegans/sequence/protein/) using SAMtools faidx. BLASTP searches using BLAST v2.2.7 were carried using each of these fasta sequences against a custom BLAST database containing all known *T. spiralis* proteins created using the WS247 *T. spiralis* protein file (available at ftp://ftp.wormbase.org/pub/wormbase/species/t_spiralis/sequence/protein/). An E value cutoff of 10^−5^ was used to establish orthology and the best match (if any) for each gene was then checked against corresponding matches to other constituents in the *C. elegans* operon.

Operon conservation was established if the *T. spiralis* orthologues were on the same scaffold, the same strand, within 5kbp of each other, and in the same orientation as the *C. elegans* operon genes. If the BLAST search for two different *C. elegans* operon constituents returned the same *T. spiralis* gene, but the homology matches were to different parts of the gene and in the correct orientation, this was also annotated as a conserved operon. In this case, this was likely an operon that had been incorrectly annotated as a single gene, as we have previously observed (Pettitt et al. 2014).

### Operon prediction

Operons were predicted via identification of uninterrupted runs of at least one strictly SL2-type spliced leader-receiving gene (*Tsp*-SL2, *Tsp*-SL10 or *Tsp*-SL12; no other *Tsp*-SLs) along each contig and strand of the reference genome. Read counts for each *Tsp*-SL were summarised across the three replicate libraries using the geometric mean while allowing for zero counts in one library. These mean counts were then added among all SL1- and SL2-types, and an SL1:SL2-type ratio was computed for each gene. All SL2-type-only genes were designated as “downstream operonic” genes, and the immediately adjacent gene upstream was designated “upstream operonic”. All other *trans*-spliced genes were designated as “monocistronic”.

### Functional enrichment of operonic genes

We carried out basic functional annotation of operonic genes to examine whether these genes may be enriched for particular biological functions. In *C. elegans*, it has been shown that 38% of genes expressed in the germline are operonic, whereas only 15% of all genes in the genome are located in operons (Reinke and Cutter 2009). Accordingly, we first tested the hypothesis whether *T. spiralis* genes involved in germline processes are more frequently located in operons than expected from the genomic background rate of operonic organisation. Identifiers of genes expressed in the *C. elegans* germline were extracted from Reinke et al. (Reinke et al. 2004) and translated to *T. spiralis* orthologues using WORMBASE PARASITE BIOMART (WBPS12; WS267). The genomic locations of the *Tsp* orthologues were then tested for overlap with the locations of all predicted operons using BEDTOOLS INTERSECT 2.26 (Quinlan and Hall 2010). The proportion of orthologues overlapping operons was tested for deviation from the expected overall proportion of operonic genes using binomial and *Chi*-square tests carried out in R.

Additionally, we carried out GeneOntology annotation and enrichment tests for operonic genes. Transcript nucleotide sequences based on each of the five sets of annotations were extracted from the reference genome using GFFREAD 0.9.9 (Trapnell et al. 2010). The sequences were queried against the UNIPROT *swissprot* and *nematoda protein* databases using BLASTX 2.6.0 (Camacho et al. 2009) and retaining up to 10 hits with an E-value cutoff of 1e-3. GeneOntology terms associated with matching UNIPROT accessions were retrieved from the Gene Ontology Annotation (GOA) database (Huntley et al. 2014). Enrichment tests were carried out with the R package GOFUNCR 1.2.0 (Grote 2018), comparing the *biological process* ontology annotations of all operonic genes against the background annotations of the whole genome using the hypergeometric test and retaining significantly enriched annotations at an FDR-corrected P-value threshold of q ≤ 0.05. Significantly enriched annotations were pooled across all datasets and semantically clustered at a similarity threshold of 0.4 (SimRel measure) using REVIGO (Supek et al. 2011).

## DATA ACCESS

RNA-Seq reads for the three *T. spiralis* libraries are available from the NCBI SRA database, accession numbers SRR8327925-SRR8327927 (bioproject PRJNA510020). All scripts used for these analyses are available at https://github.com/glewgun/Wenzel/ or upon request.

## ACKNOWLEDGEMENTS

The work was supported by the Biotechnology and Biological Sciences Research Council [Project grant BB/J007137/1]. CJ PhD studentship was funded by EASTBIO Biotechnology and Biological Sciences Research Council grant number BB/M010996/1. The authors would like to acknowledge the support of the Maxwell computer cluster funded by the University of Aberdeen.

Author contributions: BC provided the biological material; JP, BC and BM acquired funding for the project; BM, JP and BC conceived the research, and managed and coordinated the research activity; MW and CJ designed and implemented the computational analysis; MW, BM, JP and BC analysed the data; MW and JP prepared the figures and tables. JP, MW, BC and BM wrote the manuscript.

