## Supplemental Table S5 for "Deep evolutionary origin of nematode SL2 *trans*-splicing revealed by genome-wide analysis of the *Trichinella spiralis* transcriptome"

**Supplementary Table 5. GeneOntology annotation and enrichment of operonic genes.** Numbers of genes, proteins and associated GeneOntology IDs are presented for five sets of gene annotations (reference annotation + four *de novo* annotations) as well as the corresponding sets of operonic genes. Enrichment-test results are presented for all enriched GO terms in the biological process ontology across all datasets. Results of semantic clustering of enriched GO terms are presented in tabular and graphical format.

**GO Annotation Statistics**

| **Background GO annotations** | | | | | |
|---|---|---|---|---|---|
| **Annotation** | | **Genes** | **Proteins** | **GOIDs** | |
| REFERENCE | | 12,376 | 64,722 | 3,210 | |
| BRAKER | | 12,436 | 66,479 | 2,922 | |
| BRAKER+TRINITY | | 14,444 | 69,177 | 2,989 | |
| STRINGTIE | | 6,758 | 52,055 | 2,690 | |
| DENOVO_ALL | | 6,901 | 55,368 | 2,699 | |
| **GO Annotations of operonic genes** | | | | | |
| **Genome annotations** | **TSL screening stringency** | **Genes** | **Proteins** | **GO IDs** | **Enriched GO IDs (BP)** |
| REFERENCE | 10 bp | 491 | 452 | 742 | 7 |
| REFERENCE | 8 bp | 507 | 464 | 738 | 4 |
| BRAKER | 10 bp | 622 | 585 | 775 | 79 |
| BRAKER | 8 bp | 630 | 590 | 756 | 71 |
| BRAKER+TRINITY | 10 bp | 637 | 599 | 750 | 110 |
| BRAKER+TRINITY | 8 bp | 650 | 607 | 741 | 105 |
| STRINGTIE | 10 bp | 421 | 358 | 538 | - |
| STRINGTIE | 8 bp | 407 | 345 | 505 | - |
| DENOVO_ALL | 10 bp | 439 | 319 | 515 | 2 |
| DENOVO_ALL | 8 bp | 433 | 316 | 491 | 1 |

**GO BP Enrichment**

| ontology | node_id | node_name | raw_p_overrep | qval |
|---|---|---|---|---|
| biological_process | GO:0006396 | RNA processing | 1.31E-11 | 3.32E-08 |
| biological_process | GO:0016070 | RNA metabolic process | 1.89E-11 | 3.32E-08 |
| biological_process | GO:0034660 | ncRNA metabolic process | 1.16E-11 | 3.32E-08 |
| biological_process | GO:0010467 | gene expression | 1.30E-10 | 1.25E-07 |
| biological_process | GO:0051641 | cellular localization | 4.05E-10 | 3.57E-07 |
| biological_process | GO:0006091 | generation of precursor metabolites and energy | 2.34E-10 | 8.00E-07 |
| biological_process | GO:0034470 | ncRNA processing | 1.53E-09 | 1.07E-06 |
| biological_process | GO:0046907 | intracellular transport | 2.95E-09 | 1.73E-06 |
| biological_process | GO:0051649 | establishment of localization in cell | 4.73E-09 | 2.38E-06 |
| biological_process | GO:0022900 | electron transport chain | 2.83E-09 | 2.43E-06 |
| biological_process | GO:0055114 | oxidation-reduction process | 2.34E-09 | 2.43E-06 |
| biological_process | GO:0033036 | macromolecule localization | 8.32E-09 | 2.93E-06 |
| biological_process | GO:0006399 | tRNA metabolic process | 1.84E-08 | 5.88E-06 |
| biological_process | GO:0009451 | RNA modification | 1.11E-07 | 2.79E-05 |
| biological_process | GO:0009056 | catabolic process | 2.16E-07 | 4.87E-05 |
| biological_process | GO:1901575 | organic substance catabolic process | 2.22E-07 | 4.87E-05 |
| biological_process | GO:0044248 | cellular catabolic process | 2.92E-07 | 6.05E-05 |
| biological_process | GO:0016071 | mRNA metabolic process | 5.53E-07 | 0.000107802 |
| biological_process | GO:0071702 | organic substance transport | 5.82E-07 | 0.000107802 |
| biological_process | GO:0006886 | intracellular protein transport | 1.45E-06 | 0.000254937 |
| biological_process | GO:0002943 | tRNA dihydrouridine synthesis | 1.87E-06 | 0.000312983 |
| biological_process | GO:0034613 | cellular protein localization | 2.84E-06 | 0.000435096 |
| biological_process | GO:0070727 | cellular macromolecule localization | 2.84E-06 | 0.000435096 |
| biological_process | GO:0008033 | tRNA processing | 3.38E-06 | 0.000495749 |
| biological_process | GO:0044281 | small molecule metabolic process | 4.51E-06 | 0.000634542 |
| biological_process | GO:0051179 | localization | 5.23E-06 | 0.000707152 |
| biological_process | GO:0015031 | protein transport | 6.77E-06 | 0.000881698 |

| | | | | |
|---|---|---|---|---|
| biological_process | GO:0008104 | protein localization | 7.41E-06 | 0.000931559 |
| biological_process | GO:0015833 | peptide transport | 8.57E-06 | 0.000949683 |
| biological_process | GO:0032259 | methylation | 8.86E-06 | 0.000949683 |
| biological_process | GO:0051234 | establishment of localization | 8.91E-06 | 0.000949683 |
| biological_process | GO:0051643 | endoplasmic reticulum localization | 8.76E-06 | 0.000949683 |
| biological_process | GO:0061817 | endoplasmic reticulum-plasma membrane tethering | 8.76E-06 | 0.000949683 |
| biological_process | GO:0042886 | amide transport | 9.26E-06 | 0.000958303 |
| biological_process | GO:0045184 | establishment of protein localization | 1.08E-05 | 0.001085884 |
| biological_process | GO:0000956 | nuclear-transcribed mRNA catabolic process | 1.50E-05 | 0.001462955 |
| biological_process | GO:0016999 | antibiotic metabolic process | 1.33E-05 | 0.001463915 |
| biological_process | GO:0006810 | transport | 1.77E-05 | 0.001686887 |
| biological_process | GO:1901565 | organonitrogen compound catabolic process | 1.92E-05 | 0.001779566 |
| biological_process | GO:0009057 | macromolecule catabolic process | 2.10E-05 | 0.001896965 |
| biological_process | GO:0006457 | protein folding | 2.75E-05 | 0.002412207 |
| biological_process | GO:0043414 | macromolecule methylation | 2.81E-05 | 0.002412207 |
| biological_process | GO:0006379 | mRNA cleavage | 3.16E-05 | 0.002647787 |
| biological_process | GO:0043603 | cellular amide metabolic process | 2.77E-05 | 0.002672987 |
| biological_process | GO:0071705 | nitrogen compound transport | 3.78E-05 | 0.003095242 |
| biological_process | GO:0006733 | oxidoreduction coenzyme metabolic process | 3.89E-05 | 0.00311268 |
| biological_process | GO:0044267 | cellular protein metabolic process | 3.40E-05 | 0.003151043 |
| biological_process | GO:0006518 | peptide metabolic process | 5.15E-05 | 0.003913754 |
| biological_process | GO:0043436 | oxoacid metabolic process | 6.14E-05 | 0.004499583 |
| biological_process | GO:0044265 | cellular macromolecule catabolic process | 6.36E-05 | 0.004562816 |
| biological_process | GO:0006082 | organic acid metabolic process | 6.58E-05 | 0.004627049 |
| biological_process | GO:0006367 | transcription initiation from RNA polymerase II promoter | 7.85E-05 | 0.005418087 |
| biological_process | GO:0006397 | mRNA processing | 7.01E-05 | 0.005476989 |
| biological_process | GO:0019752 | carboxylic acid metabolic process | 0.000103674 | 0.007013956 |
| biological_process | GO:0009435 | NAD biosynthetic process | 0.000120321 | 0.007986598 |
| biological_process | GO:0000288 | nuclear-transcribed mRNA catabolic process, deadenylation-dependent decay | 8.18E-05 | 0.009040815 |

| | | | | |
|---|---|---|---|---|
| biological_process | GO:0070647 | protein modification by small protein conjugation or removal | 8.64E-05 | 0.009254933 |
| biological_process | GO:0006869 | lipid transport | 0.000148217 | 0.009656067 |
| biological_process | GO:0043043 | peptide biosynthetic process | 0.000152053 | 0.009725879 |
| biological_process | GO:0043604 | amide biosynthetic process | 0.000146778 | 0.009930095 |
| biological_process | GO:0006480 | N-terminal protein amino acid methylation | 0.000161553 | 0.01014896 |
| biological_process | GO:0019362 | pyridine nucleotide metabolic process | 0.000177415 | 0.010761149 |
| biological_process | GO:0046496 | nicotinamide nucleotide metabolic process | 0.000177415 | 0.010761149 |
| biological_process | GO:0046700 | heterocycle catabolic process | 0.000189258 | 0.011284892 |
| biological_process | GO:0010876 | lipid localization | 0.000208955 | 0.012059003 |
| biological_process | GO:1901361 | organic cyclic compound catabolic process | 0.000209096 | 0.012059003 |
| biological_process | GO:0090305 | nucleic acid phosphodiester bond hydrolysis | 0.000141895 | 0.013889514 |
| biological_process | GO:0019359 | nicotinamide nucleotide biosynthetic process | 0.000179571 | 0.01471119 |
| biological_process | GO:0019363 | pyridine nucleotide biosynthetic process | 0.000179571 | 0.01471119 |
| biological_process | GO:0070507 | regulation of microtubule cytoskeleton organization | 0.000155487 | 0.01471119 |
| biological_process | GO:0008299 | isoprenoid biosynthetic process | 0.000270485 | 0.015104243 |
| biological_process | GO:0001510 | RNA methylation | 0.000303763 | 0.016697474 |
| biological_process | GO:0051188 | cofactor biosynthetic process | 0.000256805 | 0.017072413 |
| biological_process | GO:0016567 | protein ubiquitination | 0.000283465 | 0.017984284 |
| biological_process | GO:0006402 | mRNA catabolic process | 0.000359769 | 0.018890552 |
| biological_process | GO:0072524 | pyridine-containing compound metabolic process | 0.000359769 | 0.018890552 |
| biological_process | GO:0006720 | isoprenoid metabolic process | 0.000377266 | 0.019235108 |
| biological_process | GO:0006412 | translation | 0.000383278 | 0.019262448 |
| biological_process | GO:0051493 | regulation of cytoskeleton organization | 0.000339682 | 0.019724579 |
| biological_process | GO:0051604 | protein maturation | 0.000330149 | 0.019724579 |
| biological_process | GO:0016072 | rRNA metabolic process | 0.000271374 | 0.019781458 |
| biological_process | GO:0019637 | organophosphate metabolic process | 0.000379346 | 0.021305552 |
| biological_process | GO:0006364 | rRNA processing | 0.000408151 | 0.0217557 |
| biological_process | GO:0098661 | inorganic anion transmembrane transport | 0.00047196 | 0.021846773 |
| biological_process | GO:0034655 | nucleobase-containing compound catabolic process | 0.000480326 | 0.021938014 |
| biological_process | GO:0006418 | tRNA aminoacylation for protein translation | 0.000510566 | 0.02205729 |

| | | | | |
|---|---|---|---|---|
| biological_process | GO:0008272 | sulfate transport | 0.000514127 | 0.02205729 |
| biological_process | GO:0051186 | cofactor metabolic process | 0.000497018 | 0.02205729 |
| biological_process | GO:1902358 | sulfate transmembrane transport | 0.000514127 | 0.02205729 |
| biological_process | GO:0000290 | deadenylation-dependent decapping of nuclear-transcribed mRNA | 0.000400678 | 0.022140676 |
| biological_process | GO:0010038 | response to metal ion | 0.00061992 | 0.025817921 |
| biological_process | GO:0043412 | macromolecule modification | 0.000623799 | 0.025817921 |
| biological_process | GO:0043038 | amino acid activation | 0.000645555 | 0.026104159 |
| biological_process | GO:0043039 | tRNA aminoacylation | 0.000645555 | 0.026104159 |
| biological_process | GO:0072525 | pyridine-containing compound biosynthetic process | 0.000613634 | 0.031077823 |
| biological_process | GO:1901566 | organonitrogen compound biosynthetic process | 0.000797127 | 0.031866977 |
| biological_process | GO:0032886 | regulation of microtubule-based process | 0.000681902 | 0.033374219 |
| biological_process | GO:0031167 | rRNA methylation | 0.000885641 | 0.034618726 |
| biological_process | GO:0072348 | sulfur compound transport | 0.000885641 | 0.034618726 |
| biological_process | GO:0010243 | response to organonitrogen compound | 0.000901211 | 0.034660125 |
| biological_process | GO:0045333 | cellular respiration | 0.000906405 | 0.034660125 |
| biological_process | GO:0006400 | tRNA modification | 0.000978953 | 0.037031786 |
| biological_process | GO:0071840 | cellular component organization or biogenesis | 0.001028589 | 0.040658169 |
| biological_process | GO:0044270 | cellular nitrogen compound catabolic process | 0.001108735 | 0.041058206 |
| biological_process | GO:0015980 | energy derivation by oxidation of organic compounds | 0.000941203 | 0.041877432 |
| biological_process | GO:0016192 | vesicle-mediated transport | 0.001175052 | 0.042616837 |
| biological_process | GO:0019439 | aromatic compound catabolic process | 0.001317397 | 0.046814164 |
| biological_process | GO:0030163 | protein catabolic process | 0.00133153 | 0.04684321 |
| biological_process | GO:0006401 | RNA catabolic process | 0.001486239 | 0.047554437 |
| biological_process | GO:0006465 | signal peptide processing | 0.001412727 | 0.047554437 |
| biological_process | GO:0008285 | negative regulation of cell proliferation | 0.001486921 | 0.047554437 |
| biological_process | GO:0010257 | NADH dehydrogenase complex assembly | 0.001486921 | 0.047554437 |
| biological_process | GO:0030433 | ubiquitin-dependent ERAD pathway | 0.001486921 | 0.047554437 |
| biological_process | GO:0032981 | mitochondrial respiratory chain complex I assembly | 0.001486921 | 0.047554437 |
| biological_process | GO:0034315 | regulation of Arp2/3 complex-mediated actin nucleation | 0.001486921 | 0.047554437 |

| | | | | |
|---|---|---|---|---|
| biological_process | GO:0051125 | regulation of actin nucleation | 0.001486921 | 0.047554437 |
| biological_process | GO:0042254 | ribosome biogenesis | 0.001324482 | 0.048536742 |

### REVIGO SimRel0.4

| term_ID | description | frequency | plot_X | plot_Y | plot_size | log10 p-value | uniqueness | dispensability | representative | eliminated |
|---|---|---|---|---|---|---|---|---|---|---|
| GO:0034660 | ncRNA metabolic process | 3.41% | 4.812 | 4.749 | 5.641 | -10.9341 | 0.717 | 0 | 34660 | 0 |
| GO:0010467 | gene expression | 19.67% | 2.708 | 7.542 | 6.402 | -9.8857 | 0.837 | 0.225 | 10467 | 0 |
| GO:0006091 | generation of precursor metabolites and energy | 1.94% | -1.549 | 6.325 | 5.396 | -9.6315 | 0.899 | 0.081 | 6091 | 0 |
| GO:0051641 | cellular localization | 2.04% | -5.489 | 3.301 | 5.418 | -9.3921 | 0.854 | 0.342 | 51641 | 0 |
| GO:0055114 | oxidation-reduction process | 15.06% | 5.231 | -3.841 | 6.286 | -8.6312 | 0.872 | 0.222 | 55114 | 0 |
| GO:0022900 | electron transport chain | 0.56% | 4.485 | -4.729 | 4.86 | -8.5475 | 0.836 | 0.07 | 22900 | 0 |
| GO:0046907 | intracellular transport | 1.56% | -5.909 | 3.469 | 5.302 | -8.5297 | 0.809 | 0 | 46907 | 0 |
| GO:0033036 | macromolecule localization | 3.03% | -5.715 | 4.451 | 5.59 | -8.0801 | 0.849 | 0.372 | 33036 | 0 |
| GO:0009056 | catabolic process | 4.82% | -6.592 | -1.802 | 5.791 | -6.6655 | 0.958 | 0.022 | 9056 | 0 |
| GO:1901575 | organic substance catabolic process | 4.61% | -2.772 | -5.431 | 5.772 | -6.6546 | 0.751 | 0.069 | 1901575 | 0 |
| GO:0016071 | mRNA metabolic process | 0.80% | 5.213 | 5.426 | 5.01 | -6.2572 | 0.755 | 0.376 | 16071 | 0 |
| GO:0051179 | localization | 18.50% | 0.285 | -1.98 | 6.375 | -5.2818 | 0.986 | 0 | 51179 | 0 |
| GO:0015031 | protein transport | 2.25% | -5.344 | 4.994 | 5.461 | -5.1696 | 0.79 | 0.377 | 15031 | 0 |
| GO:0032259 | methylation | 3.10% | -4.902 | -4.922 | 5.6 | -5.0528 | 0.96 | 0.02 | 32259 | 0 |
| GO:0016999 | antibiotic metabolic process | 0.06% | -1.932 | 1.176 | 3.846 | -4.8756 | 0.923 | 0.055 | 16999 | 0 |
| GO:0006457 | protein folding | 0.90% | -2.943 | -1.343 | 5.064 | -4.56 | 0.94 | 0.04 | 6457 | 0 |
| GO:0043603 | cellular amide metabolic process | 6.88% | 6.889 | 2.234 | 5.946 | -4.5582 | 0.808 | 0.234 | 43603 | 0 |
| GO:0044267 | cellular protein metabolic process | 14.29% | 3.027 | 6.09 | 6.263 | -4.4681 | 0.768 | 0.354 | 44267 | 0 |
| GO:0071705 | nitrogen compound transport | 1.77% | -6.07 | 3.841 | 5.355 | -4.4221 | 0.854 | 0.349 | 71705 | 0 |
| GO:0006733 | oxidoreduction coenzyme metabolic process | 1.27% | 0.952 | -4.052 | 5.213 | -4.4097 | 0.84 | 0.077 | 6733 | 0 |
| GO:0006367 | transcription initiation from RNA polymerase II promoter | 0.11% | 5.672 | 4.001 | 4.144 | -4.1049 | 0.774 | 0.302 | 6367 | 0 |
| GO:0090305 | nucleic acid phosphodiester bond hydrolysis | 2.27% | 5.431 | 4.809 | 5.464 | -3.848 | 0.756 | 0.338 | 90305 | 0 |
| GO:0006480 | N-terminal protein amino acid methylation | 0.01% | 1.267 | 5.067 | 3.103 | -3.7917 | 0.839 | 0.112 | 6480 | 0 |
| GO:0008299 | isoprenoid biosynthetic process | 0.44% | 5.726 | -3.59 | 4.754 | -3.5679 | 0.823 | 0.176 | 8299 | 0 |

| | | | | | | | | | |
|---|---|---|---|---|---|---|---|---|---|
| GO:0051604 | protein maturation | 0.29% | 1.389 | 6.791 | 4.575 | -3.4813 | 0.851 | 0.183 | 51604 | 0 |
| GO:0051493 | regulation of cytoskeleton organization | 0.23% | -0.658 | -6.429 | 4.474 | -3.4689 | 0.874 | 0.034 | 51493 | 0 |
| GO:0072524 | pyridine-containing compound metabolic process | 1.35% | 6.717 | 1.089 | 5.239 | -3.444 | 0.779 | 0.222 | 72524 | 0 |
| GO:0019637 | organophosphate metabolic process | 6.15% | 1.228 | 1.451 | 5.897 | -3.421 | 0.85 | 0.099 | 19637 | 0 |
| GO:0098661 | inorganic anion transmembrane transport | 0.47% | -4.909 | 4.944 | 4.78 | -3.3261 | 0.852 | 0.301 | 98661 | 0 |
| GO:0051186 | cofactor metabolic process | 3.99% | 3.268 | -1.315 | 5.709 | -3.3036 | 0.891 | 0.09 | 51186 | 0 |
| GO:0010038 | response to metal ion | 0.13% | -3.178 | -3.503 | 4.229 | -3.2077 | 0.966 | 0 | 10038 | 0 |
| GO:0043412 | macromolecule modification | 9.79% | 1.605 | 8.073 | 6.099 | -3.205 | 0.851 | 0.285 | 43412 | 0 |
| GO:0043038 | amino acid activation | 1.12% | 6.167 | -1.282 | 5.159 | -3.1901 | 0.744 | 0.321 | 43038 | 0 |
| GO:0032886 | regulation of microtubule-based process | 0.07% | 2.165 | -5.861 | 3.934 | -3.1663 | 0.885 | 0.203 | 32886 | 0 |
| GO:0072348 | sulfur compound transport | 0.25% | -6.211 | 2.518 | 4.504 | -3.0527 | 0.838 | 0.282 | 72348 | 0 |
| GO:0071840 | cellular component organization or biogenesis | 8.57% | -5.992 | -3.342 | 6.041 | -2.9878 | 0.984 | 0 | 71840 | 0 |
| GO:0016192 | vesicle-mediated transport | 1.09% | -5.154 | 4.097 | 5.144 | -2.9299 | 0.86 | 0.33 | 16192 | 0 |
| GO:0006465 | signal peptide processing | 0.05% | 5.078 | 2.138 | 3.809 | -2.8499 | 0.783 | 0.237 | 6465 | 0 |
| GO:0008285 | negative regulation of cell proliferation | 0.13% | 0.273 | -6.433 | 4.214 | -2.8277 | 0.914 | 0.213 | 8285 | 0 |

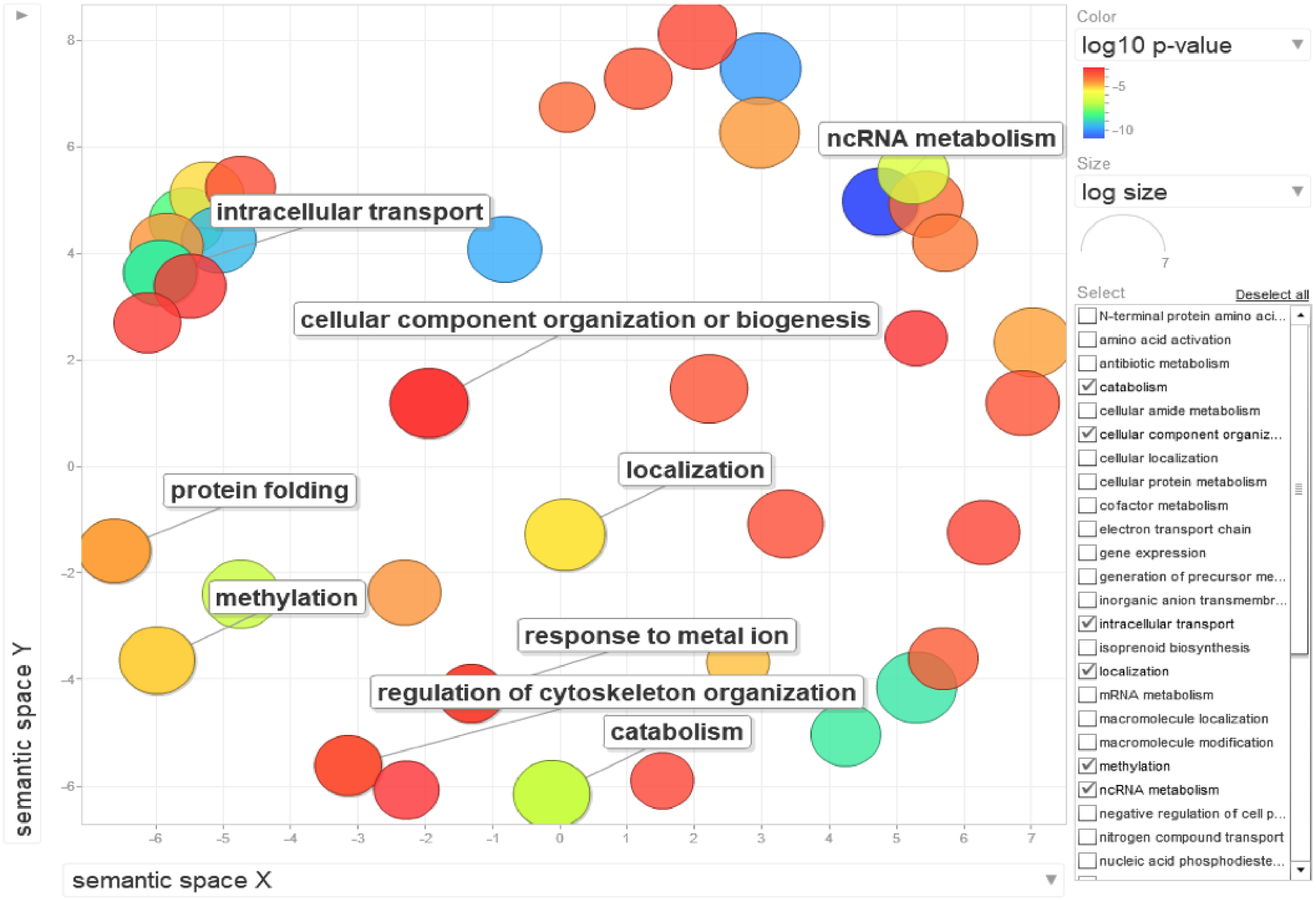

intracellular transport
ncRNA metabolism
cellular component organization or biogenesis
localization
protein folding
methylation
response to metal ion
regulation of cytoskeleton organization
catabolism
semantic space Y
semantic space X
Color
log10 p-value
-5
-10
Size
log size
7
Select
Deselect all
N-terminal protein amino aci...
amino acid activation
antibiotic metabolism
catabolism
cellular amide metabolism
cellular component organiz...
cellular localization
cellular protein metabolism
cofactor metabolism
electron transport chain
gene expression
generation of precursor me...
inorganic anion transmembr...
intracellular transport
isoprenoid biosynthesis
localization
mRNA metabolism
macromolecule localization
macromolecule modification
methylation
ncRNA metabolism
negative regulation of cell p...
nitrogen compound transport
nucleic acid phosphodieste...
