## Supplemental Table S4 for "Deep evolutionary origin of nematode SL2 *trans*-splicing revealed by genome-wide analysis of the *Trichinella spiralis* transcriptome"

**Supplementary Table 4. Identification of operonic germline genes in T. spiralis via orthology to *C. elegans*.** The Wormbase IDs of 4,235 *C. elegans* germline genes are presented alongside genomic information of 1,332 orthologous *T. spiralis* genes from ParaSite BioMart output. Numbers of intersecting *T. spiralis* operonic genes as predicted from our 20 datasets are presented with enrichment test results (*Chi*-square and binomial tests).

*C. elegans* Germline Genes
C05B5.2, F37A8.2, F40F12.3, F54A3.4, F58D5.8, K03H1.1, M70.3b, T05A7.6, T16G12.7, T23F6.3, T25C8.3, Y116A8A.6, Y23H5B.9, Y39G10AR.16, Y43F8C.5, Y46C8AL.1, Y46G5A.30, Y47D3A.22, Y50E8A.12, Y50E8A.2, Y51A2B.6a, Y52D5A.2, Y59E9AL.6, Y59E9AR.7, Y65B4A.9, Y71G12B.27, Y73B6A.1, Y73F8A.12, Y73F8A.14, C40H5.1, H12D21.1, W06A7.5, ZC412.6, ZC412.7, C25A8.1, C33F10.10, F21D9.3, W01B6.4, F36H12.13, K07F5.5, ZK484.8, C24D10.7, C24D10.8, F11G11.8, T23B7.1, C47A4.5, Y45F10B.3, C32E8.4, F09C12.7, ZK1248.4, ZK84.5, K05F1.8, ZK1248.17, F32B5.4, ZK484.5, C03B8.1, F46F5.9, ZC168.6, ZK1225.6, C35D10.11, C55C2.2, E03H12.10, F44D12.7, T27A3.3, T28H11.6, F02C9.1, C02F5.2, F36H12.4, E03A3.4, C04G2.4, C09B9.6, C33F10.9, C34F11.4, C34F11.6, F26G1.7, F32B6.6, F36H12.7, F58A6.8, K05F1.2, K05F1.7, K07F5.2, K07F5.3, R05F9.13, R05F9.8, R13H9.2, R13H9.4, T13F2.10, T13F2.11, ZK1248.6, ZK1251.6, ZK354.1, ZK354.11, ZK354.4, ZK354.5, ZK546.6, F08H9.2, C27D6.3, F35H8.4, W02D9.5, Y106G6H.13, Y37D8A.8, F35H8.1, Y67A6A.1, C18A3.7, C34D4.3, ZC412.9, C34F11.2, K03H1.12, C34G6.3, K07F5.7, F36A4.3, Y46G5A.14, K01D12.15, ZC116.2, T14D7.3, Y51B9A.5, R05D3.5, ZC262.1, C09F9.1, F29D10.1, F36A2.10, C48E7.7, T08B2.12, ZK1251.1, C25D7.1, C24D10.2, ZK617.3, F42E8.2, F52F12.5, F11G11.4, F36A4.2, F08G2.6, Y116A8A.7, T22B3.3, Y73F8A.15, C07G1.6, F36A4.4, H09I01.1, R05F9.3, T13A10.1, W09D6.4, ZK1307.3, F44G4.5, C15A11.2, D2062.4, ZK930.7, F47B8.11, C38C3.8, T15H9.5, C27F2.6, F46A8.9, T13F2.12, F32B6.7, K06A5.3, F36D1.4, T14G10.8, ZK353.3, T01H3.5, C02F5.5, Y49E10.7, C26E6.1, C05C12.5, C10G11.9, T27A3.4, F37A8.1, C26C6.6, C18H7.7, F17E9.5, F26F4.2, K01D12.7, F43E2.10, Y106G6A.4, F36H12.5, F41G3.4, C25A8.2, ZK1307.4, F02C9.4, F10D11.3, F46B3.4, Y106G6E.2, F58F12.2, C33A12.15, B0432.12, E03H12.5, B0432.11, T23B3.5, F29B9.7, T20F5.5, Y38F1A.7, C41G7.6, C01G12.3, W09C3.7, E03H12.7, F28H7.4, ZK930.4, C26B2.2, ZK945.6, K08F4.5, F56F4.7, ZK1251.3, C10H11.7, B0496.6, F33D11.1, T02E1.6, T10E9.6, Y43C5B.2, Y47G6A.3, ZC477.7, R02D5.7, F49E12.4, ZC477.1, ZK546.3, B0379.2, ZK637.12, F42A9.3, F32B4.2, R13H9.1, B0041.1, T28H11.7, Y106G6E.3, F27C1.1, F47B3.2, F46A9.1, R01H2.4, R07E5.15, T21G5.4, ZK783.3, C25G4.6, T13F2.9, C06A5.2, F32B6.5, Y38F1A.1, ZK354.7, Y45F10B.8, ZK637.15, ZK520.5, F07D3.3, C04F12.7, C55B7.10, K07F5.11, B0273.1, C04G2.5, F41G3.5, F28E10.4, F54C8.1, Y57A10B.7, C17F3.3, C29F5.3, C54G4.3, Y116A8C.37, C16A11.7, T06E4.5, F49F1.12, AH6.3, K05F1.9, F58E6.5, B0491.3, T02H6.4, F36D3.4, K07F5.8, C47E8.1, K01H12.2, T03F1.5, C17D12.5, H12D21.2, T23F11.2, R09E10.6, W03C9.1, C35D10.3, T01B11.4, F54C1.8, ZK795.2, F47B3.5, F40F9.3, K11H3.2, W09C3.6, ZC317.6, W03F11.3, T08G11.2, F59C6.2, F11G11.9, F20D6.6, F55C5.2, F21H7.5, T27E7.1, F21F3.3, W01C9.4, F38H4.6, C45G9.9, F54D1.1, C45G9.4, T08H10.4, C04G2.9, C09H5.7, C43G2.3, T05C12.1, Y1A5A.1, ZK546.7, C47E12.10, W06D4.2, F56H11.3, C08F11.10, ZK1248.5, K07A12.5, C35E7.9, F47B3.4, F36H12.8, R13H9.5, Y59E9AL.1, Y43F8C.9, ZK354.3, F44D12.4, Y81G3A.1, C35D10.2, K06A5.2, R102.3, R10E4.7, C50D2.3, C25D7.12, C01G10.14, T16A9.3, F38A5.6, F55D12.6, C17C3.11, C14C10.1, K07A1.4, C28D4.7, C28D4.8, Y69E1A.2, F53B6.7, ZK938.1, C34D4.2, F52H3.6, T28H11.1, F46A9.2, C17H12.3, C47A4.3, C33H5.16, D1081.5, W02A2.8, ZC412.5, F36H12.9, R13H9.6, ZK858.2, F13G11.2, C06A8.6, F26A3.5, AH6.2, R10E9.2, ZK512.8, T05C12.3, F36H12.3, C33F10.12, F40H6.1, T05F1.8, F42G4.6, F44F4.10, B0511.4, Y69E1A.4, Y106G6G.1, C02B10.6, F36H12.10, T16A9.5, T27A3.5, F53B6.4, C47E12.11, Y53F4B.36, F22B5.5, K02F6.2, C15H7.3, ZK84.2, Y69E1A.8, F28A10.1, F36A2.11, F18C5.4, T28H11.5, F53C3.1, F17A9.1, F10G8.1, C55B7.3, T04B2.2, C39H7.1, B0261.6, C56A3.8, F35C11.3, M05D6.1, Y38H8A.3, ZK265.3, F36H12.14, T28C6.5, ZK673.6, W03D8.10, ZK484.7, C09H10.9, R08C7.8, D2045.5, K08D10.7, B0379.7, F02E9.3, C52E4.7, B0218.7, K06H7.8, C27D8.1, C30B5.3, F58A6.5, C38C10.3, ZK930.6, K11C4.1, F44D12.6, K08C9.2, M02B1.4, F23B12.1, C15H7.4, T23B3.3, Y6E2A.9, Y106G6G.4, Y69E1A.1, M05B5.1, C29E6.3, ZK1098.6, F32H2.7, T25B9.2, C08F8.6, Y37D8A.5, T20H4.2, F48C1.7, ZK354.6, F23B2.7, C43E11.5, C50F2.5, F42C5.5, M05B5.3, C49C3.2, F47B3.1, C25A6.1, C56C10.6, ZK507.1, W09C3.8, C30A5.4, F57B9.8, C14C11.1, K08C9.1, M176.9, T01C3.5, K05F1.3, F56F3.4, K01H12.4, T28C12.3, T09B4.7, T08B2.11, C46E10.1, F13A7.1, R09E10.1, C04G2.2, T05F1.5, Y38H8A.4, F58H1.6, T09F5.10, F21F3.2, C36B1.10, C17H12.5, ZK622.1, C36H8.1, W01B6.2, T13A10.11, F26B1.5, C09B9.4, Y18D10A.21, F38A5.11, C15C6.2a, C18G1.9, F08G5.2, C32D5.4, ZK354.2, C06A1.3, C05C12.1, C09B9.2, F22D6.9, C24H11.1, F40G12.10, Y44A6D.5, T02B11.2, K08F8.5a, C33F10.8, R09E10.4, W01B6.6, H06H21.9, W02A2.4, Y45F10B.2, C31H1.5, BE10.1, F32A11.3, F54H5.2, F47B3.7, ZK550.5, R148.7, R10D12.10, C25G4.4, ZC581.7, C50F7.3, Y43C5B.3, T02E1.7, F14H3.2, F56A11.6, F56D5.2, C47D12.3, C55C3.4, F25H5.7, B0523.1, F49E11.7, C53A5.4, T28F4.3, B0252.5, M7.7, F21D9.2, W08D2.8, F44D12.8, C15H11.1, C25A8.5, D2024.1, ZC513.10, K08D10.8, ZK593.9, C27D8.2, R01H2.2, Y69A2AR.6, F26A1.3, C32C4.3, F57A8.6, T13H10.1, T09A12.1, F45E4.6, F57F4.1, F37H8.4, C24D10.1, C14A4.13, H08M01.1, C18E9.8, Y116A8C.24, Y116A8C.38, T04A8.3, C34B2.3, C09D4.3, Y106G6D.3, C43F9.6, ZK418.2, ZK354.8, C07E3.4, T06C10.6, W04E12.4, T15B12.2, Y37E11B.10, F56D6.5, F57H12.5, M05D6.3, F35C11.2, F55F8.7, F26F4.8, R11E3.1, ZK1290.6, F53B2.5, Y40H4A.2, ZK1225.4, ZK1127.2, F33D11.2, AH10.1, Y4C6A.1, F36A2.12, K09E4.1, T25B9.5, F10F2.8, C32E12.1, Y69E1A.3, T06C10.3, C01F6.2, F58A6.11, C48B6.4, C49C8.1, R05D7.2, ZK1225.5, F58G1.3, F42G8.9, F47B3.6, C05D2.3, F42G4.2, M162.7, W02B12.12, Y57G7A.5, R10H10.2, C46H11.6, C09G9.4, F47C12.4, C08F8.4, ZC581.2, W01B6.5, T11F8.4, Y57G11A.2, F59B2.5, M04G7.2, F25F2.1, T25B9.6, Y32F6A.2, F47D12.7, F02E11.1, R08C7.5, T08B6.4, ZC513.3, Y70C5C.5, F46A8.10, T21G5.1, C44B9.2, Y53F4B.11, C04E6.5, F27C8.5, C50F4.2, Y57G11B.3, R05H5.2, F10F2.7, K08A2.2, F23C8.7, C18H7.4, K09B11.5, ZC581.6, F27E5.3, F21H7.2, F53G12.6, C01G5.3, F19B6.4, T08G3.4, R09E10.3, F23C8.8, F01G10.5, R13G10.4, F59A3.8, F36H1.3, T26A5.1, C38C3.7, C55C3.3, C53B4.2, K09C6.8, T04F3.3, F49C12.15, F36D3.8, F42G8.8, C46A5.1, T06D4.4, C31H1.1, C17H12.12, C38C3.3, Y73F8A.20, F26B1.1, K09C6.7, R03D7.8, C34F11.5, F18A12.4, F58D2.2, Y49E10.10, F55H12.1, F55A11.6, F07E5.8, F55H12.2, Y57G7A.6, F44F1.3, B0393.4, ZK250.6, M28.9, T22B3.2a, T04A6.3, D1037.5, K08E4.5, K02F6.3, T10E9.4, F32H2.9, R07C3.4, M01A12.1, T22H9.3, C01G10.1, C01G12.8, F09C12.2, T08B6.5, F26D2.10, W03B1.9, F35E2.6, K02E11.1, Y116F11B.7, H05L14.1, Y43F8A.2, Y69A2AR.19, F17C8.3, R102.7, Y39B6A.30, T04A8.13, F54H12.5, C28A5.6, R13A1.3, Y113G7C.1, M70.1, F37A4.7a, C24F3.5, Y53C10A.10, F34D10.2, F47G6.4, Y59E9AL.3, ZK1128.2a, F56A4.9, Y116A8A.2, Y38E10A.17, Y38F2AR.10, Y39H10A.7, Y55B1AR.4, Y75B8A.23, Y32B12A.4, ZC155.2, W02D3.6, F09E8.1, Y50E8A.10, K02A11.2, B0457.4, F54H5.3, F20D6.1, ZK809.1, C41D11.2, Y37E11B.9, F17C8.5, Y52B11A.1, T08H10.3, C35A5.4, T21C12.4, C03C10.2, AC3.4, C06C6.6, T25D10.1, Y49E10.17, C56G2.4, C06C6.9, F36G9.15, F44G3.7, H24O09.2, Y106G6E.1, Y46G5A.22a, Y47D3A.14, Y65B4A.7, F55C12.2, F26A1.10, ZK262.4, W03B1.1, F21F8.5, Y69A2AR.8, C03B8.3, Y106G6G.3, Y57G7A.3, F35H10.2, Y44A6B.4, M70.2, H38K22.4, Y39A1A.18, T05C12.9, F46C5.7, F42A9.4, B0207.9, Y39E4A.1, W01B6.8, F26F12.2, C33C12.4, Y76A2A.1, B0496.1, F25H5.2, C25D7.2, C34B2.4, F10G8.2, B0334.10,

C28C12.11, C50E10.1, F56F4.2, W03D8.3, C28C12.1, C45G9.1, F59A3.7, M03E7.3, C10G11.10, M88.3, Y51B9A.3, F01D4.3, F36D4.1, R13F6.5, F13E9.5, Y38H6C.15, C48E7.9, R05D3.3, F44D12.11, F15H9.1, R13A5.11, F13A7.7, F59C6.6, Y57G11C.14, ZC477.2, C06E1.7, C24H11.2, F59G1.2, Y38H8A.2, F14E5.4, T08B1.5, Y51A2B.4, T25B9.4, Y75B7B.1, Y59E9AR.2, F58F12.3, K10C9.7, F10F2.6, F43G6.6, F48A9.3, H23L24.2, F13D12.9, Y6B3B.7, C54G4.2, C27B7.6, C01B12.4, ZK688.7, W03D8.1, Y18H1A.1, K12B6.2, H12D21.5, Y116A8C.33, F36G9.13, C06C6.7, W03B1.5, Y45F3A.1, F22D6.1, ZK1053.6, F59C6.3, Y116A8C.4, T05E8.1, F14E5.3, ZK418.6, T04B2.4, M28.2, Y113G7A.10, Y57G11C.23, Y71G12B.2, F14H3.6, H11L12.1, T04A8.6, Y48G1BL.3, Y62E10A.14, M28.5, F58A4.9, C34B2.2, Y48A6B.3, C35D10.13, C28C12.2, T19H12.2, B0304.2, B0414.3, ZK265.6, F59A2.3, T05B9.1, F46F11.2, F10G7.3, C18G1.5, M01E11.5, F57B10.12, Y17G7A.1, F23B2.6, ZK381.1, T25E12.5, F58A4.3, Y39A1A.14, F57C2.1, C50B8.2, C55B7.11, F44E2.8, Y23H5A.3, Y75B12B.1, K03H4.2, C27C7.1, T24D1.3, H02I12.5, T05G5.3, K10B2.3, Y18D10A.11, F41H10.10, Y102A5C.1, ZC404.8, F26D10.10, C28D4.3, C27B7.5, W02F12.3, R06F6.1, F44B9.7, T06E6.2a, F31E3.3, T19B10.6, T07G12.6, EEED8.1, C08F8.3, C09G9.6, C39E9.12, K01G5.5, C15H11.6, C07H6.5, C13G5.2, F56C9.6, B0393.6, K07A1.1, F56C9.3, C25A1.9, F57B1.2, Y45F10A.2, C27H6.2, AH6.5, T23G11.3, F54C9.6, F38E1.7, C26E6.5, W02A2.7, F26B1.3, F28C6.3, C50B6.2, C07G2.1, F44F4.2, F58A4.4, F54D5.9, T05F1.2, M01E11.6, C36B1.3, T16G12.3, K06A5.7, JC8.6a, T22D1.10, F55G1.8, C38D4.4, F52F12.4, C14B1.9, B0393.3, F32D1.10, C14B9.4a, F30F8.1, Y65B4BL.2, Y111B2A.3, R12E2.10, T21E3.1, T07C4.3a, B0035.6, B0414.5, C05D2.5, C16A11.6, D1081.7, H26D21.2, B0273.2, F18A11.1, F11E6.7, C06A1.4, F58G1.1, C16C10.3, R119.7, C41G7.3, W02D9.1, Y32F6A.1, T25G3.2, T28D9.2a, C07A9.6a, F20B4.2, T16H12.6, T19E10.1a, T28F3.3, Y110A7A.18, Y34D9A.4, Y47G6A.24, Y47G6A.25, Y50E8A.9, Y51H7C.9, Y53F4A.2, Y62E10A.3, Y69A2AR.23, Y69A2AR.24, Y71F9B.7, Y97E10AR.4, Y43C5A.1, T11G6.7, C17C3.5, T28D9.9, R09B3.2, F52B5.6, F43D9.4, R09B3.3, D2089.2, C03D6.6, F58A4.2, R148.4, W08E3.4, C50H2.7, T22F3.4, T05G5.10, F39H2.4, F46A9.4, Y37E11AL.4, F52C6.2, ZK858.3, F15H10.3, B0244.9, Y110A2AR.1, C27A2.3, F53A2.6, Y23H5B.2, C48B4.10, F55H2.3, C29A12.1, M03E7.5, K07H8.9, F36D1.6, Y75B12A.1, T01C3.9, W03A5.5, F52C6.11, Y43E12A.1, F07A5.2, T10B5.6, C03C11.2, F35G12.10, B0218.5, K01G5.3, ZK1248.7, T06C12.4, F08G5.1, W03D2.4, C01G8.6, C25G4.7, C40H1.1, Y14H12B.2, T24A6.1, F13D12.5, T24C4.1, T21B10.4, C03D6.5, C06A8.5, C10G11.8, ZC168.3, F22B7.13, M7.2, C38C10.4, K07C11.2, T27D1.1, F20A1.9, W03G9.2, F22B3.8, F54C9.8, F37E3.3, 6R55.2, ZK177.6, F42A9.6, B0273.3, T02G5.12, Y39A1A.11, B0238.9, C04E6.4, F21H12.5, F58G11.3, K07C5.4, Y53C12A.1, C18G1.4a, C06A5.6, W05F2.6, F57C9.5, K10C8.1, Y48G1BL.1, C50C3.1, C18E3.6, R07B7.2, T11F8.3, M01F1.8, C25A1.3, R06C7.1, C14B1.7, W03C9.7, W06H3.1, F23B12.8, C44E4.3, K10B2.5, C46A5.6, F55A12.1, B0025.2, T06E4.1, F33E11.3, F52B5.5, C16A11.3, ZK1055.1, F26D2.2, F10C5.1, F14D2.4, C18E3.5, W05F2.2a, Y57A10A.30a, B0348.6a, C07E3.1a, C09D4.4a, C13G3.3a, C17E7.9a, C26E6.9a, C37A2.8a, C38D4.1a, C47B2.2a, C53D5.6, C56C10.7a, D1014.8, E01A2.4, F01F1.7a, F07A5.1a, F10G7.8, F10G7.9, F12E12.7, F14H3.3, F14H3.5, F14H3.9, F29G9.7, F30H5.2, F41H10.6a, F45E4.10a, F46B6.3a, F57F4.3, F57F4.4, H06H21.7a, H12C20.2a, K07B1.7a, K08D12.1, T02G5.9a, T07A9.5a, T10C6.7, T10C6.8, T28F3.1, W03F9.2a, Y110A7A.13, Y110A7A.17, Y110A7A.19, Y111B2A.1, Y18H1A.7, Y32H12A.2, Y38B5A.1, Y38E10A.14, Y38E10A.5, Y39A3CR.8, Y39B6A.16, Y39B6A.3, Y41D4B.19a, Y43H11AL.3, Y46G5A.6, Y47G6A.12, Y48C3A.7, Y48G10A.4, Y48G1A.6, Y48G1BL.4, Y48G9A.11, Y49F6B.9, Y4C6A.3, Y50E8A.6, Y51B11A.1, Y53F4B.10, Y54E10A.12, Y54E10A.15, Y54E10BR.6, Y55B1AR.1, Y55D5A.5, Y55D9A.1a, Y55D9A.2a, Y55F3AM.12, Y57A10A.9, Y57E12AL.6, Y61A9LA.10, Y61A9LA.8, Y62E10A.17, Y62E10A.5, Y71F9B.6, Y73B6A.4, ZK484.4a, ZK688.9, F30F8.7, R09H3.2, C25A1.6, T24A6.17, F08B4.3, F10E9.5, F23F12.2, T07F10.5, C04C3.4, C29F9.12, Y54G11A.11, F27C1.12, C04G6.6, W02D9.9, B0464.9, B0391.10, K03B8.4, C46H11.1, ZC404.2, K03H1.7, F23B2.13, T27A8.4, VB0393L.2, T08B2.4, C06A1.5, T24H7.4, Y105E8A.15, ZK652.1, T02G5.6, F59H6.7, Y39A3CL.3, K10D2.5, ZK637.2, W05B10.1, T08D2.4, C15H11.8, W01G7.3, F15A4.10, H06H21.6, T07G12.8, F31E3.6, T08B2.8, C29F9.9, F59A2.5, F55F8.6, C07A9.2, F17H10.4, C12D8.4, F54D10.1, F48G7.9, W04C9.4, D1007.4, ZK287.5, M01E10.3, K02B2.6, ZK593.7, C52D10.7, Y106G6H.16, Y47D7A.1, ZK177.10, T26E3.5, C08B11.9, C30C11.1, W06D11.3, C01B12.8, K03H9.1, ZK970.3, F40F8.1, T22C1.5, K07E8.8, K02B12.2, F32H2.2, T02H6.7, E03H12.8, C52B11.4, K08D12.3a, R05D3.8, C56C10.8, R01H2.6, Y65B4BR.5, E02H1.6, F58A4.10, C41G6.12, F32A5.7, C29A12.2, F58B3.9, Y75B8A.17, T12G3.5, C36B1.7, Y54E5A.5, C48B6.2, T01C3.3, Y43F4B.3, C49C3.6, R05D11.9, C02F4.5, Y38A8.2, C24D10.6, ZK632.8, ZK678.3, C16C10.4, C50F4.4, C14A6.7, Y48E1B.11, C35D10.8, F57B10.11, T10C6.5, 3R5.1, ZK742.5, C16C8.11, W06E11.2, C44B7.1, F43G9.9, F43G9.2, C02B8.1, T20F5.2, C47A4.1, Y102A5C.2, F26H11.1, F13A2.4, C24D10.5, T12D8.2, T24A6.2, B0334.4, F29A7.6, H04D03.1, C16C8.14, F22B5.1, Y49F6C.8, C24D10.4, F59A7.4, W05H7.2, Y54E10BR.2, K08E4.6, R10H10.1, C08F11.7, H06I04.4a, F43G6.10, T06E4.11, T13F2.8, D2007.4, PAR2.1, C15H11.7, F54D5.5, F52C6.1, F39H2.1, T01B7.3, Y6G8.3, F42A6.6, ZK856.11, C02F5.9, D1007.8, B0281.5, B0416.4, T11F8.1, B0047.3, F30F8.3, T05H10.2, C47B2.4, Y54E2A.3, F23C8.9, R04D3.4, C48B6.3, T23B12.1, C25D7.8, F23F1.1, C16C8.13, C35E7.8, C32H11.7, K03B4.3a, R03H4.4, T05C12.5, F21E9.6, F39H11.5, F54C8.2, C50E3.10, T23B12.7, CD4.7, C14B1.8, C33H5.8, K08H2.3, C35D10.10, F46F6.3, W02A11.8, F25H2.4, F53B7.3, Y57A10A.3, M04F3.1, F40E3.2, ZK20.5, F26H9.4, F59H6.8, K02F3.11, F45C12.12, D1081.6, CD4.6, C14A4.1, ZK809.2, T13H5.5, R10A10.2, F45E4.2, Y17G9B.4, F35E12.3, T19B10.11, ZK1058.5, F56C9.9, B0495.9, F56A8.5, M02B7.1, T12G3.6, Y51H7BR.2, F55G1.9, T14B4.3a, K07G5.2, F32D1.7, F55G1.5, B0491.7, T28D9.1, F08B6.1, F39F10.3, T06D8.6, VW02B12L.3, C24F3.2, R07G3.5, T12F5.1, R09F10.8, C16A3.6, F45B8.2, C30A5.3, F40B1.1, C30B5.4, F52D2.2, R05G6.4, ZK353.7, F46F5.4, F52C6.10, T19C4.2, B0238.11, W04A8.5, C54G4.6, F59H6.9, T27A8.2, F45C12.7, B0035.1a, C15H11.9, C49F5.3, Y113G7B.7, H26D21.1, F17A2.13, F07C4.3, T27A1.7, C13A2.4, T28A11.8, C25F9.8, R09E12.3, F45G2.9, ZK418.8, C26B2.6, Y53C10A.13, F46B6.11, R02D3.8, F14D2.1, F56E10.1, F56G4.2, F56G4.3, T08D2.5, T20D3.8, C36B1.4, Y38H6C.21, C45G9.2, C39E9.13, Y113G7B.4, F43D9.5, C01G10.8, C14A4.11, C18E9.6, C16C4.10, F28F8.6, W05G11.2, T26A8.3, ZK1248.12, ZK1248.13, F13G3.6, R10D12.14, Y113G7B.5, ZK795.3, C56C10.9, F57G4.3, F36G9.2, ZK1248.9, T19H5.2, Y18D10A.20, C18E3.9, F26A1.1, F55G1.6, C33H5.7, C16A3.4, F07H5.10, T09A5.9, Y110A7A.11, C50H11.7, F31B9.3, F26F4.1, K06B4.9, C33H5.15, ZK39.2, Y45F10A.7, R10D12.13a, R11A5.2, R05H10.2, M18.7, C31G12.1, W03A3.3, C06A5.8, T06G6.1, Y18D10A.18, VH15N14R.1, ZC168.4, Y45F10B.5, F13E9.1, T23B12.10, B0513.2a, VF36H2L.1, DC2.4, Y53C10A.1, C56A3.5, T06C12.5, T28A11.9, F49C12.8, F36H12.2, ZK430.3, C29F7.4, T23B12.3, Y45F10B.11, F32D1.4, F08G12.10, K02F2.4, T27A3.7, R04D3.6, F32E10.6, F33H1.3, C08B11.5, Y43F11A.6, T06E6.9, F18C5.8, W08E12.7, F10E9.3, F23F12.6, F47C12.10, C02E7.11, F27D4.2, F39B2.5, C40H5.6, T10E9.1, F31D4.3, ZK20.3, M18.6, Y48A6B.5, K04C1.5, C08B11.7, ZK1307.9, B0511.7, K06H6.2, B0207.6, C34D4.10, F54C1.2, T22D1.5, C32B5.2, T04D3.1, H20J04.5, C26E6.6, C29F5.5, T19C4.3, T04D3.5, F15A4.2, B0547.1, F35G12.2, K12H4.3, B0491.1, ZK1098.5, F23F1.5, F45F2.11, C50C3.8, B0205.3, F57C9.7, T24B8.2, B0511.10, C01G5.6, ZK829.5, T27E9.3, K04F10.3, C26B2.1, ZK1127.9a, C24G6.8, F26F4.6, Y49C4A.3, F43H9.3, C42C1.15, B0024.13, ZK1010.2, H20J04.8, F56H1.4, Y69H2.8, K06H6.1, C10H11.10, T25G3.1, B0280.10, C36C5.1, C41G6.11, F21F3.4, Y102A5C.3, F35G12.5, F40G9.14, F29G9.5, K09H9.7, F57B10.8, ZK418.4, C50F7.4, T05A6.2, C41G7.4, T05H10.4, R09B3.1, F23C8.4, T23B12.2, Y64G10A.2, C33H5.17, F38E1.6, T05E11.4, M01E5.4, B0207.7, R08D7.1, T28B4.2, K08F11.4a, F52E1.10, M116.2, M116.4, W02A2.3, JC8.11, C06A5.9, ZK353.8, F29A7.1, B0035.11, B0213.8, C36C9.3, R10E4.5, C52E12.3, Y48G1C.1, Y116A8A.4, H14N18.2, F20E11.1, F19B6.2a, F43G9.10, C01G6.3, F54D5.2, M05D6.2, F56A8.4, C30C11.2, W02B12.3, ZK1248.15, C52E4.4, B0464.8, C02F5.6, F25H2.8, M18.8, F58B3.7, K04F10.6, F43G9.4, T10H9.3, W06E11.4, W01B11.3, F18A11.6, ZC308.3, C14C6.8, F23B12.6, Y45F10D.9, K05C4.4, T20H4.4, F33G12.2, F08F3.6, R12B2.4, F59A3.4, F37C12.13a, T21B10.7, W01G7.4, T10B5.3, Y53C12A.4, H21P03.2, C34D4.8, JC8.5, F15B9.4, B0432.10, C06G3.12, F13C5.3, T13H5.4, D2096.1, F53B2.1, R13A5.12, C24B9.12, T23D5.1, W03G9.6, F56D2.2, F59A1.9, T01G5.7, Y37D8A.12a, R12E2.3, K04G2.2, F37B4.6, Y6E2A.6, K01C8.10, R74.6, F29B9.1, W03G9.3, T14G10.7, C49H3.9, F25H2.12, Y56A3A.17a, T28A8.5, Y4C6B.1, F46F11.7, T10C6.10, M03C11.4, F26B1.6, D2023.6, B0334.11, Y49F6B.2, T07F10.3, F56A8.6, Y57A10A.19, F22D6.3, Y45G12C.14, Y102A5C.33, F42H10.7, D2096.4, C29F5.2, C06E7.1a, T24G10.2, C05E4.11, ZK550.4, ZK1128.5, K01G5.1, K01C8.5, F35G12.12, C34G6.5, ZK20.4, F26H9.1, C31G12.4, C16A11.4, B0024.10, F44A2.1a, K04D7.2a, Y54G9A.5, C04F5.8, F18A1.2, Y56A3A.22, F40D4.7, ZC395.6, F55B12.4, K08F11.6, F52B5.2, Y102A5C.4, F59A1.7, C47B2.1, F59B1.4, W02D9.3, C28A5.3, C25H3.4, T26G10.1, D2096.7, R11A8.1, T27C10.3, T20F5.7, F56D12.5a, C56C10.1, R04F11.3, T26A5.4, Y57A10A.5, T20H4.3, C43E11.2, K08D10.3, T26A8.4, R05D11.4, C05C8.5, F10G7.4, R05D11.6, C27H6.3, ZC317.7, C05C8.6, C14C11.2, R07G3.7, R03D7.2, F33G12.4, F16A11.2, K07C5.6, K02B9.1, C05D9.8, C23G10.4a, C33H5.14, C29H12.1, W01B6.9, Y54E2A.12, F13A2.6, F01F1.5, M04B2.2, K02F3.12, F25B5.5, C34B2.7, T21H8.3, F53F4.11, F26D10.3, ZK856.12, F55A11.4, T01G9.4, Y47H9C.8, C47E8.5, C33F10.4, Y43C5A.6, ZK430.7, T24D1.2, Y71F9AL.8, C43E11.1, C23H3.5, C50F4.11, W07G1.2, Y24D9A.2, Y55H10B.1, T07E3.5, Y57A10A.2, R06C7.7, R11G11.8, Y57A10A.6, F02E9.6, H34C03.1, Y46H3C.4, F29B9.4, R05F9.11, F10B5.2, T26A5.6, ZK546.5, C16C10.1, K09H11.2, F28C1.1, F08H9.1, K08E3.6, ZK328.4, R166.4, R11D1.7, B0205.9, C17E7.5, Y39G10AR.8, Y113G7B.24, C39E9.11, B0414.6, C03D6.3, C06G3.10, T19B10.8, F58A4.8, T24F1.2, K08F9.2, C30G12.3, M18.3, ZK858.5, F37C12.7, Y39G10AR.10, F37D6.2a, F42H11.2, C56C10.11, T04A8.15, E01A2.2,

ZK84.1, T21G5.3, F55A11.7, F22D6.6, K10B2.1, T08G11.4, W02A11.1, F31D4.1, B0495.2, M03D4.1b, C30C11.4, C35D10.9, T16A9.1, C41C4.8,
K07H8.10, C01G5.2, F52H3.4, F26D11.1, C25A1.4, F20H11.1, T10E9.2, K04G2.10, W03C9.2, F25B5.2, T12E12.3, ZK177.4, K06A5.1, F55A3.7,
C47D12.2, C10B5.1, C29H12.5, F02E9.7, F20D12.4, F22F4.2, T23B12.4, F59A2.4, T13F2.7, C44B7.2a, DC2.1, C13B4.2, D2030.6, F55F8.9, T02E1.2,
R06A4.2, K01G5.9, F07A11.3, VC5.4, F40H7.4, ZC155.3, D2089.1a, T10B5.5, F56F4.3, Y53C12B.1, C16A11.5, C54G10.2, F32A5.1, K06H7.6,
Y75B12B.7, K02B9.2, F32H2.3, F46B6.5, F48E8.6, K06B9.4, C18H2.2, T03F1.9, F48E8.2, C16C4.7, Y48B6A.1, ZK353.1a, K04F1.13, C01G6.6a,
F18A12.7, T15B7.14, C03E10.4, C06A1.1, T20H9.6, C46A5.9, F10G8.3, C05D2.10a, C25A1.10, C36B1.12, Y53C12B.3a, C32D5.11, ZK177.1, T02E1.3a,
R06C7.8, B0205.1, Y75B8A.31, F10G7.1, T01B7.5, F33E11.2, C33E10.6, T22D1.9, T12E12.2, F48A11.5, C02F5.1, C06G4.1, C07A9.7, F12F6.1,
ZK1248.10, F13H8.2, R02F2.7, K08F11.3, M01E11.3, D2030.8, H05C05.2, C07H6.4, F45E12.3, C23G10.8, C34C12.2, M01A10.1, W06E11.1, C34E10.4,
T08D2.3, C01G5.8, F19F10.9, F59H6.1, F55A12.5, C01H6.9, C01F1.1, R07E5.1, W07E6.2, K10G4.1, F09G2.4, F10C2.4, F57B10.6, F55F8.3, C16A3.7,
C50F2.3, Y74C10AR.1, T07C12.3, Y48A6B.10, Y54E10A.3, C34E10.5, F57C9.4a, F10E7.8, F30F8.8, K09H11.3, F43G9.12, F54D5.14, F18E2.3, T27F6.5,
M03C11.7, Y52D5A.1, F23F1.8, ZC404.9, C53D6.4, C25H3.6a, F27B3.5, Y38A8.3, F18A12.1, C27F2.7, R13F6.10, F18A1.4a, T12F5.3, K02F6.7, F35A5.1,
T26A5.2a, F32E10.5, F29B9.2a, W02A2.6, F55A12.8, C01H6.5a, F40F8.5, F31C3.2, F56A3.4, F21H12.1, C09H6.3, H43I07.2, F20D12.1, Y82E9BR.2,
W09D6.1, F08B4.5, ZC513.5, Y54E2A.8, F31E3.4, Y39B6A.36, Y43F4B.6, W09B6.3, K04C2.4, F52C9.8a, F45H11.3, F30A10.10, W09B6.2, D1007.7,
B0001.2, C04F5.9, C45G3.1, B0523.3, T19B4.2, C44E4.5, T05H10.1, T04H1.4, Y54E5B.2, H02I12.1, K08E3.3a, W07E6.1, B0546.2, Y65B4A.1, F11A10.1,
R06C1.1, T23H2.3, T17E9.1, H12I19.1, F20D12.2, K03C7.1, ZC395.8, C27A2.1, F54D8.6, ZK1127.11, C08F8.2, R17.2, C36B1.8, F35F11.1, F29D11.2,
D2005.4, Y55F3C.6, F56A3.1, F25H2.13, C45G3.3, E03A3.2, F07E5.5, Y37A1B.1, F21D9.1, Y48A6C.1, C25G4.5, F09G2.8, K04C2.2, T08A11.2, B0464.2,
R03G5.3, D1037.1, C07E3.2, C24H12.4a, F35G12.8, F36D4.5, F16D3.2, F55F10.1, R119.3, D2005.5, F33A8.1, T13H5.2, F56G4.4, H20J04.4, T16G12.5,
F26A3.3, C16A3.3, Y66H1B.3, T07D4.3, K04B12.3, C16A3.8, F26F12.7, C29E4.4, B0564.7, Y69H2.7, ZK328.5b, M03C11.2, F20C5.1, F10B5.7, K02C4.3,
F53C11.5, C37C3.6a, K10D2.3, EEED8.5, ZC376.6, C38D4.3, F37D6.1, Y106G6H.12, M116.5, T07C4.10a, T02H6.2, W03F9.5, Y81B9A.2, W03A3.2,
T11G6.5, W02D3.10, ZK328.7, F17A9.2, F27C1.6, C27B7.4, Y105E8A.17, Y38F2AR.4, Y45F10D.7, K10D2.1, M03C11.8, Y43F8C.14, Y23H5B.6, ZK430.1,
F20G4.3, R02F11.4, Y49F6B.4, Y41C4A.9, F56H1.5, F12F6.5, F18C5.3, C50C3.6, F56B3.4, C17E4.6, K12H4.8, R07G3.3a, C08B6.8, H06I04.1a, F54F11.2,
W03H9.4, F49D11.9, F55F10.2, C04H5.6, K12D12.2, Y37E11AM.1, C47D12.8, C04G2.6, K07E8.7, T23E7.2a, Y57A10A.1, F36A4.7, Y57A10A.31,
C36A4.8, H38K22.1, Y32B12B.4, C07G1.4a, C47E8.8, T06E4.3a, W07A8.2, Y38C9A.1, Y18H1A.6, Y39A1B.3, F25G6.2, Y32B12B.2, W04A4.5,
Y47D3A.29, D2096.3, C04A2.3a, Y47G6A.11, Y54G11A.1, R05H10.3, ZK1151.2a, H06I04.3a, Y53F4B.21, ZC101.2a, Y76B12C.2, F10G8.7, M106.1,
Y18D10A.1, F07C6.4a, Y80D3A.2, C07D10.2, C07G1.3, C41G7.1a, C53H9.2, EEED8.7a, F14D2.3, F22B5.7, F55B12.3a, K11D9.1, M01E5.5a, R13A5.1,
R151.7, T07F8.3, W04D2.6a, Y24D9A.3, Y39B6A.10, Y48G1A.4a, Y48G1A.4b, Y71G12B.9, ZC404.3, ZK637.7, C07A9.6a, F20B4.2, T16H12.6,
T19E10.1a, T28F3.3, Y110A7A.18, Y34D9A.4, Y47G6A.24, Y47G6A.25, Y50E8A.9, Y51H7C.9, Y53F4A.2, Y62E10A.3, Y69A2AR.23, Y69A2AR.24,
Y71F9B.7, Y97E10AR.4, Y43C5A.1, T11G6.7, C17C3.5, T28D9.9, R09B3.2, F52B5.6, F43D9.4, R09B3.3, D2089.2, C03D6.6, F58A4.2, R148.4, W08E3.4,
C50H2.7, T22F3.4, T05G5.10, F39H2.4, F46A9.4, Y37E11AL.4, F52C6.2, ZK858.3, F15H10.3, B0244.9, Y110A2AR.1, C27A2.3, F53A2.6, Y23H5B.2,
C48B4.10, F55H2.3, C29A12.1, M03E7.5, K07H8.9, F36D1.6, Y75B12A.1, T01C3.9, W03A5.5, F52C6.11, Y43E12A.1, F07A5.2, T10B5.6, C03C11.2,
F35G12.10, B0218.5, K01G5.3, ZK1248.7, T06C12.4, F08G5.1, W03D2.4, C01G8.6, C25G4.7, C40H1.1, Y14H12B.2, T24A6.1, F13D12.5, T24C4.1,
T21B10.4, C03D6.5, C06A8.5, C10G11.8, ZC168.3, F22B7.13, M7.2, C38C10.4, K07C11.2, T27D1.1, F20A1.9, W03G9.2, F22B3.8, F54C9.8, F37E3.3,
6R55.2, ZK177.6, F42A9.6, B0273.3, T02G5.12, Y39A1A.11, B0238.9, C04E6.4, F21H12.5, F58G11.3, K07C5.4, Y53C12A.1, C18G1.4a, C06A5.6,
W05F2.6, F57C9.5, K10C8.1, Y48G1BL.1, C50C3.1, C18E3.6, R07B7.2, T11F8.3, M01F1.8, C25A1.3, R06C7.1, C14B1.7, W03C9.7, W06H3.1, F23B12.8,
C44E4.3, K10B2.5, C46A5.6, F55A12.1, B0025.2, T06E4.1, F33E11.3, F52B5.5, C16A11.3, ZK1055.1, F26D2.2, F10C5.1, F14D2.4, C18E3.5, W05F2.2a,
Y57A10A.30a, F14H3.6, H11L12.1, T04A8.6, Y48G1BL.3, Y62E10A.14, M28.5, F58A4.9, C34B2.2, Y48A6B.3, C35D10.13, C28C12.2, T19H12.2,
B0304.2, B0414.3, ZK265.6, F59A2.3, T05B9.1, F46F11.2, F10G7.3, C18G1.5, M01E11.5, F57B10.12, Y17G7A.1, F23B2.6, ZK381.1, T25E12.5, F58A4.3,
Y39A1A.14, F57C2.1, C50B8.2, C55B7.11, F44E2.8, Y23H5A.3, Y75B12B.1, K03H4.2, C27C7.1, T24D1.3, H02I12.5, T05G5.3, K10B2.3, Y18D10A.11,
F41H10.10, Y102A5C.1, ZC404.8, F26D10.10, C28D4.3, C27B7.5, W02F12.3, R06F6.1, F44B9.7, T06E6.2a, F31E3.3, T19B10.6, T07G12.6, EEED8.1,
C08F8.3, C09G9.6, C39E9.12, K01G5.5, C15H11.6, C07H6.5, C13G5.2, F56C9.6, B0393.6, K07A1.1, F56C9.3, C25A1.9, F57B1.2, Y45F10A.2, C27H6.2,
AH6.5, T23G11.3, F54C9.6, F38E1.7, C26E6.5, W02A2.7, F26B1.3, F28C6.3, C50B6.2, C07G2.1, F44F4.2, F58A4.4, F54D5.9, T05F1.2, M01E11.6,
C36B1.3, T16G12.3, K06A5.7, JC8.6a, T22D1.10, F55G1.8, C38D4.4, F52F12.4, C14B1.9, B0393.3, F32D1.10, C14B9.4a, F30F8.1, Y65B4BL.2,
Y111B2A.3, R12E2.10, T21E3.1, T07C4.3a, B0035.6, B0414.5, C05D2.5, C16A11.6, D1081.7, H26D21.2, B0273.2, F18A11.1, F11E6.7, C06A1.4,
F58G1.1, C16C10.3, R119.7, C41G7.3, W02D9.1, Y32F6A.1, T25G3.2, T28D9.2a, B0334.8, B0511.9a, C04A2.7a, C12C8.3a, C12D8.1a, C15F1.4,
C18E9.3a, C24H12.5a, C30G12.6a, C33H5.12a, C45B11.1a, F01D4.5a, F07A11.2a, F14H3.4, F18A1.6a, F52G2.1b, F54C4.2, H04D03.2a, H35B03.2a,
R01H2.3, R05G9.3, R06F6.5a, R08C7.10a, T05G11.1, T05H4.10, T05H4.11, T05H4.14, T19B4.7, T20G5.11, T23D8.9a, W09G3.1, Y105E8A.8,
Y110A7A.16, Y111B2A.18, Y17G9B.9, Y39B6A.13, Y39G10AR.14, Y39H10A.4, Y48G1A.5, Y48G1C.8, Y51H7BR.6, Y55F3AM.15, Y65B4BL.5,
Y69A2AR.28, Y71H2AM.17, Y97E10AR.3, F54D12.4, F28F8.3, Y55H10B.2, Y71F9B.4, Y51H4A.6, Y49E10.15, F35H10.5, K04C2.3, W03C9.5, Y39A3CR.4,
F14D2.11, C09G4.3, F25H5.6, F42H10.2, C33H5.3, C04F12.5, F40F8.9, Y53C12A.6, C02B10.2, F42A9.8, B0336.8, C17E4.4, Y49E10.6, R08C7.3,
F48C1.5, F12E12.1, F40F11.3, B0035.4, Y51H7C.7, T13F2.2, T08A9.4, Y116A8C.42, F35G12.11, B0304.4, F10G2.4, JC8.4, EEED8.13, T06D8.7,
R05D11.7, R05H5.3, C13F10.2, C52D10.9, ZK1098.2a, F22E5.9, C06A8.4, R13F6.1, W08E3.1, F31C3.5, C17H11.4, F21C3.5, T12A2.3, C48B4.7,
F13A7.9, Y43F11A.4, M01D7.6, R10D12.12, C16C8.5, F53A2.5, C43E11.9, F26F4.11, F26A3.2, Y57G11C.25, F01F1.2, EEED8.3, F20D1.4, Y75D11A.2,
F53F8.3, T28D9.10, F29B9.5, C28H8.1, C17E4.5, F56D5.5, Y65B4BR.8, F23H11.1, C14B1.2, F33H2.3, F43G6.7, F11A10.2, B0035.3, ZC395.10, B0336.5,
C24G6.1, F59C6.4, C16C8.12, C16C8.4, F02H6.3a, W05F2.3, C48B4.9, R07E5.14, C05C10.5, K08E4.2, C05D11.3, F57A10.4, T02G5.11, C04H5.1,
C16A3.2, ZK856.10, B0303.15, D2024.5, M04B2.3, F32D1.6, F08B4.7, Y106G6H.15, K01G5.4, D1054.2, R06C7.2, F25H2.9, F52E1.1, F57C2.3, F10E7.5,
T10G3.6, ZK1248.11, F54D10.5, K03B4.2, Y13C8A.1, F32D8.5a, F43G9.5, C49C3.7, F56D1.3, T21C9.13, F59E12.11, R53.6, M88.2, C06G3.8, C13F10.7,
K05C4.7, C41D11.5, T24H10.3, F36F12.8, C55B7.9, Y75D11A.3, F52C6.3, ZK1127.5, C08C3.2, ZK1127.1, C02B10.4, F32E10.2, C34D4.12, C48B4.6,
B0564.1, C16C10.2, ZK643.2, T04H1.5, T26A5.7, C52E4.3, F53A2.4, R06C7.4, F31F6.3, F43D2.1, ZK652.9, Y110A7A.4, C48B6.9, C32E8.5, C50E3.5,
C32D5.5, F54D10.7, T09A5.6, C47B2.5, Y75B12B.4, F55A11.8, C01G10.7, F29G9.1, D2023.5, T24H10.1, T04A8.8, T01B7.4, T07E3.7, W09C3.4,
F21D5.6, F59H6.11, K07D4.3, Y54E5B.3a, F52C6.9, F59H5.3, T14G10.6, K06B9.3, F10E9.4, F33H1.2, T09F3.3, F45E4.9, T23G7.5, F49C12.9, C48E7.3,
B0019.2, W01A8.5, F25H8.1, F45B8.1, ZK1127.4, T09B4.2, W06D11.5, Y62E10A.2, F52C6.8, B0035.15, ZK1010.3, M01E11.1, F53F4.12, T04A8.9,
Y18D10A.17, C47E12.2, F41G3.6, F28F8.5, C02F5.4, T23G11.2, ZC308.4, C33H5.6, D1054.14, C01G8.1, F31F6.1, T01C3.7, ZK546.15, F30A10.5,
C27H8.6, W08F4.3, F02H6.2, K06H7.7, C37C3.9, F47H4.1, T12A2.7, T07A9.1, R74.7, F55H2.4, W02D9.4, R11H6.5, Y49A3A.3, R10E11.4, T23G11.7,
F31C3.4, C49H3.4, W01A11.2, R03D7.7, ZK1307.5, R07E5.3, C14B1.4, C44E4.4, C30F12.4, E02H1.5, W02B12.10, C09H10.6, R12C12.2, T06D8.8,
D1054.3, C17H12.13, C48B4.11, T07F8.4, F18A1.7, Y11D7A.7, F39H2.3, F26A3.7, F48C1.2, F58B3.6, F29F11.3, F59H6.10, D1046.2, K07A1.12,
F42A8.3, T21C9.1, Y95D11A.1, C34E10.2, F10E7.11, R04D3.2, ZC317.1, F33D11.10, W02F12.6, F18A1.3a, F26E4.4, T05G5.7, C03C10.3, C50E3.12,
C36B1.11, Y45F10C.3, D2092.2, F46F11.10, C25A1.12, T09A5.8, F52C6.4, ZK637.11, Y75B8A.18, Y14H12B.1a, E02H1.2, C09B8.5, F25H8.2, ZC513.6,
E02H4.6, C05C8.2, C47E12.7, W02D7.6, F19H6.4, F42A6.5, C08B11.6, F26E4.8, D1086.4, F02H6.4, E02H1.3, F44G4.1, Y43E12A.3, T12F5.2, ZK686.1,
T26C11.7, T24H10.4, F10B5.5, T21D12.3, F30A10.3, R04D3.3, K07A1.11, F27C8.6, ZC328.4, T01B11.3, F42G8.6, C55A6.9, C54G4.9, C26F1.3,
F55G1.7, ZK856.9, C18A3.3, C55B7.5, C53A5.3, B0361.6, F19B10.9, C50F4.12, C03H5.3, R02F2.4, F57G4.8, Y11D7A.12, F16H11.3, C17G10.2,
F54C8.4, F35H8.3, K06B9.2, F48E8.7, C50E3.4, ZK546.14, H02I12.8, B0285.4, ZC53.7, C18A3.1, F56D1.1, T02C12.2, ZK287.7, C46A5.5, F40G12.11,
C08B11.2, K11D12.2, M110.3, F46B6.4, C04B4.2, C17E7.4, R08D7.2, T10B11.8, C56C10.10, F14D7.2, K10D3.3, ZK1128.4, Y110A7A.8, R186.7,
F44G4.4, D1054.15, ZK973.9, T01H3.4, T20B12.3, C08B11.8, Y37D8A.11, C44B9.3, C16C10.6, F49E8.7, F35C11.5, F32H2.4, C27A12.6, Y75B8A.14,
D2030.7, R02D3.7, F17C11.7, T10F2.4, T20B12.2, C17F4.5, T06E6.1, F59B2.6, C17G10.1, T28A8.4, F32D1.9, F44E7.5, F28C6.2, C37C3.1, ZK809.4,
T19B10.7, C13F10.6, C01G5.5, K07F5.14, C26D10.2, F09G2.9, F22B7.6, T07G12.12, Y38A10A.6, ZC477.5, K08D10.4, EEED8.14, K01D12.6, H04D03.3,
F12A10.8, T23G4.3, C06A8.2, C43E11.4, R74.8, C27D9.1, C27A12.7, T19B4.5, C14C11.6, Y37E11B.3, C46F11.3, W03F8.3, T01G9.5, F55F8.5,

K02B12.8, D1046.3, E02H1.4, R11H6.2, C55B7.8, ZK1067.3, C05B10.1, F01F1.8a, R05F9.9, K10D2.2, F13B12.1, B0280.9, K08E7.1, C50B6.3, F14B4.2, ZK863.7, Y43C5A.5, F55F8.4, F28B3.5, K01C8.9, F59E12.13, C15C8.4, Y37A1B.3, C30G12.7, C15H11.5, T22A3.3, Y43F4B.4, R10E11.3a, W06B4.1, F56D1.7, B0280.5, F42A6.7a, T27F6.4, T05C12.7, T06D4.1, F21D5.2, B0024.11, C15C6.4, W02D3.8, T05F1.1, C17E4.3, Y17G7B.12, T24C4.5, F21D5.1, C26D10.1, K08F4.2, Y102E9.2, F26B1.2a, C43E11.10, F38A5.13, Y24F12A.1, F33G12.3, F53F4.14, F11A10.5, F49E8.1, F26G5.1, C01F6.3, C42C1.8, F01F1.11, F32B6.3, K04G7.1, F25B3.6, ZK829.6, Y113G7B.17, B0001.3, W06H3.2, C36A4.4, M01E11.2, C01F6.4, C01F6.1, B0393.2, T28A8.6, T19C3.8, T13F2.6, C56A3.4, ZK973.3, C11D2.4, C36B1.5, F52H3.2, K08F9.4, C31H1.8, Y106G6H.6, Y49E10.14, T07G12.11, T09B4.10, D2030.3, F21C3.4, C28H8.9, C14B1.5, F10B5.6, T28A8.3, E04D5.1a, Y53C10A.6, B0464.6, C56E6.3, ZK938.7, F59G1.5, K12D12.5, F35H10.7, K09H9.2, K06H7.4, T20B12.8, T25G3.3, ZK593.8, C09G4.5a, F22B3.4, Y47G6A.8, F18A1.5, T09B4.1, F45G2.3, F32A11.4, F56D6.6, C32F10.5, F44E2.7, T07A9.8, C32D5.10, ZK858.7, Y5F2A.4, W01G7.5, B0244.8, K01C8.3, DY3.4b, T16H12.1, F56D2.6, ZK863.4, Y62E10A.15, C26E6.7a, F58B3.4, R06F6.4, C41D11.4, C01G8.3, F58G11.5, F52C9.7, B0336.6, C48E7.2, Y39B6A.35, C23G10.7, F28B12.3, C47G2.4, K07H8.1, ZK381.4, F10D11.2, F58G11.6, C27A12.3, R05D11.8, T12E12.4a, F45E12.2, B0336.3, F57B9.10, B0336.7, F45C12.15, ZK973.11, K08E7.3, C08B6.9, F57B10.4, E02D9.1a, F48A11.4, C30B5.1, F49E11.1a, C33H5.4, C48A7.2, F21H12.4, C27A2.6, R119.4, M05B5.5, F55H2.7, F35G12.4a, F28C6.6, M151.4, K07H8.2a, F59E12.1, ZK858.4, C01H6.7, F56F3.1, F02E8.4, C29A12.3a, K04G2.3, ZK1127.7, Y49E10.3, T09A5.10, W02D3.9, ZC302.1, D1007.5, R151.8, T20F7.2, Y39A1A.12, T01B7.6, F36A2.1, F26F4.10, K07C5.8, R10E4.4, B0001.7, C10C6.5, F43G6.9, K02E7.3, W03G1.4, F32E10.1, D1046.1, F52C12.2, W04D2.4, C01B10.8, K05C4.6, F12F6.7, R11E3.7, F52D2.4, T03F6.2, F43C1.3, ZC155.4, C16C8.16, F49E8.2, F39B2.1, Y39E4B.5, C25A11.1, C05C10.6a, ZK858.1, C37A2.4, C35D10.7a, F32A7.4, C06G3.2, F25H5.5, C49H3.6a, K06A5.4, B0035.12, F10C2.6, Y87G2A.7, F38B7.5, Y37D8A.9, M03F8.3, ZK328.2, B0414.8, F23B12.7, K08D10.1, K08F11.2, T01C3.1, R11D1.1a, F55F8.2, C08B6.7, T10B11.3, JC8.7, F53F10.5, T09E8.2, C36C9.1, C05D11.9, K09H9.6, F52C12.1, F59A2.1, C55B7.1, F14D2.8, R10H10.7, Y47G6A.2, F54C9.9, ZK686.4, H28O16.2, F26H11.4, F59E10.1, C32E8.8, F32A7.5, Y39G10AR.11, T02C12.3, R07E5.8, T02G6.5, F56A3.2, F49D11.1, F37E3.1, C17E4.10, R05D3.4, F53F8.5, C38D4.6, B0365.1, F08F3.2, K07F5.13a, F32H2.1a, C04E6.11, F54E12.2, F55C5.4, F17C11.10, Y52B11A.2, T10F2.3, W02A11.4a, T23D8.7, B0285.5, T09F3.4, Y54E10A.6, F44B9.6, Y71F9B.2, T08G5.5, F59A3.2, C33F10.2, F26F4.7, R09A8.2, T11G6.8, F11A10.3, F45F2.10, K08F4.1, R11A5.1a, F57C9.3, C36A4.5, F25D7.4, ZK632.5, Y40D12A.1, F10E9.8, W09C5.2, Y2H9A.1, T05F1.6a, F59A6.5, ZK632.1, C29E4.2, K07A12.2, F18C5.2, C27A12.2, T24C4.7, F26E4.10, T05E7.3, C06A5.3a, Y71H2B.2, F44C4.4, T20B12.1, D1081.8, C25A1.7a, T07D4.4a, F33H2.2, T27F2.1, T14B4.1, C14A4.4, R74.5, T24D1.1, F25G6.9, T20F5.6, W04A8.1, C03D6.4, C28A5.1, C28A5.2, T19A5.1, T26A5.5a, T07A9.6, Y52B11A.9, F33H1.4, W08D2.7, K05C4.5, F53G12.5a, F52C12.3, D1044.6, C34E10.8, Y55B1BR.2, C44B7.10, Y17G7B.5, F42A6.3, F33D11.9b, W06D4.6, T23F6.4, ZK546.13, F54F2.5, T04A8.14, F21G4.2, K04G2.8b, K04D7.5, F53A3.2, K12D12.1, Y48G8AL.7, F26G1.1, F14B4.3, C41D11.7, R05D3.11, Y51H7C.6a, K11H3.4, F52B5.3, W07E6.4, C42D4.8, T28A8.7, Y76B12C.6, Y119D3B.11, F33H2.1, D1043.1, F52B11.1, B0240.2, F16D3.4, F37A4.8, R11E3.6, F54C8.3, F28B3.7, R13H4.4, C46C11.1, F56A3.3a, C37A2.2, R05D3.1, Y116A8C.36, T23H2.1, C50F2.2, W04A8.6, Y55F3AM.3a, T23B5.1, R119.1, F56F11.4, Y47G6A.6, Y102A5C.18, C49H3.10, T05E8.3, W04B5.5, F02A9.6, F29G9.2, ZK1251.9, W10C6.1, F09F7.3, D2045.2, T04A11.6, C32F10.2, R06A4.4a, T22A3.5, C26C6.1, F02E9.4, Y53C10A.12, F53A2.8, T23E1.2, ZK742.1, F10C2.2, C37H5.5, F09E8.3, Y47D3A.28, R09A1.1, K02B12.5, F54D11.2, F36A2.13, Y37D8A.13, Y41E3.9, F09D1.1, C44B9.4, Y87G2A.1, F54E7.3a, W03G1.6, C16C2.3, Y116A8C.13, F15D4.1, Y47D3A.26, B0361.2, M03A1.1a, K02F2.3, C01F6.8a, F36H1.4, T01H8.1a, T05C12.6a, T07A9.5b, Y39G10AR.19, Y44E3B.1, Y54H5A.2, Y56A3A.29a, Y57E12AL.1, Y62F5A.1a, Y69A2AR.30, Y71F9B.12, Y71G12B.14, Y77E11A.7, ZK546.16, C05B5.2, F37A8.2, F40F12.3, F54A3.4, F58D5.8, K03H1.1, M70.3b, T05A7.6, T16G12.7, T23F6.3, T25C8.3, Y116A8A.6, Y23H5B.9, Y39G10AR.16, Y43F8C.5, Y46C8AL.1, Y46G5A.30, Y47D3A.22, Y50E8A.12, Y50E8A.2, Y51A2B.6a, Y52D5A.2, Y59E9AL.6, Y59E9AR.7, Y65B4A.9, Y71G12B.27, Y73B6A.1, Y73F8A.12, Y73F8A.14, C40H5.1, H12D21.1, W06A7.5, ZC412.6, ZC412.7, C25A8.1, C33F10.10, F21D9.3, W01B6.4, F36H12.13, K07F5.5, ZK484.8, C24D10.7, C24D10.8, F11G11.8, T23B7.1, C47A4.5, Y45F10B.3, C32E8.4, F09C12.7, ZK1248.4, ZK84.5, K05F1.8, ZK1248.17, F32B5.4, ZK484.5, C03B8.1, F46F5.9, ZC168.6, ZK1225.6, C35D10.11, C55C2.2, E03H12.10, F44D12.7, T27A3.3, T28H11.6, F02C9.1, C02F5.2, F36H12.4, E03A3.4, C04G2.4, C09B9.6, C33F10.9, C34F11.4, C34F11.6, F26G1.7, F32B6.6, F36H12.7, F58A6.8, K05F1.2, K05F1.7, K07F5.2, K07F5.3, R05F9.13, R05F9.8, R13H9.2, R13H9.4, T13F2.10, T13F2.11, ZK1248.6, ZK1251.6, ZK354.1, ZK354.11, ZK354.4, ZK354.5, ZK546.6, F08H9.2, C27D6.3, F35H8.4, W02D9.5, Y106G6H.13, Y37D8A.8, F35H8.1, Y67A6A.1, C18A3.7, C34D4.3, ZC412.9, C34F11.2, K03H1.12, C34G6.3, K07F5.7, F36A4.3, Y46G5A.14, K01D12.15, ZC116.2, T14D7.3, Y51B9A.5, R05D3.5, ZC262.1, C09F9.1, F29D10.1, F36A2.10, C48E7.7, T08B2.12, ZK1251.1, C25D7.1, C24D10.2, ZK617.3, F42E8.2, F52F12.5, F11G11.4, F36A4.2, F08G2.6, Y116A8A.7, T22B3.3, Y73F8A.15, C07G1.6, F36A4.4, H09I01.1, R05F9.3, T13A10.1, W09D6.4, ZK1307.3, F44G4.5, C15A11.2, D2062.4, ZK930.7, F47B8.11, C38C3.8, T15H9.5, C27F2.6, F46A8.9, T13F2.12, F32B6.7, K06A5.3, F36D1.4, T14G10.8, ZK353.3, T01H3.5, C02F5.5, Y49E10.7, C26E6.1, C05C12.5, C10G11.9, T27A3.4, F37A8.1, C26C6.6, C18H7.7, F17E9.5, F26F4.2, K01D12.7, F43E2.10, Y106G6A.4, F36H12.5, F41G3.4, C25A8.2, ZK1307.4, F02C9.4, F10D11.3, F46B3.4, Y106G6E.2, F58F12.2, C33A12.15, B0432.12, E03H12.5, B0432.11, T23B3.5, F29B9.7, T20F5.5, Y38F1A.7, C41G7.6, C01G12.3, W09C3.7, E03H12.7, F28H7.4, ZK930.4, C26B2.2, ZK945.6, K08F4.5, F56F4.7, ZK1251.3, C10H11.7, B0496.6, F33D11.1, T02E1.6, T10E9.6, Y43C5B.2, Y47G6A.3, ZC477.7, R02D5.7, F49E12.4, ZC477.1, ZK546.3, B0379.2, ZK637.12, F42A9.3, F32B4.2, R13H9.1, B0041.1, T28H11.7, Y106G6E.3, F27C1.1, F47B3.2, F46A9.1, R01H2.4, R07E5.15, T21G5.4, ZK783.3, C25G4.6, T13F2.9, C06A5.2, F32B6.5, Y38F1A.1, ZK354.7, Y45F10B.8, ZK637.15, ZK520.5, F07D3.3, C04F12.7, C55B7.10, K07F5.11, B0273.1, C04G2.5, F41G3.5, F28E10.4, F54C8.1, Y57A10B.7, C17F3.3, C29F5.3, C54G4.3, Y116A8C.37, C16A11.7, T06E4.5, F49F1.12, AH6.3, K05F1.9, F58E6.5, B0491.3, T02H6.4, F36D3.4, K07F5.8, C47E8.1, K01H12.2, T03F1.5, C17D12.5, H12D21.2, T23F11.2, R09E10.6, W03C9.1, C35D10.3, T01B11.4, F54C1.8, ZK795.2, F47B3.5, F40F9.3, K11H3.2, W09C3.6, ZC317.6, W03F11.3, T08G11.2, F59C6.2, F11G11.9, F20D6.6, F55C5.2, F21H7.5, T27E7.1, F21F3.3, W01C9.4, F38H4.6, C45G9.9, F54D1.1, C45G9.4, T08H10.4, C04G2.9, C09H5.7, C43G2.3, T05C12.1, Y1A5A.1, ZK546.7, C47E12.10, W06D4.2, F56H11.3, C08F11.10, ZK1248.5, K07A12.5, C35E7.9, F47B3.4, F36H12.8, R13H9.5, Y59E9AL.1, Y43F8C.9, ZK354.3, F44D12.4, Y81G3A.1, C35D10.2, K06A5.2, R102.3, R10E4.7, C50D2.3, C25D7.12, C01G10.14, T16A9.3, F38A5.6, F55D12.6, C17C3.11, C14C10.1, K07A1.4, C28D4.7, C28D4.8, Y69E1A.2, F53B6.7, ZK938.1, C34D4.2, F52H3.6, T28H11.1, F46A9.2, C17H12.3, C47A4.3, C33H5.16, D1081.5, W02A2.8, ZC412.5, F36H12.9, R13H9.6, ZK858.2, F13G11.2, C06A8.6, F26A3.5, AH6.2, R10E9.2, ZK512.8, T05C12.3, F36H12.3, C33F10.12, F40H6.1, T05F1.8, F42G4.6, F44F4.10, B0511.4, Y69E1A.4, Y106G6G.1, C02B10.6, F36H12.10, T16A9.5, T27A3.5, F53B6.4, C47E12.11, Y53F4B.36, F22B5.5, K02F6.2, C15H7.3, ZK84.2, Y69E1A.8, F28A10.1, F36A2.11, F18C5.4, T28H11.5, F53C3.1, F17A9.1, F10G8.1, C55B7.3, T04B2.2, C39H7.1, B0261.6, C56A3.8, F35C11.3, M05D6.1, Y38H8A.3, ZK265.3, F36H12.14, T28C6.5, ZK673.6, W03D8.10, ZK484.7, C09H10.9, R08C7.8, D2045.5, K08D10.7, B0379.7, F02E9.3, C52E4.7, B0218.7, K06H7.8, C27D8.1, C30B5.3, F58A6.5, C38C10.3, ZK930.6, K11C4.1, F44D12.6, K08C9.2, M02B1.4, F23B12.1, C15H7.4, T23B3.3, Y6E2A.9, Y106G6G.4, Y69E1A.1, M05B5.1, C29E6.3, ZK1098.6, F32H2.7, T25B9.2, C08F8.6, Y37D8A.5, T20H4.2, F48C1.7, ZK354.6, F23B2.7, C43E11.5, C50F2.5, F42C5.5, M05B5.3, C49C3.2, F47B3.1, C25A6.1, C56C10.6, ZK507.1, W09C3.8, C30A5.4, F57B9.8, C14C11.1, K08C9.1, M176.9, T01C3.5, K05F1.3, F56F3.4, K01H12.4, T28C12.3, T09B4.7, T08B2.11, C46E10.1, F13A7.1, R09E10.1, C04G2.2, T05F1.5, Y38H8A.4, F58H1.6, T09F5.10, F21F3.2, C36B1.10, C17H12.5, ZK622.1, C36H8.1, W01B6.2, T13A10.11, F26B1.5, C09B9.4, Y18D10A.21, F38A5.11, C15C6.2a, C18G1.9, F08G5.2, C32D5.4, ZK354.2, C06A1.3, C05C12.1, C09B9.2, F22D6.9, C24H11.1, F40G12.10, Y44A6D.5, T02B11.2, K08F8.5a, C33F10.8, R09E10.4, W01B6.6, H06H21.9, W02A2.4, Y45F10B.2, C31H1.5, BE10.1, F32A11.3, F54H5.2, F47B3.7, ZK550.5, R148.7, R10D12.10, C25G4.4, ZC581.7, C50F7.3, Y43C5B.3, T02E1.7, F14H3.2, F56A11.6, F56D5.2, C47D12.3, C55C3.4, F25H5.7, B0523.1, F49E11.7, C53A5.4, T28F4.3, B0252.5, M7.7, F21D9.2, W08D2.8, F44D12.8, C15H11.1, C25A8.5, D2024.1, ZC513.10, K08D10.8, ZK593.9, C27D8.2, R01H2.2, Y69A2AR.6, F26A1.3, C32C4.3, F57A8.6, T13H10.1, T09A12.1, F45E4.6, F57F4.1, F37H8.4, C24D10.1, C14A4.13, H08M01.1, C18E9.8, Y116A8C.24, Y116A8C.38, T04A8.3, C34B2.3, C09D4.3, Y106G6D.3, C43F9.6, ZK418.2, ZK354.8, C07E3.4, T06C10.6, W04E12.4, T15B12.2, Y37E11B.10, F56D6.5, F57H12.5, M05D6.3, F35C11.2, F55F8.7, F26F4.8, R11E3.1, ZK1290.6, F53B2.5, Y40H4A.2, ZK1225.4, ZK1127.2, F33D11.2, AH10.1, Y4C6A.1, F36A2.12, K09E4.1, T25B9.5, F10F2.8, C32E12.1, Y69E1A.3, T06C10.3, C01F6.2, F58A6.11, C48B6.4, C49C8.1, R05D7.2, ZK1225.5, F58G1.3, F42G8.9, F47B3.6, C05D2.3, F42G4.2, M162.7, W02B12.12, Y57G7A.5, R10H10.2, C46H11.6, C09G9.4, F47C12.4, C08F8.4, ZC581.2, W01B6.5, T11F8.4, Y57G11A.2, F59B2.5, M04G7.2, F25F2.1, T25B9.6, Y32F6A.2, F47D12.7, F02E11.1, R08C7.5, T08B6.4, ZC513.3, Y70C5C.5, F46A8.10, T21G5.1, C44B9.2, Y53F4B.11, C04E6.5, F27C8.5, C50F4.2, Y57G11B.3, R05H5.2, F10F2.7, K08A2.2, F23C8.7, C18H7.4, K09B11.5, ZC581.6, F27E5.3, F21H7.2, F53G12.6, C01G5.3, F19B6.4, T08G3.4, R09E10.3, F23C8.8, F01G10.5, R13G10.4, F59A3.8,

F36H1.3, T26A5.1, C38C3.7, C55C3.3, C53B4.2, K09C6.8, T04F3.3, F49C12.15, F36D3.8, F42G8.8, C46A5.1, T06D4.4, C31H1.1, C17H12.12, C38C3.3, Y73F8A.20, F26B1.1, K09C6.7, R03D7.8, C34F11.5, F18A12.4, F58D2.2, Y49E10.10, F55H12.1, F55A11.6, F07E5.8, F55H12.2, Y57G7A.6, F44F1.3, B0393.4, ZK250.6, M28.9, T22B3.2a, T04A6.3, D1037.5, K08E4.5, K02F6.3, T10E9.4, F32H2.9, R07C3.4, M01A12.1, T22H9.3, C01G10.1, C01G12.8, F09C12.2, T08B6.5, F26D2.10, W03B1.9, F35E2.6, K02E11.1, Y116F11B.7, H05L14.1, Y43F8A.2, Y69A2AR.19, F17C8.3, R102.7, Y39B6A.30, T04A8.13, F54H12.5, C28A5.6, R13A1.3, Y113G7C.1, M70.1, F37A4.7a, C24F3.5, Y53C10A.10, F34D10.2, F47G6.4, Y59E9AL.3, C05B5.1, C15F1.5, Y110A7A.12, Y22D7AR.12, Y39G10AR.15, Y40C5A.3, Y46G5A.25, Y47D9A.5, Y47G6A.26, Y49F6B.8, Y53F4B.35, Y59E9AL.5, Y69A2AR.27, Y73B6A.2, Y76B12C.4, Y77E11A.10, ZK666.2, R02F2.6, F58G1.9, K10H10.7, W03F9.3, R04B5.11, Y11D7A.16, K08E7.4, Y53C10A.2, F59A6.2, K06A4.6, AC8.2, C03C11.1, F19C7.5, C01G6.2, Y54E2A.9, D2062.5, ZK484.6, K03H1.8, C17C3.9, ZK930.5, R02F2.5, ZK945.7, C34F11.1, T21E12.5, F38H4.5, ZK1290.7, B0457.3, C24A11.1, Y54E2A.7, Y47D9A.3, Y53F4B.34, C10G11.1, B0511.3, Y53G8AM.1, W01D2.3, H06O01.4, F53G12.8, ZK353.4, F19B6.3, ZK1251.5, Y41C4A.7, F14F7.4, F46F5.2, C07A9.5, ZK1290.9, F32B6.10, F21A3.4, F09C6.5, F09C6.4, Y57G11C.8, W03A5.2, F59E12.3, C09G5.7, Y43C5A.4, T05H4.2, C18E3.1, T15H9.4, F42H11.1, ZK354.9, R03D7.5, F10E9.2, C10C6.3, Y69A2AR.9, F23C8.1, K07F5.6, F28A10.4, F20H11.4, F26E4.5, W03G1.2, F38E1.3, Y24D9B.1, Y54E2A.5, Y39G8C.2, Y53G8AR.6, K07F5.4, F13H8.8, F42G9.1, F38H4.4, C56G2.5, D2092.7, ZK973.8, Y27F2A.3, W09C3.2, F12A10.3, C31H1.2, C23G10.1a, F28H7.6, F07F6.1, C37A2.3, W03G9.5, T22C1.8, C16D2.1, F35E2.5, K01C8.8, T27F6.1, C50E10.2, B0207.1, C18G1.3, T05D4.5, M04F3.3, F59A6.4, C09B9.7, C49G7.1, C17E4.1, B0513.6, Y66A7A.4, F46F5.11, B0280.11, R09E10.2, C48E7.8, B0001.1, F18A12.5, T22A3.6, W03D8.5, H32C10.1, W02B12.7, T27E4.6, M110.7, F48F5.1, R155.3, F18A1.1, F54F12.1, K09F6.3, Y53G8AR.3, F30F8.2, K01A11.4, F47F6.5, R06B10.2, H32C10.3, F15D3.4, C18H2.4, W01B11.2, W03F11.4, F56H1.3, F35E2.9, Y80D3A.8, Y49E10.19, R155.2, Y53C10A.9, C09B9.3, T28B8.4, Y53G8AR.7, F43G9.6, D1081.3, C18H2.1

| Genome project | Gene stable ID | Chromosome/ scaffold name | Gene start (bp) | Gene end (bp) | Strand | Trichinella spiralis (PRJNA12603) gene stable ID | Trichinella spiralis (PRJNA12603) gene name | Trichinella spiralis (PRJNA12603) start (bp) | Trichinella spiralis (PRJNA12603) chromosome/ scaffold | Trichinella spiralis (PRJNA12603) end (bp) |
|---|---|---|---|---|---|---|---|---|---|---|
| caenorhabditis_elegans_prjna13758 | WBGene00007065 | III | 13780220 | 13781133 | 1 | Tsp_03716 | | 220783 | GL622788 | 222607 |
| caenorhabditis_elegans_prjna13758 | WBGene00007082 | V | 14146499 | 14148784 | 1 | Tsp_10497 | | 1167057 | GL622791 | 1168176 |
| caenorhabditis_elegans_prjna13758 | WBGene00007082 | V | 14146499 | 14148784 | 1 | Tsp_10496 | | 1164873 | GL622791 | 1166542 |
| caenorhabditis_elegans_prjna13758 | WBGene00007080 | II | 9517567 | 9518876 | -1 | Tsp_11550 | | 735716 | GL622790 | 741190 |
| caenorhabditis_elegans_prjna13758 | WBGene00007080 | II | 9517567 | 9518876 | -1 | Tsp_12889 | | 454 | GL626579 | 1719 |
| caenorhabditis_elegans_prjna13758 | WBGene00003231 | II | 9524174 | 9526390 | 1 | Tsp_12414 | | 8096 | GL624002 | 8753 |
| caenorhabditis_elegans_prjna13758 | WBGene00003231 | II | 9524174 | 9526390 | 1 | Tsp_08560 | zfp36l2 | 4699442 | GL622788 | 4700099 |
| caenorhabditis_elegans_prjna13758 | WBGene00003231 | II | 9524174 | 9526390 | 1 | Tsp_08559 | Zfp36l3 | 4695882 | GL622788 | 4696511 |
| caenorhabditis_elegans_prjna13758 | WBGene00003231 | II | 9524174 | 9526390 | 1 | Tsp_12415 | | 11512 | GL624002 | 12169 |
| caenorhabditis_elegans_prjna13758 | WBGene00003231 | II | 9524174 | 9526390 | 1 | Tsp_12413.2 | | 5119 | GL624002 | 5437 |
| caenorhabditis_elegans_prjna13758 | WBGene00003231 | II | 9524174 | 9526390 | 1 | Tsp_08561 | | 4702854 | GL622788 | 4703511 |
| caenorhabditis_elegans_prjna13758 | WBGene00007100 | V | 10315245 | 10317394 | -1 | Tsp_03004 | | 2295904 | GL622785 | 2298406 |
| caenorhabditis_elegans_prjna13758 | WBGene00007100 | V | 10315245 | 10317394 | -1 | Tsp_14353 | | 1440 | GL625580 | 1821 |
| caenorhabditis_elegans_prjna13758 | WBGene00007101 | V | 10317490 | 10319534 | -1 | Tsp_03113 | | 2821299 | GL622785 | 2826101 |
| caenorhabditis_elegans_prjna13758 | WBGene00000814 | I | 6022660 | 6026874 | 1 | Tsp_12920 | | 51 | GL624478 | 1550 |
| caenorhabditis_elegans_prjna13758 | WBGene00000814 | I | 6022660 | 6026874 | 1 | Tsp_05853 | | 5553253 | GL622785 | 5554722 |
| caenorhabditis_elegans_prjna13758 | WBGene00007110 | IV | 11327576 | 11329369 | -1 | Tsp_05978 | | 6065507 | GL622785 | 6068114 |
| caenorhabditis_elegans_prjna13758 | WBGene00007111 | IV | 11329466 | 11332969 | -1 | Tsp_06855 | | 8187026 | GL622785 | 8193801 |
| caenorhabditis_elegans_prjna13758 | WBGene00007106 | IV | 11314078 | 11315116 | -1 | Tsp_07852 | | 10925950 | GL622785 | 10926905 |
| caenorhabditis_elegans_prjna13758 | WBGene00004466 | I | 10730712 | 10732448 | -1 | Tsp_01500 | | 5935932 | GL622787 | 5938440 |
| caenorhabditis_elegans_prjna13758 | WBGene00015029 | I | 5935370 | 5936957 | -1 | Tsp_01904 | | 7798250 | GL622787 | 7799356 |
| caenorhabditis_elegans_prjna13758 | WBGene00015030 | I | 5945581 | 5947620 | -1 | Tsp_00226 | | 857479 | GL622787 | 859843 |
| caenorhabditis_elegans_prjna13758 | WBGene00015049 | IV | 8155211 | 8156587 | -1 | Tsp_02051 | | 8391493 | GL622787 | 8393242 |

| | | | | | | | | | | |
|---|---|---|---|---|---|---|---|---|---|---|
| caenorhabditis_elegans_prjna13758 | WBGene00015049 | IV | 8155211 | 8156587 | -1 | Tsp_06549 | | 11192741 | GL622787 | 11195348 |
| caenorhabditis_elegans_prjna13758 | WBGene00015049 | IV | 8155211 | 8156587 | -1 | Tsp_00662 | Ttbk2 | 2539389 | GL622787 | 2540670 |
| caenorhabditis_elegans_prjna13758 | WBGene00015049 | IV | 8155211 | 8156587 | -1 | Tsp_04341 | Ttbk1 | 3010978 | GL622788 | 3012100 |
| caenorhabditis_elegans_prjna13758 | WBGene00015049 | IV | 8155211 | 8156587 | -1 | Tsp_06889.2 | | 8369982 | GL622785 | 8372542 |
| caenorhabditis_elegans_prjna13758 | WBGene00007117 | V | 11733733 | 11736968 | -1 | Tsp_06250 | | 10255084 | GL622787 | 10258408 |
| caenorhabditis_elegans_prjna13758 | WBGene00015104 | III | 7120905 | 7122895 | -1 | Tsp_01679 | | 6868296 | GL622787 | 6869462 |
| caenorhabditis_elegans_prjna13758 | WBGene00015104 | III | 7120905 | 7122895 | -1 | Tsp_15885 | | 149 | GL627398 | 1250 |
| caenorhabditis_elegans_prjna13758 | WBGene00007137 | III | 4343446 | 4345366 | 1 | Tsp_08223 | LCMT1 | 2490491 | GL622789 | 2491537 |
| caenorhabditis_elegans_prjna13758 | WBGene00002003 | III | 4345339 | 4350079 | 1 | Tsp_05727 | | 4821785 | GL622792 | 4823901 |
| caenorhabditis_elegans_prjna13758 | WBGene00015133 | III | 8717644 | 8718567 | 1 | Tsp_06271 | | 10162361 | GL622787 | 10164468 |
| caenorhabditis_elegans_prjna13758 | WBGene00007144 | II | 11495692 | 11496663 | 1 | Tsp_07093 | | 86989 | GL622784 | 88108 |
| caenorhabditis_elegans_prjna13758 | WBGene00015143 | III | 5711548 | 5714846 | 1 | Tsp_11096 | | 2198678 | GL622784 | 2208838 |
| caenorhabditis_elegans_prjna13758 | WBGene00015146 | III | 5690107 | 5692740 | -1 | Tsp_00560 | Abi2 | 2211480 | GL622787 | 2212736 |
| caenorhabditis_elegans_prjna13758 | WBGene00015156 | III | 7293632 | 7308364 | 1 | Tsp_12298 | | 1887792 | GL622784 | 1890642 |
| caenorhabditis_elegans_prjna13758 | WBGene00015156 | III | 7293632 | 7308364 | 1 | Tsp_07121 | | 159973 | GL622784 | 161466 |
| caenorhabditis_elegans_prjna13758 | WBGene00015160 | III | 7275502 | 7277274 | -1 | Tsp_01157 | | 4491416 | GL622787 | 4495426 |
| caenorhabditis_elegans_prjna13758 | WBGene00007150 | V | 13116488 | 13120167 | 1 | Tsp_01728 | | 7071336 | GL622787 | 7076470 |
| caenorhabditis_elegans_prjna13758 | WBGene00000772 | I | 5788634 | 5791752 | -1 | Tsp_10790 | | 3209270 | GL622789 | 3212283 |
| caenorhabditis_elegans_prjna13758 | WBGene00001600 | I | 5792324 | 5795114 | -1 | Tsp_12124 | | 535552 | GL622790 | 536415 |
| caenorhabditis_elegans_prjna13758 | WBGene00001600 | I | 5792324 | 5795114 | -1 | Tsp_15219 | | 119 | GL623636 | 662 |
| caenorhabditis_elegans_prjna13758 | WBGene00015176 | I | 5803959 | 5807291 | -1 | Tsp_10687 | | 2836358 | GL622789 | 2837831 |
| caenorhabditis_elegans_prjna13758 | WBGene00015176 | I | 5803959 | 5807291 | -1 | Tsp_10689 | | 2835087 | GL622789 | 2835732 |
| caenorhabditis_elegans_prjna13758 | WBGene00007184 | III | 9491640 | 9497044 | 1 | Tsp_02172 | | 8895404 | GL622787 | 8901352 |
| caenorhabditis_elegans_prjna13758 | WBGene00007188 | III | 9465767 | 9467177 | 1 | Tsp_01169 | | 4704851 | GL622787 | 4706709 |
| caenorhabditis_elegans_prjna13758 | WBGene00007189 | II | 11348430 | 11349946 | 1 | Tsp_10673 | | 2716682 | GL622789 | 2718806 |
| caenorhabditis_elegans_prjna13758 | WBGene00007194 | II | 11350192 | 11351373 | 1 | Tsp_10674 | | 2714732 | GL622789 | 2715945 |

| | | | | | | | | | | |
|---|---|---|---|---|---|---|---|---|---|---|
| caenorhabditis_elegans_prjna13758 | WBGene00015203 | II | 7699254 | 7702012 | 1 | Tsp_10993 | | 9685516 | GL622785 | 9689677 |
| caenorhabditis_elegans_prjna13758 | WBGene00001228 | I | 10640465 | 10642281 | -1 | Tsp_04424.2 | | 3410600 | GL622788 | 3411573 |
| caenorhabditis_elegans_prjna13758 | WBGene00001228 | I | 10640465 | 10642281 | -1 | Tsp_15148 | | 13 | GL627104 | 977 |
| caenorhabditis_elegans_prjna13758 | WBGene00001228 | I | 10640465 | 10642281 | -1 | Tsp_12760 | | 96 | GL623059 | 1190 |
| caenorhabditis_elegans_prjna13758 | WBGene00001228 | I | 10640465 | 10642281 | -1 | Tsp_15643 | | 323 | GL623586 | 1112 |
| caenorhabditis_elegans_prjna13758 | WBGene00015233 | I | 10631665 | 10633150 | -1 | Tsp_08168 | | 2204578 | GL622789 | 2205585 |
| caenorhabditis_elegans_prjna13758 | WBGene00015249 | IV | 3373387 | 3378130 | -1 | Tsp_04017 | | 1510331 | GL622788 | 1526730 |
| caenorhabditis_elegans_prjna13758 | WBGene00044083 | IV | 13098978 | 13099859 | 1 | Tsp_00048 | | 285640 | GL622787 | 287331 |
| caenorhabditis_elegans_prjna13758 | WBGene00015282 | IV | 6635039 | 6638155 | 1 | Tsp_10574 | | 6049893 | GL622792 | 6050676 |
| caenorhabditis_elegans_prjna13758 | WBGene00015282 | IV | 6635039 | 6638155 | 1 | Tsp_05984 | | 6316252 | GL622785 | 6318664 |
| caenorhabditis_elegans_prjna13758 | WBGene00015282 | IV | 6635039 | 6638155 | 1 | Tsp_10580.2 | | 6078786 | GL622792 | 6080403 |
| caenorhabditis_elegans_prjna13758 | WBGene00015282 | IV | 6635039 | 6638155 | 1 | Tsp_10580.1 | | 6141807 | GL622792 | 6144068 |
| caenorhabditis_elegans_prjna13758 | WBGene00015287 | II | 6664 | 9486 | 1 | Tsp_04355 | | 3070205 | GL622788 | 3072576 |
| caenorhabditis_elegans_prjna13758 | WBGene00015287 | II | 6664 | 9486 | 1 | Tsp_13862 | | 585 | GL628400 | 1618 |
| caenorhabditis_elegans_prjna13758 | WBGene00015287 | II | 6664 | 9486 | 1 | Tsp_02752 | | 1212813 | GL622785 | 1214864 |
| caenorhabditis_elegans_prjna13758 | WBGene00015296 | II | 4301607 | 4305287 | 1 | Tsp_01672 | | 6547172 | GL622787 | 6549416 |
| caenorhabditis_elegans_prjna13758 | WBGene00007234 | V | 15085747 | 15087064 | 1 | Tsp_09267 | Clybl | 5203037 | GL622788 | 5204011 |
| caenorhabditis_elegans_prjna13758 | WBGene00007235 | V | 15084059 | 15085382 | -1 | Tsp_02541 | | 366241 | GL622785 | 368940 |
| caenorhabditis_elegans_prjna13758 | WBGene00015308 | IV | 6522874 | 6524568 | 1 | Tsp_05024 | | 1779167 | GL624340 | 1781017 |
| caenorhabditis_elegans_prjna13758 | WBGene00015310 | IV | 6548818 | 6552511 | 1 | Tsp_08014 | | 1411158 | GL622789 | 1414870 |
| caenorhabditis_elegans_prjna13758 | WBGene00000965 | I | 5278803 | 5281526 | -1 | Tsp_08944.2 | | 2513297 | GL622784 | 2514018 |
| caenorhabditis_elegans_prjna13758 | WBGene00000965 | I | 5278803 | 5281526 | -1 | Tsp_04022 | | 1550849 | GL622788 | 1551646 |
| caenorhabditis_elegans_prjna13758 | WBGene00007258 | I | 7229433 | 7233121 | -1 | Tsp_06552 | GSG2 | 11176441 | GL622787 | 11180078 |
| caenorhabditis_elegans_prjna13758 | WBGene00015347 | III | 8244445 | 8245794 | 1 | Tsp_02036 | | 8319421 | GL622787 | 8320983 |
| caenorhabditis_elegans_prjna13758 | WBGene00003952 | III | 8243122 | 8244092 | -1 | Tsp_04234 | | 2533154 | GL622788 | 2534654 |
| caenorhabditis_elegans_prjna13758 | WBGene00007269 | III | 4091126 | 4093311 | 1 | Tsp_05227 | Ttbk1 | 2180440 | GL622792 | 2181339 |

| | | | | | | | | | | |
|---|---|---|---|---|---|---|---|---|---|---|
| caenorhabditis_elegans_prjna13758 | WBGene00007269 | III | 4091126 | 4093311 | 1 | Tsp_12209 | | 4940141 | GL622792 | 4941163 |
| caenorhabditis_elegans_prjna13758 | WBGene00007269 | III | 4091126 | 4093311 | 1 | Tsp_11208 | | 5439495 | GL622792 | 5440343 |
| caenorhabditis_elegans_prjna13758 | WBGene00007269 | III | 4091126 | 4093311 | 1 | Tsp_12401 | | 2248 | GL623394 | 3147 |
| caenorhabditis_elegans_prjna13758 | WBGene00007269 | III | 4091126 | 4093311 | 1 | Tsp_12402 | | 5826 | GL623394 | 6369 |
| caenorhabditis_elegans_prjna13758 | WBGene00007269 | III | 4091126 | 4093311 | 1 | Tsp_13461 | | 44 | GL626854 | 695 |
| caenorhabditis_elegans_prjna13758 | WBGene00004392 | III | 4093306 | 4094921 | -1 | Tsp_04092 | Rrm2 | 1929339 | GL622788 | 1930804 |
| caenorhabditis_elegans_prjna13758 | WBGene00000466 | I | 9665194 | 9668664 | 1 | Tsp_02609 | | 646372 | GL622785 | 649805 |
| caenorhabditis_elegans_prjna13758 | WBGene00007277 | I | 9658274 | 9660080 | 1 | Tsp_02412 | | 9731931 | GL622787 | 9733118 |
| caenorhabditis_elegans_prjna13758 | WBGene00001645 | V | 11282572 | 11286039 | 1 | Tsp_00241 | | 1019798 | GL622787 | 1022295 |
| caenorhabditis_elegans_prjna13758 | WBGene00007290 | X | 12283285 | 12285307 | 1 | Tsp_10949 | | 9477882 | GL622785 | 9479166 |
| caenorhabditis_elegans_prjna13758 | WBGene00003432 | IV | 10095398 | 10095851 | 1 | Tsp_00768 | | 3048065 | GL622787 | 3048698 |
| caenorhabditis_elegans_prjna13758 | WBGene00003432 | IV | 10095398 | 10095851 | 1 | Tsp_00761 | | 3076026 | GL622787 | 3077501 |
| caenorhabditis_elegans_prjna13758 | WBGene00003432 | IV | 10095398 | 10095851 | 1 | Tsp_00756 | | 2908432 | GL622787 | 2908941 |
| caenorhabditis_elegans_prjna13758 | WBGene00003432 | IV | 10095398 | 10095851 | 1 | Tsp_13425 | | 585 | GL623126 | 965 |
| caenorhabditis_elegans_prjna13758 | WBGene00003432 | IV | 10095398 | 10095851 | 1 | Tsp_13917 | | 820 | GL624354 | 1264 |
| caenorhabditis_elegans_prjna13758 | WBGene00001001 | IV | 10097196 | 10106556 | 1 | Tsp_05346 | | 2799977 | GL622792 | 2802923 |
| caenorhabditis_elegans_prjna13758 | WBGene00003043 | IV | 6842964 | 6845805 | 1 | Tsp_02993 | | 2249992 | GL622785 | 2250555 |
| caenorhabditis_elegans_prjna13758 | WBGene00003043 | IV | 6842964 | 6845805 | 1 | Tsp_12613 | | 1943 | GL622995 | 3807 |
| caenorhabditis_elegans_prjna13758 | WBGene00015461 | V | 7231190 | 7232749 | 1 | Tsp_09134 | | 3268911 | GL622784 | 3270436 |
| caenorhabditis_elegans_prjna13758 | WBGene00015461 | V | 7231190 | 7232749 | 1 | Tsp_09138.1 | | 3282445 | GL622784 | 3283294 |
| caenorhabditis_elegans_prjna13758 | WBGene00015462 | V | 7232434 | 7234826 | -1 | Tsp_09594 | | 5463128 | GL622789 | 5465428 |
| caenorhabditis_elegans_prjna13758 | WBGene00015486 | III | 6427529 | 6430934 | -1 | Tsp_00685 | | 2736947 | GL622787 | 2746227 |
| caenorhabditis_elegans_prjna13758 | WBGene00015467 | III | 5614972 | 5617257 | 1 | Tsp_00610 | Ddc | 2492021 | GL622787 | 2494539 |
| caenorhabditis_elegans_prjna13758 | WBGene00007352 | II | 10560018 | 10563407 | -1 | Tsp_09503 | | 755059 | GL622792 | 760398 |
| caenorhabditis_elegans_prjna13758 | WBGene00007354 | II | 10572247 | 10574067 | 1 | Tsp_06387 | | 10716569 | GL622787 | 10717856 |
| caenorhabditis_elegans_prjna13758 | WBGene00007354 | II | 10572247 | 10574067 | 1 | Tsp_13670 | | 166 | GL627984 | 1026 |

| | | | | | | | | | | |
|---|---|---|---|---|---|---|---|---|---|---|
| caenorhabditis_elegans_prjna13758 | WBGene00007354 | II | 10572247 | 10574067 | 1 | Tsp_11877 | | 139964 | GL622790 | 141286 |
| caenorhabditis_elegans_prjna13758 | WBGene00007354 | II | 10572247 | 10574067 | 1 | Tsp_13257 | | 551 | GL628954 | 1584 |
| caenorhabditis_elegans_prjna13758 | WBGene00007354 | II | 10572247 | 10574067 | 1 | Tsp_01419 | | 5756161 | GL622787 | 5757550 |
| caenorhabditis_elegans_prjna13758 | WBGene00007354 | II | 10572247 | 10574067 | 1 | Tsp_06318.1 | | 10514671 | GL622787 | 10515097 |
| caenorhabditis_elegans_prjna13758 | WBGene00007354 | II | 10572247 | 10574067 | 1 | Tsp_12521 | | 1192 | GL627057 | 2036 |
| caenorhabditis_elegans_prjna13758 | WBGene00007354 | II | 10572247 | 10574067 | 1 | Tsp_06319.2 | | 10512081 | GL622787 | 10512342 |
| caenorhabditis_elegans_prjna13758 | WBGene00007354 | II | 10572247 | 10574067 | 1 | Tsp_13571 | | 383 | GL628396 | 1344 |
| caenorhabditis_elegans_prjna13758 | WBGene00007355 | II | 10579366 | 10579955 | 1 | Tsp_02387 | | 9835641 | GL622787 | 9837057 |
| caenorhabditis_elegans_prjna13758 | WBGene00015516 | II | 7772499 | 7773914 | 1 | Tsp_06375 | Ppp1r7 | 10759467 | GL622787 | 10760411 |
| caenorhabditis_elegans_prjna13758 | WBGene00002228 | IV | 7040211 | 7043694 | 1 | Tsp_01664 | | 6573483 | GL622787 | 6580525 |
| caenorhabditis_elegans_prjna13758 | WBGene00007400 | III | 9694668 | 9696517 | -1 | Tsp_01774 | | 7169067 | GL622787 | 7169886 |
| caenorhabditis_elegans_prjna13758 | WBGene00007413 | II | 10339692 | 10345112 | -1 | Tsp_01959 | | 8025472 | GL622787 | 8028622 |
| caenorhabditis_elegans_prjna13758 | WBGene00007413 | II | 10339692 | 10345112 | -1 | Tsp_01951 | | 7973980 | GL622787 | 7977765 |
| caenorhabditis_elegans_prjna13758 | WBGene00003961 | IV | 8176479 | 8188879 | 1 | Tsp_03730 | | 258841 | GL622788 | 260503 |
| caenorhabditis_elegans_prjna13758 | WBGene00015581 | III | 7499677 | 7503241 | 1 | Tsp_08098 | | 1850745 | GL622789 | 1854313 |
| caenorhabditis_elegans_prjna13758 | WBGene00000479 | III | 7495320 | 7497832 | -1 | Tsp_11234 | | 27730 | GL624139 | 30106 |
| caenorhabditis_elegans_prjna13758 | WBGene00004723 | II | 8034250 | 8035767 | -1 | Tsp_02373 | | 9881075 | GL622787 | 9883145 |
| caenorhabditis_elegans_prjna13758 | WBGene00007434 | II | 8035869 | 8037346 | -1 | Tsp_14182 | | 50 | GL628616 | 421 |
| caenorhabditis_elegans_prjna13758 | WBGene00007434 | II | 8035869 | 8037346 | -1 | Tsp_04580 | ACTR6 | 405642 | GL624340 | 406844 |
| caenorhabditis_elegans_prjna13758 | WBGene00007435 | II | 8040006 | 8041732 | 1 | Tsp_07541 | | 1728064 | GL622784 | 1729981 |
| caenorhabditis_elegans_prjna13758 | WBGene00007428 | V | 10130405 | 10134148 | -1 | Tsp_04413 | | 3360270 | GL622788 | 3362330 |
| caenorhabditis_elegans_prjna13758 | WBGene00000142 | V | 10127492 | 10130295 | -1 | Tsp_00060 | | 250336 | GL622787 | 253752 |
| caenorhabditis_elegans_prjna13758 | WBGene00007444 | IV | 11149189 | 11154334 | 1 | Tsp_02851 | | 1603517 | GL622785 | 1605942 |
| caenorhabditis_elegans_prjna13758 | WBGene00007448 | IV | 11162768 | 11164622 | -1 | Tsp_02051 | | 8391493 | GL622787 | 8393242 |
| caenorhabditis_elegans_prjna13758 | WBGene00007448 | IV | 11162768 | 11164622 | -1 | Tsp_06549 | | 11192741 | GL622787 | 11195348 |
| caenorhabditis_elegans_prjna13758 | WBGene00007448 | IV | 11162768 | 11164622 | -1 | Tsp_00662 | Ttbk2 | 2539389 | GL622787 | 2540670 |

| | | | | | | | | | | |
|---|---|---|---|---|---|---|---|---|---|---|
| caenorhabditis_elegans_prjna13758 | WBGene00007448 | IV | 11162768 | 11164622 | -1 | Tsp_04341 | Ttbk1 | 3010978 | GL622788 | 3012100 |
| caenorhabditis_elegans_prjna13758 | WBGene00007448 | IV | 11162768 | 11164622 | -1 | Tsp_06889.2 | | 8369982 | GL622785 | 8372542 |
| caenorhabditis_elegans_prjna13758 | WBGene00003448 | IV | 5055593 | 5056049 | -1 | Tsp_00768 | | 3048065 | GL622787 | 3048698 |
| caenorhabditis_elegans_prjna13758 | WBGene00003448 | IV | 5055593 | 5056049 | -1 | Tsp_00761 | | 3076026 | GL622787 | 3077501 |
| caenorhabditis_elegans_prjna13758 | WBGene00003448 | IV | 5055593 | 5056049 | -1 | Tsp_00756 | | 2908432 | GL622787 | 2908941 |
| caenorhabditis_elegans_prjna13758 | WBGene00003448 | IV | 5055593 | 5056049 | -1 | Tsp_13425 | | 585 | GL623126 | 965 |
| caenorhabditis_elegans_prjna13758 | WBGene00003448 | IV | 5055593 | 5056049 | -1 | Tsp_13917 | | 820 | GL624354 | 1264 |
| caenorhabditis_elegans_prjna13758 | WBGene00001051 | IV | 8513919 | 8514725 | 1 | Tsp_07202 | | 442421 | GL622784 | 442780 |
| caenorhabditis_elegans_prjna13758 | WBGene00003864 | IV | 8888976 | 8890999 | 1 | Tsp_12414 | | 8096 | GL624002 | 8753 |
| caenorhabditis_elegans_prjna13758 | WBGene00003864 | IV | 8888976 | 8890999 | 1 | Tsp_08560 | zfp36l2 | 4699442 | GL622788 | 4700099 |
| caenorhabditis_elegans_prjna13758 | WBGene00003864 | IV | 8888976 | 8890999 | 1 | Tsp_08559 | Zfp36l3 | 4695882 | GL622788 | 4696511 |
| caenorhabditis_elegans_prjna13758 | WBGene00003864 | IV | 8888976 | 8890999 | 1 | Tsp_12415 | | 11512 | GL624002 | 12169 |
| caenorhabditis_elegans_prjna13758 | WBGene00003864 | IV | 8888976 | 8890999 | 1 | Tsp_12413.2 | | 5119 | GL624002 | 5437 |
| caenorhabditis_elegans_prjna13758 | WBGene00003864 | IV | 8888976 | 8890999 | 1 | Tsp_08561 | | 4702854 | GL622788 | 4703511 |
| caenorhabditis_elegans_prjna13758 | WBGene00015702 | IV | 6155492 | 6156960 | -1 | Tsp_06051 | | 6454142 | GL622785 | 6456063 |
| caenorhabditis_elegans_prjna13758 | WBGene00006856 | II | 14485624 | 14488688 | -1 | Tsp_04252 | USP14 | 2615257 | GL622788 | 2618051 |
| caenorhabditis_elegans_prjna13758 | WBGene00007555 | II | 10579955 | 10581045 | 1 | Tsp_00421 | | 1613198 | GL622787 | 1614897 |
| caenorhabditis_elegans_prjna13758 | WBGene00007561 | II | 10605717 | 10607275 | -1 | Tsp_10672 | | 2721132 | GL622789 | 2721788 |
| caenorhabditis_elegans_prjna13758 | WBGene00006474 | III | 3705265 | 3706784 | 1 | Tsp_09085 | | 3079631 | GL622784 | 3081934 |
| caenorhabditis_elegans_prjna13758 | WBGene00007584 | V | 12588309 | 12589644 | 1 | Tsp_02716 | | 1067687 | GL622785 | 1074809 |
| caenorhabditis_elegans_prjna13758 | WBGene00007603 | I | 12213109 | 12215184 | 1 | Tsp_09827 | | 110653 | GL623393 | 115169 |
| caenorhabditis_elegans_prjna13758 | WBGene00003835 | V | 14416067 | 14417835 | -1 | Tsp_03999 | | 1419239 | GL622788 | 1422640 |
| caenorhabditis_elegans_prjna13758 | WBGene00003922 | V | 14421566 | 14422618 | -1 | Tsp_06994 | | 8866864 | GL622785 | 8868285 |
| caenorhabditis_elegans_prjna13758 | WBGene00007617 | V | 14423776 | 14425008 | -1 | Tsp_00837 | | 3539588 | GL622787 | 3542931 |
| caenorhabditis_elegans_prjna13758 | WBGene00015807 | III | 6389833 | 6390841 | 1 | Tsp_00458 | | 1488317 | GL622787 | 1491676 |
| caenorhabditis_elegans_prjna13758 | WBGene00002850 | III | 6383552 | 6389401 | 1 | Tsp_00218 | | 883551 | GL622787 | 892365 |

| | | | | | | | | | | |
|---|---|---|---|---|---|---|---|---|---|---|
| caenorhabditis_elegans_prjna13758 | WBGene00015813 | III | 6368042 | 6373904 | 1 | Tsp_07461 | | 1402624 | GL622784 | 1407615 |
| caenorhabditis_elegans_prjna13758 | WBGene00015813 | III | 6368042 | 6373904 | 1 | Tsp_07460 | | 1407617 | GL622784 | 1411127 |
| caenorhabditis_elegans_prjna13758 | WBGene00007622 | III | 4178783 | 4181438 | -1 | Tsp_00302 | slc25a40 | 1347682 | GL622787 | 1350630 |
| caenorhabditis_elegans_prjna13758 | WBGene00007623 | III | 4177532 | 4178664 | -1 | Tsp_00872 | C16C10.2 | 3407126 | GL622787 | 3408661 |
| caenorhabditis_elegans_prjna13758 | WBGene00007625 | III | 4171336 | 4172223 | 1 | Tsp_00407 | | 1651900 | GL622787 | 1652758 |
| caenorhabditis_elegans_prjna13758 | WBGene00007627 | III | 4166944 | 4168765 | -1 | Tsp_06474.1 | | 11164489 | GL622787 | 11165490 |
| caenorhabditis_elegans_prjna13758 | WBGene00007627 | III | 4166944 | 4168765 | -1 | Tsp_06465 | | 10872780 | GL622787 | 10875108 |
| caenorhabditis_elegans_prjna13758 | WBGene00007620 | I | 9715321 | 9728108 | -1 | Tsp_10203 | | 6548030 | GL622792 | 6553334 |
| caenorhabditis_elegans_prjna13758 | WBGene00007643 | I | 9416704 | 9418884 | -1 | Tsp_09446 | | 261775 | GL622792 | 262882 |
| caenorhabditis_elegans_prjna13758 | WBGene00003904 | I | 9420907 | 9421992 | -1 | Tsp_05477 | | 3588400 | GL622792 | 3591433 |
| caenorhabditis_elegans_prjna13758 | WBGene00003904 | I | 9420907 | 9421992 | -1 | Tsp_06460 | | 10887662 | GL622787 | 10895079 |
| caenorhabditis_elegans_prjna13758 | WBGene00015916 | II | 5594714 | 5596463 | 1 | Tsp_01896 | | 7780804 | GL622787 | 7782635 |
| caenorhabditis_elegans_prjna13758 | WBGene00015924 | X | 8181031 | 8181640 | -1 | Tsp_03988 | arih1l | 1378658 | GL622788 | 1380369 |
| caenorhabditis_elegans_prjna13758 | WBGene00015941 | II | 5716307 | 5718038 | 1 | Tsp_01503 | | 5926545 | GL622787 | 5928283 |
| caenorhabditis_elegans_prjna13758 | WBGene00015941 | II | 5716307 | 5718038 | 1 | Tsp_14896 | | 896 | GL623976 | 1443 |
| caenorhabditis_elegans_prjna13758 | WBGene00015974 | I | 4193241 | 4203303 | 1 | Tsp_07793 | | 10664838 | GL622785 | 10667739 |
| caenorhabditis_elegans_prjna13758 | WBGene00015975 | I | 4190142 | 4193129 | 1 | Tsp_04087 | | 1917334 | GL622788 | 1919052 |
| caenorhabditis_elegans_prjna13758 | WBGene00007686 | II | 8971965 | 8973566 | -1 | Tsp_01518 | | 6144799 | GL622787 | 6147126 |
| caenorhabditis_elegans_prjna13758 | WBGene00016014 | III | 6195984 | 6198414 | -1 | Tsp_06632 | | 11585804 | GL622787 | 11589512 |
| caenorhabditis_elegans_prjna13758 | WBGene00016015 | III | 6202619 | 6206211 | -1 | Tsp_09877 | | 286531 | GL623393 | 290269 |
| caenorhabditis_elegans_prjna13758 | WBGene00016015 | III | 6202619 | 6206211 | -1 | Tsp_00836 | | 3543110 | GL622787 | 3548373 |
| caenorhabditis_elegans_prjna13758 | WBGene00016015 | III | 6202619 | 6206211 | -1 | Tsp_13268 | | 606 | GL627345 | 2400 |
| caenorhabditis_elegans_prjna13758 | WBGene00016062 | V | 5534131 | 5535837 | -1 | Tsp_02906 | | 1861679 | GL622785 | 1862687 |
| caenorhabditis_elegans_prjna13758 | WBGene00007699 | III | 11787690 | 11789496 | 1 | Tsp_01125 | | 4353662 | GL622787 | 4354442 |
| caenorhabditis_elegans_prjna13758 | WBGene00007700 | III | 11790497 | 11792295 | -1 | Tsp_01125 | | 4353662 | GL622787 | 4354442 |
| caenorhabditis_elegans_prjna13758 | WBGene00007711 | I | 10191273 | 10192894 | -1 | Tsp_06765 | ABHD4 | 7795671 | GL622785 | 7797598 |

| | | | | | | | | | | |
|---|---|---|---|---|---|---|---|---|---|---|
| caenorhabditis_elegans_prjna13758 | WBGene00006447 | I | 10159982 | 10161883 | -1 | Tsp_08271 | rnmt | 3577564 | GL622788 | 3578604 |
| caenorhabditis_elegans_prjna13758 | WBGene00045383 | I | 10169639 | 10172645 | -1 | Tsp_11602 | | 1878928 | GL622792 | 1883015 |
| caenorhabditis_elegans_prjna13758 | WBGene00007718 | V | 15066298 | 15067464 | 1 | Tsp_01660 | | 6592686 | GL622787 | 6593634 |
| caenorhabditis_elegans_prjna13758 | WBGene00016124 | IV | 8018130 | 8019808 | -1 | Tsp_04132 | | 2112017 | GL622788 | 2115828 |
| caenorhabditis_elegans_prjna13758 | WBGene00007042 | I | 7495437 | 7503577 | 1 | Tsp_06604 | | 11723760 | GL622787 | 11724725 |
| caenorhabditis_elegans_prjna13758 | WBGene00004304 | II | 8322612 | 8325052 | -1 | Tsp_06544 | | 11210634 | GL622787 | 11216207 |
| caenorhabditis_elegans_prjna13758 | WBGene00001840 | II | 8326852 | 8328870 | -1 | Tsp_09354 | | 5551190 | GL622788 | 5554956 |
| caenorhabditis_elegans_prjna13758 | WBGene00001840 | II | 8326852 | 8328870 | -1 | Tsp_12657 | | 7580 | GL625015 | 11111 |
| caenorhabditis_elegans_prjna13758 | WBGene00001499 | III | 4937322 | 4939453 | 1 | Tsp_03679 | fsn-1 | 95659 | GL622788 | 98014 |
| caenorhabditis_elegans_prjna13758 | WBGene00016142 | III | 4935578 | 4937223 | 1 | Tsp_05414 | | 3282267 | GL622792 | 3287835 |
| caenorhabditis_elegans_prjna13758 | WBGene00016151 | II | 5062129 | 5067116 | 1 | Tsp_08511 | | 4518222 | GL622788 | 4523362 |
| caenorhabditis_elegans_prjna13758 | WBGene00016151 | II | 5062129 | 5067116 | 1 | Tsp_13083 | | 2428 | GL629160 | 3225 |
| caenorhabditis_elegans_prjna13758 | WBGene00001102 | II | 5078235 | 5081161 | -1 | Tsp_08621 | | 3641980 | GL622789 | 3645765 |
| caenorhabditis_elegans_prjna13758 | WBGene00007763 | IV | 8903222 | 8906205 | -1 | Tsp_01125 | | 4353662 | GL622787 | 4354442 |
| caenorhabditis_elegans_prjna13758 | WBGene00001607 | IV | 9739219 | 9740908 | -1 | Tsp_04186 | | 2339558 | GL622788 | 2341969 |
| caenorhabditis_elegans_prjna13758 | WBGene00016192 | III | 5926086 | 5926987 | 1 | Tsp_01303 | C28H8.1 | 5127608 | GL622787 | 5128684 |
| caenorhabditis_elegans_prjna13758 | WBGene00016200 | III | 5903576 | 5906122 | -1 | Tsp_04288 | | 2767119 | GL622788 | 2770620 |
| caenorhabditis_elegans_prjna13758 | WBGene00004680 | II | 6125559 | 6127942 | 1 | Tsp_11858 | | 56958 | GL622790 | 61153 |
| caenorhabditis_elegans_prjna13758 | WBGene00016238 | III | 8225097 | 8226386 | 1 | Tsp_05146 | Mob4 | 2199381 | GL624340 | 2200742 |
| caenorhabditis_elegans_prjna13758 | WBGene00016239 | III | 8222226 | 8223792 | 1 | Tsp_06294 | | 10331217 | GL622787 | 10333834 |
| caenorhabditis_elegans_prjna13758 | WBGene00016245 | II | 6195813 | 6196959 | -1 | Tsp_08389 | | 4043238 | GL622788 | 4044742 |
| caenorhabditis_elegans_prjna13758 | WBGene00004460 | III | 8447353 | 8449346 | 1 | Tsp_00955 | Psmd3 | 3892714 | GL622787 | 3895439 |
| caenorhabditis_elegans_prjna13758 | WBGene00016250 | III | 8443917 | 8446920 | -1 | Tsp_00561 | | 2206125 | GL622787 | 2210532 |
| caenorhabditis_elegans_prjna13758 | WBGene00004244 | II | 7296110 | 7298247 | -1 | Tsp_04240 | | 2554050 | GL622788 | 2555842 |
| caenorhabditis_elegans_prjna13758 | WBGene00016323 | I | 3782118 | 3783294 | 1 | Tsp_05204 | | 2446915 | GL624340 | 2448276 |
| caenorhabditis_elegans_prjna13758 | WBGene00004217 | I | 3789309 | 3792797 | -1 | Tsp_01463 | | 6095018 | GL622787 | 6100354 |

| | | | | | | | | | | |
|---|---|---|---|---|---|---|---|---|---|---|
| caenorhabditis_elegans_prjna13758 | WBGene00004217 | I | 3789309 | 3792797 | -1 | Tsp_07377 | | 932107 | GL622784 | 944288 |
| caenorhabditis_elegans_prjna13758 | WBGene00004217 | I | 3789309 | 3792797 | -1 | Tsp_06808 | | 7963255 | GL622785 | 7966665 |
| caenorhabditis_elegans_prjna13758 | WBGene00004217 | I | 3789309 | 3792797 | -1 | Tsp_02133 | | 8604279 | GL622787 | 8606182 |
| caenorhabditis_elegans_prjna13758 | WBGene00001973 | I | 5815609 | 5818132 | -1 | Tsp_07406 | | 1229278 | GL622784 | 1234117 |
| caenorhabditis_elegans_prjna13758 | WBGene00016358 | II | 4836304 | 4837759 | -1 | Tsp_02716 | | 1067687 | GL622785 | 1074809 |
| caenorhabditis_elegans_prjna13758 | WBGene00016352 | II | 4828560 | 4832469 | 1 | Tsp_01646 | | 6625315 | GL622787 | 6629865 |
| caenorhabditis_elegans_prjna13758 | WBGene00016352 | II | 4828560 | 4832469 | 1 | Tsp_15670 | | 242 | GL623305 | 859 |
| caenorhabditis_elegans_prjna13758 | WBGene00003435 | II | 4825380 | 4825837 | 1 | Tsp_00768 | | 3048065 | GL622787 | 3048698 |
| caenorhabditis_elegans_prjna13758 | WBGene00003435 | II | 4825380 | 4825837 | 1 | Tsp_00761 | | 3076026 | GL622787 | 3077501 |
| caenorhabditis_elegans_prjna13758 | WBGene00003435 | II | 4825380 | 4825837 | 1 | Tsp_00756 | | 2908432 | GL622787 | 2908941 |
| caenorhabditis_elegans_prjna13758 | WBGene00003435 | II | 4825380 | 4825837 | 1 | Tsp_13425 | | 585 | GL623126 | 965 |
| caenorhabditis_elegans_prjna13758 | WBGene00003435 | II | 4825380 | 4825837 | 1 | Tsp_13917 | | 820 | GL624354 | 1264 |
| caenorhabditis_elegans_prjna13758 | WBGene00016380 | IV | 7792349 | 7794809 | -1 | Tsp_01927 | | 7869736 | GL622787 | 7873897 |
| caenorhabditis_elegans_prjna13758 | WBGene00016380 | IV | 7792349 | 7794809 | -1 | Tsp_01928 | | 7867951 | GL622787 | 7869171 |
| caenorhabditis_elegans_prjna13758 | WBGene00016373 | IV | 7776025 | 7778022 | 1 | Tsp_12380 | | 8229 | GL628879 | 9959 |
| caenorhabditis_elegans_prjna13758 | WBGene00016373 | IV | 7776025 | 7778022 | 1 | Tsp_02441 | | 9639288 | GL622787 | 9641840 |
| caenorhabditis_elegans_prjna13758 | WBGene00016373 | IV | 7776025 | 7778022 | 1 | Tsp_12379 | | 4716 | GL628879 | 6017 |
| caenorhabditis_elegans_prjna13758 | WBGene00016374 | IV | 7774411 | 7775956 | 1 | Tsp_02440 | | 9642505 | GL622787 | 9644648 |
| caenorhabditis_elegans_prjna13758 | WBGene00016374 | IV | 7774411 | 7775956 | 1 | Tsp_10182 | WDR82 | 6456671 | GL622792 | 6458439 |
| caenorhabditis_elegans_prjna13758 | WBGene00016374 | IV | 7774411 | 7775956 | 1 | Tsp_08028 | wdr82 | 1461714 | GL622789 | 1463096 |
| caenorhabditis_elegans_prjna13758 | WBGene00016388 | I | 10672737 | 10674864 | 1 | Tsp_02051 | | 8391493 | GL622787 | 8393242 |
| caenorhabditis_elegans_prjna13758 | WBGene00016388 | I | 10672737 | 10674864 | 1 | Tsp_06549 | | 11192741 | GL622787 | 11195348 |
| caenorhabditis_elegans_prjna13758 | WBGene00016388 | I | 10672737 | 10674864 | 1 | Tsp_00662 | Ttbk2 | 2539389 | GL622787 | 2540670 |
| caenorhabditis_elegans_prjna13758 | WBGene00016388 | I | 10672737 | 10674864 | 1 | Tsp_04341 | Ttbk1 | 3010978 | GL622788 | 3012100 |
| caenorhabditis_elegans_prjna13758 | WBGene00016388 | I | 10672737 | 10674864 | 1 | Tsp_06889.2 | | 8369982 | GL622785 | 8372542 |
| caenorhabditis_elegans_prjna13758 | WBGene00016392 | I | 10683270 | 10685664 | -1 | Tsp_06974 | | 8734959 | GL622785 | 8739582 |

| | | | | | | | | | | |
|---|---|---|---|---|---|---|---|---|---|---|
| caenorhabditis_elegans_prjna13758 | WBGene00000888 | IV | 7132132 | 7133217 | -1 | Tsp_11356 | Ppil1 | 7175329 | GL622792 | 7176101 |
| caenorhabditis_elegans_prjna13758 | WBGene00016408 | III | 5230555 | 5234530 | 1 | Tsp_08085 | | 1807874 | GL622789 | 1812874 |
| caenorhabditis_elegans_prjna13758 | WBGene00016408 | III | 5230555 | 5234530 | 1 | Tsp_08087 | | 1813695 | GL622789 | 1815636 |
| caenorhabditis_elegans_prjna13758 | WBGene00003443 | II | 5199763 | 5200223 | 1 | Tsp_00768 | | 3048065 | GL622787 | 3048698 |
| caenorhabditis_elegans_prjna13758 | WBGene00003443 | II | 5199763 | 5200223 | 1 | Tsp_00761 | | 3076026 | GL622787 | 3077501 |
| caenorhabditis_elegans_prjna13758 | WBGene00003443 | II | 5199763 | 5200223 | 1 | Tsp_00756 | | 2908432 | GL622787 | 2908941 |
| caenorhabditis_elegans_prjna13758 | WBGene00003443 | II | 5199763 | 5200223 | 1 | Tsp_13425 | | 585 | GL623126 | 965 |
| caenorhabditis_elegans_prjna13758 | WBGene00003443 | II | 5199763 | 5200223 | 1 | Tsp_13917 | | 820 | GL624354 | 1264 |
| caenorhabditis_elegans_prjna13758 | WBGene00016416 | II | 5195804 | 5199257 | 1 | Tsp_08852 | | 4870180 | GL622789 | 4874051 |
| caenorhabditis_elegans_prjna13758 | WBGene00016416 | II | 5195804 | 5199257 | 1 | Tsp_08682 | | 3923843 | GL622789 | 3926450 |
| caenorhabditis_elegans_prjna13758 | WBGene00016416 | II | 5195804 | 5199257 | 1 | Tsp_08683 | | 3927517 | GL622789 | 3929569 |
| caenorhabditis_elegans_prjna13758 | WBGene00016421 | I | 5872174 | 5874343 | 1 | Tsp_08585 | | 3412775 | GL622789 | 3414893 |
| caenorhabditis_elegans_prjna13758 | WBGene00016446 | III | 4860669 | 4861794 | -1 | Tsp_02096 | | 8768127 | GL622787 | 8769622 |
| caenorhabditis_elegans_prjna13758 | WBGene00000418 | III | 4852313 | 4855177 | 1 | Tsp_01495 | | 5944733 | GL622787 | 5951134 |
| caenorhabditis_elegans_prjna13758 | WBGene00007965 | III | 3841262 | 3843614 | -1 | Tsp_05595.2 | | 4125734 | GL622792 | 4125949 |
| caenorhabditis_elegans_prjna13758 | WBGene00007979 | I | 8752236 | 8755678 | 1 | Tsp_08733 | | 4178521 | GL622789 | 4180786 |
| caenorhabditis_elegans_prjna13758 | WBGene00007971 | I | 8728687 | 8730054 | 1 | Tsp_11760 | | 5042635 | GL622789 | 5043652 |
| caenorhabditis_elegans_prjna13758 | WBGene00007971 | I | 8728687 | 8730054 | 1 | Tsp_09380 | | 748 | GL622792 | 5462 |
| caenorhabditis_elegans_prjna13758 | WBGene00003925 | I | 8727214 | 8728497 | -1 | Tsp_09781 | psma7 | 6273678 | GL622789 | 6276581 |
| caenorhabditis_elegans_prjna13758 | WBGene00007972 | I | 8733413 | 8735717 | -1 | Tsp_03946 | | 1182003 | GL622788 | 1186580 |
| caenorhabditis_elegans_prjna13758 | WBGene00016508 | V | 4827360 | 4836025 | 1 | Tsp_01458 | | 5870508 | GL622787 | 5876278 |
| caenorhabditis_elegans_prjna13758 | WBGene00003912 | III | 4804708 | 4809489 | -1 | Tsp_00888 | | 3332437 | GL622787 | 3336228 |
| caenorhabditis_elegans_prjna13758 | WBGene00008034 | IV | 13091187 | 13093853 | -1 | Tsp_02448 | PRCC | 9619085 | GL622787 | 9619940 |
| caenorhabditis_elegans_prjna13758 | WBGene00004339 | IV | 13095762 | 13097065 | 1 | Tsp_10836 | RFC3 | 110448 | GL623868 | 111895 |
| caenorhabditis_elegans_prjna13758 | WBGene00016541 | IV | 5631895 | 5633379 | 1 | Tsp_02051 | | 8391493 | GL622787 | 8393242 |
| caenorhabditis_elegans_prjna13758 | WBGene00016541 | IV | 5631895 | 5633379 | 1 | Tsp_06549 | | 11192741 | GL622787 | 11195348 |

| | | | | | | | | | | |
|---|---|---|---|---|---|---|---|---|---|---|
| caenorhabditis_elegans_prjna13758 | WBGene00016541 | IV | 5631895 | 5633379 | 1 | Tsp_00662 | Ttbk2 | 2539389 | GL622787 | 2540670 |
| caenorhabditis_elegans_prjna13758 | WBGene00016541 | IV | 5631895 | 5633379 | 1 | Tsp_04341 | Ttbk1 | 3010978 | GL622788 | 3012100 |
| caenorhabditis_elegans_prjna13758 | WBGene00016541 | IV | 5631895 | 5633379 | 1 | Tsp_06889.2 | | 8369982 | GL622785 | 8372542 |
| caenorhabditis_elegans_prjna13758 | WBGene00008053 | II | 8137880 | 8141276 | 1 | Tsp_09503 | | 755059 | GL622792 | 760398 |
| caenorhabditis_elegans_prjna13758 | WBGene00016592 | IV | 12287928 | 12289392 | 1 | Tsp_03442 | | 4267278 | GL622785 | 4268827 |
| caenorhabditis_elegans_prjna13758 | WBGene00016586 | IV | 12291046 | 12293317 | 1 | Tsp_02920 | | 1923099 | GL622785 | 1924691 |
| caenorhabditis_elegans_prjna13758 | WBGene00016601 | I | 4270697 | 4273332 | 1 | Tsp_09565 | | 1018674 | GL622792 | 1025486 |
| caenorhabditis_elegans_prjna13758 | WBGene00000382 | I | 4268070 | 4270287 | -1 | Tsp_03181 | | 3139679 | GL622785 | 3141619 |
| caenorhabditis_elegans_prjna13758 | WBGene00016602 | I | 4265109 | 4267238 | 1 | Tsp_04894 | | 1389752 | GL624340 | 1392007 |
| caenorhabditis_elegans_prjna13758 | WBGene00007001 | I | 4247754 | 4249415 | 1 | Tsp_07198 | | 452845 | GL622784 | 454391 |
| caenorhabditis_elegans_prjna13758 | WBGene00016630 | II | 6888553 | 6890418 | -1 | Tsp_05518 | | 3789879 | GL622792 | 3792123 |
| caenorhabditis_elegans_prjna13758 | WBGene00016652 | I | 4637432 | 4640882 | -1 | Tsp_04835 | | 1208323 | GL624340 | 1209451 |
| caenorhabditis_elegans_prjna13758 | WBGene00016653 | I | 4634103 | 4635620 | 1 | Tsp_01639 | | 6662197 | GL622787 | 6666292 |
| caenorhabditis_elegans_prjna13758 | WBGene00016653 | I | 4634103 | 4635620 | 1 | Tsp_01638 | | 6666339 | GL622787 | 6669011 |
| caenorhabditis_elegans_prjna13758 | WBGene00016654 | I | 4605655 | 4610202 | 1 | Tsp_09955 | | 512546 | GL623393 | 515010 |
| caenorhabditis_elegans_prjna13758 | WBGene00008107 | I | 9221834 | 9226390 | -1 | Tsp_06395 | | 10671288 | GL622787 | 10678487 |
| caenorhabditis_elegans_prjna13758 | WBGene00001590 | I | 9226905 | 9228184 | 1 | Tsp_07794 | | 10663549 | GL622785 | 10664675 |
| caenorhabditis_elegans_prjna13758 | WBGene00001827 | IV | 7770770 | 7774275 | -1 | Tsp_06154 | | 7076076 | GL622785 | 7080215 |
| caenorhabditis_elegans_prjna13758 | WBGene00016704 | X | 4637242 | 4643695 | 1 | Tsp_02264 | | 9164081 | GL622787 | 9168128 |
| caenorhabditis_elegans_prjna13758 | WBGene00008122 | IV | 13715906 | 13716379 | -1 | Tsp_05810 | | 5414347 | GL622785 | 5418315 |
| caenorhabditis_elegans_prjna13758 | WBGene00003948 | I | 12967216 | 12968280 | -1 | Tsp_09780 | | 6276721 | GL622789 | 6278023 |
| caenorhabditis_elegans_prjna13758 | WBGene00001234 | I | 12968921 | 12970238 | 1 | Tsp_07767 | eIF6 | 10620410 | GL622785 | 10621420 |
| caenorhabditis_elegans_prjna13758 | WBGene00008136 | II | 11676306 | 11679197 | 1 | Tsp_10156 | | 823308 | GL622789 | 825816 |
| caenorhabditis_elegans_prjna13758 | WBGene00008136 | II | 11676306 | 11679197 | 1 | Tsp_10155 | | 822423 | GL622789 | 823217 |
| caenorhabditis_elegans_prjna13758 | WBGene00008137 | II | 11679091 | 11681068 | -1 | Tsp_11550 | | 735716 | GL622790 | 741190 |
| caenorhabditis_elegans_prjna13758 | WBGene00008137 | II | 11679091 | 11681068 | -1 | Tsp_12889 | | 454 | GL626579 | 1719 |

| | | | | | | | | | | |
|---|---|---|---|---|---|---|---|---|---|---|
| caenorhabditis_elegans_prjna13758 | WBGene00008140 | II | 11696171 | 11705371 | 1 | Tsp_10891 | ERCC4 | 339563 | GL623868 | 343050 |
| caenorhabditis_elegans_prjna13758 | WBGene00008151 | IV | 9980724 | 9982276 | -1 | Tsp_07494 | | 1654442 | GL622784 | 1657778 |
| caenorhabditis_elegans_prjna13758 | WBGene00000915 | V | 14684918 | 14688543 | 1 | Tsp_03752 | | 367161 | GL622788 | 370645 |
| caenorhabditis_elegans_prjna13758 | WBGene00008165 | II | 11290843 | 11293606 | -1 | Tsp_12091 | lmbrd2 | 384749 | GL622791 | 387956 |
| caenorhabditis_elegans_prjna13758 | WBGene00016739 | IV | 7393221 | 7397555 | 1 | Tsp_07959 | | 1027152 | GL622789 | 1030721 |
| caenorhabditis_elegans_prjna13758 | WBGene00016739 | IV | 7393221 | 7397555 | 1 | Tsp_11958 | Slc20a1 | 5504283 | GL622792 | 5508416 |
| caenorhabditis_elegans_prjna13758 | WBGene00016750 | I | 6255035 | 6257791 | 1 | Tsp_00826 | | 3195493 | GL622787 | 3196776 |
| caenorhabditis_elegans_prjna13758 | WBGene00016750 | I | 6255035 | 6257791 | 1 | Tsp_00827 | | 3194050 | GL622787 | 3195371 |
| caenorhabditis_elegans_prjna13758 | WBGene00003059 | I | 6253701 | 6254947 | 1 | Tsp_02451 | | 9608276 | GL622787 | 9611679 |
| caenorhabditis_elegans_prjna13758 | WBGene00004187 | III | 8164373 | 8172702 | 1 | Tsp_01598 | | 6461911 | GL622787 | 6471744 |
| caenorhabditis_elegans_prjna13758 | WBGene00004187 | III | 8164373 | 8172702 | 1 | Tsp_01597 | | 6473846 | GL622787 | 6476539 |
| caenorhabditis_elegans_prjna13758 | WBGene00016837 | I | 3877063 | 3880747 | 1 | Tsp_03302 | | 3697203 | GL622785 | 3701072 |
| caenorhabditis_elegans_prjna13758 | WBGene00016844 | IV | 7728913 | 7730645 | 1 | Tsp_06006 | | 6212360 | GL622785 | 6214429 |
| caenorhabditis_elegans_prjna13758 | WBGene00016844 | IV | 7728913 | 7730645 | 1 | Tsp_06005 | | 6214612 | GL622785 | 6217088 |
| caenorhabditis_elegans_prjna13758 | WBGene00004917 | V | 11978518 | 11979970 | -1 | Tsp_05371 | | 3064819 | GL622792 | 3065388 |
| caenorhabditis_elegans_prjna13758 | WBGene00004501 | V | 11980502 | 11982450 | -1 | Tsp_09237 | | 5058306 | GL622788 | 5060511 |
| caenorhabditis_elegans_prjna13758 | WBGene00001834 | V | 14525389 | 14527411 | 1 | Tsp_11412 | | 7474289 | GL622792 | 7477169 |
| caenorhabditis_elegans_prjna13758 | WBGene00004337 | V | 14642739 | 14645979 | -1 | Tsp_06737 | | 7662853 | GL622785 | 7668574 |
| caenorhabditis_elegans_prjna13758 | WBGene00008316 | I | 8019578 | 8020901 | -1 | Tsp_11052 | | 2051118 | GL622784 | 2054649 |
| caenorhabditis_elegans_prjna13758 | WBGene00001599 | I | 6513928 | 6517507 | 1 | Tsp_12124 | | 535552 | GL622790 | 536415 |
| caenorhabditis_elegans_prjna13758 | WBGene00001599 | I | 6513928 | 6517507 | 1 | Tsp_15219 | | 119 | GL623636 | 662 |
| caenorhabditis_elegans_prjna13758 | WBGene00000937 | I | 6490716 | 6492727 | -1 | Tsp_09915 | dbr1 | 422293 | GL623393 | 423600 |
| caenorhabditis_elegans_prjna13758 | WBGene00007018 | I | 6492722 | 6494754 | -1 | Tsp_01368 | MED18 | 5570577 | GL622787 | 5571227 |
| caenorhabditis_elegans_prjna13758 | WBGene00016960 | II | 6601216 | 6603449 | 1 | Tsp_04103 | VPS33B | 1976725 | GL622788 | 1978601 |
| caenorhabditis_elegans_prjna13758 | WBGene00016960 | II | 6601216 | 6603449 | 1 | Tsp_13297 | | 9853 | GL624545 | 10311 |
| caenorhabditis_elegans_prjna13758 | WBGene00002045 | II | 6587577 | 6588679 | -1 | Tsp_04407 | btf3l4 | 3340677 | GL622788 | 3341156 |

| | | | | | | | | | | |
|---|---|---|---|---|---|---|---|---|---|---|
| caenorhabditis_elegans_prjna13758 | WBGene00002045 | II | 6587577 | 6588679 | -1 | Tsp_14188 | btf3l4 | 764 | GL623344 | 1243 |
| caenorhabditis_elegans_prjna13758 | WBGene00016979 | III | 6331732 | 6335829 | 1 | Tsp_01584 | | 6514059 | GL622787 | 6516157 |
| caenorhabditis_elegans_prjna13758 | WBGene00016979 | III | 6331732 | 6335829 | 1 | Tsp_01583 | | 6519313 | GL622787 | 6522386 |
| caenorhabditis_elegans_prjna13758 | WBGene00003927 | V | 5583161 | 5584343 | -1 | Tsp_13296.2 | | 8 | GL624545 | 1334 |
| caenorhabditis_elegans_prjna13758 | WBGene00003927 | V | 5583161 | 5584343 | -1 | Tsp_13296.1 | | 3253 | GL624545 | 5169 |
| caenorhabditis_elegans_prjna13758 | WBGene00003927 | V | 5583161 | 5584343 | -1 | Tsp_04102 | | 1965842 | GL622788 | 1975530 |
| caenorhabditis_elegans_prjna13758 | WBGene00016992 | V | 5595653 | 5596551 | -1 | Tsp_04522 | | 277411 | GL624340 | 279035 |
| caenorhabditis_elegans_prjna13758 | WBGene00017003 | I | 4580692 | 4583810 | -1 | Tsp_10250 | | 6750462 | GL622792 | 6759147 |
| caenorhabditis_elegans_prjna13758 | WBGene00017003 | I | 4580692 | 4583810 | -1 | Tsp_10251 | | 6759243 | GL622792 | 6759915 |
| caenorhabditis_elegans_prjna13758 | WBGene00008364 | IV | 8930442 | 8932791 | 1 | Tsp_00380 | | 1750203 | GL622787 | 1751447 |
| caenorhabditis_elegans_prjna13758 | WBGene00008364 | IV | 8930442 | 8932791 | 1 | Tsp_00381 | | 1746280 | GL622787 | 1749039 |
| caenorhabditis_elegans_prjna13758 | WBGene00008380 | V | 10804721 | 10806034 | -1 | Tsp_07315 | | 865644 | GL622784 | 869411 |
| caenorhabditis_elegans_prjna13758 | WBGene00006481 | V | 10806131 | 10807973 | -1 | Tsp_06269 | PLRG1 | 10169502 | GL622787 | 10171991 |
| caenorhabditis_elegans_prjna13758 | WBGene00003923 | V | 10770321 | 10771351 | 1 | Tsp_01907 | | 7843691 | GL622787 | 7844750 |
| caenorhabditis_elegans_prjna13758 | WBGene00008371 | V | 10771451 | 10772832 | 1 | Tsp_04513 | | 256229 | GL624340 | 257473 |
| caenorhabditis_elegans_prjna13758 | WBGene00008386 | I | 8491291 | 8495855 | 1 | Tsp_07418 | CDC5L | 1204211 | GL622784 | 1208389 |
| caenorhabditis_elegans_prjna13758 | WBGene00008400 | I | 7820831 | 7826433 | 1 | Tsp_13512 | | 1465 | GL626788 | 3218 |
| caenorhabditis_elegans_prjna13758 | WBGene00017044 | III | 8145015 | 8145974 | 1 | Tsp_05627 | mrpl-18 | 4244919 | GL622792 | 4245966 |
| caenorhabditis_elegans_prjna13758 | WBGene00008410 | V | 11827306 | 11829370 | 1 | Tsp_03160 | | 3060130 | GL622785 | 3065973 |
| caenorhabditis_elegans_prjna13758 | WBGene00017064 | I | 6625066 | 6626788 | -1 | Tsp_03970 | | 1299342 | GL622788 | 1300440 |
| caenorhabditis_elegans_prjna13758 | WBGene00017069 | IV | 8396493 | 8397950 | 1 | Tsp_05774 | | 5260531 | GL622785 | 5261385 |
| caenorhabditis_elegans_prjna13758 | WBGene00017071 | IV | 8366539 | 8375371 | 1 | Tsp_03950 | | 1215067 | GL622788 | 1217793 |
| caenorhabditis_elegans_prjna13758 | WBGene00017071 | IV | 8366539 | 8375371 | 1 | Tsp_10814 | | 31395 | GL623868 | 32801 |
| caenorhabditis_elegans_prjna13758 | WBGene00017071 | IV | 8366539 | 8375371 | 1 | Tsp_01057 | | 4310767 | GL622787 | 4313727 |
| caenorhabditis_elegans_prjna13758 | WBGene00005019 | IV | 8361089 | 8363370 | 1 | Tsp_07365 | UXS1 | 998848 | GL622784 | 1004783 |
| caenorhabditis_elegans_prjna13758 | WBGene00017085 | I | 4132500 | 4135239 | 1 | Tsp_12101 | | 4759167 | GL622788 | 4759659 |

| | | | | | | | | | | |
|---|---|---|---|---|---|---|---|---|---|---|
| caenorhabditis_elegans_prjna13758 | WBGene00008456 | II | 9588212 | 9589767 | -1 | Tsp_03649 | eral1 | 5100617 | GL622785 | 5102338 |
| caenorhabditis_elegans_prjna13758 | WBGene00008456 | II | 9588212 | 9589767 | -1 | Tsp_03641 | | 5070861 | GL622785 | 5072270 |
| caenorhabditis_elegans_prjna13758 | WBGene00006473 | II | 9591292 | 9592936 | 1 | Tsp_01719 | | 7098996 | GL622787 | 7101349 |
| caenorhabditis_elegans_prjna13758 | WBGene00008458 | II | 9596701 | 9597528 | 1 | Tsp_09539 | | 827685 | GL622792 | 830727 |
| caenorhabditis_elegans_prjna13758 | WBGene00004322 | III | 4052807 | 4057976 | -1 | Tsp_00046 | | 291103 | GL622787 | 292849 |
| caenorhabditis_elegans_prjna13758 | WBGene00004322 | III | 4052807 | 4057976 | -1 | Tsp_14514 | | 36 | GL625389 | 1122 |
| caenorhabditis_elegans_prjna13758 | WBGene00001944 | III | 4059647 | 4060172 | -1 | Tsp_07554 | | 1811190 | GL622784 | 1811549 |
| caenorhabditis_elegans_prjna13758 | WBGene00001944 | III | 4059647 | 4060172 | -1 | Tsp_13713 | | 1978 | GL627003 | 2179 |
| caenorhabditis_elegans_prjna13758 | WBGene00001944 | III | 4059647 | 4060172 | -1 | Tsp_14200 | | 502 | GL627296 | 861 |
| caenorhabditis_elegans_prjna13758 | WBGene00001944 | III | 4059647 | 4060172 | -1 | Tsp_15676 | | 26 | GL624861 | 358 |
| caenorhabditis_elegans_prjna13758 | WBGene00001944 | III | 4059647 | 4060172 | -1 | Tsp_14173 | | 746 | GL627080 | 1105 |
| caenorhabditis_elegans_prjna13758 | WBGene00001944 | III | 4059647 | 4060172 | -1 | Tsp_15409 | | 148 | GL628791 | 507 |
| caenorhabditis_elegans_prjna13758 | WBGene00001944 | III | 4059647 | 4060172 | -1 | Tsp_13682 | | 1156 | GL629101 | 1341 |
| caenorhabditis_elegans_prjna13758 | WBGene00001944 | III | 4059647 | 4060172 | -1 | Tsp_07127 | | 228155 | GL622784 | 228526 |
| caenorhabditis_elegans_prjna13758 | WBGene00003393 | II | 5399538 | 5405810 | 1 | Tsp_01590 | | 6493038 | GL622787 | 6497729 |
| caenorhabditis_elegans_prjna13758 | WBGene00017159 | III | 5878521 | 5879415 | 1 | Tsp_03399 | | 4080753 | GL622785 | 4081272 |
| caenorhabditis_elegans_prjna13758 | WBGene00001609 | III | 9092241 | 9099698 | 1 | Tsp_06168 | | 7229241 | GL622785 | 7239006 |
| caenorhabditis_elegans_prjna13758 | WBGene00004117 | I | 8416831 | 8425042 | 1 | Tsp_07206 | | 416135 | GL622784 | 424507 |
| caenorhabditis_elegans_prjna13758 | WBGene00004117 | I | 8416831 | 8425042 | 1 | Tsp_01129 | SIN3B | 4621350 | GL622787 | 4625674 |
| caenorhabditis_elegans_prjna13758 | WBGene00004117 | I | 8416831 | 8425042 | 1 | Tsp_12058 | | 4952407 | GL622792 | 4958010 |
| caenorhabditis_elegans_prjna13758 | WBGene00017210 | II | 2053919 | 2059072 | -1 | Tsp_06410 | ZCCHC9 | 11092044 | GL622787 | 11093416 |
| caenorhabditis_elegans_prjna13758 | WBGene00017237 | IV | 8682239 | 8686610 | -1 | Tsp_07893 | | 11197807 | GL622785 | 11200528 |
| caenorhabditis_elegans_prjna13758 | WBGene00002124 | X | 11294962 | 11296731 | -1 | Tsp_00312 | | 1305355 | GL622787 | 1307253 |
| caenorhabditis_elegans_prjna13758 | WBGene00017280 | II | 2565275 | 2575930 | 1 | Tsp_07711 | USP39 | 10249880 | GL622785 | 10251496 |
| caenorhabditis_elegans_prjna13758 | WBGene00003421 | IV | 13155228 | 13164056 | 1 | Tsp_05453 | | 3383721 | GL622792 | 3385351 |
| caenorhabditis_elegans_prjna13758 | WBGene00017300 | III | 5557344 | 5564837 | 1 | Tsp_08925 | | 2427381 | GL622784 | 2435787 |

| | | | | | | | | | | |
|---|---|---|---|---|---|---|---|---|---|---|
| caenorhabditis_elegans_prjna13758 | WBGene00017316 | V | 7188332 | 7193662 | -1 | Tsp_03137 | | 2930909 | GL622785 | 2933057 |
| caenorhabditis_elegans_prjna13758 | WBGene00008639 | II | 8144762 | 8147236 | 1 | Tsp_04815 | | 1149156 | GL624340 | 1151570 |
| caenorhabditis_elegans_prjna13758 | WBGene00001281 | II | 8159865 | 8162170 | -1 | Tsp_06453 | | 10917593 | GL622787 | 10919905 |
| caenorhabditis_elegans_prjna13758 | WBGene00004510 | II | 8162465 | 8168644 | 1 | Tsp_01517 | | 6147594 | GL622787 | 6152275 |
| caenorhabditis_elegans_prjna13758 | WBGene00003134 | III | 475454 | 482322 | 1 | Tsp_13280 | | 1755 | GL623464 | 3140 |
| caenorhabditis_elegans_prjna13758 | WBGene00003134 | III | 475454 | 482322 | 1 | Tsp_06918 | APC8 | 8476673 | GL622785 | 8478121 |
| caenorhabditis_elegans_prjna13758 | WBGene00003134 | III | 475454 | 482322 | 1 | Tsp_13279 | | 333 | GL623464 | 1722 |
| caenorhabditis_elegans_prjna13758 | WBGene00017347 | II | 7111978 | 7112982 | 1 | Tsp_04233 | | 2531948 | GL622788 | 2532675 |
| caenorhabditis_elegans_prjna13758 | WBGene00017347 | II | 7111978 | 7112982 | 1 | Tsp_13341 | | 3744 | GL628955 | 4656 |
| caenorhabditis_elegans_prjna13758 | WBGene00017349 | II | 7101218 | 7105030 | -1 | Tsp_11771 | | 5078226 | GL622789 | 5082133 |
| caenorhabditis_elegans_prjna13758 | WBGene00006497 | II | 4710277 | 4713690 | 1 | Tsp_03479 | | 4420915 | GL622785 | 4425684 |
| caenorhabditis_elegans_prjna13758 | WBGene00006817 | II | 4702069 | 4703219 | 1 | Tsp_02412 | | 9731931 | GL622787 | 9733118 |
| caenorhabditis_elegans_prjna13758 | WBGene00004737 | II | 4698721 | 4700994 | 1 | Tsp_10008 | | 17546 | GL622789 | 19771 |
| caenorhabditis_elegans_prjna13758 | WBGene00004737 | II | 4698721 | 4700994 | 1 | Tsp_10009 | | 20161 | GL622789 | 24646 |
| caenorhabditis_elegans_prjna13758 | WBGene00004461 | II | 4696419 | 4698427 | -1 | Tsp_10788 | | 3215559 | GL622789 | 3218916 |
| caenorhabditis_elegans_prjna13758 | WBGene00003803 | I | 10026188 | 10030058 | 1 | Tsp_06746 | | 7696941 | GL622785 | 7700732 |
| caenorhabditis_elegans_prjna13758 | WBGene00008665 | I | 10030174 | 10046760 | 1 | Tsp_01641 | | 6648533 | GL622787 | 6649795 |
| caenorhabditis_elegans_prjna13758 | WBGene00008683 | IV | 12084478 | 12085320 | 1 | Tsp_13057 | | 3232 | GL627484 | 3550 |
| caenorhabditis_elegans_prjna13758 | WBGene00008683 | IV | 12084478 | 12085320 | 1 | Tsp_12601 | | 2217 | GL625084 | 3045 |
| caenorhabditis_elegans_prjna13758 | WBGene00008683 | IV | 12084478 | 12085320 | 1 | Tsp_07705 | Sf3a2 | 10275043 | GL622785 | 10276553 |
| caenorhabditis_elegans_prjna13758 | WBGene00008684 | IV | 12089247 | 12092213 | -1 | Tsp_06575 | | 11851450 | GL622787 | 11853175 |
| caenorhabditis_elegans_prjna13758 | WBGene00008684 | IV | 12089247 | 12092213 | -1 | Tsp_06574 | Nudcd2 | 11853470 | GL622787 | 11854094 |
| caenorhabditis_elegans_prjna13758 | WBGene00008686 | IV | 12094372 | 12096692 | 1 | Tsp_06127 | | 6924891 | GL622785 | 6927569 |
| caenorhabditis_elegans_prjna13758 | WBGene00006406 | IV | 11570500 | 11578699 | 1 | Tsp_03919 | | 1086385 | GL622788 | 1087934 |
| caenorhabditis_elegans_prjna13758 | WBGene00008722 | IV | 11558737 | 11561891 | -1 | Tsp_13254 | | 39 | GL628818 | 2609 |
| caenorhabditis_elegans_prjna13758 | WBGene00008722 | IV | 11558737 | 11561891 | -1 | Tsp_02376 | pold2 | 9865544 | GL622787 | 9867840 |

| | | | | | | | | | | |
|---|---|---|---|---|---|---|---|---|---|---|
| caenorhabditis_elegans_prjna13758 | WBGene00008729 | IV | 10422610 | 10424635 | -1 | Tsp_06200 | | 7121182 | GL622785 | 7124403 |
| caenorhabditis_elegans_prjna13758 | WBGene00017435 | II | 6270370 | 6274005 | 1 | Tsp_00117 | | 409474 | GL622787 | 414216 |
| caenorhabditis_elegans_prjna13758 | WBGene00000274 | II | 13200165 | 13215701 | 1 | Tsp_00622 | | 2434038 | GL622787 | 2440613 |
| caenorhabditis_elegans_prjna13758 | WBGene00000144 | V | 10425362 | 10426399 | -1 | Tsp_11955 | | 5497175 | GL622792 | 5497997 |
| caenorhabditis_elegans_prjna13758 | WBGene00000144 | V | 10425362 | 10426399 | -1 | Tsp_11890 | | 5601643 | GL622792 | 5605498 |
| caenorhabditis_elegans_prjna13758 | WBGene00008877 | I | 9390829 | 9393095 | -1 | Tsp_12314 | | 4972 | GL629016 | 7462 |
| caenorhabditis_elegans_prjna13758 | WBGene00008877 | I | 9390829 | 9393095 | -1 | Tsp_05733 | | 5136740 | GL622785 | 5137657 |
| caenorhabditis_elegans_prjna13758 | WBGene00008877 | I | 9390829 | 9393095 | -1 | Tsp_12406.2 | | 1893 | GL623659 | 4067 |
| caenorhabditis_elegans_prjna13758 | WBGene00008877 | I | 9390829 | 9393095 | -1 | Tsp_12406.1 | | 1089 | GL623659 | 1582 |
| caenorhabditis_elegans_prjna13758 | WBGene00008887 | I | 8363954 | 8369917 | -1 | Tsp_03497 | | 4513535 | GL622785 | 4517189 |
| caenorhabditis_elegans_prjna13758 | WBGene00017532 | X | 4647526 | 4649197 | -1 | Tsp_07410 | SLC29A1 | 1218333 | GL622784 | 1221797 |
| caenorhabditis_elegans_prjna13758 | WBGene00017534 | V | 6107828 | 6114856 | 1 | Tsp_07622 | | 9924704 | GL622785 | 9928000 |
| caenorhabditis_elegans_prjna13758 | WBGene00008921 | V | 10961722 | 10965562 | -1 | Tsp_06946 | | 8591211 | GL622785 | 8595323 |
| caenorhabditis_elegans_prjna13758 | WBGene00008921 | V | 10961722 | 10965562 | -1 | Tsp_12651 | | 9284976 | GL622785 | 9288009 |
| caenorhabditis_elegans_prjna13758 | WBGene00008918 | V | 10956910 | 10958821 | -1 | Tsp_12474.1 | | 829 | GL625490 | 2017 |
| caenorhabditis_elegans_prjna13758 | WBGene00008918 | V | 10956910 | 10958821 | -1 | Tsp_09346 | | 5496482 | GL622788 | 5500792 |
| caenorhabditis_elegans_prjna13758 | WBGene00008918 | V | 10956910 | 10958821 | -1 | Tsp_12474.2 | | 3905 | GL625490 | 4269 |
| caenorhabditis_elegans_prjna13758 | WBGene00017546 | II | 7664582 | 7667110 | 1 | Tsp_07124.2 | | 151434 | GL622784 | 154416 |
| caenorhabditis_elegans_prjna13758 | WBGene00017546 | II | 7664582 | 7667110 | 1 | Tsp_07124.1 | | 167984 | GL622784 | 170125 |
| caenorhabditis_elegans_prjna13758 | WBGene00007050 | II | 6536380 | 6554846 | 1 | Tsp_09914 | | 411585 | GL623393 | 421962 |
| caenorhabditis_elegans_prjna13758 | WBGene00004738 | V | 12763945 | 12767933 | 1 | Tsp_06183 | | 7183340 | GL622785 | 7189114 |
| caenorhabditis_elegans_prjna13758 | WBGene00017605 | V | 7571950 | 7575715 | -1 | Tsp_03550 | | 4698106 | GL622785 | 4705469 |
| caenorhabditis_elegans_prjna13758 | WBGene00008976 | X | 14982140 | 14982804 | -1 | Tsp_09335 | | 5458150 | GL622788 | 5459221 |
| caenorhabditis_elegans_prjna13758 | WBGene00017642 | IV | 7948643 | 7953548 | 1 | Tsp_05363 | | 3087341 | GL622792 | 3098224 |
| caenorhabditis_elegans_prjna13758 | WBGene00003777 | I | 7926623 | 7934577 | -1 | Tsp_01842 | | 7505262 | GL622787 | 7520561 |
| caenorhabditis_elegans_prjna13758 | WBGene00017646 | III | 6589377 | 6592175 | -1 | Tsp_10942 | | 9447722 | GL622785 | 9449933 |

| | | | | | | | | | | |
|---|---|---|---|---|---|---|---|---|---|---|
| caenorhabditis_elegans_prjna13758 | WBGene00009006 | IV | 8725722 | 8728060 | 1 | Tsp_14771 | | 13 | GL628150 | 1434 |
| caenorhabditis_elegans_prjna13758 | WBGene00009006 | IV | 8725722 | 8728060 | 1 | Tsp_06157 | | 7063504 | GL622785 | 7068278 |
| caenorhabditis_elegans_prjna13758 | WBGene00009007 | IV | 8728139 | 8730510 | 1 | Tsp_04332 | | 2974022 | GL622788 | 2976272 |
| caenorhabditis_elegans_prjna13758 | WBGene00017673 | I | 4901918 | 4903264 | 1 | Tsp_01153 | icmt | 4502606 | GL622787 | 4505263 |
| caenorhabditis_elegans_prjna13758 | WBGene00017683 | II | 6097830 | 6099663 | 1 | Tsp_15201 | | 795 | GL626006 | 1163 |
| caenorhabditis_elegans_prjna13758 | WBGene00009035 | IV | 11410228 | 11413001 | 1 | Tsp_02495 | | 133746 | GL622785 | 138843 |
| caenorhabditis_elegans_prjna13758 | WBGene00001358 | II | 8444307 | 8445177 | 1 | Tsp_10876 | | 277277 | GL623868 | 277783 |
| caenorhabditis_elegans_prjna13758 | WBGene00006994 | II | 8457818 | 8464517 | 1 | Tsp_01109 | | 4426992 | GL622787 | 4430371 |
| caenorhabditis_elegans_prjna13758 | WBGene00002198 | I | 7075918 | 7079566 | -1 | Tsp_12784 | | 3362 | GL624279 | 5110 |
| caenorhabditis_elegans_prjna13758 | WBGene00002198 | I | 7075918 | 7079566 | -1 | Tsp_06641 | Fps85D | 11546255 | GL622787 | 11548147 |
| caenorhabditis_elegans_prjna13758 | WBGene00002198 | I | 7075918 | 7079566 | -1 | Tsp_11870 | | 113432 | GL622790 | 117458 |
| caenorhabditis_elegans_prjna13758 | WBGene00003815 | I | 7088030 | 7090062 | -1 | Tsp_06285 | | 10294625 | GL622787 | 10300413 |
| caenorhabditis_elegans_prjna13758 | WBGene00002125 | X | 5996156 | 5998243 | 1 | Tsp_00791 | inx | 3163819 | GL622787 | 3165960 |
| caenorhabditis_elegans_prjna13758 | WBGene00002125 | X | 5996156 | 5998243 | 1 | Tsp_01271 | unc-7 | 5023274 | GL622787 | 5024989 |
| caenorhabditis_elegans_prjna13758 | WBGene00006465 | V | 14447629 | 14449587 | 1 | Tsp_08562 | | 4705074 | GL622788 | 4707939 |
| caenorhabditis_elegans_prjna13758 | WBGene00006465 | V | 14447629 | 14449587 | 1 | Tsp_12506.1 | | 6445 | GL626364 | 8968 |
| caenorhabditis_elegans_prjna13758 | WBGene00009084 | V | 14449697 | 14452971 | 1 | Tsp_03554 | | 4710431 | GL622785 | 4714022 |
| caenorhabditis_elegans_prjna13758 | WBGene00009078 | IV | 9146337 | 9146869 | -1 | Tsp_06400 | | 11141020 | GL622787 | 11142227 |
| caenorhabditis_elegans_prjna13758 | WBGene00017738 | I | 2432570 | 2433755 | -1 | Tsp_10710 | | 2870115 | GL622789 | 2872807 |
| caenorhabditis_elegans_prjna13758 | WBGene00017742 | II | 43854 | 44981 | 1 | Tsp_00857 | Nfyc | 3472922 | GL622787 | 3475612 |
| caenorhabditis_elegans_prjna13758 | WBGene00017746 | II | 30061 | 31584 | -1 | Tsp_05332 | | 2840603 | GL622792 | 2841670 |
| caenorhabditis_elegans_prjna13758 | WBGene00004504 | II | 42181 | 46216 | -1 | Tsp_02519 | | 268193 | GL622785 | 271290 |
| caenorhabditis_elegans_prjna13758 | WBGene00004503 | III | 6489793 | 6491282 | 1 | Tsp_05241 | | 2327589 | GL622792 | 2329779 |
| caenorhabditis_elegans_prjna13758 | WBGene00017757 | III | 911261 | 912243 | 1 | Tsp_01392 | | 5845574 | GL622787 | 5848374 |
| caenorhabditis_elegans_prjna13758 | WBGene00017757 | III | 911261 | 912243 | 1 | Tsp_09562 | | 1033523 | GL622792 | 1036030 |
| caenorhabditis_elegans_prjna13758 | WBGene00009103 | V | 9580041 | 9582644 | -1 | Tsp_11259 | | 104837 | GL624139 | 107891 |

| | | | | | | | | | | |
|---|---|---|---|---|---|---|---|---|---|---|
| caenorhabditis_elegans_prjna13758 | WBGene00017797 | V | 8570311 | 8581397 | 1 | Tsp_00894 | | 3305668 | GL622787 | 3308802 |
| caenorhabditis_elegans_prjna13758 | WBGene00017797 | V | 8570311 | 8581397 | 1 | Tsp_00135 | | 333673 | GL622787 | 336825 |
| caenorhabditis_elegans_prjna13758 | WBGene00009124 | I | 10571954 | 10577135 | 1 | Tsp_03837 | | 736434 | GL622788 | 741493 |
| caenorhabditis_elegans_prjna13758 | WBGene00009118 | I | 10551972 | 10553035 | 1 | Tsp_07510 | | 1596784 | GL622784 | 1597125 |
| caenorhabditis_elegans_prjna13758 | WBGene00006720 | I | 10562968 | 10564914 | 1 | Tsp_03600 | | 4927475 | GL622785 | 4928458 |
| caenorhabditis_elegans_prjna13758 | WBGene00003926 | I | 10566350 | 10567300 | 1 | Tsp_05973 | | 6082260 | GL622785 | 6084946 |
| caenorhabditis_elegans_prjna13758 | WBGene00009131 | IV | 9930899 | 9932240 | -1 | Tsp_05742 | | 5205401 | GL622785 | 5206086 |
| caenorhabditis_elegans_prjna13758 | WBGene00009132 | IV | 9932235 | 9933879 | 1 | Tsp_06225 | | 10025740 | GL622787 | 10027485 |
| caenorhabditis_elegans_prjna13758 | WBGene00009141 | I | 7642270 | 7643328 | -1 | Tsp_00254 | | 974785 | GL622787 | 975670 |
| caenorhabditis_elegans_prjna13758 | WBGene00009145 | I | 7674734 | 7676127 | -1 | Tsp_03692 | | 146971 | GL622788 | 148736 |
| caenorhabditis_elegans_prjna13758 | WBGene00017817 | I | 6309114 | 6310781 | 1 | Tsp_01125 | | 4353662 | GL622787 | 4354442 |
| caenorhabditis_elegans_prjna13758 | WBGene00001606 | IV | 17272231 | 17273652 | -1 | Tsp_04186 | | 2339558 | GL622788 | 2341969 |
| caenorhabditis_elegans_prjna13758 | WBGene00002005 | IV | 17278959 | 17281330 | 1 | Tsp_06317 | | 10497528 | GL622787 | 10504226 |
| caenorhabditis_elegans_prjna13758 | WBGene00009163 | I | 9781338 | 9785722 | -1 | Tsp_06192 | | 7144005 | GL622785 | 7149790 |
| caenorhabditis_elegans_prjna13758 | WBGene00009160 | I | 9768836 | 9771068 | -1 | Tsp_12784 | | 3362 | GL624279 | 5110 |
| caenorhabditis_elegans_prjna13758 | WBGene00009160 | I | 9768836 | 9771068 | -1 | Tsp_06641 | Fps85D | 11546255 | GL622787 | 11548147 |
| caenorhabditis_elegans_prjna13758 | WBGene00009160 | I | 9768836 | 9771068 | -1 | Tsp_11870 | | 113432 | GL622790 | 117458 |
| caenorhabditis_elegans_prjna13758 | WBGene00006528 | I | 9785778 | 9787624 | -1 | Tsp_00847 | | 3510190 | GL622787 | 3512482 |
| caenorhabditis_elegans_prjna13758 | WBGene00006528 | I | 9785778 | 9787624 | -1 | Tsp_08041 | | 1530059 | GL622789 | 1533849 |
| caenorhabditis_elegans_prjna13758 | WBGene00006528 | I | 9785778 | 9787624 | -1 | Tsp_01451 | | 5594555 | GL622787 | 5596616 |
| caenorhabditis_elegans_prjna13758 | WBGene00006528 | I | 9785778 | 9787624 | -1 | Tsp_01450 | | 5597665 | GL622787 | 5599719 |
| caenorhabditis_elegans_prjna13758 | WBGene00002637 | V | 5826833 | 5832866 | 1 | Tsp_01163 | | 4736419 | GL622787 | 4746481 |
| caenorhabditis_elegans_prjna13758 | WBGene00017825 | III | 4912207 | 4914133 | 1 | Tsp_03313 | | 3747890 | GL622785 | 3749025 |
| caenorhabditis_elegans_prjna13758 | WBGene00004679 | III | 4914096 | 4917129 | -1 | Tsp_00564 | | 2185971 | GL622787 | 2192857 |
| caenorhabditis_elegans_prjna13758 | WBGene00017830 | III | 4917104 | 4918008 | -1 | Tsp_00326 | rpb-8 | 1256879 | GL622787 | 1257391 |
| caenorhabditis_elegans_prjna13758 | WBGene00003598 | III | 4894647 | 4899085 | -1 | Tsp_00130 | | 350877 | GL622787 | 354980 |

| | | | | | | | | | | |
|---|---|---|---|---|---|---|---|---|---|---|
| caenorhabditis_elegans_prjna13758 | WBGene00003424 | II | 4785861 | 4786316 | -1 | Tsp_00768 | | 3048065 | GL622787 | 3048698 |
| caenorhabditis_elegans_prjna13758 | WBGene00003424 | II | 4785861 | 4786316 | -1 | Tsp_00761 | | 3076026 | GL622787 | 3077501 |
| caenorhabditis_elegans_prjna13758 | WBGene00003424 | II | 4785861 | 4786316 | -1 | Tsp_00756 | | 2908432 | GL622787 | 2908941 |
| caenorhabditis_elegans_prjna13758 | WBGene00003424 | II | 4785861 | 4786316 | -1 | Tsp_13425 | | 585 | GL623126 | 965 |
| caenorhabditis_elegans_prjna13758 | WBGene00003424 | II | 4785861 | 4786316 | -1 | Tsp_13917 | | 820 | GL624354 | 1264 |
| caenorhabditis_elegans_prjna13758 | WBGene00017855 | I | 5421313 | 5428457 | -1 | Tsp_08374 | | 3994427 | GL622788 | 3996569 |
| caenorhabditis_elegans_prjna13758 | WBGene00017855 | I | 5421313 | 5428457 | -1 | Tsp_12801 | | 158 | GL625223 | 2853 |
| caenorhabditis_elegans_prjna13758 | WBGene00000773 | II | 8600435 | 8602479 | -1 | Tsp_09726 | | 5956901 | GL622789 | 5960109 |
| caenorhabditis_elegans_prjna13758 | WBGene00003079 | V | 15569935 | 15570742 | 1 | Tsp_12500 | | 7983 | GL626231 | 8261 |
| caenorhabditis_elegans_prjna13758 | WBGene00003079 | V | 15569935 | 15570742 | 1 | Tsp_07532 | | 1764319 | GL622784 | 1764597 |
| caenorhabditis_elegans_prjna13758 | WBGene00003079 | V | 15569935 | 15570742 | 1 | Tsp_12470 | | 11449 | GL625295 | 11637 |
| caenorhabditis_elegans_prjna13758 | WBGene00006446 | V | 15575498 | 15577443 | -1 | Tsp_03879 | | 913303 | GL622788 | 917445 |
| caenorhabditis_elegans_prjna13758 | WBGene00009238 | V | 10746548 | 10747337 | -1 | Tsp_03801 | selt2 | 591187 | GL622788 | 592064 |
| caenorhabditis_elegans_prjna13758 | WBGene00017919 | IV | 4665251 | 4667293 | 1 | Tsp_09375 | | 5650153 | GL622788 | 5651781 |
| caenorhabditis_elegans_prjna13758 | WBGene00004205 | IV | 4653293 | 4655848 | 1 | Tsp_09337 | | 5461164 | GL622788 | 5461819 |
| caenorhabditis_elegans_prjna13758 | WBGene00004205 | IV | 4653293 | 4655848 | 1 | Tsp_09336 | | 5459876 | GL622788 | 5461157 |
| caenorhabditis_elegans_prjna13758 | WBGene00004502 | V | 6016049 | 6017669 | -1 | Tsp_07726 | | 10378436 | GL622785 | 10383009 |
| caenorhabditis_elegans_prjna13758 | WBGene00009262 | I | 9492615 | 9494589 | -1 | Tsp_08677 | | 3898689 | GL622789 | 3900819 |
| caenorhabditis_elegans_prjna13758 | WBGene00009271 | I | 7826442 | 7831629 | -1 | Tsp_04399 | | 3312503 | GL622788 | 3316282 |
| caenorhabditis_elegans_prjna13758 | WBGene00009272 | I | 7835814 | 7836967 | 1 | Tsp_11578 | | 1748715 | GL622792 | 1750509 |
| caenorhabditis_elegans_prjna13758 | WBGene00006386 | I | 7848975 | 7853631 | 1 | Tsp_00348 | | 1898060 | GL622787 | 1902226 |
| caenorhabditis_elegans_prjna13758 | WBGene00009284 | I | 15036373 | 15040487 | 1 | Tsp_15629 | | 229 | GL626767 | 1104 |
| caenorhabditis_elegans_prjna13758 | WBGene00009284 | I | 15036373 | 15040487 | 1 | Tsp_07735 | | 10507826 | GL622785 | 10510690 |
| caenorhabditis_elegans_prjna13758 | WBGene00009289 | V | 20838865 | 20841582 | 1 | Tsp_06288 | EXOSC7 | 10285524 | GL622787 | 10286984 |
| caenorhabditis_elegans_prjna13758 | WBGene00004340 | III | 6972210 | 6973717 | 1 | Tsp_15369 | | 351 | GL628433 | 1381 |
| caenorhabditis_elegans_prjna13758 | WBGene00004340 | III | 6972210 | 6973717 | 1 | Tsp_04544 | | 327745 | GL624340 | 329249 |

| | | | | | | | | | |
|---|---|---|---|---|---|---|---|---|---|
| caenorhabditis_elegans_prjna13758 | WBGene00017951 | III | 6963620 | 6968130 | -1 | Tsp_13525 | 2749 | GL628822 | 3862 |
| caenorhabditis_elegans_prjna13758 | WBGene00017951 | III | 6963620 | 6968130 | -1 | Tsp_02196 | 9046158 | GL622787 | 9049562 |
| caenorhabditis_elegans_prjna13758 | WBGene00017951 | III | 6963620 | 6968130 | -1 | Tsp_02199 | 9034981 | GL622787 | 9039033 |
| caenorhabditis_elegans_prjna13758 | WBGene00017951 | III | 6963620 | 6968130 | -1 | Tsp_15940 | 1240 | GL626840 | 1362 |
| caenorhabditis_elegans_prjna13758 | WBGene00003078 | II | 7252230 | 7253222 | -1 | Tsp_08226 | 2551400 | GL622789 | 2553265 |
| caenorhabditis_elegans_prjna13758 | WBGene00009305 | I | 14839856 | 14843128 | 1 | Tsp_14744 | 18 | GL625327 | 970 |
| caenorhabditis_elegans_prjna13758 | WBGene00009305 | I | 14839856 | 14843128 | 1 | Tsp_04543 | 325949 | GL624340 | 327484 |
| caenorhabditis_elegans_prjna13758 | WBGene00009320 | IV | 9882132 | 9884383 | -1 | Tsp_15116 | 157 | GL623434 | 645 |
| caenorhabditis_elegans_prjna13758 | WBGene00009320 | IV | 9882132 | 9884383 | -1 | Tsp_02989 | 2238350 | GL622785 | 2240127 |
| caenorhabditis_elegans_prjna13758 | WBGene00003464 | IV | 9894596 | 9895038 | 1 | Tsp_00768 | 3048065 | GL622787 | 3048698 |
| caenorhabditis_elegans_prjna13758 | WBGene00003464 | IV | 9894596 | 9895038 | 1 | Tsp_00761 | 3076026 | GL622787 | 3077501 |
| caenorhabditis_elegans_prjna13758 | WBGene00003464 | IV | 9894596 | 9895038 | 1 | Tsp_00756 | 2908432 | GL622787 | 2908941 |
| caenorhabditis_elegans_prjna13758 | WBGene00003464 | IV | 9894596 | 9895038 | 1 | Tsp_13425 | 585 | GL623126 | 965 |
| caenorhabditis_elegans_prjna13758 | WBGene00003464 | IV | 9894596 | 9895038 | 1 | Tsp_13917 | 820 | GL624354 | 1264 |
| caenorhabditis_elegans_prjna13758 | WBGene00017988 | V | 4371807 | 4373811 | 1 | Tsp_00518 | 2087982 | GL622787 | 2090396 |
| caenorhabditis_elegans_prjna13758 | WBGene00017989 | IV | 7580597 | 7583726 | 1 | Tsp_04317 | 2886252 | GL622788 | 2889189 |
| caenorhabditis_elegans_prjna13758 | WBGene00009341 | I | 8971139 | 8973063 | 1 | Tsp_00088 | 522850 | GL622787 | 524091 |
| caenorhabditis_elegans_prjna13758 | WBGene00002957 | II | 11022370 | 11027909 | 1 | Tsp_05710 | 4875900 | GL622792 | 4880650 |
| caenorhabditis_elegans_prjna13758 | WBGene00018007 | I | 5866514 | 5868139 | -1 | Tsp_11820 | 1266762 | GL622792 | 1274356 |
| caenorhabditis_elegans_prjna13758 | WBGene00018011 | V | 300015 | 304713 | -1 | Tsp_03086 | 2665460 | GL622785 | 2668484 |
| caenorhabditis_elegans_prjna13758 | WBGene00018011 | V | 300015 | 304713 | -1 | Tsp_14384 | 225 | GL622820 | 992 |
| caenorhabditis_elegans_prjna13758 | WBGene00018014 | II | 5029942 | 5031712 | -1 | Tsp_07670 | 10142015 | GL622785 | 10142409 |
| caenorhabditis_elegans_prjna13758 | WBGene00001049 | I | 15017283 | 15023017 | -1 | Tsp_11487 | 427717 | GL622790 | 435286 |
| caenorhabditis_elegans_prjna13758 | WBGene00001049 | I | 15017283 | 15023017 | -1 | Tsp_00165 | 827512 | GL622787 | 832124 |
| caenorhabditis_elegans_prjna13758 | WBGene00001049 | I | 15017283 | 15023017 | -1 | Tsp_12342 | 480 | GL629576 | 4064 |
| caenorhabditis_elegans_prjna13758 | WBGene00009366 | I | 15023092 | 15027761 | 1 | Tsp_10409 | 667296 | GL622791 | 669373 |

| | | | | | | | | | | |
|---|---|---|---|---|---|---|---|---|---|---|
| caenorhabditis_elegans_prjna13758 | WBGene00009366 | I | 15023092 | 15027761 | 1 | Tsp_10410 | | 669398 | GL622791 | 670921 |
| caenorhabditis_elegans_prjna13758 | WBGene00009372 | III | 3729493 | 3735039 | -1 | Tsp_11153 | | 5082926 | GL622792 | 5085426 |
| caenorhabditis_elegans_prjna13758 | WBGene00018024 | X | 3807460 | 3816429 | 1 | Tsp_07362 | | 1011392 | GL622784 | 1017763 |
| caenorhabditis_elegans_prjna13758 | WBGene00018064 | IV | 4042575 | 4047786 | 1 | Tsp_04052 | | 1717923 | GL622788 | 1720646 |
| caenorhabditis_elegans_prjna13758 | WBGene00000206 | III | 4594543 | 4595944 | -1 | Tsp_08439 | | 4230250 | GL622788 | 4231393 |
| caenorhabditis_elegans_prjna13758 | WBGene00009445 | III | 4581795 | 4583893 | 1 | Tsp_11317 | | 266245 | GL624139 | 268344 |
| caenorhabditis_elegans_prjna13758 | WBGene00009445 | III | 4581795 | 4583893 | 1 | Tsp_12959 | | 226 | GL627483 | 1749 |
| caenorhabditis_elegans_prjna13758 | WBGene00009451 | I | 8804799 | 8808081 | -1 | Tsp_05507 | | 3710865 | GL622792 | 3714265 |
| caenorhabditis_elegans_prjna13758 | WBGene00009460 | I | 8828048 | 8838719 | -1 | Tsp_04055 | | 1732308 | GL622788 | 1744146 |
| caenorhabditis_elegans_prjna13758 | WBGene00009460 | I | 8828048 | 8838719 | -1 | Tsp_13089 | | 13 | GL624346 | 1544 |
| caenorhabditis_elegans_prjna13758 | WBGene00009460 | I | 8828048 | 8838719 | -1 | Tsp_04054 | | 1722974 | GL622788 | 1730499 |
| caenorhabditis_elegans_prjna13758 | WBGene00018090 | V | 9419292 | 9425762 | -1 | Tsp_03413 | | 4136756 | GL622785 | 4137814 |
| caenorhabditis_elegans_prjna13758 | WBGene00009492 | IV | 11050128 | 11052430 | 1 | Tsp_07545 | | 1711702 | GL622784 | 1712427 |
| caenorhabditis_elegans_prjna13758 | WBGene00003426 | IV | 5260554 | 5261010 | 1 | Tsp_00768 | | 3048065 | GL622787 | 3048698 |
| caenorhabditis_elegans_prjna13758 | WBGene00003426 | IV | 5260554 | 5261010 | 1 | Tsp_00761 | | 3076026 | GL622787 | 3077501 |
| caenorhabditis_elegans_prjna13758 | WBGene00003426 | IV | 5260554 | 5261010 | 1 | Tsp_00756 | | 2908432 | GL622787 | 2908941 |
| caenorhabditis_elegans_prjna13758 | WBGene00003426 | IV | 5260554 | 5261010 | 1 | Tsp_13425 | | 585 | GL623126 | 965 |
| caenorhabditis_elegans_prjna13758 | WBGene00003426 | IV | 5260554 | 5261010 | 1 | Tsp_13917 | | 820 | GL624354 | 1264 |
| caenorhabditis_elegans_prjna13758 | WBGene00018122 | IV | 5258685 | 5259985 | 1 | Tsp_02051 | | 8391493 | GL622787 | 8393242 |
| caenorhabditis_elegans_prjna13758 | WBGene00018122 | IV | 5258685 | 5259985 | 1 | Tsp_06549 | | 11192741 | GL622787 | 11195348 |
| caenorhabditis_elegans_prjna13758 | WBGene00018122 | IV | 5258685 | 5259985 | 1 | Tsp_00662 | Ttbk2 | 2539389 | GL622787 | 2540670 |
| caenorhabditis_elegans_prjna13758 | WBGene00018122 | IV | 5258685 | 5259985 | 1 | Tsp_04341 | Ttbk1 | 3010978 | GL622788 | 3012100 |
| caenorhabditis_elegans_prjna13758 | WBGene00018122 | IV | 5258685 | 5259985 | 1 | Tsp_06889.2 | | 8369982 | GL622785 | 8372542 |
| caenorhabditis_elegans_prjna13758 | WBGene00018123 | IV | 5257020 | 5258352 | 1 | Tsp_02051 | | 8391493 | GL622787 | 8393242 |
| caenorhabditis_elegans_prjna13758 | WBGene00018123 | IV | 5257020 | 5258352 | 1 | Tsp_06549 | | 11192741 | GL622787 | 11195348 |
| caenorhabditis_elegans_prjna13758 | WBGene00018123 | IV | 5257020 | 5258352 | 1 | Tsp_00662 | Ttbk2 | 2539389 | GL622787 | 2540670 |

| | | | | | | | | | | |
|---|---|---|---|---|---|---|---|---|---|---|
| caenorhabditis_elegans_prjna13758 | WBGene00018123 | IV | 5257020 | 5258352 | 1 | Tsp_04341 | Ttbk1 | 3010978 | GL622788 | 3012100 |
| caenorhabditis_elegans_prjna13758 | WBGene00018123 | IV | 5257020 | 5258352 | 1 | Tsp_06889.2 | | 8369982 | GL622785 | 8372542 |
| caenorhabditis_elegans_prjna13758 | WBGene00002169 | III | 6703993 | 6710084 | -1 | Tsp_01296 | | 5156433 | GL622787 | 5161344 |
| caenorhabditis_elegans_prjna13758 | WBGene00018152 | III | 7167838 | 7170990 | 1 | Tsp_03822 | | 670149 | GL622788 | 676252 |
| caenorhabditis_elegans_prjna13758 | WBGene00009507 | I | 10479283 | 10484969 | -1 | Tsp_01290 | | 5174096 | GL622787 | 5176570 |
| caenorhabditis_elegans_prjna13758 | WBGene00018156 | I | 6421978 | 6425794 | 1 | Tsp_04884 | | 1352039 | GL624340 | 1356937 |
| caenorhabditis_elegans_prjna13758 | WBGene00009513 | II | 11186028 | 11188080 | 1 | Tsp_11550 | | 735716 | GL622790 | 741190 |
| caenorhabditis_elegans_prjna13758 | WBGene00009513 | II | 11186028 | 11188080 | 1 | Tsp_12889 | | 454 | GL626579 | 1719 |
| caenorhabditis_elegans_prjna13758 | WBGene00001029 | IV | 6585871 | 6588186 | 1 | Tsp_10571 | | 6060079 | GL622792 | 6064007 |
| caenorhabditis_elegans_prjna13758 | WBGene00009549 | IV | 11852193 | 11853146 | 1 | Tsp_00259 | | 940785 | GL622787 | 941576 |
| caenorhabditis_elegans_prjna13758 | WBGene00009553 | I | 14791040 | 14794371 | 1 | Tsp_10446 | | 885846 | GL622791 | 888634 |
| caenorhabditis_elegans_prjna13758 | WBGene00009553 | I | 14791040 | 14794371 | 1 | Tsp_10052 | | 207235 | GL622789 | 209212 |
| caenorhabditis_elegans_prjna13758 | WBGene00009556 | I | 14775184 | 14776663 | -1 | Tsp_09733 | | 5932024 | GL622789 | 5932869 |
| caenorhabditis_elegans_prjna13758 | WBGene00009563 | I | 8664925 | 8666499 | 1 | Tsp_05593 | | 4131064 | GL622792 | 4134195 |
| caenorhabditis_elegans_prjna13758 | WBGene00009575 | II | 11127634 | 11129256 | 1 | Tsp_00848 | | 3507418 | GL622787 | 3509438 |
| caenorhabditis_elegans_prjna13758 | WBGene00003076 | II | 11126932 | 11127530 | 1 | Tsp_07966 | LSM1 | 1072097 | GL622789 | 1073170 |
| caenorhabditis_elegans_prjna13758 | WBGene00001424 | II | 6755136 | 6755965 | 1 | Tsp_05902 | | 5832694 | GL622785 | 5833548 |
| caenorhabditis_elegans_prjna13758 | WBGene00001424 | II | 6755136 | 6755965 | 1 | Tsp_05905 | | 5911946 | GL622785 | 5912800 |
| caenorhabditis_elegans_prjna13758 | WBGene00018301 | II | 6746704 | 6748586 | 1 | Tsp_02051 | | 8391493 | GL622787 | 8393242 |
| caenorhabditis_elegans_prjna13758 | WBGene00018301 | II | 6746704 | 6748586 | 1 | Tsp_06549 | | 11192741 | GL622787 | 11195348 |
| caenorhabditis_elegans_prjna13758 | WBGene00018301 | II | 6746704 | 6748586 | 1 | Tsp_00662 | Ttbk2 | 2539389 | GL622787 | 2540670 |
| caenorhabditis_elegans_prjna13758 | WBGene00018301 | II | 6746704 | 6748586 | 1 | Tsp_04341 | Ttbk1 | 3010978 | GL622788 | 3012100 |
| caenorhabditis_elegans_prjna13758 | WBGene00018301 | II | 6746704 | 6748586 | 1 | Tsp_06889.2 | | 8369982 | GL622785 | 8372542 |
| caenorhabditis_elegans_prjna13758 | WBGene00018328 | IV | 3334042 | 3336868 | -1 | Tsp_05800 | | 5447878 | GL622785 | 5448817 |
| caenorhabditis_elegans_prjna13758 | WBGene00018357 | IV | 8130561 | 8132213 | 1 | Tsp_04285 | | 2756117 | GL622788 | 2758674 |
| caenorhabditis_elegans_prjna13758 | WBGene00018359 | IV | 8119090 | 8121340 | 1 | Tsp_01125 | | 4353662 | GL622787 | 4354442 |

| | | | | | | | | | | |
|---|---|---|---|---|---|---|---|---|---|---|
| caenorhabditis_elegans_prjna13758 | WBGene00018362 | III | 783714 | 786092 | 1 | Tsp_00466 | | 1441954 | GL622787 | 1445723 |
| caenorhabditis_elegans_prjna13758 | WBGene00018371 | III | 8488344 | 8490347 | -1 | Tsp_09133 | | 3265854 | GL622784 | 3267896 |
| caenorhabditis_elegans_prjna13758 | WBGene00002276 | I | 9578526 | 9581267 | -1 | Tsp_11594 | | 1838874 | GL622792 | 1841692 |
| caenorhabditis_elegans_prjna13758 | WBGene00004798 | III | 10506106 | 10506801 | -1 | Tsp_11071 | HSPB1 | 2099620 | GL622784 | 2100114 |
| caenorhabditis_elegans_prjna13758 | WBGene00009672 | I | 8640508 | 8644399 | 1 | Tsp_03565 | | 4753109 | GL622785 | 4757778 |
| caenorhabditis_elegans_prjna13758 | WBGene00009668 | I | 8615388 | 8616420 | 1 | Tsp_03672 | cpsf5 | 80781 | GL622788 | 81446 |
| caenorhabditis_elegans_prjna13758 | WBGene00000777 | I | 8636123 | 8637971 | -1 | Tsp_11680 | | 1673192 | GL622792 | 1676515 |
| caenorhabditis_elegans_prjna13758 | WBGene00009711 | II | 8994636 | 8996269 | 1 | Tsp_02353 | | 9504876 | GL622787 | 9506760 |
| caenorhabditis_elegans_prjna13758 | WBGene00000271 | II | 7305861 | 7308738 | 1 | Tsp_00909 | Brf1 | 3649792 | GL622787 | 3652172 |
| caenorhabditis_elegans_prjna13758 | WBGene00000839 | II | 7302101 | 7305760 | 1 | Tsp_12317 | | 15021 | GL629088 | 15965 |
| caenorhabditis_elegans_prjna13758 | WBGene00000839 | II | 7302101 | 7305760 | 1 | Tsp_12318 | | 18017 | GL629088 | 21365 |
| caenorhabditis_elegans_prjna13758 | WBGene00004046 | IV | 7633360 | 7634657 | 1 | Tsp_09335 | | 5458150 | GL622788 | 5459221 |
| caenorhabditis_elegans_prjna13758 | WBGene00009769 | V | 9777863 | 9779518 | -1 | Tsp_07064 | | 9233699 | GL622785 | 9235737 |
| caenorhabditis_elegans_prjna13758 | WBGene00000473 | I | 5619569 | 5620668 | 1 | Tsp_05554 | | 3860189 | GL622792 | 3860985 |
| caenorhabditis_elegans_prjna13758 | WBGene00018609 | III | 5464381 | 5467992 | 1 | Tsp_04512 | | 255388 | GL624340 | 256188 |
| caenorhabditis_elegans_prjna13758 | WBGene00018620 | V | 628059 | 628623 | -1 | Tsp_14455 | | 12 | GL627789 | 993 |
| caenorhabditis_elegans_prjna13758 | WBGene00018620 | V | 628059 | 628623 | -1 | Tsp_08348 | | 3897196 | GL622788 | 3899177 |
| caenorhabditis_elegans_prjna13758 | WBGene00004463 | IV | 9316639 | 9318632 | 1 | Tsp_03448 | | 4286620 | GL622785 | 4292416 |
| caenorhabditis_elegans_prjna13758 | WBGene00018625 | I | 10936819 | 10940521 | 1 | Tsp_12048 | | 3290107 | GL622789 | 3292973 |
| caenorhabditis_elegans_prjna13758 | WBGene00018632 | I | 10933810 | 10935712 | -1 | Tsp_13923 | | 1145 | GL623265 | 2303 |
| caenorhabditis_elegans_prjna13758 | WBGene00018632 | I | 10933810 | 10935712 | -1 | Tsp_05647 | | 4323175 | GL622792 | 4324333 |
| caenorhabditis_elegans_prjna13758 | WBGene00009893 | IV | 13053297 | 13055154 | 1 | Tsp_01125 | | 4353662 | GL622787 | 4354442 |
| caenorhabditis_elegans_prjna13758 | WBGene00018635 | IV | 7557209 | 7558749 | 1 | Tsp_06912 | | 8462800 | GL622785 | 8464239 |
| caenorhabditis_elegans_prjna13758 | WBGene00009924 | IV | 14077441 | 14083559 | 1 | Tsp_02372 | | 9884200 | GL622787 | 9888244 |
| caenorhabditis_elegans_prjna13758 | WBGene00009922 | I | 8318347 | 8323987 | -1 | Tsp_05179 | DHX30 | 2312105 | GL624340 | 2317055 |
| caenorhabditis_elegans_prjna13758 | WBGene00004439 | I | 8329062 | 8329684 | -1 | Tsp_06633 | rpl-25.1 | 11584026 | GL622787 | 11585153 |

| | | | | | | | | | | |
|---|---|---|---|---|---|---|---|---|---|---|
| caenorhabditis_elegans_prjna13758 | WBGene00018678 | IV | 1959854 | 1963240 | 1 | Tsp_10827 | | 67053 | GL623868 | 68898 |
| caenorhabditis_elegans_prjna13758 | WBGene00018679 | IV | 1953579 | 1956704 | 1 | Tsp_00957 | | 3885991 | GL622787 | 3888629 |
| caenorhabditis_elegans_prjna13758 | WBGene00003391 | III | 5308600 | 5311140 | 1 | Tsp_04666 | | 638450 | GL624340 | 640659 |
| caenorhabditis_elegans_prjna13758 | WBGene00004078 | V | 8414085 | 8415254 | 1 | Tsp_12414 | | 8096 | GL624002 | 8753 |
| caenorhabditis_elegans_prjna13758 | WBGene00004078 | V | 8414085 | 8415254 | 1 | Tsp_08560 | zfp36l2 | 4699442 | GL622788 | 4700099 |
| caenorhabditis_elegans_prjna13758 | WBGene00004078 | V | 8414085 | 8415254 | 1 | Tsp_08559 | Zfp36l3 | 4695882 | GL622788 | 4696511 |
| caenorhabditis_elegans_prjna13758 | WBGene00004078 | V | 8414085 | 8415254 | 1 | Tsp_12415 | | 11512 | GL624002 | 12169 |
| caenorhabditis_elegans_prjna13758 | WBGene00004078 | V | 8414085 | 8415254 | 1 | Tsp_12413.2 | | 5119 | GL624002 | 5437 |
| caenorhabditis_elegans_prjna13758 | WBGene00004078 | V | 8414085 | 8415254 | 1 | Tsp_08561 | | 4702854 | GL622788 | 4703511 |
| caenorhabditis_elegans_prjna13758 | WBGene00018698 | V | 8411775 | 8413513 | -1 | Tsp_02924 | | 1933686 | GL622785 | 1936548 |
| caenorhabditis_elegans_prjna13758 | WBGene00009937 | I | 9901255 | 9902929 | 1 | Tsp_07065 | | 9238570 | GL622785 | 9239571 |
| caenorhabditis_elegans_prjna13758 | WBGene00003829 | III | 13341527 | 13342607 | 1 | Tsp_09025 | | 2847793 | GL622784 | 2850161 |
| caenorhabditis_elegans_prjna13758 | WBGene00001092 | III | 13342684 | 13343337 | 1 | Tsp_04254 | | 2628862 | GL622788 | 2630134 |
| caenorhabditis_elegans_prjna13758 | WBGene00003478 | III | 13347085 | 13355683 | 1 | Tsp_01469 | | 6056187 | GL622787 | 6062157 |
| caenorhabditis_elegans_prjna13758 | WBGene00018721 | III | 1945058 | 1950270 | -1 | Tsp_02614 | | 663101 | GL622785 | 665513 |
| caenorhabditis_elegans_prjna13758 | WBGene00003797 | I | 3817996 | 3821505 | 1 | Tsp_03459 | | 4322494 | GL622785 | 4323615 |
| caenorhabditis_elegans_prjna13758 | WBGene00003797 | I | 3817996 | 3821505 | 1 | Tsp_03460 | | 4323626 | GL622785 | 4325933 |
| caenorhabditis_elegans_prjna13758 | WBGene00009993 | V | 13622107 | 13624525 | -1 | Tsp_11884 | | 162113 | GL622790 | 165236 |
| caenorhabditis_elegans_prjna13758 | WBGene00009993 | V | 13622107 | 13624525 | -1 | Tsp_12449 | | 6865 | GL624672 | 11654 |
| caenorhabditis_elegans_prjna13758 | WBGene00009993 | V | 13622107 | 13624525 | -1 | Tsp_14350 | | 881 | GL625180 | 1261 |
| caenorhabditis_elegans_prjna13758 | WBGene00004962 | I | 118109 | 120870 | 1 | Tsp_12784 | | 3362 | GL624279 | 5110 |
| caenorhabditis_elegans_prjna13758 | WBGene00004962 | I | 118109 | 120870 | 1 | Tsp_06641 | Fps85D | 11546255 | GL622787 | 11548147 |
| caenorhabditis_elegans_prjna13758 | WBGene00004962 | I | 118109 | 120870 | 1 | Tsp_11870 | | 113432 | GL622790 | 117458 |
| caenorhabditis_elegans_prjna13758 | WBGene00018783 | II | 2220464 | 2223567 | -1 | Tsp_13505 | | 5245 | GL625767 | 6522 |
| caenorhabditis_elegans_prjna13758 | WBGene00018783 | II | 2220464 | 2223567 | -1 | Tsp_00354 | | 1878189 | GL622787 | 1880417 |
| caenorhabditis_elegans_prjna13758 | WBGene00005014 | III | 69073 | 69817 | -1 | Tsp_00705 | | 2665180 | GL622787 | 2666898 |

| | | | | | | | | | | |
|---|---|---|---|---|---|---|---|---|---|---|
| caenorhabditis_elegans_prjna13758 | WBGene00010042 | II | 8571085 | 8572904 | 1 | Tsp_03371 | | 3998760 | GL622785 | 4000485 |
| caenorhabditis_elegans_prjna13758 | WBGene00010042 | II | 8571085 | 8572904 | 1 | Tsp_13180 | | 98 | GL623602 | 2248 |
| caenorhabditis_elegans_prjna13758 | WBGene00010044 | II | 8578049 | 8581513 | 1 | Tsp_03018 | | 2340062 | GL622785 | 2342971 |
| caenorhabditis_elegans_prjna13758 | WBGene00010056 | II | 11578716 | 11582758 | 1 | Tsp_15102 | | 14 | GL623569 | 1058 |
| caenorhabditis_elegans_prjna13758 | WBGene00010061 | IV | 11333079 | 11337184 | -1 | Tsp_06531 | | 11268490 | GL622787 | 11275448 |
| caenorhabditis_elegans_prjna13758 | WBGene00010070 | II | 13507806 | 13516479 | 1 | Tsp_00149 | | 622785 | GL622787 | 625202 |
| caenorhabditis_elegans_prjna13758 | WBGene00010070 | II | 13507806 | 13516479 | 1 | Tsp_00147 | | 629010 | GL622787 | 632541 |
| caenorhabditis_elegans_prjna13758 | WBGene00010077 | V | 11770456 | 11773495 | 1 | Tsp_03773 | | 478107 | GL622788 | 481060 |
| caenorhabditis_elegans_prjna13758 | WBGene00018866 | I | 5349646 | 5353767 | -1 | Tsp_15260 | | 711 | GL624258 | 1270 |
| caenorhabditis_elegans_prjna13758 | WBGene00018866 | I | 5349646 | 5353767 | -1 | Tsp_06939.1 | | 8568267 | GL622785 | 8574128 |
| caenorhabditis_elegans_prjna13758 | WBGene00018866 | I | 5349646 | 5353767 | -1 | Tsp_06936 | | 8551277 | GL622785 | 8560464 |
| caenorhabditis_elegans_prjna13758 | WBGene00010089 | V | 13823052 | 13825222 | 1 | Tsp_04511 | | 249933 | GL624340 | 254935 |
| caenorhabditis_elegans_prjna13758 | WBGene00018898 | IV | 4360837 | 4386533 | 1 | Tsp_14647 | | 80 | GL627512 | 997 |
| caenorhabditis_elegans_prjna13758 | WBGene00018890 | I | 5646369 | 5649656 | -1 | Tsp_07031 | | 9076906 | GL622785 | 9079718 |
| caenorhabditis_elegans_prjna13758 | WBGene00018891 | I | 5649756 | 5653648 | -1 | Tsp_02374 | | 9876314 | GL622787 | 9880130 |
| caenorhabditis_elegans_prjna13758 | WBGene00018892 | I | 5653804 | 5655857 | -1 | Tsp_01477 | | 6029372 | GL622787 | 6030745 |
| caenorhabditis_elegans_prjna13758 | WBGene00018893 | I | 5655950 | 5658118 | -1 | Tsp_05703 | | 4901399 | GL622792 | 4903999 |
| caenorhabditis_elegans_prjna13758 | WBGene00018897 | I | 5669950 | 5673352 | -1 | Tsp_07648 | | 10026727 | GL622785 | 10027824 |
| caenorhabditis_elegans_prjna13758 | WBGene00018901 | IV | 7475041 | 7476215 | 1 | Tsp_01596 | | 6478192 | GL622787 | 6481504 |
| caenorhabditis_elegans_prjna13758 | WBGene00004044 | IV | 7470860 | 7473340 | -1 | Tsp_02561 | | 467619 | GL622785 | 471249 |
| caenorhabditis_elegans_prjna13758 | WBGene00018909 | I | 5182736 | 5186234 | 1 | Tsp_14543 | | 273 | GL623557 | 503 |
| caenorhabditis_elegans_prjna13758 | WBGene00018909 | I | 5182736 | 5186234 | 1 | Tsp_01025 | | 4154465 | GL622787 | 4156787 |
| caenorhabditis_elegans_prjna13758 | WBGene00000774 | III | 13266874 | 13268897 | -1 | Tsp_04334 | | 2986791 | GL622788 | 2987560 |
| caenorhabditis_elegans_prjna13758 | WBGene00000774 | III | 13266874 | 13268897 | -1 | Tsp_11546 | | 723221 | GL622790 | 725196 |
| caenorhabditis_elegans_prjna13758 | WBGene00000774 | III | 13266874 | 13268897 | -1 | Tsp_05173 | CSTF2 | 2287060 | GL624340 | 2289325 |
| caenorhabditis_elegans_prjna13758 | WBGene00000774 | III | 13266874 | 13268897 | -1 | Tsp_13276 | | 430 | GL623256 | 1901 |

| | | | | | | | | | | |
|---|---|---|---|---|---|---|---|---|---|---|
| caenorhabditis_elegans_prjna13758 | WBGene00018948 | III | 7311290 | 7313195 | 1 | Tsp_05610 | | 4300271 | GL622792 | 4301996 |
| caenorhabditis_elegans_prjna13758 | WBGene00018961 | II | 5474373 | 5475475 | 1 | Tsp_04008 | | 1450880 | GL622788 | 1452934 |
| caenorhabditis_elegans_prjna13758 | WBGene00018967 | III | 5589036 | 5591718 | -1 | Tsp_09577 | | 971683 | GL622792 | 975729 |
| caenorhabditis_elegans_prjna13758 | WBGene00018991 | III | 2855554 | 2865497 | 1 | Tsp_06519 | | 11325864 | GL622787 | 11327745 |
| caenorhabditis_elegans_prjna13758 | WBGene00004505 | I | 5741868 | 5743571 | -1 | Tsp_07392 | Psmc3 | 1144544 | GL622784 | 1147073 |
| caenorhabditis_elegans_prjna13758 | WBGene00018995 | I | 5754312 | 5760803 | -1 | Tsp_05380 | | 3020373 | GL622792 | 3022641 |
| caenorhabditis_elegans_prjna13758 | WBGene00018995 | I | 5754312 | 5760803 | -1 | Tsp_05381 | | 3017015 | GL622792 | 3019837 |
| caenorhabditis_elegans_prjna13758 | WBGene00018995 | I | 5754312 | 5760803 | -1 | Tsp_13206 | | 2538 | GL623735 | 4750 |
| caenorhabditis_elegans_prjna13758 | WBGene00019004 | I | 6558696 | 6562346 | 1 | Tsp_05200 | | 2434192 | GL624340 | 2438463 |
| caenorhabditis_elegans_prjna13758 | WBGene00019005 | I | 6548644 | 6550447 | -1 | Tsp_08616 | | 3591318 | GL622789 | 3592032 |
| caenorhabditis_elegans_prjna13758 | WBGene00004462 | III | 6959110 | 6962027 | -1 | Tsp_00047 | | 287363 | GL622787 | 289891 |
| caenorhabditis_elegans_prjna13758 | WBGene00010195 | II | 14524173 | 14525112 | 1 | Tsp_03716 | | 220783 | GL622788 | 222607 |
| caenorhabditis_elegans_prjna13758 | WBGene00006704 | III | 9630085 | 9631167 | -1 | Tsp_00934 | | 3970973 | GL622787 | 3971965 |
| caenorhabditis_elegans_prjna13758 | WBGene00006540 | III | 9626532 | 9629154 | -1 | Tsp_06382 | | 10728828 | GL622787 | 10731564 |
| caenorhabditis_elegans_prjna13758 | WBGene00003438 | II | 5156700 | 5157083 | -1 | Tsp_00768 | | 3048065 | GL622787 | 3048698 |
| caenorhabditis_elegans_prjna13758 | WBGene00003438 | II | 5156700 | 5157083 | -1 | Tsp_00761 | | 3076026 | GL622787 | 3077501 |
| caenorhabditis_elegans_prjna13758 | WBGene00003438 | II | 5156700 | 5157083 | -1 | Tsp_00756 | | 2908432 | GL622787 | 2908941 |
| caenorhabditis_elegans_prjna13758 | WBGene00003438 | II | 5156700 | 5157083 | -1 | Tsp_13425 | | 585 | GL623126 | 965 |
| caenorhabditis_elegans_prjna13758 | WBGene00003438 | II | 5156700 | 5157083 | -1 | Tsp_13917 | | 820 | GL624354 | 1264 |
| caenorhabditis_elegans_prjna13758 | WBGene00010231 | IV | 11628757 | 11631850 | -1 | Tsp_09293 | | 5317929 | GL622788 | 5321394 |
| caenorhabditis_elegans_prjna13758 | WBGene00010231 | IV | 11628757 | 11631850 | -1 | Tsp_09294 | | 5322024 | GL622788 | 5325072 |
| caenorhabditis_elegans_prjna13758 | WBGene00010232 | IV | 11636648 | 11638208 | 1 | Tsp_06042 | | 6498393 | GL622785 | 6501082 |
| caenorhabditis_elegans_prjna13758 | WBGene00010233 | IV | 11638316 | 11640097 | 1 | Tsp_11979 | RBM17 | 1170972 | GL622792 | 1172454 |
| caenorhabditis_elegans_prjna13758 | WBGene00010265 | II | 12925981 | 12928341 | -1 | Tsp_06387 | | 10716569 | GL622787 | 10717856 |
| caenorhabditis_elegans_prjna13758 | WBGene00010265 | II | 12925981 | 12928341 | -1 | Tsp_13670 | | 166 | GL627984 | 1026 |
| caenorhabditis_elegans_prjna13758 | WBGene00010265 | II | 12925981 | 12928341 | -1 | Tsp_11877 | | 139964 | GL622790 | 141286 |

| | | | | | | | | | | |
|---|---|---|---|---|---|---|---|---|---|---|
| caenorhabditis_elegans_prjna13758 | WBGene00010265 | II | 12925981 | 12928341 | -1 | Tsp_13257 | | 551 | GL628954 | 1584 |
| caenorhabditis_elegans_prjna13758 | WBGene00010265 | II | 12925981 | 12928341 | -1 | Tsp_01419 | | 5756161 | GL622787 | 5757550 |
| caenorhabditis_elegans_prjna13758 | WBGene00010265 | II | 12925981 | 12928341 | -1 | Tsp_06318.1 | | 10514671 | GL622787 | 10515097 |
| caenorhabditis_elegans_prjna13758 | WBGene00010265 | II | 12925981 | 12928341 | -1 | Tsp_12521 | | 1192 | GL627057 | 2036 |
| caenorhabditis_elegans_prjna13758 | WBGene00010265 | II | 12925981 | 12928341 | -1 | Tsp_06319.2 | | 10512081 | GL622787 | 10512342 |
| caenorhabditis_elegans_prjna13758 | WBGene00010265 | II | 12925981 | 12928341 | -1 | Tsp_13571 | | 383 | GL628396 | 1344 |
| caenorhabditis_elegans_prjna13758 | WBGene00006442 | V | 13675988 | 13678598 | -1 | Tsp_05629 | | 4238319 | GL622792 | 4241988 |
| caenorhabditis_elegans_prjna13758 | WBGene00010283 | V | 13678875 | 13681539 | 1 | Tsp_02024.1 | | 8254887 | GL622787 | 8255598 |
| caenorhabditis_elegans_prjna13758 | WBGene00003795 | III | 3400644 | 3404479 | -1 | Tsp_00378 | RGPD1 | 1756676 | GL622787 | 1761582 |
| caenorhabditis_elegans_prjna13758 | WBGene00010303 | III | 3396585 | 3397675 | 1 | Tsp_00845 | | 3514822 | GL622787 | 3515922 |
| caenorhabditis_elegans_prjna13758 | WBGene00010304 | III | 3397743 | 3400644 | 1 | Tsp_03638 | | 5057157 | GL622785 | 5060680 |
| caenorhabditis_elegans_prjna13758 | WBGene00010304 | III | 3397743 | 3400644 | 1 | Tsp_03622.2 | | 5018385 | GL622785 | 5020473 |
| caenorhabditis_elegans_prjna13758 | WBGene00019077 | I | 5507123 | 5509288 | -1 | Tsp_10418 | | 691906 | GL622791 | 695963 |
| caenorhabditis_elegans_prjna13758 | WBGene00010309 | III | 8999828 | 9001973 | -1 | Tsp_00047 | | 287363 | GL622787 | 289891 |
| caenorhabditis_elegans_prjna13758 | WBGene00003882 | II | 10873931 | 10877934 | 1 | Tsp_01471 | | 6050226 | GL622787 | 6051401 |
| caenorhabditis_elegans_prjna13758 | WBGene00004214 | II | 5918626 | 5921617 | 1 | Tsp_10590 | | 6094173 | GL622792 | 6098496 |
| caenorhabditis_elegans_prjna13758 | WBGene00019163 | IV | 4807430 | 4809487 | -1 | Tsp_03591 | | 4888229 | GL622785 | 4890150 |
| caenorhabditis_elegans_prjna13758 | WBGene00019219 | II | 4329023 | 4334484 | 1 | Tsp_01918 | | 7804220 | GL622787 | 7807598 |
| caenorhabditis_elegans_prjna13758 | WBGene00019220 | II | 4326720 | 4328284 | 1 | Tsp_05977 | | 6068453 | GL622785 | 6068833 |
| caenorhabditis_elegans_prjna13758 | WBGene00003390 | II | 4324361 | 4326089 | 1 | Tsp_09506 | | 741868 | GL622792 | 747766 |
| caenorhabditis_elegans_prjna13758 | WBGene00010420 | I | 12647376 | 12652087 | -1 | Tsp_07041 | | 9117232 | GL622785 | 9120198 |
| caenorhabditis_elegans_prjna13758 | WBGene00001352 | III | 4303096 | 4313128 | -1 | Tsp_11871 | | 117623 | GL622790 | 124444 |
| caenorhabditis_elegans_prjna13758 | WBGene00019275 | V | 4199485 | 4203696 | 1 | Tsp_06988 | | 8830202 | GL622785 | 8831589 |
| caenorhabditis_elegans_prjna13758 | WBGene00010436 | IV | 13243673 | 13244296 | 1 | Tsp_04322 | | 2900039 | GL622788 | 2902850 |
| caenorhabditis_elegans_prjna13758 | WBGene00004976 | III | 3439023 | 3444207 | -1 | Tsp_12037 | | 107023 | GL622786 | 113886 |
| caenorhabditis_elegans_prjna13758 | WBGene00004976 | III | 3439023 | 3444207 | -1 | Tsp_03836 | | 734725 | GL622788 | 735836 |

| | | | | | | | | | | |
|---|---|---|---|---|---|---|---|---|---|---|
| caenorhabditis_elegans_prjna13758 | WBGene00000379 | II | 8283667 | 8285711 | 1 | Tsp_08448 | | 4259107 | GL622788 | 4261513 |
| caenorhabditis_elegans_prjna13758 | WBGene00003821 | II | 8280855 | 8283208 | -1 | Tsp_13264 | | 280 | GL623603 | 3012 |
| caenorhabditis_elegans_prjna13758 | WBGene00003821 | II | 8280855 | 8283208 | -1 | Tsp_02671 | | 865708 | GL622785 | 866935 |
| caenorhabditis_elegans_prjna13758 | WBGene00004302 | III | 10746581 | 10747728 | -1 | Tsp_04091 | | 1927840 | GL622788 | 1928623 |
| caenorhabditis_elegans_prjna13758 | WBGene00010478 | III | 10748429 | 10750080 | 1 | Tsp_00865 | Nop60B | 3427074 | GL622787 | 3430649 |
| caenorhabditis_elegans_prjna13758 | WBGene00010485 | IV | 9708170 | 9709305 | -1 | Tsp_01983 | | 8105151 | GL622787 | 8107104 |
| caenorhabditis_elegans_prjna13758 | WBGene00010510 | V | 14232281 | 14236076 | -1 | Tsp_07410 | SLC29A1 | 1218333 | GL622784 | 1221797 |
| caenorhabditis_elegans_prjna13758 | WBGene00019323 | I | 6814511 | 6834687 | 1 | Tsp_05143 | | 2177446 | GL624340 | 2183897 |
| caenorhabditis_elegans_prjna13758 | WBGene00004388 | III | 852801 | 853909 | -1 | Tsp_15741 | | 417 | GL628032 | 1597 |
| caenorhabditis_elegans_prjna13758 | WBGene00004388 | III | 852801 | 853909 | -1 | Tsp_07086 | | 27419 | GL622784 | 29003 |
| caenorhabditis_elegans_prjna13758 | WBGene00019334 | III | 856560 | 860090 | 1 | Tsp_01640 | K02F3.12 | 6650027 | GL622787 | 6652417 |
| caenorhabditis_elegans_prjna13758 | WBGene00001603 | III | 9943243 | 9944823 | -1 | Tsp_04186 | | 2339558 | GL622788 | 2341969 |
| caenorhabditis_elegans_prjna13758 | WBGene00010551 | II | 14441409 | 14447219 | 1 | Tsp_09040 | | 2890172 | GL622784 | 2897954 |
| caenorhabditis_elegans_prjna13758 | WBGene00010551 | II | 14441409 | 14447219 | 1 | Tsp_12993.1 | | 4471 | GL622797 | 6233 |
| caenorhabditis_elegans_prjna13758 | WBGene00001979 | I | 14738882 | 14742657 | 1 | Tsp_14258 | | 154 | GL628835 | 1177 |
| caenorhabditis_elegans_prjna13758 | WBGene00001979 | I | 14738882 | 14742657 | 1 | Tsp_08140 | | 2086529 | GL622789 | 2089990 |
| caenorhabditis_elegans_prjna13758 | WBGene00003469 | II | 5797893 | 5798341 | 1 | Tsp_00768 | | 3048065 | GL622787 | 3048698 |
| caenorhabditis_elegans_prjna13758 | WBGene00003469 | II | 5797893 | 5798341 | 1 | Tsp_00761 | | 3076026 | GL622787 | 3077501 |
| caenorhabditis_elegans_prjna13758 | WBGene00003469 | II | 5797893 | 5798341 | 1 | Tsp_00756 | | 2908432 | GL622787 | 2908941 |
| caenorhabditis_elegans_prjna13758 | WBGene00003469 | II | 5797893 | 5798341 | 1 | Tsp_13425 | | 585 | GL623126 | 965 |
| caenorhabditis_elegans_prjna13758 | WBGene00003469 | II | 5797893 | 5798341 | 1 | Tsp_13917 | | 820 | GL624354 | 1264 |
| caenorhabditis_elegans_prjna13758 | WBGene00019406 | II | 5794495 | 5796185 | 1 | Tsp_04159 | | 2252924 | GL622788 | 2254844 |
| caenorhabditis_elegans_prjna13758 | WBGene00000386 | I | 6469386 | 6471995 | -1 | Tsp_13015.2 | | 3440 | GL624006 | 4762 |
| caenorhabditis_elegans_prjna13758 | WBGene00000386 | I | 6469386 | 6471995 | -1 | Tsp_13015.1 | | 4885 | GL624006 | 5857 |
| caenorhabditis_elegans_prjna13758 | WBGene00001743 | III | 8085705 | 8088449 | 1 | Tsp_07750 | | 10455066 | GL622785 | 10459060 |
| caenorhabditis_elegans_prjna13758 | WBGene00001743 | III | 8085705 | 8088449 | 1 | Tsp_07751 | | 10450077 | GL622785 | 10454827 |

| | | | | | | | | | | |
|---|---|---|---|---|---|---|---|---|---|---|
| caenorhabditis_elegans_prjna13758 | WBGene00004312 | I | 9616508 | 9618107 | -1 | Tsp_09779 | Caf1 | 6278927 | GL622789 | 6281525 |
| caenorhabditis_elegans_prjna13758 | WBGene00003036 | I | 9618207 | 9619760 | -1 | Tsp_05180 | | 2317375 | GL624340 | 2320982 |
| caenorhabditis_elegans_prjna13758 | WBGene00000098 | V | 8221410 | 8223587 | 1 | Tsp_11544 | | 715159 | GL622790 | 716583 |
| caenorhabditis_elegans_prjna13758 | WBGene00000098 | V | 8221410 | 8223587 | 1 | Tsp_07501 | | 1630867 | GL622784 | 1634897 |
| caenorhabditis_elegans_prjna13758 | WBGene00010627 | V | 10350471 | 10352978 | -1 | Tsp_01355 | | 5425299 | GL622787 | 5431142 |
| caenorhabditis_elegans_prjna13758 | WBGene00010629 | V | 10353584 | 10355837 | -1 | Tsp_12584 | | 126 | GL629434 | 472 |
| caenorhabditis_elegans_prjna13758 | WBGene00010631 | V | 10362261 | 10366014 | 1 | Tsp_07628 | | 10016715 | GL622785 | 10019021 |
| caenorhabditis_elegans_prjna13758 | WBGene00010631 | V | 10362261 | 10366014 | 1 | Tsp_12829 | | 86 | GL626847 | 4124 |
| caenorhabditis_elegans_prjna13758 | WBGene00010631 | V | 10362261 | 10366014 | 1 | Tsp_13376 | | 1642 | GL625564 | 3433 |
| caenorhabditis_elegans_prjna13758 | WBGene00010631 | V | 10362261 | 10366014 | 1 | Tsp_16111 | | 44 | GL625891 | 868 |
| caenorhabditis_elegans_prjna13758 | WBGene00004467 | II | 4043059 | 4044330 | -1 | Tsp_02191 | | 9067370 | GL622787 | 9073376 |
| caenorhabditis_elegans_prjna13758 | WBGene00019498 | II | 641010 | 650367 | -1 | Tsp_04053 | | 1720811 | GL622788 | 1722025 |
| caenorhabditis_elegans_prjna13758 | WBGene00003425 | IV | 9837616 | 9838065 | 1 | Tsp_00768 | | 3048065 | GL622787 | 3048698 |
| caenorhabditis_elegans_prjna13758 | WBGene00003425 | IV | 9837616 | 9838065 | 1 | Tsp_00761 | | 3076026 | GL622787 | 3077501 |
| caenorhabditis_elegans_prjna13758 | WBGene00003425 | IV | 9837616 | 9838065 | 1 | Tsp_00756 | | 2908432 | GL622787 | 2908941 |
| caenorhabditis_elegans_prjna13758 | WBGene00003425 | IV | 9837616 | 9838065 | 1 | Tsp_13425 | | 585 | GL623126 | 965 |
| caenorhabditis_elegans_prjna13758 | WBGene00003425 | IV | 9837616 | 9838065 | 1 | Tsp_13917 | | 820 | GL624354 | 1264 |
| caenorhabditis_elegans_prjna13758 | WBGene00003449 | IV | 9841443 | 9841961 | 1 | Tsp_00768 | | 3048065 | GL622787 | 3048698 |
| caenorhabditis_elegans_prjna13758 | WBGene00003449 | IV | 9841443 | 9841961 | 1 | Tsp_00761 | | 3076026 | GL622787 | 3077501 |
| caenorhabditis_elegans_prjna13758 | WBGene00003449 | IV | 9841443 | 9841961 | 1 | Tsp_00756 | | 2908432 | GL622787 | 2908941 |
| caenorhabditis_elegans_prjna13758 | WBGene00003449 | IV | 9841443 | 9841961 | 1 | Tsp_13425 | | 585 | GL623126 | 965 |
| caenorhabditis_elegans_prjna13758 | WBGene00003449 | IV | 9841443 | 9841961 | 1 | Tsp_13917 | | 820 | GL624354 | 1264 |
| caenorhabditis_elegans_prjna13758 | WBGene00002206 | IV | 9839057 | 9841084 | -1 | Tsp_12784 | | 3362 | GL624279 | 5110 |
| caenorhabditis_elegans_prjna13758 | WBGene00002206 | IV | 9839057 | 9841084 | -1 | Tsp_06641 | Fps85D | 11546255 | GL622787 | 11548147 |
| caenorhabditis_elegans_prjna13758 | WBGene00002206 | IV | 9839057 | 9841084 | -1 | Tsp_11870 | | 113432 | GL622790 | 117458 |
| caenorhabditis_elegans_prjna13758 | WBGene00006963 | I | 7159714 | 7160917 | 1 | Tsp_10304 | | 6956512 | GL622792 | 6958770 |

| | | | | | | | | | | |
|---|---|---|---|---|---|---|---|---|---|---|
| caenorhabditis_elegans_prjna13758 | WBGene00006963 | I | 7159714 | 7160917 | 1 | Tsp_10305 | | 6958884 | GL622792 | 6959468 |
| caenorhabditis_elegans_prjna13758 | WBGene00019503 | IV | 8294690 | 8297283 | 1 | Tsp_02571 | | 502140 | GL622785 | 505244 |
| caenorhabditis_elegans_prjna13758 | WBGene00019510 | IV | 8286587 | 8289531 | -1 | Tsp_06160 | | 7052599 | GL622785 | 7056347 |
| caenorhabditis_elegans_prjna13758 | WBGene00004386 | IV | 4174089 | 4176279 | 1 | Tsp_09888 | | 320650 | GL623393 | 321722 |
| caenorhabditis_elegans_prjna13758 | WBGene00004385 | IV | 4172076 | 4173988 | 1 | Tsp_09888 | | 320650 | GL623393 | 321722 |
| caenorhabditis_elegans_prjna13758 | WBGene00000875 | III | 13768541 | 13771242 | -1 | Tsp_00839 | | 3530850 | GL622787 | 3535555 |
| caenorhabditis_elegans_prjna13758 | WBGene00000875 | III | 13768541 | 13771242 | -1 | Tsp_00838 | | 3537055 | GL622787 | 3539471 |
| caenorhabditis_elegans_prjna13758 | WBGene00000875 | III | 13768541 | 13771242 | -1 | Tsp_13966 | | 308 | GL623889 | 616 |
| caenorhabditis_elegans_prjna13758 | WBGene00000875 | III | 13768541 | 13771242 | -1 | Tsp_02108 | | 8728385 | GL622787 | 8731257 |
| caenorhabditis_elegans_prjna13758 | WBGene00010670 | IV | 12077107 | 12078162 | -1 | Tsp_08838 | tmed1 | 4759576 | GL622789 | 4760415 |
| caenorhabditis_elegans_prjna13758 | WBGene00010670 | IV | 12077107 | 12078162 | -1 | Tsp_00224 | tmed1 | 861185 | GL622787 | 861871 |
| caenorhabditis_elegans_prjna13758 | WBGene00019543 | IV | 4706777 | 4710755 | -1 | Tsp_09339 | | 5463970 | GL622788 | 5465245 |
| caenorhabditis_elegans_prjna13758 | WBGene00010676 | IV | 10122120 | 10126675 | -1 | Tsp_07918 | | 11289600 | GL622785 | 11296617 |
| caenorhabditis_elegans_prjna13758 | WBGene00010685 | V | 15134188 | 15136801 | -1 | Tsp_12433 | | 10336 | GL624403 | 12537 |
| caenorhabditis_elegans_prjna13758 | WBGene00010685 | V | 15134188 | 15136801 | -1 | Tsp_11547 | | 725507 | GL622790 | 728534 |
| caenorhabditis_elegans_prjna13758 | WBGene00003062 | I | 3144410 | 3147793 | -1 | Tsp_06836 | | 8088205 | GL622785 | 8091458 |
| caenorhabditis_elegans_prjna13758 | WBGene00003009 | II | 6371295 | 6374544 | 1 | Tsp_05213 | | 2467064 | GL624340 | 2470007 |
| caenorhabditis_elegans_prjna13758 | WBGene00019608 | II | 6361159 | 6365367 | -1 | Tsp_13631 | | 243 | GL623607 | 1460 |
| caenorhabditis_elegans_prjna13758 | WBGene00019627 | III | 5191131 | 5196028 | 1 | Tsp_09906 | | 383359 | GL623393 | 387190 |
| caenorhabditis_elegans_prjna13758 | WBGene00019642 | V | 6896999 | 6898497 | 1 | Tsp_02051 | | 8391493 | GL622787 | 8393242 |
| caenorhabditis_elegans_prjna13758 | WBGene00019642 | V | 6896999 | 6898497 | 1 | Tsp_06549 | | 11192741 | GL622787 | 11195348 |
| caenorhabditis_elegans_prjna13758 | WBGene00019642 | V | 6896999 | 6898497 | 1 | Tsp_00662 | Ttbk2 | 2539389 | GL622787 | 2540670 |
| caenorhabditis_elegans_prjna13758 | WBGene00019642 | V | 6896999 | 6898497 | 1 | Tsp_04341 | Ttbk1 | 3010978 | GL622788 | 3012100 |
| caenorhabditis_elegans_prjna13758 | WBGene00019642 | V | 6896999 | 6898497 | 1 | Tsp_06889.2 | | 8369982 | GL622785 | 8372542 |
| caenorhabditis_elegans_prjna13758 | WBGene00002219 | III | 10805930 | 10810999 | 1 | Tsp_05539 | KIF2A | 3914776 | GL622792 | 3918307 |
| caenorhabditis_elegans_prjna13758 | WBGene00019667 | V | 6281359 | 6284863 | 1 | Tsp_15868 | | 366 | GL626428 | 1089 |

| | | | | | | | | | | |
|---|---|---|---|---|---|---|---|---|---|---|
| caenorhabditis_elegans_prjna13758 | WBGene00019667 | V | 6281359 | 6284863 | 1 | Tsp_06873 | | 8284855 | GL622785 | 8289393 |
| caenorhabditis_elegans_prjna13758 | WBGene00010785 | II | 11875145 | 11880588 | 1 | Tsp_00990 | | 3763911 | GL622787 | 3769982 |
| caenorhabditis_elegans_prjna13758 | WBGene00019678 | III | 8046034 | 8047612 | 1 | Tsp_04514 | brix1 | 257791 | GL624340 | 259110 |
| caenorhabditis_elegans_prjna13758 | WBGene00000939 | III | 8071916 | 8080281 | -1 | Tsp_00125 | | 371594 | GL622787 | 379675 |
| caenorhabditis_elegans_prjna13758 | WBGene00019692 | I | 5563827 | 5566265 | 1 | Tsp_03295 | | 3637445 | GL622785 | 3640633 |
| caenorhabditis_elegans_prjna13758 | WBGene00019710 | I | 5582005 | 5583471 | 1 | Tsp_01153 | icmt | 4502606 | GL622787 | 4505263 |
| caenorhabditis_elegans_prjna13758 | WBGene00019711 | I | 5579666 | 5581979 | 1 | Tsp_13994 | | 386 | GL628611 | 1208 |
| caenorhabditis_elegans_prjna13758 | WBGene00019711 | I | 5579666 | 5581979 | 1 | Tsp_03894 | | 962857 | GL622788 | 964935 |
| caenorhabditis_elegans_prjna13758 | WBGene00000474 | I | 5571787 | 5573019 | -1 | Tsp_05554 | | 3860189 | GL622792 | 3860985 |
| caenorhabditis_elegans_prjna13758 | WBGene00010839 | III | 10399174 | 10405240 | -1 | Tsp_00137 | | 326191 | GL622787 | 330540 |
| caenorhabditis_elegans_prjna13758 | WBGene00010839 | III | 10399174 | 10405240 | -1 | Tsp_16035 | | 32 | GL628115 | 884 |
| caenorhabditis_elegans_prjna13758 | WBGene00044069 | III | 10405248 | 10407248 | -1 | Tsp_06631 | | 11592279 | GL622787 | 11594851 |
| caenorhabditis_elegans_prjna13758 | WBGene00010844 | III | 10424112 | 10427982 | -1 | Tsp_07851 | | 10927633 | GL622785 | 10932264 |
| caenorhabditis_elegans_prjna13758 | WBGene00010845 | III | 10428334 | 10435988 | -1 | Tsp_00134 | | 339183 | GL622787 | 343527 |
| caenorhabditis_elegans_prjna13758 | WBGene00019762 | V | 5940024 | 5943362 | 1 | Tsp_09327 | CRNKL1 | 5429731 | GL622788 | 5433475 |
| caenorhabditis_elegans_prjna13758 | WBGene00010846 | IV | 11528052 | 11530715 | 1 | Tsp_07029 | | 9070841 | GL622785 | 9073953 |
| caenorhabditis_elegans_prjna13758 | WBGene00001585 | IV | 11521290 | 11522214 | 1 | Tsp_07529 | | 1777139 | GL622784 | 1777910 |
| caenorhabditis_elegans_prjna13758 | WBGene00001585 | IV | 11521290 | 11522214 | 1 | Tsp_12526 | | 5307 | GL627130 | 6078 |
| caenorhabditis_elegans_prjna13758 | WBGene00001585 | IV | 11521290 | 11522214 | 1 | Tsp_08715 | | 4120133 | GL622789 | 4121280 |
| caenorhabditis_elegans_prjna13758 | WBGene00019773 | IV | 458032 | 461029 | -1 | Tsp_07545 | | 1711702 | GL622784 | 1712427 |
| caenorhabditis_elegans_prjna13758 | WBGene00001949 | I | 7190834 | 7193924 | -1 | Tsp_11731 | | 198798 | GL622791 | 201603 |
| caenorhabditis_elegans_prjna13758 | WBGene00003367 | II | 10824294 | 10841424 | 1 | Tsp_04889 | | 1371239 | GL624340 | 1376762 |
| caenorhabditis_elegans_prjna13758 | WBGene00010915 | II | 8232789 | 8237202 | 1 | Tsp_02642 | | 764349 | GL622785 | 772025 |
| caenorhabditis_elegans_prjna13758 | WBGene00010891 | IV | 12121266 | 12122881 | 1 | Tsp_03934 | | 1147658 | GL622788 | 1148366 |
| caenorhabditis_elegans_prjna13758 | WBGene00010892 | IV | 12124952 | 12126785 | 1 | Tsp_02740 | | 1138372 | GL622785 | 1138702 |
| caenorhabditis_elegans_prjna13758 | WBGene00002214 | IV | 11083735 | 11086050 | 1 | Tsp_03007 | | 2305837 | GL622785 | 2308811 |

| | | | | | | | | | |
|---|---|---|---|---|---|---|---|---|---|
| caenorhabditis_elegans_prjna13758 | WBGene00010883 | IV | 11089797 | 11091856 | 1 | Tsp_00744 | 2962847 | GL622787 | 2964285 |
| caenorhabditis_elegans_prjna13758 | WBGene00010905 | III | 4541326 | 4542452 | -1 | Tsp_07613 | 9891940 | GL622785 | 9892530 |
| caenorhabditis_elegans_prjna13758 | WBGene00006713 | III | 7067686 | 7068570 | -1 | Tsp_00154 | 608812 | GL622787 | 609386 |
| caenorhabditis_elegans_prjna13758 | WBGene00019825 | IV | 255966 | 257199 | -1 | Tsp_02314 | 9376279 | GL622787 | 9377897 |
| caenorhabditis_elegans_prjna13758 | WBGene00011032 | I | 8598658 | 8600923 | 1 | Tsp_09154 | 3318544 | GL622784 | 3321887 |
| caenorhabditis_elegans_prjna13758 | WBGene00011035 | I | 8601339 | 8602034 | 1 | Tsp_00800 | 3132609 | GL622787 | 3133685 |
| caenorhabditis_elegans_prjna13758 | WBGene00019878 | III | 8364078 | 8366538 | 1 | Tsp_00277 | 1139368 | GL622787 | 1142464 |
| caenorhabditis_elegans_prjna13758 | WBGene00007008 | III | 8358625 | 8362496 | 1 | Tsp_15935 | 551 | GL625682 | 979 |
| caenorhabditis_elegans_prjna13758 | WBGene00007008 | III | 8358625 | 8362496 | 1 | Tsp_13975 | 206 | GL625313 | 1093 |
| caenorhabditis_elegans_prjna13758 | WBGene00007008 | III | 8358625 | 8362496 | 1 | Tsp_02037 | 8310458 | GL622787 | 8315262 |
| caenorhabditis_elegans_prjna13758 | WBGene00003429 | II | 4916179 | 4916562 | -1 | Tsp_00768 | 3048065 | GL622787 | 3048698 |
| caenorhabditis_elegans_prjna13758 | WBGene00003429 | II | 4916179 | 4916562 | -1 | Tsp_00761 | 3076026 | GL622787 | 3077501 |
| caenorhabditis_elegans_prjna13758 | WBGene00003429 | II | 4916179 | 4916562 | -1 | Tsp_00756 | 2908432 | GL622787 | 2908941 |
| caenorhabditis_elegans_prjna13758 | WBGene00003429 | II | 4916179 | 4916562 | -1 | Tsp_13425 | 585 | GL623126 | 965 |
| caenorhabditis_elegans_prjna13758 | WBGene00003429 | II | 4916179 | 4916562 | -1 | Tsp_13917 | 820 | GL624354 | 1264 |
| caenorhabditis_elegans_prjna13758 | WBGene00003430 | II | 4898616 | 4899188 | 1 | Tsp_00768 | 3048065 | GL622787 | 3048698 |
| caenorhabditis_elegans_prjna13758 | WBGene00003430 | II | 4898616 | 4899188 | 1 | Tsp_00761 | 3076026 | GL622787 | 3077501 |
| caenorhabditis_elegans_prjna13758 | WBGene00003430 | II | 4898616 | 4899188 | 1 | Tsp_00756 | 2908432 | GL622787 | 2908941 |
| caenorhabditis_elegans_prjna13758 | WBGene00003430 | II | 4898616 | 4899188 | 1 | Tsp_13425 | 585 | GL623126 | 965 |
| caenorhabditis_elegans_prjna13758 | WBGene00003430 | II | 4898616 | 4899188 | 1 | Tsp_13917 | 820 | GL624354 | 1264 |
| caenorhabditis_elegans_prjna13758 | WBGene00003431 | II | 4892270 | 4892728 | -1 | Tsp_00768 | 3048065 | GL622787 | 3048698 |
| caenorhabditis_elegans_prjna13758 | WBGene00003431 | II | 4892270 | 4892728 | -1 | Tsp_00761 | 3076026 | GL622787 | 3077501 |
| caenorhabditis_elegans_prjna13758 | WBGene00003431 | II | 4892270 | 4892728 | -1 | Tsp_00756 | 2908432 | GL622787 | 2908941 |
| caenorhabditis_elegans_prjna13758 | WBGene00003431 | II | 4892270 | 4892728 | -1 | Tsp_13425 | 585 | GL623126 | 965 |
| caenorhabditis_elegans_prjna13758 | WBGene00003431 | II | 4892270 | 4892728 | -1 | Tsp_13917 | 820 | GL624354 | 1264 |
| caenorhabditis_elegans_prjna13758 | WBGene00011043 | II | 14857259 | 14866636 | 1 | Tsp_06219 | 10047449 | GL622787 | 10049824 |

| | | | | | | | | | | |
|---|---|---|---|---|---|---|---|---|---|---|
| caenorhabditis_elegans_prjna13758 | WBGene00000389 | II | 10188263 | 10189469 | -1 | Tsp_13015.2 | | 3440 | GL624006 | 4762 |
| caenorhabditis_elegans_prjna13758 | WBGene00000389 | II | 10188263 | 10189469 | -1 | Tsp_13015.1 | | 4885 | GL624006 | 5857 |
| caenorhabditis_elegans_prjna13758 | WBGene00001836 | I | 11912791 | 11917719 | -1 | Tsp_11412 | | 7474289 | GL622792 | 7477169 |
| caenorhabditis_elegans_prjna13758 | WBGene00000275 | I | 7251941 | 7255223 | 1 | Tsp_01159 | | 4485369 | GL622787 | 4489700 |
| caenorhabditis_elegans_prjna13758 | WBGene00000411 | II | 10785452 | 10787272 | 1 | Tsp_10275 | | 6849929 | GL622792 | 6851808 |
| caenorhabditis_elegans_prjna13758 | WBGene00011109 | III | 4408551 | 4412149 | 1 | Tsp_08239 | NTH1 | 2594879 | GL622789 | 2601583 |
| caenorhabditis_elegans_prjna13758 | WBGene00004387 | III | 4406286 | 4407326 | 1 | Tsp_04007 | | 1449927 | GL622788 | 1450781 |
| caenorhabditis_elegans_prjna13758 | WBGene00011111 | III | 4404786 | 4406285 | 1 | Tsp_06227 | | 10021142 | GL622787 | 10022072 |
| caenorhabditis_elegans_prjna13758 | WBGene00011111 | III | 4404786 | 4406285 | 1 | Tsp_06226 | | 10022152 | GL622787 | 10023344 |
| caenorhabditis_elegans_prjna13758 | WBGene00019941 | II | 7597545 | 7598752 | -1 | Tsp_00519 | | 2085953 | GL622787 | 2087490 |
| caenorhabditis_elegans_prjna13758 | WBGene00019951 | IV | 4433049 | 4434541 | -1 | Tsp_01125 | | 4353662 | GL622787 | 4354442 |
| caenorhabditis_elegans_prjna13758 | WBGene00011142 | III | 8966022 | 8967727 | -1 | Tsp_07526 | | 1784443 | GL622784 | 1786567 |
| caenorhabditis_elegans_prjna13758 | WBGene00019983 | V | 773518 | 774854 | 1 | Tsp_03199 | | 3225988 | GL622785 | 3228198 |
| caenorhabditis_elegans_prjna13758 | WBGene00003157 | III | 4288264 | 4291283 | -1 | Tsp_07749 | | 10460904 | GL622785 | 10464747 |
| caenorhabditis_elegans_prjna13758 | WBGene00003064 | IV | 10387413 | 10388345 | 1 | Tsp_04117 | | 2045990 | GL622788 | 2048480 |
| caenorhabditis_elegans_prjna13758 | WBGene00001324 | IV | 4793202 | 4799378 | -1 | Tsp_13826 | | 4 | GL628678 | 2233 |
| caenorhabditis_elegans_prjna13758 | WBGene00011250 | V | 14597939 | 14600092 | 1 | Tsp_03377 | | 4015733 | GL622785 | 4019537 |
| caenorhabditis_elegans_prjna13758 | WBGene00000846 | III | 7584181 | 7591570 | 1 | Tsp_10515 | Mcoln3 | 5789558 | GL622792 | 5793570 |
| caenorhabditis_elegans_prjna13758 | WBGene00020053 | III | 7591570 | 7593337 | -1 | Tsp_06294 | | 10331217 | GL622787 | 10333834 |
| caenorhabditis_elegans_prjna13758 | WBGene00003063 | III | 7573350 | 7575263 | 1 | Tsp_02259 | | 9182235 | GL622787 | 9184741 |
| caenorhabditis_elegans_prjna13758 | WBGene00001978 | V | 11845877 | 11852165 | 1 | Tsp_10127 | | 594342 | GL622789 | 599708 |
| caenorhabditis_elegans_prjna13758 | WBGene00003450 | IV | 5062534 | 5062995 | 1 | Tsp_00768 | | 3048065 | GL622787 | 3048698 |
| caenorhabditis_elegans_prjna13758 | WBGene00003450 | IV | 5062534 | 5062995 | 1 | Tsp_00761 | | 3076026 | GL622787 | 3077501 |
| caenorhabditis_elegans_prjna13758 | WBGene00003450 | IV | 5062534 | 5062995 | 1 | Tsp_00756 | | 2908432 | GL622787 | 2908941 |
| caenorhabditis_elegans_prjna13758 | WBGene00003450 | IV | 5062534 | 5062995 | 1 | Tsp_13425 | | 585 | GL623126 | 965 |
| caenorhabditis_elegans_prjna13758 | WBGene00003450 | IV | 5062534 | 5062995 | 1 | Tsp_13917 | | 820 | GL624354 | 1264 |

| | | | | | | | | | | |
|---|---|---|---|---|---|---|---|---|---|---|
| caenorhabditis_elegans_prjna13758 | WBGene00003446 | IV | 5064643 | 5065153 | -1 | Tsp_00768 | | 3048065 | GL622787 | 3048698 |
| caenorhabditis_elegans_prjna13758 | WBGene00003446 | IV | 5064643 | 5065153 | -1 | Tsp_00761 | | 3076026 | GL622787 | 3077501 |
| caenorhabditis_elegans_prjna13758 | WBGene00003446 | IV | 5064643 | 5065153 | -1 | Tsp_00756 | | 2908432 | GL622787 | 2908941 |
| caenorhabditis_elegans_prjna13758 | WBGene00003446 | IV | 5064643 | 5065153 | -1 | Tsp_13425 | | 585 | GL623126 | 965 |
| caenorhabditis_elegans_prjna13758 | WBGene00003446 | IV | 5064643 | 5065153 | -1 | Tsp_13917 | | 820 | GL624354 | 1264 |
| caenorhabditis_elegans_prjna13758 | WBGene00020071 | IV | 5065746 | 5066987 | -1 | Tsp_02051 | | 8391493 | GL622787 | 8393242 |
| caenorhabditis_elegans_prjna13758 | WBGene00020071 | IV | 5065746 | 5066987 | -1 | Tsp_06549 | | 11192741 | GL622787 | 11195348 |
| caenorhabditis_elegans_prjna13758 | WBGene00020071 | IV | 5065746 | 5066987 | -1 | Tsp_00662 | Ttbk2 | 2539389 | GL622787 | 2540670 |
| caenorhabditis_elegans_prjna13758 | WBGene00020071 | IV | 5065746 | 5066987 | -1 | Tsp_04341 | Ttbk1 | 3010978 | GL622788 | 3012100 |
| caenorhabditis_elegans_prjna13758 | WBGene00020071 | IV | 5065746 | 5066987 | -1 | Tsp_06889.2 | | 8369982 | GL622785 | 8372542 |
| caenorhabditis_elegans_prjna13758 | WBGene00020072 | IV | 5067320 | 5068652 | -1 | Tsp_02051 | | 8391493 | GL622787 | 8393242 |
| caenorhabditis_elegans_prjna13758 | WBGene00020072 | IV | 5067320 | 5068652 | -1 | Tsp_06549 | | 11192741 | GL622787 | 11195348 |
| caenorhabditis_elegans_prjna13758 | WBGene00020072 | IV | 5067320 | 5068652 | -1 | Tsp_00662 | Ttbk2 | 2539389 | GL622787 | 2540670 |
| caenorhabditis_elegans_prjna13758 | WBGene00020072 | IV | 5067320 | 5068652 | -1 | Tsp_04341 | Ttbk1 | 3010978 | GL622788 | 3012100 |
| caenorhabditis_elegans_prjna13758 | WBGene00020072 | IV | 5067320 | 5068652 | -1 | Tsp_06889.2 | | 8369982 | GL622785 | 8372542 |
| caenorhabditis_elegans_prjna13758 | WBGene00020111 | III | 7226820 | 7229787 | -1 | Tsp_01453 | | 5911834 | GL622787 | 5914253 |
| caenorhabditis_elegans_prjna13758 | WBGene00004185 | II | 10539438 | 10541996 | -1 | Tsp_02266 | | 9157033 | GL622787 | 9161566 |
| caenorhabditis_elegans_prjna13758 | WBGene00011268 | III | 10840023 | 10845020 | -1 | Tsp_09418 | | 21073 | GL622792 | 24189 |
| caenorhabditis_elegans_prjna13758 | WBGene00011308 | V | 12971110 | 12973282 | 1 | Tsp_02832 | | 1526439 | GL622785 | 1530205 |
| caenorhabditis_elegans_prjna13758 | WBGene00011280 | III | 4202253 | 4204269 | -1 | Tsp_13129 | | 2797 | GL625762 | 3977 |
| caenorhabditis_elegans_prjna13758 | WBGene00020140 | IV | 8459178 | 8460327 | -1 | Tsp_01983 | | 8105151 | GL622787 | 8107104 |
| caenorhabditis_elegans_prjna13758 | WBGene00004279 | II | 8713992 | 8715013 | -1 | Tsp_09592 | | 5469268 | GL622789 | 5470173 |
| caenorhabditis_elegans_prjna13758 | WBGene00000887 | II | 8715132 | 8716281 | -1 | Tsp_05613 | | 4295788 | GL622792 | 4296699 |
| caenorhabditis_elegans_prjna13758 | WBGene00011311 | II | 8716834 | 8721691 | 1 | Tsp_11258 | | 102345 | GL624139 | 104537 |
| caenorhabditis_elegans_prjna13758 | WBGene00011318 | V | 14987556 | 14991024 | -1 | Tsp_03558 | | 4720946 | GL622785 | 4726663 |
| caenorhabditis_elegans_prjna13758 | WBGene00001423 | V | 15000653 | 15002273 | -1 | Tsp_01344 | | 5476678 | GL622787 | 5479277 |

| | | | | | | | | | | |
|---|---|---|---|---|---|---|---|---|---|---|
| caenorhabditis_elegans_prjna13758 | WBGene00003183 | I | 8294881 | 8296940 | 1 | Tsp_10633 | KATNA1 | 6350258 | GL622792 | 6352735 |
| caenorhabditis_elegans_prjna13758 | WBGene00011368 | III | 4022166 | 4025658 | 1 | Tsp_02682 | | 907792 | GL622785 | 910499 |
| caenorhabditis_elegans_prjna13758 | WBGene00020172 | II | 702107 | 708608 | 1 | Tsp_12746 | | 5304 | GL624343 | 6632 |
| caenorhabditis_elegans_prjna13758 | WBGene00001865 | IV | 12490791 | 12495298 | 1 | Tsp_10696 | | 2821007 | GL622789 | 2823425 |
| caenorhabditis_elegans_prjna13758 | WBGene00001259 | III | 4715293 | 4720670 | -1 | Tsp_02371 | | 9889426 | GL622787 | 9896445 |
| caenorhabditis_elegans_prjna13758 | WBGene00011415 | III | 4712572 | 4715188 | -1 | Tsp_01912 | | 7829760 | GL622787 | 7834408 |
| caenorhabditis_elegans_prjna13758 | WBGene00001487 | IV | 10043183 | 10045739 | 1 | Tsp_12784 | | 3362 | GL624279 | 5110 |
| caenorhabditis_elegans_prjna13758 | WBGene00001487 | IV | 10043183 | 10045739 | 1 | Tsp_06641 | Fps85D | 11546255 | GL622787 | 11548147 |
| caenorhabditis_elegans_prjna13758 | WBGene00001487 | IV | 10043183 | 10045739 | 1 | Tsp_11870 | | 113432 | GL622790 | 117458 |
| caenorhabditis_elegans_prjna13758 | WBGene00004296 | V | 12245594 | 12251246 | 1 | Tsp_00253 | | 980447 | GL622787 | 985944 |
| caenorhabditis_elegans_prjna13758 | WBGene00011451 | V | 12251341 | 12252407 | 1 | Tsp_13301 | | 27 | GL624879 | 1157 |
| caenorhabditis_elegans_prjna13758 | WBGene00011466 | II | 8173680 | 8174866 | 1 | Tsp_05227 | Ttbk1 | 2180440 | GL622792 | 2181339 |
| caenorhabditis_elegans_prjna13758 | WBGene00011466 | II | 8173680 | 8174866 | 1 | Tsp_12209 | | 4940141 | GL622792 | 4941163 |
| caenorhabditis_elegans_prjna13758 | WBGene00011466 | II | 8173680 | 8174866 | 1 | Tsp_11208 | | 5439495 | GL622792 | 5440343 |
| caenorhabditis_elegans_prjna13758 | WBGene00011466 | II | 8173680 | 8174866 | 1 | Tsp_12401 | | 2248 | GL623394 | 3147 |
| caenorhabditis_elegans_prjna13758 | WBGene00011466 | II | 8173680 | 8174866 | 1 | Tsp_12402 | | 5826 | GL623394 | 6369 |
| caenorhabditis_elegans_prjna13758 | WBGene00011466 | II | 8173680 | 8174866 | 1 | Tsp_13461 | | 44 | GL626854 | 695 |
| caenorhabditis_elegans_prjna13758 | WBGene00000377 | II | 8185443 | 8187601 | 1 | Tsp_04075 | | 1856639 | GL622788 | 1860687 |
| caenorhabditis_elegans_prjna13758 | WBGene00020263 | I | 5472321 | 5479438 | -1 | Tsp_10024 | Dhx33 | 88818 | GL622789 | 90968 |
| caenorhabditis_elegans_prjna13758 | WBGene00011488 | I | 9620233 | 9622370 | 1 | Tsp_11215 | | 5412548 | GL622792 | 5416240 |
| caenorhabditis_elegans_prjna13758 | WBGene00011493 | I | 9642344 | 9643804 | 1 | Tsp_02716 | | 1067687 | GL622785 | 1074809 |
| caenorhabditis_elegans_prjna13758 | WBGene00002064 | III | 9745864 | 9747193 | -1 | Tsp_01882 | | 7726849 | GL622787 | 7727199 |
| caenorhabditis_elegans_prjna13758 | WBGene00000405 | III | 9747432 | 9748852 | 1 | Tsp_11762 | | 5045814 | GL622789 | 5047211 |
| caenorhabditis_elegans_prjna13758 | WBGene00000151 | II | 8047564 | 8049105 | 1 | Tsp_01310 | | 5302456 | GL622787 | 5306555 |
| caenorhabditis_elegans_prjna13758 | WBGene00000151 | II | 8047564 | 8049105 | 1 | Tsp_13029 | | 1642 | GL625299 | 2581 |
| caenorhabditis_elegans_prjna13758 | WBGene00020273 | V | 6417495 | 6424155 | -1 | Tsp_06685 | | 7380175 | GL622785 | 7386332 |

| | | | | | | | | | | |
|---|---|---|---|---|---|---|---|---|---|---|
| caenorhabditis_elegans_prjna13758 | WBGene00001513 | V | 6431331 | 6433496 | -1 | Tsp_03285 | | 3600633 | GL622785 | 3603195 |
| caenorhabditis_elegans_prjna13758 | WBGene00011527 | II | 11235053 | 11236450 | -1 | Tsp_01197 | | 4796829 | GL622787 | 4797680 |
| caenorhabditis_elegans_prjna13758 | WBGene00011527 | II | 11235053 | 11236450 | -1 | Tsp_01200 | | 4812869 | GL622787 | 4813486 |
| caenorhabditis_elegans_prjna13758 | WBGene00011527 | II | 11235053 | 11236450 | -1 | Tsp_01207 | | 4826543 | GL622787 | 4827437 |
| caenorhabditis_elegans_prjna13758 | WBGene00011527 | II | 11235053 | 11236450 | -1 | Tsp_01210 | | 4834824 | GL622787 | 4835438 |
| caenorhabditis_elegans_prjna13758 | WBGene00004465 | II | 11237447 | 11239092 | -1 | Tsp_00727 | PSMD13 | 2883240 | GL622787 | 2885478 |
| caenorhabditis_elegans_prjna13758 | WBGene00011538 | V | 15393108 | 15394957 | -1 | Tsp_08265 | | 3561911 | GL622788 | 3563551 |
| caenorhabditis_elegans_prjna13758 | WBGene00011538 | V | 15393108 | 15394957 | -1 | Tsp_12764 | | 1624 | GL623251 | 3277 |
| caenorhabditis_elegans_prjna13758 | WBGene00011538 | V | 15393108 | 15394957 | -1 | Tsp_12765 | | 3307 | GL623251 | 4368 |
| caenorhabditis_elegans_prjna13758 | WBGene00000913 | IV | 419938 | 425181 | -1 | Tsp_07635 | | 9991554 | GL622785 | 9993423 |
| caenorhabditis_elegans_prjna13758 | WBGene00011589 | V | 12868026 | 12870035 | -1 | Tsp_13758 | | 192 | GL626449 | 656 |
| caenorhabditis_elegans_prjna13758 | WBGene00011605 | III | 4263785 | 4269270 | -1 | Tsp_00780 | | 2994595 | GL622787 | 3001862 |
| caenorhabditis_elegans_prjna13758 | WBGene00011631 | I | 8902747 | 8905534 | -1 | Tsp_06900 | | 8416946 | GL622785 | 8418938 |
| caenorhabditis_elegans_prjna13758 | WBGene00011625 | V | 14034323 | 14038372 | -1 | Tsp_09220 | | 5006995 | GL622788 | 5010666 |
| caenorhabditis_elegans_prjna13758 | WBGene00007014 | II | 7853880 | 7855042 | -1 | Tsp_14529 | | 179 | GL628073 | 898 |
| caenorhabditis_elegans_prjna13758 | WBGene00007014 | II | 7853880 | 7855042 | -1 | Tsp_07623 | med10 | 9923229 | GL622785 | 9924651 |
| caenorhabditis_elegans_prjna13758 | WBGene00007014 | II | 7853880 | 7855042 | -1 | Tsp_13812 | | 317 | GL626659 | 1144 |
| caenorhabditis_elegans_prjna13758 | WBGene00011637 | II | 7858401 | 7859781 | 1 | Tsp_06375 | Ppp1r7 | 10759467 | GL622787 | 10760411 |
| caenorhabditis_elegans_prjna13758 | WBGene00020375 | I | 6185462 | 6188121 | 1 | Tsp_07331 | | 1126768 | GL622784 | 1128999 |
| caenorhabditis_elegans_prjna13758 | WBGene00000500 | I | 6181314 | 6183913 | -1 | Tsp_15308 | | 141 | GL627810 | 849 |
| caenorhabditis_elegans_prjna13758 | WBGene00020391 | V | 1872246 | 1875526 | 1 | Tsp_00784 | | 3091478 | GL622787 | 3098883 |
| caenorhabditis_elegans_prjna13758 | WBGene00011687 | V | 16023351 | 16024185 | 1 | Tsp_10929 | | 9359671 | GL622785 | 9363737 |
| caenorhabditis_elegans_prjna13758 | WBGene00006736 | III | 5165659 | 5169799 | -1 | Tsp_07219 | | 643234 | GL622784 | 648554 |
| caenorhabditis_elegans_prjna13758 | WBGene00020423 | III | 5169933 | 5172045 | -1 | Tsp_00442 | | 1536056 | GL622787 | 1541396 |
| caenorhabditis_elegans_prjna13758 | WBGene00020425 | V | 6653613 | 6655449 | -1 | Tsp_01998 | | 8050910 | GL622787 | 8052800 |
| caenorhabditis_elegans_prjna13758 | WBGene00004374 | IV | 5469961 | 5473174 | 1 | Tsp_13554 | | 1224 | GL626375 | 1574 |

| | | | | | | | | | | |
|---|---|---|---|---|---|---|---|---|---|---|
| caenorhabditis_elegans_prjna13758 | WBGene00004374 | IV | 5469961 | 5473174 | 1 | Tsp_04363 | | 3099503 | GL622788 | 3102792 |
| caenorhabditis_elegans_prjna13758 | WBGene00011722 | IV | 10861402 | 10864232 | 1 | Tsp_04398 | | 3309670 | GL622788 | 3311602 |
| caenorhabditis_elegans_prjna13758 | WBGene00011722 | IV | 10861402 | 10864232 | 1 | Tsp_08298 | | 3677242 | GL622788 | 3680117 |
| caenorhabditis_elegans_prjna13758 | WBGene00011722 | IV | 10861402 | 10864232 | 1 | Tsp_13535 | | 298 | GL624218 | 1630 |
| caenorhabditis_elegans_prjna13758 | WBGene00020441 | III | 6244672 | 6246039 | -1 | Tsp_11517 | | 614600 | GL622790 | 615764 |
| caenorhabditis_elegans_prjna13758 | WBGene00011730 | III | 13642420 | 13643570 | -1 | Tsp_06589 | | 11799459 | GL622787 | 11801458 |
| caenorhabditis_elegans_prjna13758 | WBGene00011740 | IV | 12031314 | 12032154 | 1 | Tsp_01276 | mrpl-51 | 5108724 | GL622787 | 5110278 |
| caenorhabditis_elegans_prjna13758 | WBGene00006416 | IV | 6277435 | 6279572 | -1 | Tsp_05882 | | 5764868 | GL622785 | 5767620 |
| caenorhabditis_elegans_prjna13758 | WBGene00003466 | IV | 9766745 | 9767187 | -1 | Tsp_00768 | | 3048065 | GL622787 | 3048698 |
| caenorhabditis_elegans_prjna13758 | WBGene00003466 | IV | 9766745 | 9767187 | -1 | Tsp_00761 | | 3076026 | GL622787 | 3077501 |
| caenorhabditis_elegans_prjna13758 | WBGene00003466 | IV | 9766745 | 9767187 | -1 | Tsp_00756 | | 2908432 | GL622787 | 2908941 |
| caenorhabditis_elegans_prjna13758 | WBGene00003466 | IV | 9766745 | 9767187 | -1 | Tsp_13425 | | 585 | GL623126 | 965 |
| caenorhabditis_elegans_prjna13758 | WBGene00003466 | IV | 9766745 | 9767187 | -1 | Tsp_13917 | | 820 | GL624354 | 1264 |
| caenorhabditis_elegans_prjna13758 | WBGene00003465 | IV | 9770013 | 9770459 | 1 | Tsp_00768 | | 3048065 | GL622787 | 3048698 |
| caenorhabditis_elegans_prjna13758 | WBGene00003465 | IV | 9770013 | 9770459 | 1 | Tsp_00761 | | 3076026 | GL622787 | 3077501 |
| caenorhabditis_elegans_prjna13758 | WBGene00003465 | IV | 9770013 | 9770459 | 1 | Tsp_00756 | | 2908432 | GL622787 | 2908941 |
| caenorhabditis_elegans_prjna13758 | WBGene00003465 | IV | 9770013 | 9770459 | 1 | Tsp_13425 | | 585 | GL623126 | 965 |
| caenorhabditis_elegans_prjna13758 | WBGene00003465 | IV | 9770013 | 9770459 | 1 | Tsp_13917 | | 820 | GL624354 | 1264 |
| caenorhabditis_elegans_prjna13758 | WBGene00000301 | IV | 9772510 | 9773470 | 1 | Tsp_03271 | | 3538704 | GL622785 | 3539738 |
| caenorhabditis_elegans_prjna13758 | WBGene00011758 | II | 8523494 | 8525396 | -1 | Tsp_00418 | | 1621214 | GL622787 | 1625524 |
| caenorhabditis_elegans_prjna13758 | WBGene00011759 | II | 8525685 | 8526804 | 1 | Tsp_06024 | | 6373191 | GL622785 | 6374711 |
| caenorhabditis_elegans_prjna13758 | WBGene00006638 | IV | 10149166 | 10150602 | -1 | Tsp_03722 | | 237591 | GL622788 | 241095 |
| caenorhabditis_elegans_prjna13758 | WBGene00020553 | V | 8957045 | 8961935 | 1 | Tsp_13235 | | 21 | GL626784 | 1247 |
| caenorhabditis_elegans_prjna13758 | WBGene00011834 | V | 11237438 | 11239331 | -1 | Tsp_09283 | | 5266919 | GL622788 | 5271526 |
| caenorhabditis_elegans_prjna13758 | WBGene00006776 | I | 5679844 | 5691503 | -1 | Tsp_07995 | | 1261860 | GL622789 | 1269189 |
| caenorhabditis_elegans_prjna13758 | WBGene00001412 | III | 631938 | 634645 | -1 | Tsp_01554 | | 6339512 | GL622787 | 6341568 |

| | | | | | | | | | | |
|---|---|---|---|---|---|---|---|---|---|---|
| caenorhabditis_elegans_prjna13758 | WBGene00020600 | III | 7386229 | 7391135 | 1 | Tsp_05597 | TTC27 | 4153534 | GL622792 | 4155909 |
| caenorhabditis_elegans_prjna13758 | WBGene00006542 | III | 7384033 | 7386132 | 1 | Tsp_11535 | tbp-1 | 675579 | GL622790 | 676518 |
| caenorhabditis_elegans_prjna13758 | WBGene00020601 | III | 7382115 | 7383906 | 1 | Tsp_13943 | | 323 | GL626864 | 1558 |
| caenorhabditis_elegans_prjna13758 | WBGene00001974 | III | 7379143 | 7381609 | -1 | Tsp_07406 | | 1229278 | GL622784 | 1234117 |
| caenorhabditis_elegans_prjna13758 | WBGene00011860 | IV | 9341964 | 9343152 | -1 | Tsp_03777 | | 501559 | GL622788 | 503461 |
| caenorhabditis_elegans_prjna13758 | WBGene00003950 | I | 3908747 | 3909701 | 1 | Tsp_03495 | | 4501515 | GL622785 | 4502571 |
| caenorhabditis_elegans_prjna13758 | WBGene00004189 | III | 7238607 | 7241284 | 1 | Tsp_08341 | | 3872943 | GL622788 | 3879674 |
| caenorhabditis_elegans_prjna13758 | WBGene00004189 | III | 7238607 | 7241284 | 1 | Tsp_08340 | | 3871479 | GL622788 | 3872882 |
| caenorhabditis_elegans_prjna13758 | WBGene00004189 | III | 7238607 | 7241284 | 1 | Tsp_08339 | | 3870520 | GL622788 | 3871414 |
| caenorhabditis_elegans_prjna13758 | WBGene00000080 | III | 7230964 | 7232882 | -1 | Tsp_11657 | | 1556612 | GL622792 | 1558284 |
| caenorhabditis_elegans_prjna13758 | WBGene00000080 | III | 7230964 | 7232882 | -1 | Tsp_11655 | | 1541277 | GL622792 | 1542316 |
| caenorhabditis_elegans_prjna13758 | WBGene00000378 | II | 8927741 | 8929820 | 1 | Tsp_00786 | | 3179779 | GL622787 | 3182376 |
| caenorhabditis_elegans_prjna13758 | WBGene00020647 | IV | 259970 | 261683 | -1 | Tsp_00320 | | 1280085 | GL622787 | 1281609 |
| caenorhabditis_elegans_prjna13758 | WBGene00020659 | I | 6874304 | 6876928 | 1 | Tsp_12784 | | 3362 | GL624279 | 5110 |
| caenorhabditis_elegans_prjna13758 | WBGene00020659 | I | 6874304 | 6876928 | 1 | Tsp_06641 | Fps85D | 11546255 | GL622787 | 11548147 |
| caenorhabditis_elegans_prjna13758 | WBGene00020659 | I | 6874304 | 6876928 | 1 | Tsp_11870 | | 113432 | GL622790 | 117458 |
| caenorhabditis_elegans_prjna13758 | WBGene00001598 | I | 6854688 | 6857529 | 1 | Tsp_12124 | | 535552 | GL622790 | 536415 |
| caenorhabditis_elegans_prjna13758 | WBGene00001598 | I | 6854688 | 6857529 | 1 | Tsp_15219 | | 119 | GL623636 | 662 |
| caenorhabditis_elegans_prjna13758 | WBGene00011908 | I | 10598673 | 10606620 | 1 | Tsp_01393 | | 5843030 | GL622787 | 5844813 |
| caenorhabditis_elegans_prjna13758 | WBGene00011918 | I | 7955977 | 7958505 | -1 | Tsp_07545 | | 1711702 | GL622784 | 1712427 |
| caenorhabditis_elegans_prjna13758 | WBGene00004458 | IV | 6906415 | 6909913 | -1 | Tsp_03599 | | 4918846 | GL622785 | 4924517 |
| caenorhabditis_elegans_prjna13758 | WBGene00004422 | V | 3585313 | 3586064 | -1 | Tsp_02925 | | 1936917 | GL622785 | 1937747 |
| caenorhabditis_elegans_prjna13758 | WBGene00001040 | V | 8461096 | 8462116 | -1 | Tsp_05421 | | 3255936 | GL622792 | 3257269 |
| caenorhabditis_elegans_prjna13758 | WBGene00004315 | IV | 12724948 | 12730095 | 1 | Tsp_06041 | | 6501519 | GL622785 | 6503864 |
| caenorhabditis_elegans_prjna13758 | WBGene00001647 | I | 7699543 | 7700958 | 1 | Tsp_07744 | | 10481184 | GL622785 | 10483857 |
| caenorhabditis_elegans_prjna13758 | WBGene00020742 | I | 6441637 | 6446465 | -1 | Tsp_06531 | | 11268490 | GL622787 | 11275448 |

| | | | | | | | | | | |
|---|---|---|---|---|---|---|---|---|---|---|
| caenorhabditis_elegans_prjna13758 | WBGene00005023 | I | 9947812 | 9953387 | 1 | Tsp_10489 | | 1137481 | GL622791 | 1141837 |
| caenorhabditis_elegans_prjna13758 | WBGene00012000 | II | 9099371 | 9100639 | 1 | Tsp_03675 | | 87300 | GL622788 | 88464 |
| caenorhabditis_elegans_prjna13758 | WBGene00001041 | II | 9107382 | 9108600 | -1 | Tsp_00823 | Dnajc9 | 3203073 | GL622787 | 3206123 |
| caenorhabditis_elegans_prjna13758 | WBGene00012020 | V | 16745080 | 16746258 | -1 | Tsp_05820 | Gyg1 | 5538646 | GL622785 | 5541470 |
| caenorhabditis_elegans_prjna13758 | WBGene00000496 | I | 7554855 | 7559916 | 1 | Tsp_04418 | | 3371013 | GL622788 | 3384125 |
| caenorhabditis_elegans_prjna13758 | WBGene00012030 | I | 7559839 | 7562349 | 1 | Tsp_03870 | | 891698 | GL622788 | 894870 |
| caenorhabditis_elegans_prjna13758 | WBGene00020820 | III | 6460120 | 6462222 | 1 | Tsp_00267 | | 1169613 | GL622787 | 1171175 |
| caenorhabditis_elegans_prjna13758 | WBGene00020822 | III | 6448075 | 6450696 | -1 | Tsp_00901 | | 3667172 | GL622787 | 3668573 |
| caenorhabditis_elegans_prjna13758 | WBGene00020822 | III | 6448075 | 6450696 | -1 | Tsp_00899 | | 3671885 | GL622787 | 3674176 |
| caenorhabditis_elegans_prjna13758 | WBGene00020822 | III | 6448075 | 6450696 | -1 | Tsp_00900 | | 3668753 | GL622787 | 3671181 |
| caenorhabditis_elegans_prjna13758 | WBGene00020822 | III | 6448075 | 6450696 | -1 | Tsp_00902 | | 3664496 | GL622787 | 3667072 |
| caenorhabditis_elegans_prjna13758 | WBGene00004781 | III | 6450792 | 6452372 | -1 | Tsp_14266 | | 829 | GL626598 | 1008 |
| caenorhabditis_elegans_prjna13758 | WBGene00004781 | III | 6450792 | 6452372 | -1 | Tsp_09846 | | 171180 | GL623393 | 172450 |
| caenorhabditis_elegans_prjna13758 | WBGene00020827 | IV | 8428961 | 8431493 | -1 | Tsp_03718 | | 226016 | GL622788 | 229652 |
| caenorhabditis_elegans_prjna13758 | WBGene00012059 | III | 9404484 | 9406641 | -1 | Tsp_02129 | | 8617204 | GL622787 | 8620872 |
| caenorhabditis_elegans_prjna13758 | WBGene00000407 | III | 13464687 | 13466390 | -1 | Tsp_01981 | | 8110096 | GL622787 | 8110974 |
| caenorhabditis_elegans_prjna13758 | WBGene00000407 | III | 13464687 | 13466390 | -1 | Tsp_01982 | | 8107303 | GL622787 | 8109300 |
| caenorhabditis_elegans_prjna13758 | WBGene00004806 | V | 11645481 | 11650090 | 1 | Tsp_10826 | | 62522 | GL623868 | 66454 |
| caenorhabditis_elegans_prjna13758 | WBGene00004190 | I | 12486872 | 12490931 | 1 | Tsp_01086 | | 4175522 | GL622787 | 4177225 |
| caenorhabditis_elegans_prjna13758 | WBGene00003373 | III | 13509301 | 13514570 | -1 | Tsp_09579 | MLH1 | 966316 | GL622792 | 968894 |
| caenorhabditis_elegans_prjna13758 | WBGene00012115 | I | 8160512 | 8168643 | 1 | Tsp_02645 | | 778233 | GL622785 | 783085 |
| caenorhabditis_elegans_prjna13758 | WBGene00012115 | I | 8160512 | 8168643 | 1 | Tsp_02647 | | 783973 | GL622785 | 790394 |
| caenorhabditis_elegans_prjna13758 | WBGene00012115 | I | 8160512 | 8168643 | 1 | Tsp_02648 | | 795405 | GL622785 | 796460 |
| caenorhabditis_elegans_prjna13758 | WBGene00012115 | I | 8160512 | 8168643 | 1 | Tsp_13859 | | 181 | GL627559 | 1047 |
| caenorhabditis_elegans_prjna13758 | WBGene00004916 | II | 6488830 | 6489761 | -1 | Tsp_09652 | | 5579145 | GL622789 | 5579507 |
| caenorhabditis_elegans_prjna13758 | WBGene00004916 | II | 6488830 | 6489761 | -1 | Tsp_09653 | | 5577113 | GL622789 | 5578169 |

| | | | | | | | | | | |
|---|---|---|---|---|---|---|---|---|---|---|
| caenorhabditis_elegans_prjna13758 | WBGene00007029 | V | 7081537 | 7084662 | -1 | Tsp_05023 | | 1776772 | GL624340 | 1778882 |
| caenorhabditis_elegans_prjna13758 | WBGene00012156 | II | 11442693 | 11444009 | 1 | Tsp_09764 | | 6150922 | GL622789 | 6154975 |
| caenorhabditis_elegans_prjna13758 | WBGene00020910 | V | 6497110 | 6498484 | 1 | Tsp_09229 | | 5031558 | GL622788 | 5033357 |
| caenorhabditis_elegans_prjna13758 | WBGene00020910 | V | 6497110 | 6498484 | 1 | Tsp_03947 | mogat2-a | 1187191 | GL622788 | 1189940 |
| caenorhabditis_elegans_prjna13758 | WBGene00020914 | I | 3287336 | 3293127 | 1 | Tsp_05440 | | 3427410 | GL622792 | 3431341 |
| caenorhabditis_elegans_prjna13758 | WBGene00020915 | I | 3267094 | 3269116 | 1 | Tsp_10405 | NOP58 | 647747 | GL622791 | 655042 |
| caenorhabditis_elegans_prjna13758 | WBGene00012187 | II | 14060061 | 14060624 | 1 | Tsp_01995 | | 8059169 | GL622787 | 8059729 |
| caenorhabditis_elegans_prjna13758 | WBGene00002275 | II | 14063450 | 14066170 | 1 | Tsp_07804 | | 10708106 | GL622785 | 10711427 |
| caenorhabditis_elegans_prjna13758 | WBGene00012192 | I | 12730011 | 12732753 | -1 | Tsp_05165 | | 2256160 | GL624340 | 2258019 |
| caenorhabditis_elegans_prjna13758 | WBGene00004156 | IV | 13329508 | 13331406 | -1 | Tsp_09987 | | 653677 | GL623393 | 654755 |
| caenorhabditis_elegans_prjna13758 | WBGene00004156 | IV | 13329508 | 13331406 | -1 | Tsp_09988 | | 654805 | GL623393 | 655588 |
| caenorhabditis_elegans_prjna13758 | WBGene00003230 | IV | 13353723 | 13356104 | 1 | Tsp_12414 | | 8096 | GL624002 | 8753 |
| caenorhabditis_elegans_prjna13758 | WBGene00003230 | IV | 13353723 | 13356104 | 1 | Tsp_08560 | zfp36l2 | 4699442 | GL622788 | 4700099 |
| caenorhabditis_elegans_prjna13758 | WBGene00003230 | IV | 13353723 | 13356104 | 1 | Tsp_08559 | Zfp36l3 | 4695882 | GL622788 | 4696511 |
| caenorhabditis_elegans_prjna13758 | WBGene00003230 | IV | 13353723 | 13356104 | 1 | Tsp_12415 | | 11512 | GL624002 | 12169 |
| caenorhabditis_elegans_prjna13758 | WBGene00003230 | IV | 13353723 | 13356104 | 1 | Tsp_12413.2 | | 5119 | GL624002 | 5437 |
| caenorhabditis_elegans_prjna13758 | WBGene00003230 | IV | 13353723 | 13356104 | 1 | Tsp_08561 | | 4702854 | GL622788 | 4703511 |
| caenorhabditis_elegans_prjna13758 | WBGene00012205 | II | 11472135 | 11473571 | -1 | Tsp_02600 | Mettl1 | 614187 | GL622785 | 615351 |
| caenorhabditis_elegans_prjna13758 | WBGene00004698 | II | 11453362 | 11455864 | 1 | Tsp_03996 | | 1408766 | GL622788 | 1410375 |
| caenorhabditis_elegans_prjna13758 | WBGene00004698 | II | 11453362 | 11455864 | 1 | Tsp_03995 | | 1405654 | GL622788 | 1408513 |
| caenorhabditis_elegans_prjna13758 | WBGene00007057 | I | 6729996 | 6730959 | -1 | Tsp_01983 | | 8105151 | GL622787 | 8107104 |
| caenorhabditis_elegans_prjna13758 | WBGene00006773 | I | 6742293 | 6745341 | -1 | Tsp_16036 | | 1 | GL628249 | 863 |
| caenorhabditis_elegans_prjna13758 | WBGene00006773 | I | 6742293 | 6745341 | -1 | Tsp_10041 | | 137529 | GL622789 | 141957 |
| caenorhabditis_elegans_prjna13758 | WBGene00004181 | I | 12528312 | 12532182 | -1 | Tsp_04784 | | 1053910 | GL624340 | 1056625 |
| caenorhabditis_elegans_prjna13758 | WBGene00004181 | I | 12528312 | 12532182 | -1 | Tsp_04535 | | 306928 | GL624340 | 308973 |
| caenorhabditis_elegans_prjna13758 | WBGene00004181 | I | 12528312 | 12532182 | -1 | Tsp_04532 | | 305202 | GL624340 | 305543 |

| | | | | | | | | | | |
|---|---|---|---|---|---|---|---|---|---|---|
| caenorhabditis_elegans_prjna13758 | WBGene00020964 | III | 5791171 | 5803042 | -1 | Tsp_04157.2 | | 2242742 | GL622788 | 2245386 |
| caenorhabditis_elegans_prjna13758 | WBGene00020964 | III | 5791171 | 5803042 | -1 | Tsp_04158 | | 2245398 | GL622788 | 2252119 |
| caenorhabditis_elegans_prjna13758 | WBGene00020964 | III | 5791171 | 5803042 | -1 | Tsp_04157.1 | | 2239278 | GL622788 | 2239478 |
| caenorhabditis_elegans_prjna13758 | WBGene00003228 | II | 11968637 | 11972334 | 1 | Tsp_12414 | | 8096 | GL624002 | 8753 |
| caenorhabditis_elegans_prjna13758 | WBGene00003228 | II | 11968637 | 11972334 | 1 | Tsp_08560 | zfp36l2 | 4699442 | GL622788 | 4700099 |
| caenorhabditis_elegans_prjna13758 | WBGene00003228 | II | 11968637 | 11972334 | 1 | Tsp_08559 | Zfp36l3 | 4695882 | GL622788 | 4696511 |
| caenorhabditis_elegans_prjna13758 | WBGene00003228 | II | 11968637 | 11972334 | 1 | Tsp_12415 | | 11512 | GL624002 | 12169 |
| caenorhabditis_elegans_prjna13758 | WBGene00003228 | II | 11968637 | 11972334 | 1 | Tsp_12413.2 | | 5119 | GL624002 | 5437 |
| caenorhabditis_elegans_prjna13758 | WBGene00003228 | II | 11968637 | 11972334 | 1 | Tsp_08561 | | 4702854 | GL622788 | 4703511 |
| caenorhabditis_elegans_prjna13758 | WBGene00003955 | IV | 4047937 | 4049448 | -1 | Tsp_05750 | | 5173811 | GL622785 | 5175282 |
| caenorhabditis_elegans_prjna13758 | WBGene00021012 | IV | 506737 | 519457 | -1 | Tsp_03506 | | 4545440 | GL622785 | 4548962 |
| caenorhabditis_elegans_prjna13758 | WBGene00003906 | I | 4984760 | 4986739 | -1 | Tsp_03685 | | 110977 | GL622788 | 113440 |
| caenorhabditis_elegans_prjna13758 | WBGene00012230 | II | 14133397 | 14142289 | -1 | Tsp_08110 | | 1924019 | GL622789 | 1929762 |
| caenorhabditis_elegans_prjna13758 | WBGene00004298 | I | 9065087 | 9070264 | 1 | Tsp_09697 | | 5782043 | GL622789 | 5783275 |
| caenorhabditis_elegans_prjna13758 | WBGene00021063 | III | 636479 | 638419 | -1 | Tsp_05118 | | 2087244 | GL624340 | 2089259 |
| caenorhabditis_elegans_prjna13758 | WBGene00012315 | V | 16789267 | 16792771 | 1 | Tsp_00799 | | 3135386 | GL622787 | 3138525 |
| caenorhabditis_elegans_prjna13758 | WBGene00004248 | V | 16792764 | 16795007 | -1 | Tsp_09367 | | 5605503 | GL622788 | 5608171 |
| caenorhabditis_elegans_prjna13758 | WBGene00012319 | V | 20717872 | 20729271 | -1 | Tsp_02795 | | 1404505 | GL622785 | 1408361 |
| caenorhabditis_elegans_prjna13758 | WBGene00021073 | II | 471247 | 475940 | -1 | Tsp_05558 | | 4083046 | GL622792 | 4085415 |
| caenorhabditis_elegans_prjna13758 | WBGene00021074 | II | 480722 | 484386 | -1 | Tsp_06445 | | 10948700 | GL622787 | 10950757 |
| caenorhabditis_elegans_prjna13758 | WBGene00004188 | II | 484866 | 490403 | 1 | Tsp_02916 | | 1899354 | GL622785 | 1902598 |
| caenorhabditis_elegans_prjna13758 | WBGene00012342 | IV | 9825694 | 9830620 | 1 | Tsp_00364 | | 1830917 | GL622787 | 1836729 |
| caenorhabditis_elegans_prjna13758 | WBGene00012342 | IV | 9825694 | 9830620 | 1 | Tsp_12792.1 | | 388 | GL624745 | 3303 |
| caenorhabditis_elegans_prjna13758 | WBGene00012342 | IV | 9825694 | 9830620 | 1 | Tsp_12792.2 | | 4806 | GL624745 | 5386 |
| caenorhabditis_elegans_prjna13758 | WBGene00021088 | IV | 3295135 | 3297225 | 1 | Tsp_10838 | | 116320 | GL623868 | 118039 |
| caenorhabditis_elegans_prjna13758 | WBGene00004915 | I | 13339397 | 13340224 | 1 | Tsp_10755 | SNRPB | 3121111 | GL622789 | 3123425 |

| | | | | | | | | | | |
|---|---|---|---|---|---|---|---|---|---|---|
| caenorhabditis_elegans_prjna13758 | WBGene00004915 | I | 13339397 | 13340224 | 1 | Tsp_08631 | SNRPB | 3696553 | GL622789 | 3698374 |
| caenorhabditis_elegans_prjna13758 | WBGene00021093 | II | 580602 | 582014 | 1 | Tsp_07973 | | 1117673 | GL622789 | 1119094 |
| caenorhabditis_elegans_prjna13758 | WBGene00006387 | II | 1127976 | 1132352 | 1 | Tsp_03956 | | 1246348 | GL622788 | 1250032 |
| caenorhabditis_elegans_prjna13758 | WBGene00006387 | II | 1127976 | 1132352 | 1 | Tsp_14056 | | 1275 | GL627148 | 1634 |
| caenorhabditis_elegans_prjna13758 | WBGene00021112 | I | 4704598 | 4705881 | 1 | Tsp_00416 | | 1629186 | GL622787 | 1630317 |
| caenorhabditis_elegans_prjna13758 | WBGene00006793 | I | 13629227 | 13633494 | 1 | Tsp_10676 | | 2704969 | GL622789 | 2706937 |
| caenorhabditis_elegans_prjna13758 | WBGene00003133 | II | 11112719 | 11120788 | 1 | Tsp_11030 | | 1984517 | GL622784 | 1987050 |
| caenorhabditis_elegans_prjna13758 | WBGene00003133 | II | 11112719 | 11120788 | 1 | Tsp_11032 | | 1991174 | GL622784 | 1996568 |
| caenorhabditis_elegans_prjna13758 | WBGene00013669 | I | 14400679 | 14405649 | -1 | Tsp_10937 | | 9393302 | GL622785 | 9396848 |
| caenorhabditis_elegans_prjna13758 | WBGene00004959 | I | 5101682 | 5103467 | -1 | Tsp_06498 | | 11411948 | GL622787 | 11414270 |
| caenorhabditis_elegans_prjna13758 | WBGene00000502 | I | 5103581 | 5104956 | -1 | Tsp_11591 | | 1828987 | GL622792 | 1830416 |
| caenorhabditis_elegans_prjna13758 | WBGene00003132 | I | 5123916 | 5127199 | -1 | Tsp_10065 | | 270000 | GL622789 | 274096 |
| caenorhabditis_elegans_prjna13758 | WBGene00022455 | I | 5138256 | 5139382 | 1 | Tsp_03568 | ts | 4764689 | GL622785 | 4765912 |
| caenorhabditis_elegans_prjna13758 | WBGene00022458 | I | 5127828 | 5129611 | 1 | Tsp_02161 | | 8458286 | GL622787 | 8462805 |
| caenorhabditis_elegans_prjna13758 | WBGene00004700 | III | 12686243 | 12687916 | -1 | Tsp_02088 | | 8813576 | GL622787 | 8815674 |
| caenorhabditis_elegans_prjna13758 | WBGene00013766 | V | 20267377 | 20269711 | -1 | Tsp_04802 | | 1101096 | GL624340 | 1101765 |
| caenorhabditis_elegans_prjna13758 | WBGene00013766 | V | 20267377 | 20269711 | -1 | Tsp_04801 | | 1099760 | GL624340 | 1100904 |
| caenorhabditis_elegans_prjna13758 | WBGene00013777 | IV | 16830208 | 16830790 | 1 | Tsp_00768 | | 3048065 | GL622787 | 3048698 |
| caenorhabditis_elegans_prjna13758 | WBGene00013777 | IV | 16830208 | 16830790 | 1 | Tsp_00761 | | 3076026 | GL622787 | 3077501 |
| caenorhabditis_elegans_prjna13758 | WBGene00013777 | IV | 16830208 | 16830790 | 1 | Tsp_00756 | | 2908432 | GL622787 | 2908941 |
| caenorhabditis_elegans_prjna13758 | WBGene00013777 | IV | 16830208 | 16830790 | 1 | Tsp_13425 | | 585 | GL623126 | 965 |
| caenorhabditis_elegans_prjna13758 | WBGene00013777 | IV | 16830208 | 16830790 | 1 | Tsp_13917 | | 820 | GL624354 | 1264 |
| caenorhabditis_elegans_prjna13758 | WBGene00013801 | IV | 17068795 | 17070903 | -1 | Tsp_12784 | | 3362 | GL624279 | 5110 |
| caenorhabditis_elegans_prjna13758 | WBGene00013801 | IV | 17068795 | 17070903 | -1 | Tsp_06641 | Fps85D | 11546255 | GL622787 | 11548147 |
| caenorhabditis_elegans_prjna13758 | WBGene00013801 | IV | 17068795 | 17070903 | -1 | Tsp_11870 | | 113432 | GL622790 | 117458 |
| caenorhabditis_elegans_prjna13758 | WBGene00006405 | IV | 17119684 | 17126266 | 1 | Tsp_03846 | | 770778 | GL622788 | 778681 |

| | | | | | | | | | | |
|---|---|---|---|---|---|---|---|---|---|---|
| caenorhabditis_elegans_prjna13758 | WBGene00013811 | IV | 17131545 | 17133657 | 1 | Tsp_12784 | | 3362 | GL624279 | 5110 |
| caenorhabditis_elegans_prjna13758 | WBGene00013811 | IV | 17131545 | 17133657 | 1 | Tsp_06641 | Fps85D | 11546255 | GL622787 | 11548147 |
| caenorhabditis_elegans_prjna13758 | WBGene00013811 | IV | 17131545 | 17133657 | 1 | Tsp_11870 | | 113432 | GL622790 | 117458 |
| caenorhabditis_elegans_prjna13758 | WBGene00004914 | IV | 17088058 | 17088834 | 1 | Tsp_11736 | | 218573 | GL622791 | 219179 |
| caenorhabditis_elegans_prjna13758 | WBGene00004914 | IV | 17088058 | 17088834 | 1 | Tsp_11739 | | 225900 | GL622791 | 226506 |
| caenorhabditis_elegans_prjna13758 | WBGene00012465 | II | 12023861 | 12025932 | 1 | Tsp_09279 | Rexo4 | 5248668 | GL622788 | 5249877 |
| caenorhabditis_elegans_prjna13758 | WBGene00003154 | II | 12001510 | 12006625 | -1 | Tsp_00580 | | 2328053 | GL622787 | 2331856 |
| caenorhabditis_elegans_prjna13758 | WBGene00012484 | I | 12904470 | 12906212 | 1 | Tsp_05623 | | 4260352 | GL622792 | 4262001 |
| caenorhabditis_elegans_prjna13758 | WBGene00021277 | I | 2489976 | 2497371 | 1 | Tsp_07169 | DDX10 | 324481 | GL622784 | 329165 |
| caenorhabditis_elegans_prjna13758 | WBGene00004952 | I | 1030629 | 1033436 | -1 | Tsp_00833 | | 3553536 | GL622787 | 3560018 |
| caenorhabditis_elegans_prjna13758 | WBGene00006804 | III | 12881037 | 12891738 | -1 | Tsp_01232 | | 4935976 | GL622787 | 4940997 |
| caenorhabditis_elegans_prjna13758 | WBGene00012637 | IV | 13472497 | 13474005 | 1 | Tsp_02051 | | 8391493 | GL622787 | 8393242 |
| caenorhabditis_elegans_prjna13758 | WBGene00012637 | IV | 13472497 | 13474005 | 1 | Tsp_06549 | | 11192741 | GL622787 | 11195348 |
| caenorhabditis_elegans_prjna13758 | WBGene00012637 | IV | 13472497 | 13474005 | 1 | Tsp_00662 | Ttbk2 | 2539389 | GL622787 | 2540670 |
| caenorhabditis_elegans_prjna13758 | WBGene00012637 | IV | 13472497 | 13474005 | 1 | Tsp_04341 | Ttbk1 | 3010978 | GL622788 | 3012100 |
| caenorhabditis_elegans_prjna13758 | WBGene00012637 | IV | 13472497 | 13474005 | 1 | Tsp_06889.2 | | 8369982 | GL622785 | 8372542 |
| caenorhabditis_elegans_prjna13758 | WBGene00012650 | III | 10643002 | 10646107 | 1 | Tsp_03951 | | 1220486 | GL622788 | 1228626 |
| caenorhabditis_elegans_prjna13758 | WBGene00012652 | III | 10649519 | 10650908 | 1 | Tsp_04549 | | 340546 | GL624340 | 342173 |
| caenorhabditis_elegans_prjna13758 | WBGene00001087 | III | 10763131 | 10777343 | 1 | Tsp_00260 | | 928727 | GL622787 | 939037 |
| caenorhabditis_elegans_prjna13758 | WBGene00012666 | V | 19189246 | 19190035 | -1 | Tsp_03047 | | 2471224 | GL622785 | 2473944 |
| caenorhabditis_elegans_prjna13758 | WBGene00006567 | V | 18998345 | 19000918 | 1 | Tsp_04280 | | 2744775 | GL622788 | 2747369 |
| caenorhabditis_elegans_prjna13758 | WBGene00006567 | V | 18998345 | 19000918 | 1 | Tsp_04275 | | 2729827 | GL622788 | 2732391 |
| caenorhabditis_elegans_prjna13758 | WBGene00012694 | V | 18993940 | 18998200 | 1 | Tsp_03298 | | 3649630 | GL622785 | 3650438 |
| caenorhabditis_elegans_prjna13758 | WBGene00003156 | I | 2318291 | 2321847 | -1 | Tsp_06355 | | 10844784 | GL622787 | 10849995 |
| caenorhabditis_elegans_prjna13758 | WBGene00021466 | I | 2313271 | 2316003 | 1 | Tsp_00365 | | 1827442 | GL622787 | 1829578 |
| caenorhabditis_elegans_prjna13758 | WBGene00021466 | I | 2313271 | 2316003 | 1 | Tsp_12793.2 | | 6714 | GL624745 | 7124 |

| | | | | | | | | | | |
|---|---|---|---|---|---|---|---|---|---|---|
| caenorhabditis_elegans_prjna13758 | WBGene00021466 | I | 2313271 | 2316003 | 1 | Tsp_12793.1 | | 8298 | GL624745 | 9197 |
| caenorhabditis_elegans_prjna13758 | WBGene00000498 | V | 3757047 | 3759469 | -1 | Tsp_05158 | | 2233908 | GL624340 | 2235301 |
| caenorhabditis_elegans_prjna13758 | WBGene00000498 | V | 3757047 | 3759469 | -1 | Tsp_05157 | | 2232578 | GL624340 | 2233717 |
| caenorhabditis_elegans_prjna13758 | WBGene00012756 | III | 11710827 | 11718741 | -1 | Tsp_04583 | | 411430 | GL624340 | 415497 |
| caenorhabditis_elegans_prjna13758 | WBGene00004297 | IV | 10281637 | 10284244 | -1 | Tsp_02540 | | 363654 | GL622785 | 365300 |
| caenorhabditis_elegans_prjna13758 | WBGene00012786 | IV | 10353167 | 10355120 | 1 | Tsp_09675 | | 5634045 | GL622789 | 5639304 |
| caenorhabditis_elegans_prjna13758 | WBGene00000866 | IV | 10985985 | 10987426 | 1 | Tsp_05726 | ccnb2 | 4824667 | GL622792 | 4826249 |
| caenorhabditis_elegans_prjna13758 | WBGene00003804 | III | 13296853 | 13299122 | 1 | Tsp_05449 | seh1l | 3397253 | GL622792 | 3398909 |
| caenorhabditis_elegans_prjna13758 | WBGene00002229 | III | 13302743 | 13307259 | -1 | Tsp_01222 | KIF4A | 4867462 | GL622787 | 4872793 |
| caenorhabditis_elegans_prjna13758 | WBGene00012835 | V | 19697560 | 19705212 | 1 | Tsp_13631 | | 243 | GL623607 | 1460 |
| caenorhabditis_elegans_prjna13758 | WBGene00012855 | V | 20801184 | 20803006 | -1 | Tsp_00422 | | 1608385 | GL622787 | 1610657 |
| caenorhabditis_elegans_prjna13758 | WBGene00012927 | III | 11235581 | 11236868 | -1 | Tsp_09117 | | 3203898 | GL622784 | 3205025 |
| caenorhabditis_elegans_prjna13758 | WBGene00012927 | III | 11235581 | 11236868 | -1 | Tsp_09116 | | 3203269 | GL622784 | 3203820 |
| caenorhabditis_elegans_prjna13758 | WBGene00012927 | III | 11235581 | 11236868 | -1 | Tsp_09115 | | 3202042 | GL622784 | 3203090 |
| caenorhabditis_elegans_prjna13758 | WBGene00004873 | III | 11290911 | 11306145 | 1 | Tsp_00968 | | 3848230 | GL622787 | 3850133 |
| caenorhabditis_elegans_prjna13758 | WBGene00004873 | III | 11290911 | 11306145 | 1 | Tsp_00966 | | 3851850 | GL622787 | 3862430 |
| caenorhabditis_elegans_prjna13758 | WBGene00012935 | III | 11317570 | 11326664 | 1 | Tsp_13222 | | 112 | GL624949 | 2869 |
| caenorhabditis_elegans_prjna13758 | WBGene00012936 | III | 11142448 | 11154287 | 1 | Tsp_09727 | | 5949651 | GL622789 | 5955057 |
| caenorhabditis_elegans_prjna13758 | WBGene00003422 | I | 3449326 | 3462177 | 1 | Tsp_12822 | | 307 | GL626366 | 2673 |
| caenorhabditis_elegans_prjna13758 | WBGene00003422 | I | 3449326 | 3462177 | 1 | Tsp_09358 | | 5575956 | GL622788 | 5581903 |
| caenorhabditis_elegans_prjna13758 | WBGene00004775 | I | 3433157 | 3438543 | -1 | Tsp_02125 | | 8631029 | GL622787 | 8635834 |
| caenorhabditis_elegans_prjna13758 | WBGene00021636 | I | 3516450 | 3523387 | 1 | Tsp_05985 | | 6312102 | GL622785 | 6316074 |
| caenorhabditis_elegans_prjna13758 | WBGene00000794 | I | 3481987 | 3484499 | 1 | Tsp_05685 | FEN1 | 4587160 | GL622792 | 4589476 |
| caenorhabditis_elegans_prjna13758 | WBGene00012964 | III | 11008222 | 11008934 | 1 | Tsp_13708 | | 206 | GL626378 | 560 |
| caenorhabditis_elegans_prjna13758 | WBGene00012964 | III | 11008222 | 11008934 | 1 | Tsp_12947 | | 916 | GL626646 | 1694 |
| caenorhabditis_elegans_prjna13758 | WBGene00012978 | II | 14151040 | 14154373 | -1 | Tsp_13422 | | 2679 | GL622935 | 3128 |

| | | | | | | | | | | |
|---|---|---|---|---|---|---|---|---|---|---|
| caenorhabditis_elegans_prjna13758 | WBGene00012978 | II | 14151040 | 14154373 | -1 | Tsp_13421 | | 231 | GL622935 | 1354 |
| caenorhabditis_elegans_prjna13758 | WBGene00012978 | II | 14151040 | 14154373 | -1 | Tsp_07475 | | 1509884 | GL622784 | 1513322 |
| caenorhabditis_elegans_prjna13758 | WBGene00003119 | II | 13311459 | 13320911 | -1 | Tsp_09896 | | 357590 | GL623393 | 362388 |
| caenorhabditis_elegans_prjna13758 | WBGene00002079 | I | 318390 | 323352 | -1 | Tsp_08627 | | 3679011 | GL622789 | 3682663 |
| caenorhabditis_elegans_prjna13758 | WBGene00004896 | II | 3482408 | 3490143 | 1 | Tsp_03955 | | 1243374 | GL622788 | 1245173 |
| caenorhabditis_elegans_prjna13758 | WBGene00013047 | V | 14728441 | 14728854 | -1 | Tsp_00768 | | 3048065 | GL622787 | 3048698 |
| caenorhabditis_elegans_prjna13758 | WBGene00013047 | V | 14728441 | 14728854 | -1 | Tsp_00761 | | 3076026 | GL622787 | 3077501 |
| caenorhabditis_elegans_prjna13758 | WBGene00013047 | V | 14728441 | 14728854 | -1 | Tsp_00756 | | 2908432 | GL622787 | 2908941 |
| caenorhabditis_elegans_prjna13758 | WBGene00013047 | V | 14728441 | 14728854 | -1 | Tsp_13425 | | 585 | GL623126 | 965 |
| caenorhabditis_elegans_prjna13758 | WBGene00013047 | V | 14728441 | 14728854 | -1 | Tsp_13917 | | 820 | GL624354 | 1264 |
| caenorhabditis_elegans_prjna13758 | WBGene00021787 | II | 1409184 | 1411472 | -1 | Tsp_09634 | | 5527680 | GL622789 | 5529663 |
| caenorhabditis_elegans_prjna13758 | WBGene00013122 | I | 10966394 | 10970166 | -1 | Tsp_11425 | impact | 7542673 | GL622792 | 7543503 |
| caenorhabditis_elegans_prjna13758 | WBGene00002004 | I | 11953512 | 11961984 | -1 | Tsp_00094 | | 507428 | GL622787 | 512236 |
| caenorhabditis_elegans_prjna13758 | WBGene00000023 | I | 12011195 | 12018668 | 1 | Tsp_06590 | | 11775201 | GL622787 | 11796990 |
| caenorhabditis_elegans_prjna13758 | WBGene00006940 | II | 9713737 | 9716467 | -1 | Tsp_05938 | | 5998194 | GL622785 | 6000995 |
| caenorhabditis_elegans_prjna13758 | WBGene00013140 | II | 9701402 | 9703732 | -1 | Tsp_06938 | | 8565136 | GL622785 | 8567450 |
| caenorhabditis_elegans_prjna13758 | WBGene00013140 | II | 9701402 | 9703732 | -1 | Tsp_06943 | | 8584761 | GL622785 | 8585504 |
| caenorhabditis_elegans_prjna13758 | WBGene00013140 | II | 9701402 | 9703732 | -1 | Tsp_06939.2 | | 8567768 | GL622785 | 8567890 |
| caenorhabditis_elegans_prjna13758 | WBGene00013140 | II | 9701402 | 9703732 | -1 | Tsp_06942 | | 8582145 | GL622785 | 8584447 |
| caenorhabditis_elegans_prjna13758 | WBGene00013143 | II | 9742982 | 9746074 | -1 | Tsp_06424 | | 11041269 | GL622787 | 11045551 |
| caenorhabditis_elegans_prjna13758 | WBGene00021811 | III | 3311423 | 3317713 | 1 | Tsp_01642 | | 6637730 | GL622787 | 6646025 |
| caenorhabditis_elegans_prjna13758 | WBGene00021814 | III | 3282308 | 3290215 | 1 | Tsp_09207 | | 4925930 | GL622788 | 4927002 |
| caenorhabditis_elegans_prjna13758 | WBGene00021814 | III | 3282308 | 3290215 | 1 | Tsp_09192 | | 4857003 | GL622788 | 4858075 |
| caenorhabditis_elegans_prjna13758 | WBGene00021814 | III | 3282308 | 3290215 | 1 | Tsp_09206.2 | | 4924089 | GL622788 | 4924572 |
| caenorhabditis_elegans_prjna13758 | WBGene00021814 | III | 3282308 | 3290215 | 1 | Tsp_09206.1 | | 4925043 | GL622788 | 4925893 |
| caenorhabditis_elegans_prjna13758 | WBGene00021814 | III | 3282308 | 3290215 | 1 | Tsp_09191 | | 4855319 | GL622788 | 4856966 |

| | | | | | | | | | | |
|---|---|---|---|---|---|---|---|---|---|---|
| caenorhabditis_elegans_prjna13758 | WBGene00021828 | I | 3163688 | 3167925 | 1 | Tsp_00550 | | 1997048 | GL622787 | 1998165 |
| caenorhabditis_elegans_prjna13758 | WBGene00021841 | I | 3032935 | 3034761 | 1 | Tsp_08023 | | 1448659 | GL622789 | 1449744 |
| caenorhabditis_elegans_prjna13758 | WBGene00021845 | I | 3031212 | 3032938 | -1 | Tsp_11644 | Polr2g | 1478439 | GL622792 | 1478957 |
| caenorhabditis_elegans_prjna13758 | WBGene00013196 | II | 14798472 | 14800971 | 1 | Tsp_08058 | | 1661691 | GL622789 | 1662674 |
| caenorhabditis_elegans_prjna13758 | WBGene00013200 | I | 14707059 | 14707875 | -1 | Tsp_01825 | Pbdc1 | 7401206 | GL622787 | 7401862 |
| caenorhabditis_elegans_prjna13758 | WBGene00013219 | II | 14349783 | 14350795 | 1 | Tsp_00542 | | 2024885 | GL622787 | 2025323 |
| caenorhabditis_elegans_prjna13758 | WBGene00000898 | III | 2995487 | 3040860 | -1 | Tsp_12077 | | 2107174 | GL622792 | 2113811 |
| caenorhabditis_elegans_prjna13758 | WBGene00000898 | III | 2995487 | 3040860 | -1 | Tsp_06967 | | 8699073 | GL622785 | 8703440 |
| caenorhabditis_elegans_prjna13758 | WBGene00021929 | IV | 1052524 | 1054794 | -1 | Tsp_10887 | Dcp1a | 329192 | GL623868 | 330995 |
| caenorhabditis_elegans_prjna13758 | WBGene00000816 | IV | 1077463 | 1087013 | -1 | Tsp_07915 | | 11304115 | GL622785 | 11305845 |
| caenorhabditis_elegans_prjna13758 | WBGene00021956 | V | 7060811 | 7062695 | 1 | Tsp_03377 | | 4015733 | GL622785 | 4019537 |
| caenorhabditis_elegans_prjna13758 | WBGene00013319 | IV | 14919610 | 14920421 | 1 | Tsp_12414 | | 8096 | GL624002 | 8753 |
| caenorhabditis_elegans_prjna13758 | WBGene00013319 | IV | 14919610 | 14920421 | 1 | Tsp_08560 | zfp36l2 | 4699442 | GL622788 | 4700099 |
| caenorhabditis_elegans_prjna13758 | WBGene00013319 | IV | 14919610 | 14920421 | 1 | Tsp_08559 | Zfp36l3 | 4695882 | GL622788 | 4696511 |
| caenorhabditis_elegans_prjna13758 | WBGene00013319 | IV | 14919610 | 14920421 | 1 | Tsp_12415 | | 11512 | GL624002 | 12169 |
| caenorhabditis_elegans_prjna13758 | WBGene00013319 | IV | 14919610 | 14920421 | 1 | Tsp_12413.2 | | 5119 | GL624002 | 5437 |
| caenorhabditis_elegans_prjna13758 | WBGene00013319 | IV | 14919610 | 14920421 | 1 | Tsp_08561 | | 4702854 | GL622788 | 4703511 |
| caenorhabditis_elegans_prjna13758 | WBGene00022002 | IV | 5241364 | 5241794 | -1 | Tsp_00768 | | 3048065 | GL622787 | 3048698 |
| caenorhabditis_elegans_prjna13758 | WBGene00022002 | IV | 5241364 | 5241794 | -1 | Tsp_00761 | | 3076026 | GL622787 | 3077501 |
| caenorhabditis_elegans_prjna13758 | WBGene00022002 | IV | 5241364 | 5241794 | -1 | Tsp_00756 | | 2908432 | GL622787 | 2908941 |
| caenorhabditis_elegans_prjna13758 | WBGene00022002 | IV | 5241364 | 5241794 | -1 | Tsp_13425 | | 585 | GL623126 | 965 |
| caenorhabditis_elegans_prjna13758 | WBGene00022002 | IV | 5241364 | 5241794 | -1 | Tsp_13917 | | 820 | GL624354 | 1264 |
| caenorhabditis_elegans_prjna13758 | WBGene00012385 | IV | 11076085 | 11078582 | -1 | Tsp_09190 | | 4853799 | GL622788 | 4854471 |
| caenorhabditis_elegans_prjna13758 | WBGene00022021 | V | 4607039 | 4615817 | -1 | Tsp_03461 | | 4326372 | GL622785 | 4335227 |
| caenorhabditis_elegans_prjna13758 | WBGene00022025 | I | 654090 | 658810 | 1 | Tsp_11355 | | 7171282 | GL622792 | 7174505 |
| caenorhabditis_elegans_prjna13758 | WBGene00022037 | I | 511052 | 519800 | -1 | Tsp_06708 | | 7499688 | GL622785 | 7504626 |

| | | | | | | | | | | |
|---|---|---|---|---|---|---|---|---|---|---|
| caenorhabditis_elegans_prjna13758 | WBGene00022042 | I | 535791 | 536670 | 1 | Tsp_00702 | | 2675235 | GL622787 | 2677852 |
| caenorhabditis_elegans_prjna13758 | WBGene00012391 | I | 13746673 | 13750483 | 1 | Tsp_11383 | | 7327147 | GL622792 | 7328601 |
| caenorhabditis_elegans_prjna13758 | WBGene00012402 | V | 15726841 | 15728353 | 1 | Tsp_11550 | | 735716 | GL622790 | 741190 |
| caenorhabditis_elegans_prjna13758 | WBGene00012402 | V | 15726841 | 15728353 | 1 | Tsp_12889 | | 454 | GL626579 | 1719 |
| caenorhabditis_elegans_prjna13758 | WBGene00022126 | I | 2731562 | 2735371 | 1 | Tsp_10027 | | 94668 | GL622789 | 98251 |
| caenorhabditis_elegans_prjna13758 | WBGene00004920 | I | 2727014 | 2727545 | 1 | Tsp_03714 | snr-7 | 215870 | GL622788 | 216325 |
| caenorhabditis_elegans_prjna13758 | WBGene00022128 | I | 2721879 | 2726724 | -1 | Tsp_09776 | | 6291739 | GL622789 | 6296265 |
| caenorhabditis_elegans_prjna13758 | WBGene00004043 | I | 2755540 | 2758204 | -1 | Tsp_02561 | | 467619 | GL622785 | 471249 |
| caenorhabditis_elegans_prjna13758 | WBGene00022192 | III | 2623170 | 2627771 | 1 | Tsp_01402 | PHAX | 5820536 | GL622787 | 5822576 |
| caenorhabditis_elegans_prjna13758 | WBGene00001232 | I | 2485867 | 2489780 | 1 | Tsp_07361 | | 1018445 | GL622784 | 1019613 |
| caenorhabditis_elegans_prjna13758 | WBGene00013550 | III | 12216779 | 12218634 | -1 | Tsp_03194 | | 3204042 | GL622785 | 3205360 |
| caenorhabditis_elegans_prjna13758 | WBGene00000546 | IV | 1431272 | 1435039 | -1 | Tsp_13576 | | 4450 | GL629167 | 5113 |
| caenorhabditis_elegans_prjna13758 | WBGene00001258 | V | 18860287 | 18894974 | -1 | Tsp_05465 | | 3450375 | GL622792 | 3483209 |
| caenorhabditis_elegans_prjna13758 | WBGene00001258 | V | 18860287 | 18894974 | -1 | Tsp_05466 | | 3450172 | GL622792 | 3450309 |
| caenorhabditis_elegans_prjna13758 | WBGene00013857 | IV | 10732832 | 10734632 | -1 | Tsp_09564 | | 1025525 | GL622792 | 1029397 |
| caenorhabditis_elegans_prjna13758 | WBGene00013857 | IV | 10732832 | 10734632 | -1 | Tsp_09563 | | 1030763 | GL622792 | 1032469 |
| caenorhabditis_elegans_prjna13758 | WBGene00000865 | IV | 10735259 | 10737177 | 1 | Tsp_05726 | ccnb2 | 4824667 | GL622792 | 4826249 |
| caenorhabditis_elegans_prjna13758 | WBGene00003405 | V | 10733931 | 10736892 | -1 | Tsp_12012 | | 14229 | GL625294 | 17458 |
| caenorhabditis_elegans_prjna13758 | WBGene00013877 | V | 14185691 | 14192156 | 1 | Tsp_09975 | | 594908 | GL623393 | 604925 |
| caenorhabditis_elegans_prjna13758 | WBGene00004984 | V | 6783187 | 6784922 | -1 | Tsp_05134 | | 2147856 | GL624340 | 2154149 |
| caenorhabditis_elegans_prjna13758 | WBGene00004984 | V | 6783187 | 6784922 | -1 | Tsp_13755 | | 174 | GL626245 | 1096 |
| caenorhabditis_elegans_prjna13758 | WBGene00022603 | V | 6788449 | 6792665 | -1 | Tsp_12597 | | 307 | GL624342 | 2452 |
| caenorhabditis_elegans_prjna13758 | WBGene00022603 | V | 6788449 | 6792665 | -1 | Tsp_04021 | | 1542782 | GL622788 | 1547914 |
| caenorhabditis_elegans_prjna13758 | WBGene00022617 | IV | 7108221 | 7109912 | 1 | Tsp_01125 | | 4353662 | GL622787 | 4354442 |
| caenorhabditis_elegans_prjna13758 | WBGene00022629 | V | 8035111 | 8039506 | 1 | Tsp_15122 | | 143 | GL626550 | 1214 |
| caenorhabditis_elegans_prjna13758 | WBGene00003865 | V | 8029439 | 8031214 | 1 | Tsp_12414 | | 8096 | GL624002 | 8753 |

| | | | | | | | | | | |
|---|---|---|---|---|---|---|---|---|---|---|
| caenorhabditis_elegans_prjna13758 | WBGene00003865 | V | 8029439 | 8031214 | 1 | Tsp_08560 | zfp36l2 | 4699442 | GL622788 | 4700099 |
| caenorhabditis_elegans_prjna13758 | WBGene00003865 | V | 8029439 | 8031214 | 1 | Tsp_08559 | Zfp36l3 | 4695882 | GL622788 | 4696511 |
| caenorhabditis_elegans_prjna13758 | WBGene00003865 | V | 8029439 | 8031214 | 1 | Tsp_12415 | | 11512 | GL624002 | 12169 |
| caenorhabditis_elegans_prjna13758 | WBGene00003865 | V | 8029439 | 8031214 | 1 | Tsp_12413.2 | | 5119 | GL624002 | 5437 |
| caenorhabditis_elegans_prjna13758 | WBGene00003865 | V | 8029439 | 8031214 | 1 | Tsp_08561 | | 4702854 | GL622788 | 4703511 |
| caenorhabditis_elegans_prjna13758 | WBGene00022632 | I | 6655608 | 6658371 | 1 | Tsp_05227 | Ttbk1 | 2180440 | GL622792 | 2181339 |
| caenorhabditis_elegans_prjna13758 | WBGene00022632 | I | 6655608 | 6658371 | 1 | Tsp_12209 | | 4940141 | GL622792 | 4941163 |
| caenorhabditis_elegans_prjna13758 | WBGene00022632 | I | 6655608 | 6658371 | 1 | Tsp_11208 | | 5439495 | GL622792 | 5440343 |
| caenorhabditis_elegans_prjna13758 | WBGene00022632 | I | 6655608 | 6658371 | 1 | Tsp_12401 | | 2248 | GL623394 | 3147 |
| caenorhabditis_elegans_prjna13758 | WBGene00022632 | I | 6655608 | 6658371 | 1 | Tsp_12402 | | 5826 | GL623394 | 6369 |
| caenorhabditis_elegans_prjna13758 | WBGene00022632 | I | 6655608 | 6658371 | 1 | Tsp_13461 | | 44 | GL626854 | 695 |
| caenorhabditis_elegans_prjna13758 | WBGene00014176 | III | 12975068 | 12977862 | -1 | Tsp_11677 | | 1652387 | GL622792 | 1653655 |
| caenorhabditis_elegans_prjna13758 | WBGene00014177 | III | 12979162 | 12980163 | 1 | Tsp_01655 | | 6600982 | GL622787 | 6602170 |
| caenorhabditis_elegans_prjna13758 | WBGene00014201 | I | 13238540 | 13240726 | 1 | Tsp_05976 | | 6070604 | GL622785 | 6074765 |
| caenorhabditis_elegans_prjna13758 | WBGene00001872 | II | 7065020 | 7069966 | -1 | Tsp_11619 | MSH4 | 2002538 | GL622792 | 2007277 |
| caenorhabditis_elegans_prjna13758 | WBGene00022849 | II | 7061999 | 7064777 | 1 | Tsp_10497 | | 1167057 | GL622791 | 1168176 |
| caenorhabditis_elegans_prjna13758 | WBGene00022849 | II | 7061999 | 7064777 | 1 | Tsp_10496 | | 1164873 | GL622791 | 1166542 |
| caenorhabditis_elegans_prjna13758 | WBGene00022852 | II | 7050672 | 7052001 | 1 | Tsp_02777 | | 1312024 | GL622785 | 1313226 |
| caenorhabditis_elegans_prjna13758 | WBGene00022854 | II | 7038292 | 7041685 | 1 | Tsp_00990 | | 3763911 | GL622787 | 3769982 |
| caenorhabditis_elegans_prjna13758 | WBGene00044072 | III | 10120781 | 10122928 | 1 | Tsp_00956 | | 3889019 | GL622787 | 3891266 |
| caenorhabditis_elegans_prjna13758 | WBGene00022880 | II | 5823816 | 5827443 | -1 | Tsp_08861 | | 4897332 | GL622789 | 4901350 |
| caenorhabditis_elegans_prjna13758 | WBGene00022884 | II | 5845254 | 5845541 | -1 | Tsp_00768 | | 3048065 | GL622787 | 3048698 |
| caenorhabditis_elegans_prjna13758 | WBGene00022884 | II | 5845254 | 5845541 | -1 | Tsp_00761 | | 3076026 | GL622787 | 3077501 |
| caenorhabditis_elegans_prjna13758 | WBGene00022884 | II | 5845254 | 5845541 | -1 | Tsp_00756 | | 2908432 | GL622787 | 2908941 |
| caenorhabditis_elegans_prjna13758 | WBGene00022884 | II | 5845254 | 5845541 | -1 | Tsp_13425 | | 585 | GL623126 | 965 |
| caenorhabditis_elegans_prjna13758 | WBGene00022884 | II | 5845254 | 5845541 | -1 | Tsp_13917 | | 820 | GL624354 | 1264 |

| | | | | | | | | | |
|---|---|---|---|---|---|---|---|---|---|
| caenorhabditis_elegans_prjna13758 | WBGene00003457 | II | 5809917 | 5810342 | -1 | Tsp_00768 | 3048065 | GL622787 | 3048698 |
| caenorhabditis_elegans_prjna13758 | WBGene00003457 | II | 5809917 | 5810342 | -1 | Tsp_00761 | 3076026 | GL622787 | 3077501 |
| caenorhabditis_elegans_prjna13758 | WBGene00003457 | II | 5809917 | 5810342 | -1 | Tsp_00756 | 2908432 | GL622787 | 2908941 |
| caenorhabditis_elegans_prjna13758 | WBGene00003457 | II | 5809917 | 5810342 | -1 | Tsp_13425 | 585 | GL623126 | 965 |
| caenorhabditis_elegans_prjna13758 | WBGene00003457 | II | 5809917 | 5810342 | -1 | Tsp_13917 | 820 | GL624354 | 1264 |
| caenorhabditis_elegans_prjna13758 | WBGene00014243 | IV | 9696989 | 9706723 | -1 | Tsp_03111 | 2810811 | GL622785 | 2818818 |
| caenorhabditis_elegans_prjna13758 | WBGene00005026 | II | 9647887 | 9649367 | 1 | Tsp_07764 | 10629322 | GL622785 | 10630280 |
| caenorhabditis_elegans_prjna13758 | WBGene00014250 | II | 9649491 | 9650959 | 1 | Tsp_04920 | 1480361 | GL624340 | 1482288 |
| caenorhabditis_elegans_prjna13758 | WBGene00001511 | II | 5501633 | 5503908 | 1 | Tsp_12087 | 366340 | GL622791 | 368652 |
| caenorhabditis_elegans_prjna13758 | WBGene00013924 | II | 11651261 | 11652906 | -1 | Tsp_13625 | 78 | GL629375 | 1028 |
| caenorhabditis_elegans_prjna13758 | WBGene00013924 | II | 11651261 | 11652906 | -1 | Tsp_10709 | 2874046 | GL622789 | 2878218 |
| caenorhabditis_elegans_prjna13758 | WBGene00013925 | II | 11649096 | 11651213 | -1 | Tsp_11373 | 7246704 | GL622792 | 7247524 |
| caenorhabditis_elegans_prjna13758 | WBGene00004468 | II | 11653096 | 11654194 | 1 | Tsp_00908 | 3653333 | GL622787 | 3654809 |
| caenorhabditis_elegans_prjna13758 | WBGene00001166 | III | 6009550 | 6012839 | 1 | Tsp_01994 | 8059960 | GL622787 | 8064520 |
| caenorhabditis_elegans_prjna13758 | WBGene00022694 | III | 6006344 | 6009084 | -1 | Tsp_00755 | 2910337 | GL622787 | 2914521 |
| caenorhabditis_elegans_prjna13758 | WBGene00022696 | III | 6032696 | 6039475 | -1 | Tsp_01556 | 6326754 | GL622787 | 6335621 |
| caenorhabditis_elegans_prjna13758 | WBGene00022703 | III | 8403715 | 8405562 | -1 | Tsp_01730 | 7062934 | GL622787 | 7064886 |
| caenorhabditis_elegans_prjna13758 | WBGene00003458 | IV | 5316487 | 5316944 | 1 | Tsp_00768 | 3048065 | GL622787 | 3048698 |
| caenorhabditis_elegans_prjna13758 | WBGene00003458 | IV | 5316487 | 5316944 | 1 | Tsp_00761 | 3076026 | GL622787 | 3077501 |
| caenorhabditis_elegans_prjna13758 | WBGene00003458 | IV | 5316487 | 5316944 | 1 | Tsp_00756 | 2908432 | GL622787 | 2908941 |
| caenorhabditis_elegans_prjna13758 | WBGene00003458 | IV | 5316487 | 5316944 | 1 | Tsp_13425 | 585 | GL623126 | 965 |
| caenorhabditis_elegans_prjna13758 | WBGene00003458 | IV | 5316487 | 5316944 | 1 | Tsp_13917 | 820 | GL624354 | 1264 |
| caenorhabditis_elegans_prjna13758 | WBGene00003452 | IV | 5312693 | 5313149 | -1 | Tsp_00768 | 3048065 | GL622787 | 3048698 |
| caenorhabditis_elegans_prjna13758 | WBGene00003452 | IV | 5312693 | 5313149 | -1 | Tsp_00761 | 3076026 | GL622787 | 3077501 |
| caenorhabditis_elegans_prjna13758 | WBGene00003452 | IV | 5312693 | 5313149 | -1 | Tsp_00756 | 2908432 | GL622787 | 2908941 |
| caenorhabditis_elegans_prjna13758 | WBGene00003452 | IV | 5312693 | 5313149 | -1 | Tsp_13425 | 585 | GL623126 | 965 |

| | | | | | | | | | | |
|---|---|---|---|---|---|---|---|---|---|---|
| caenorhabditis_elegans_prjna13758 | WBGene00003452 | IV | 5312693 | 5313149 | -1 | Tsp_13917 | | 820 | GL624354 | 1264 |
| caenorhabditis_elegans_prjna13758 | WBGene00003468 | IV | 5297927 | 5298380 | 1 | Tsp_00768 | | 3048065 | GL622787 | 3048698 |
| caenorhabditis_elegans_prjna13758 | WBGene00003468 | IV | 5297927 | 5298380 | 1 | Tsp_00761 | | 3076026 | GL622787 | 3077501 |
| caenorhabditis_elegans_prjna13758 | WBGene00003468 | IV | 5297927 | 5298380 | 1 | Tsp_00756 | | 2908432 | GL622787 | 2908941 |
| caenorhabditis_elegans_prjna13758 | WBGene00003468 | IV | 5297927 | 5298380 | 1 | Tsp_13425 | | 585 | GL623126 | 965 |
| caenorhabditis_elegans_prjna13758 | WBGene00003468 | IV | 5297927 | 5298380 | 1 | Tsp_13917 | | 820 | GL624354 | 1264 |
| caenorhabditis_elegans_prjna13758 | WBGene00003444 | IV | 5294922 | 5295375 | -1 | Tsp_00768 | | 3048065 | GL622787 | 3048698 |
| caenorhabditis_elegans_prjna13758 | WBGene00003444 | IV | 5294922 | 5295375 | -1 | Tsp_00761 | | 3076026 | GL622787 | 3077501 |
| caenorhabditis_elegans_prjna13758 | WBGene00003444 | IV | 5294922 | 5295375 | -1 | Tsp_00756 | | 2908432 | GL622787 | 2908941 |
| caenorhabditis_elegans_prjna13758 | WBGene00003444 | IV | 5294922 | 5295375 | -1 | Tsp_13425 | | 585 | GL623126 | 965 |
| caenorhabditis_elegans_prjna13758 | WBGene00003444 | IV | 5294922 | 5295375 | -1 | Tsp_13917 | | 820 | GL624354 | 1264 |
| caenorhabditis_elegans_prjna13758 | WBGene00022707 | IV | 5295931 | 5297487 | -1 | Tsp_05227 | Ttbk1 | 2180440 | GL622792 | 2181339 |
| caenorhabditis_elegans_prjna13758 | WBGene00022707 | IV | 5295931 | 5297487 | -1 | Tsp_12209 | | 4940141 | GL622792 | 4941163 |
| caenorhabditis_elegans_prjna13758 | WBGene00022707 | IV | 5295931 | 5297487 | -1 | Tsp_11208 | | 5439495 | GL622792 | 5440343 |
| caenorhabditis_elegans_prjna13758 | WBGene00022707 | IV | 5295931 | 5297487 | -1 | Tsp_12401 | | 2248 | GL623394 | 3147 |
| caenorhabditis_elegans_prjna13758 | WBGene00022707 | IV | 5295931 | 5297487 | -1 | Tsp_12402 | | 5826 | GL623394 | 6369 |
| caenorhabditis_elegans_prjna13758 | WBGene00022707 | IV | 5295931 | 5297487 | -1 | Tsp_13461 | | 44 | GL626854 | 695 |
| caenorhabditis_elegans_prjna13758 | WBGene00022710 | IV | 5306599 | 5308146 | -1 | Tsp_06387 | | 10716569 | GL622787 | 10717856 |
| caenorhabditis_elegans_prjna13758 | WBGene00022710 | IV | 5306599 | 5308146 | -1 | Tsp_13670 | | 166 | GL627984 | 1026 |
| caenorhabditis_elegans_prjna13758 | WBGene00022710 | IV | 5306599 | 5308146 | -1 | Tsp_11877 | | 139964 | GL622790 | 141286 |
| caenorhabditis_elegans_prjna13758 | WBGene00022710 | IV | 5306599 | 5308146 | -1 | Tsp_13257 | | 551 | GL628954 | 1584 |
| caenorhabditis_elegans_prjna13758 | WBGene00022710 | IV | 5306599 | 5308146 | -1 | Tsp_01419 | | 5756161 | GL622787 | 5757550 |
| caenorhabditis_elegans_prjna13758 | WBGene00022710 | IV | 5306599 | 5308146 | -1 | Tsp_06318.1 | | 10514671 | GL622787 | 10515097 |
| caenorhabditis_elegans_prjna13758 | WBGene00022710 | IV | 5306599 | 5308146 | -1 | Tsp_12521 | | 1192 | GL627057 | 2036 |
| caenorhabditis_elegans_prjna13758 | WBGene00022710 | IV | 5306599 | 5308146 | -1 | Tsp_06319.2 | | 10512081 | GL622787 | 10512342 |
| caenorhabditis_elegans_prjna13758 | WBGene00022710 | IV | 5306599 | 5308146 | -1 | Tsp_13571 | | 383 | GL628396 | 1344 |

| | | | | | | | | | | |
|---|---|---|---|---|---|---|---|---|---|---|
| caenorhabditis_elegans_prjna13758 | WBGene00022739 | II | 4425167 | 4432613 | 1 | Tsp_13147.1 | | 3784 | GL627412 | 5245 |
| caenorhabditis_elegans_prjna13758 | WBGene00000878 | III | 13692668 | 13693811 | 1 | Tsp_08266 | | 3565239 | GL622788 | 3566924 |
| caenorhabditis_elegans_prjna13758 | WBGene00000878 | III | 13692668 | 13693811 | 1 | Tsp_12763 | | 195 | GL623251 | 1038 |
| caenorhabditis_elegans_prjna13758 | WBGene00007012 | II | 4942107 | 4943949 | -1 | Tsp_05745 | MED4 | 5197978 | GL622785 | 5198903 |
| caenorhabditis_elegans_prjna13758 | WBGene00022765 | II | 4944042 | 4945838 | -1 | Tsp_07070 | | 9254796 | GL622785 | 9255221 |
| caenorhabditis_elegans_prjna13758 | WBGene00022765 | II | 4944042 | 4945838 | -1 | Tsp_07069 | | 9251286 | GL622785 | 9254773 |
| caenorhabditis_elegans_prjna13758 | WBGene00006619 | II | 4948154 | 4949527 | -1 | Tsp_01988 | | 8087677 | GL622787 | 8088330 |
| caenorhabditis_elegans_prjna13758 | WBGene00022762 | II | 4935563 | 4937956 | 1 | Tsp_03248 | | 3453015 | GL622785 | 3458565 |
| caenorhabditis_elegans_prjna13758 | WBGene00003470 | II | 4932991 | 4933450 | 1 | Tsp_00768 | | 3048065 | GL622787 | 3048698 |
| caenorhabditis_elegans_prjna13758 | WBGene00003470 | II | 4932991 | 4933450 | 1 | Tsp_00761 | | 3076026 | GL622787 | 3077501 |
| caenorhabditis_elegans_prjna13758 | WBGene00003470 | II | 4932991 | 4933450 | 1 | Tsp_00756 | | 2908432 | GL622787 | 2908941 |
| caenorhabditis_elegans_prjna13758 | WBGene00003470 | II | 4932991 | 4933450 | 1 | Tsp_13425 | | 585 | GL623126 | 965 |
| caenorhabditis_elegans_prjna13758 | WBGene00003470 | II | 4932991 | 4933450 | 1 | Tsp_13917 | | 820 | GL624354 | 1264 |
| caenorhabditis_elegans_prjna13758 | WBGene00013998 | IV | 17255301 | 17257562 | 1 | Tsp_03661 | | 31464 | GL622788 | 33027 |
| caenorhabditis_elegans_prjna13758 | WBGene00003158 | III | 9794367 | 9798643 | 1 | Tsp_15939 | MCM7 | 765 | GL626773 | 1142 |
| caenorhabditis_elegans_prjna13758 | WBGene00000189 | III | 9823242 | 9824132 | 1 | Tsp_00680 | | 2764513 | GL622787 | 2765975 |
| caenorhabditis_elegans_prjna13758 | WBGene00000388 | III | 8916351 | 8917929 | -1 | Tsp_13015.2 | | 3440 | GL624006 | 4762 |
| caenorhabditis_elegans_prjna13758 | WBGene00000388 | III | 8916351 | 8917929 | -1 | Tsp_13015.1 | | 4885 | GL624006 | 5857 |
| caenorhabditis_elegans_prjna13758 | WBGene00002998 | III | 8900237 | 8904160 | -1 | Tsp_00779 | | 3003601 | GL622787 | 3006181 |
| caenorhabditis_elegans_prjna13758 | WBGene00014034 | III | 8956459 | 8957731 | 1 | Tsp_05426 | | 3233283 | GL622792 | 3233950 |
| caenorhabditis_elegans_prjna13758 | WBGene00004918 | III | 7862070 | 7862577 | 1 | Tsp_11073 | | 2103313 | GL622784 | 2103615 |
| caenorhabditis_elegans_prjna13758 | WBGene00022794 | III | 7769165 | 7770149 | -1 | Tsp_12911 | | 920 | GL623940 | 1583 |
| caenorhabditis_elegans_prjna13758 | WBGene00022794 | III | 7769165 | 7770149 | -1 | Tsp_09103 | | 3145942 | GL622784 | 3147332 |
| caenorhabditis_elegans_prjna13758 | WBGene00022794 | III | 7769165 | 7770149 | -1 | Tsp_13434 | | 171 | GL624286 | 529 |
| caenorhabditis_elegans_prjna13758 | WBGene00002078 | V | 7805064 | 7814656 | 1 | Tsp_13954 | | 179 | GL628336 | 1739 |
| caenorhabditis_elegans_prjna13758 | WBGene00002078 | V | 7805064 | 7814656 | 1 | Tsp_03075 | Xpo1 | 2598395 | GL622785 | 2603924 |

| | | | | | | | | | | |
|---|---|---|---|---|---|---|---|---|---|---|
| caenorhabditis_elegans_prjna13758 | WBGene00014083 | IV | 12559365 | 12560719 | 1 | Tsp_06916 | lmp4 | 8472387 | GL622785 | 8474298 |
| caenorhabditis_elegans_prjna13758 | WBGene00006491 | IV | 11650715 | 11652033 | -1 | Tsp_06712 | | 7521599 | GL622785 | 7522854 |
| caenorhabditis_elegans_prjna13758 | WBGene00001320 | IV | 11653422 | 11655470 | -1 | Tsp_07410 | SLC29A1 | 1218333 | GL622784 | 1221797 |
| caenorhabditis_elegans_prjna13758 | WBGene00006566 | IV | 11957547 | 11959804 | -1 | Tsp_06742 | qtrt1 | 7682056 | GL622785 | 7683321 |
| caenorhabditis_elegans_prjna13758 | WBGene00014111 | V | 10205683 | 10206602 | -1 | Tsp_10965 | | 9548975 | GL622785 | 9550199 |
| caenorhabditis_elegans_prjna13758 | WBGene00003209 | I | 9132538 | 9135753 | 1 | Tsp_10426 | | 734334 | GL622791 | 743797 |
| caenorhabditis_elegans_prjna13758 | WBGene00014120 | I | 9141971 | 9144376 | -1 | Tsp_00570 | | 2358356 | GL622787 | 2359589 |
| caenorhabditis_elegans_prjna13758 | WBGene00022836 | I | 4379325 | 4381948 | -1 | Tsp_05307 | | 2709213 | GL622792 | 2711039 |
| caenorhabditis_elegans_prjna13758 | WBGene00022832 | I | 4368089 | 4370504 | 1 | Tsp_08181 | | 2258587 | GL622789 | 2260403 |

**Total Tsp germline genes:** **1332**

| Genes overlapping operons | % of total | Annotations | Expressed genes | Operonic genes | % of expressed | P-values Chi-square | P-values Binomial |
|---|---|---|---|---|---|---|---|
| 245 | 18% | Operons_BRAKER_BAM.featureCounts.10bp.exon.conv.gff3 | 14903 | 758 | 5.09% | 2.88E-108 | 1.27E-67 |
| 212 | 16% | Operons_BRAKER_BAM.featureCounts.10bp.gene.gff3 | 14561 | 622 | 4.17% | 4.72E-98 | 3.76E-60 |
| 250 | 19% | Operons_BRAKER_BAM.featureCounts.8bp.exon.conv.gff3 | 15013 | 795 | 5.33% | 7.26E-107 | 1.98E-67 |
| 209 | 16% | Operons_BRAKER_BAM.featureCounts.8bp.gene.gff3 | 14561 | 630 | 4.23% | 2.26E-92 | 2.16E-57 |
| 246 | 18% | Operons_BRAKER_TRINITY.featureCounts.10bp.exon.conv.gff3 | 17925 | 752 | 5.05% | 7.57E-149 | 1.93E-84 |
| 227 | 17% | Operons_BRAKER_TRINITY.featureCounts.10bp.gene.gff3 | 17633 | 637 | 4.27% | 4.68E-152 | 4.20E-83 |
| 252 | 19% | Operons_BRAKER_TRINITY.featureCounts.8bp.exon.conv.gff3 | 18031 | 791 | 5.31% | 7.29E-148 | 8.16E-85 |
| 220 | 17% | Operons_BRAKER_TRINITY.featureCounts.8bp.gene.gff3 | 17633 | 650 | 4.36% | 2.50E-136 | 2.04E-76 |
| 338 | 25% | Operons_DENOVO_ALL.featureCounts.10bp.exon.conv.gff3 | 18119 | 792 | 5.31% | 1.12E-307 | 2.25E-153 |
| 207 | 16% | Operons_DENOVO_ALL.featureCounts.10bp.gene.gff3 | 17244 | 439 | 2.95% | 3.62E-199 | 6.91E-95 |
| 360 | 27% | Operons_DENOVO_ALL.featureCounts.8bp.exon.conv.gff3 | 18347 | 845 | 5.67% | 0.00E+00 | 7.79E-166 |
| 207 | 16% | Operons_DENOVO_ALL.featureCounts.8bp.gene.gff3 | 17244 | 433 | 2.91% | 6.73E-203 | 5.96E-96 |
| 187 | 14% | Operons_PRJNA12603.featureCounts.10bp.exon.conv.gff3 | 15273 | 608 | 4.08% | 1.19E-78 | 1.96E-49 |
| 148 | 11% | Operons_PRJNA12603.featureCounts.10bp.gene.gff3 | 15024 | 491 | 3.29% | 2.59E-58 | 3.16E-37 |
| 191 | 14% | Operons_PRJNA12603.featureCounts.8bp.exon.conv.gff3 | 15364 | 645 | 4.33% | 4.70E-76 | 1.50E-48 |
| 152 | 11% | Operons_PRJNA12603.featureCounts.8bp.gene.gff3 | 15025 | 507 | 3.40% | 2.45E-59 | 5.56E-38 |
| 312 | 23% | Operons_STRINGTIE.featureCounts.10bp.exon.conv.gff3 | 13677 | 706 | 4.74% | 2.47E-199 | 2.03E-112 |
| 212 | 16% | Operons_STRINGTIE.featureCounts.10bp.gene.gff3 | 12973 | 421 | 2.82% | 3.91E-150 | 2.68E-80 |
| 328 | 25% | Operons_STRINGTIE.featureCounts.8bp.exon.conv.gff3 | 13866 | 752 | 5.05% | 3.19E-210 | 8.07E-119 |
| 209 | 16% | Operons_STRINGTIE.featureCounts.8bp.gene.gff3 | 12973 | 407 | 2.73% | 3.08E-152 | 1.40E-80 |

BEDTOOLS Intersect
