## Supplemental Table S3 for "Deep evolutionary origin of nematode SL2 *trans*-splicing revealed by genome-wide analysis of the *Trichinella spiralis* transcriptome"

**Supplementary Table 3. Detailed summaries of genome annotation approaches (reference annotation + four *de novo* annotations), gene classification by *Tsp*-SL trans-splicing events and operon prediction across all 20 datasets.** Numbers of reads and genes in particular classes are presented alongside numbers and sizes of predicted operons and results of Welch's *t*-tests among datasets grouped by annotation type, *Tsp*-SL read-screening stringency and exon-based annotation correction.

### Operon Identification Overview

**Gene annotation summary**

| Genome annotation | SL screening stringency | Genes | Corrected genes | Genes gained |
|---|---|---|---|---|
| BRAKER | 10 bp | 16,312 | 16,657 | 345 |
| BRAKER | 8 bp | 16,312 | 16,767 | 455 |
| BRAKER +TRINITY | 10 bp | 20,083 | 20,381 | 298 |
| BRAKER +TRINITY | 8 bp | 20,083 | 20,487 | 404 |
| DENOVO_ALL | 10 bp | 19,573 | 20,448 | 875 |
| DENOVO_ALL | 8 bp | 19,573 | 20,676 | 1,103 |
| REFERENCE | 10 bp | 16,380 | 16,630 | 250 |
| REFERENCE | 8 bp | 16,380 | 16,720 | 340 |
| STRINGTIE | 10 bp | 13,060 | 13,765 | 705 |
| STRINGTIE | 8 bp | 13,060 | 13,954 | 894 |

**SL reads summary**

| Genome annotation | SL screening stringency | Annotation correction | SL1 reads | SL2 reads | Genes | Expressed genes | Trans-spliced genes | SL1 genes | SL2 genes | SL1+SL2 genes | No SL genes | Operons | Operonic genes | Operon sizes | | | | |
|---|---|---|---|---|---|---|---|---|---|---|---|---|---|---|---|---|---|---|
| | | | | | | | | | | | | | | 1 | 2 | 3 | 4 | 5 |
| BRAKER | 10 bp | corrected | 238,144 | 47,900 | 16,657 | 14,903 | 2,615 | 1,577 | 399 | 639 | 14,042 | 362 | 758 | 3 | 328 | 26 | 4 | 1 |

| | | | | | | | | | | | | | | | | | | |
|---|---|---|---|---|---|---|---|---|---|---|---|---|---|---|---|---|---|---|
| BRAKER | 10 bp | raw | 239,278 | 48,108 | 16,312 | 14,561 | 2,454 | 1,432 | 323 | 699 | 13,858 | 302 | 622 | 3 | 281 | 16 | 1 | 1 |
| BRAKER | 8 bp | corrected | 349,571 | 66,675 | 16,767 | 15,013 | 2,981 | 1,598 | 419 | 964 | 13,786 | 380 | 795 | 4 | 344 | 26 | 5 | 1 |
| BRAKER | 8 bp | raw | 351,226 | 67,047 | 16,312 | 14,561 | 2,779 | 1,433 | 328 | 1,018 | 13,533 | 306 | 630 | 4 | 283 | 17 | 1 | 1 |
| BRAKER +TRINITY | 10 bp | corrected | 240,225 | 47,957 | 20,381 | 17,925 | 2,593 | 1,556 | 397 | 640 | 17,788 | 358 | 752 | 3 | 322 | 28 | 4 | 1 |
| BRAKER +TRINITY | 10 bp | raw | 241,289 | 48,166 | 20,083 | 17,633 | 2,461 | 1,433 | 334 | 694 | 17,622 | 306 | 637 | 3 | 277 | 24 | 2 | |
| BRAKER +TRINITY | 8 bp | corrected | 352,023 | 66,726 | 20,487 | 18,031 | 2,957 | 1,573 | 417 | 967 | 17,530 | 378 | 791 | 4 | 342 | 26 | 5 | 1 |
| BRAKER +TRINITY | 8 bp | raw | 353,586 | 67,085 | 20,083 | 17,633 | 2,788 | 1,429 | 341 | 1,018 | 17,295 | 313 | 650 | 4 | 283 | 24 | 2 | |
| DENOVO_ALL | 10 bp | corrected | 264,320 | 52,973 | 20,448 | 18,119 | 2,922 | 1,746 | 416 | 760 | 17,526 | 382 | 792 | 6 | 345 | 28 | 3 | |
| DENOVO_ALL | 10 bp | raw | 266,192 | 53,320 | 19,573 | 17,244 | 2,417 | 1,336 | 225 | 856 | 17,156 | 220 | 439 | 6 | 209 | 5 | | |
| DENOVO_ALL | 8 bp | corrected | 386,471 | 73,704 | 20,676 | 18,347 | 3,362 | 1,807 | 444 | 1,111 | 17,314 | 408 | 845 | 7 | 368 | 30 | 3 | |
| DENOVO_ALL | 8 bp | raw | 388,839 | 74,208 | 19,573 | 17,244 | 2,709 | 1,307 | 223 | 1,179 | 16,864 | 216 | 433 | 6 | 203 | 7 | | |
| REFERENCE | 10 bp | corrected | 216,267 | 41,529 | 16,630 | 15,273 | 2,228 | 1,353 | 314 | 561 | 14,402 | 296 | 608 | 6 | 313 | 22 | 2 | |
| REFERENCE | 10 bp | raw | 217,007 | 41,707 | 16,380 | 15,024 | 2,112 | 1,255 | 250 | 607 | 14,268 | 243 | 491 | 6 | 197 | 7 | | |
| REFERENCE | 8 bp | corrected | 317,337 | 58,129 | 16,720 | 15,364 | 2,571 | 1,398 | 334 | 839 | 14,149 | 313 | 645 | 7 | 331 | 25 | 2 | |
| REFERENCE | 8 bp | raw | 318,358 | 58,387 | 16,380 | 15,025 | 2,417 | 1,280 | 259 | 878 | 13,963 | 250 | 507 | 6 | 190 | 7 | | |
| STRINGTIE | 10 bp | corrected | 254,819 | 50,309 | 13,765 | 13,677 | 2,755 | 1,658 | 369 | 728 | 11,010 | 343 | 706 | 2 | 277 | 16 | 1 | |
| STRINGTIE | 10 bp | raw | 256,550 | 50,632 | 13,060 | 12,973 | 2,370 | 1,343 | 217 | 810 | 10,690 | 210 | 421 | 2 | 234 | 7 | | |
| STRINGTIE | 8 bp | corrected | 372,570 | 70,140 | 13,954 | 13,866 | 3,169 | 1,709 | 394 | 1,066 | 10,785 | 365 | 752 | 2 | 291 | 19 | 1 | |
| STRINGTIE | 8 bp | raw | 374,657 | 70,587 | 13,060 | 12,973 | 2,656 | 1,313 | 210 | 1,133 | 10,404 | 203 | 407 | 2 | 239 | 9 | | |

### Statistical Tests

| | SL1 reads | SL2 reads | Total genes | Expressed genes | Trans-spliced genes | SL1 genes | SL2 genes | SL1+SL2 genes | No SL genes | Operonic genes | Operons |
|---|---|---|---|---|---|---|---|---|---|---|---|
| Total | 299,936 ± 61,680 | 57,764 ± 10,821 | 17,365 ± 2,630 | 15,769 ± 1,816 | 2,666 ± 311 | 1,477 ± 165 | 331 ± 77 | 858 ± 190 | 14,699 ± 2,578 | 634 ± 142 | 308 ± 65 |
| **Genome annotation** | | | | | | | | | | | |
| de novo | 308,110 ± 61,479 | 59,721 ± 10,461 | 17,574 ± 2,920 | 15,919 ± 2,012 | 2,749 ± 278 | 1,516 ± 160 | 341 ± 81 | 893 ± 186 | 14,825 ± 2,886 | 652 ± 150 | 316 ± 69 |
| reference | 267,242 ± 58,436 | 49,938 ± 9,608 | 16,528 ± 174 | 15,172 ± 174 | 2,332 ± 203 | 1,322 ± 66 | 289 ± 41 | 721 ± 160 | 14,196 ± 186 | 563 ± 75 | 276 ± 34 |
| P-value (Welch's t-test) | 0.273 | 0.134 | 0.174 | 0.163 | 0.014 | 0.003 | 0.104 | 0.121 | 0.4 | 0.125 | 0.129 |
| **TSL screening stringency** | | | | | | | | | | | |
| 10 bp | 243,409 ± 17,488 | 48,260 ± 4,030 | 17,329 ± 2,697 | 15,733 ± 1,856 | 2,493 ± 240 | 1,469 ± 159 | 324 ± 73 | 699 ± 92 | 14,836 ± 2,646 | 623 ± 135 | 302 ± 61 |
| 8 bp | 356,464 ± 25,000 | 67,269 ± 5,497 | 17,401 ± 2,708 | 15,806 ± 1,873 | 2,839 ± 283 | 1,485 ± 180 | 337 ± 84 | 1,017 ± 109 | 14,562 ± 2,644 | 646 ± 155 | 313 ± 71 |
| P-value (Welch's t-test) | 0 | 0 | 0.953 | 0.932 | 0.009 | 0.837 | 0.727 | 0 | 0.819 | 0.728 | 0.716 |
| **Annotation correction** | | | | | | | | | | | |
| corrected | 299,175 ± 63,180 | 57,604 ± 11,068 | 17,649 ± 2,686 | 16,052 ± 1,852 | 2,815 ± 330 | 1,598 ± 143 | 390 ± 40 | 828 ± 192 | 14,833 ± 2,641 | 744 ± 73 | 358 ± 34 |
| raw | 300,698 ± 63,549 | 57,925 ± 11,165 | 17,082 ± 2,686 | 15,487 ± 1,831 | 2,516 ± 214 | 1,356 ± 70 | 271 ± 54 | 889 ± 194 | 14,565 ± 2,649 | 524 ± 100 | 257 ± 45 |
| P-value (Welch's t-test) | 0.958 | 0.949 | 0.643 | 0.502 | 0.029 | 0 | 0 | 0.484 | 0.823 | 0 | 0 |
