## Supplemental Table S1 for "Deep evolutionary origin of nematode SL2 *trans*-splicing revealed by genome-wide analysis of the *Trichinella spiralis* transcriptome"

**Supplementary Table 1. Detailed summaries of *Tsp*-SL read-screening, genome mapping and read quantification results for each screening stringency (10 bp or 8 bp) and set of gene annotations.** Results are broken down for each of the three replicate RNASeq libraries and each of the 15 TSL types. The quantification results detail the numbers of alignments that were uniquely assigned to a feature, represented multi-mapping reads or did not overlap a single feature.

*Tsp*-SL-containing read screening

| | LIB7135 | LIB7136 | LIB7137 | Total | % Total |
|---|---|---|---|---|---|
| read pairs | 68,557,151 | 63,199,696 | 69,408,020 | 201,164,867 | |
| mapped | 88.44% | 88.07% | 86.41% | 87.62% | |
| properly paired | 85.11% | 84.76% | 82.76% | 84.19% | |
| TSP-SL candidate pairs | 1,914,678 | 1,777,562 | 2,047,880 | 5,740,120 | |
| | 2.79% | 2.81% | 2.95% | 2.85% | |
| **10 bp stringency** | | | | | |
| TSP-SL1: 5'-AGGTATTTACCAGATCTAAAAG | 5,052 | 4,874 | 3,827 | 13,753 | 0.007% |
| TSP-SL2: 5'-AGGTATTTACCGAATTAAAAAG | 5,762 | 6,281 | 4,472 | 16,515 | 0.008% |
| TSP-SL3: 5'-GGTTATTTACCGAACTTAAAAG | 19,557 | 18,483 | 16,826 | 54,866 | 0.027% |
| TSP-SL4: 5'-GCGATTGTTCGAATTTACTTGAAG | 15,722 | 16,256 | 15,461 | 47,439 | 0.024% |
| TSP-SL5: 5'-AAATACCTTTCAATTTGTTTGAAG | 1,889 | 2,435 | 2,237 | 6,561 | 0.003% |
| TSP-SL6: 5'-AACCTTTGCGCATCGTTTAAAG | 13,655 | 17,055 | 11,730 | 42,440 | 0.021% |
| TSP-SL7: 5'-AACCTGCACGACTTGTTCGAAG | 18,306 | 18,453 | 19,491 | 56,250 | 0.028% |
| TSP-SL8: 5'-ATCTGTCGGTATTCCTGAAAG | 1,365 | 1,053 | 798 | 3,216 | 0.002% |
| TSP-SL9: 5'-AGACGTGGTTATTTATTGAAG | 397 | 487 | 406 | 1,290 | 0.001% |

| | | | | | |
|---|---|---|---|---|---|
| TSP-SL10: 5'-GGTAATATTTACTGAATTCAAG | 5,100 | 4,902 | 4,923 | 14,925 | 0.007% |
| TSP-SL11: 5'-AACCTTTGAACCCACTTCAAG | 20,130 | 20,282 | 18,426 | 58,838 | 0.029% |
| TSP-SL12: 5'-ACGAATTTACCGTATTTGTCAAG | 10,786 | 10,473 | 8,755 | 30,014 | 0.015% |
| TSP-SL13: 5'-TACCATTCAATTTATTTTGAAG | 640 | 682 | 706 | 2,028 | 0.001% |
| TSP-SL14: 5'-TACCGTTCAATTAATTTTGAAG | 1,014 | 958 | 1,001 | 2,973 | 0.001% |
| TSP-SL15: 5'-TACCGTTCAATTCATTTTGAAG | 486 | 504 | 626 | 1,616 | 0.001% |
| **Total TSP-SL** | **119,861** | **123,178** | **109,685** | **352,724** | 0.175% |
| **% total** | **0.17%** | **0.19%** | **0.16%** | **0.18%** | |
| **% candidates** | **6.26%** | **6.93%** | **5.36%** | **6.14%** | |
| **8 bp stringency** | | | | | |
| TSP-SL1: 5'-AGGTATTTACCAGATCTAAAAG | 6,059 | 6,007 | 4,564 | 16,630 | 0.008% |
| TSP-SL2: 5'-AGGTATTTACCGAATTAAAAAG | 11,469 | 12,426 | 8,619 | 32,514 | 0.016% |
| TSP-SL3: 5'-GGTTATTTACCGAACTTAAAAG | 32,022 | 30,907 | 28,983 | 91,912 | 0.046% |
| TSP-SL4: 5'-GCGATTGTTCGAATTTACTTGAAG | 16,480 | 17,099 | 16,151 | 49,730 | 0.025% |
| TSP-SL5: 5'-AAATACCTTTCAATTTGTTTGAAG | 3,293 | 3,875 | 4,073 | 11,241 | 0.006% |
| TSP-SL6: 5'-AACCTTTGCGCATCGTTTAAAG | 31,355 | 36,258 | 30,117 | 97,730 | 0.049% |
| TSP-SL7: 5'-AACCTGCACGACTTGTTCGAAG | 22,904 | 22,865 | 24,254 | 70,023 | 0.035% |

| | | | | | |
|---|---|---|---|---|---|
| TSP-SL8: 5'-ATCTGTCGGTATTCCTGAAAG | 4,221 | 3,202 | 2,668 | 10,091 | 0.005% |
| TSP-SL9: 5'-AGACGTGGTTATTTATTGAAG | 533 | 613 | 576 | 1,722 | 0.001% |
| TSP-SL10: 5'-GGTAATATTTACTGAATTCAAG | 6,335 | 6,121 | 6,133 | 18,589 | 0.009% |
| TSP-SL11: 5'-AACCTTTGAACCCACTTCAAG | 22,269 | 22,554 | 20,481 | 65,304 | 0.032% |
| TSP-SL12: 5'-ACGAATTTACCGTATTTGTCAAG | 12,881 | 12,128 | 10,295 | 35,304 | 0.018% |
| TSP-SL13: 5'-TACCATTCAATTTATTTTGAAG | 2,100 | 2,105 | 2,567 | 6,772 | 0.003% |
| TSP-SL14: 5'-TACCGTTCAATTAATTTTGAAG | 1,111 | 1,047 | 1,086 | 3,244 | 0.002% |
| TSP-SL15: 5'-TACCGTTCAATTCATTTTGAAG | 588 | 683 | 769 | 2,040 | 0.001% |
| **Total TSP-SL** | **173,620** | **177,890** | **161,336** | **512,846** | 0.255% |
| **% total** | **0.25%** | **0.28%** | **0.23%** | **0.25%** | |
| **% candidates** | **9.07%** | **10.01%** | **7.88%** | **8.93%** | |

TSP-SL13, 14 and 15 are not reliably distinguishable at 8 bp stringency.

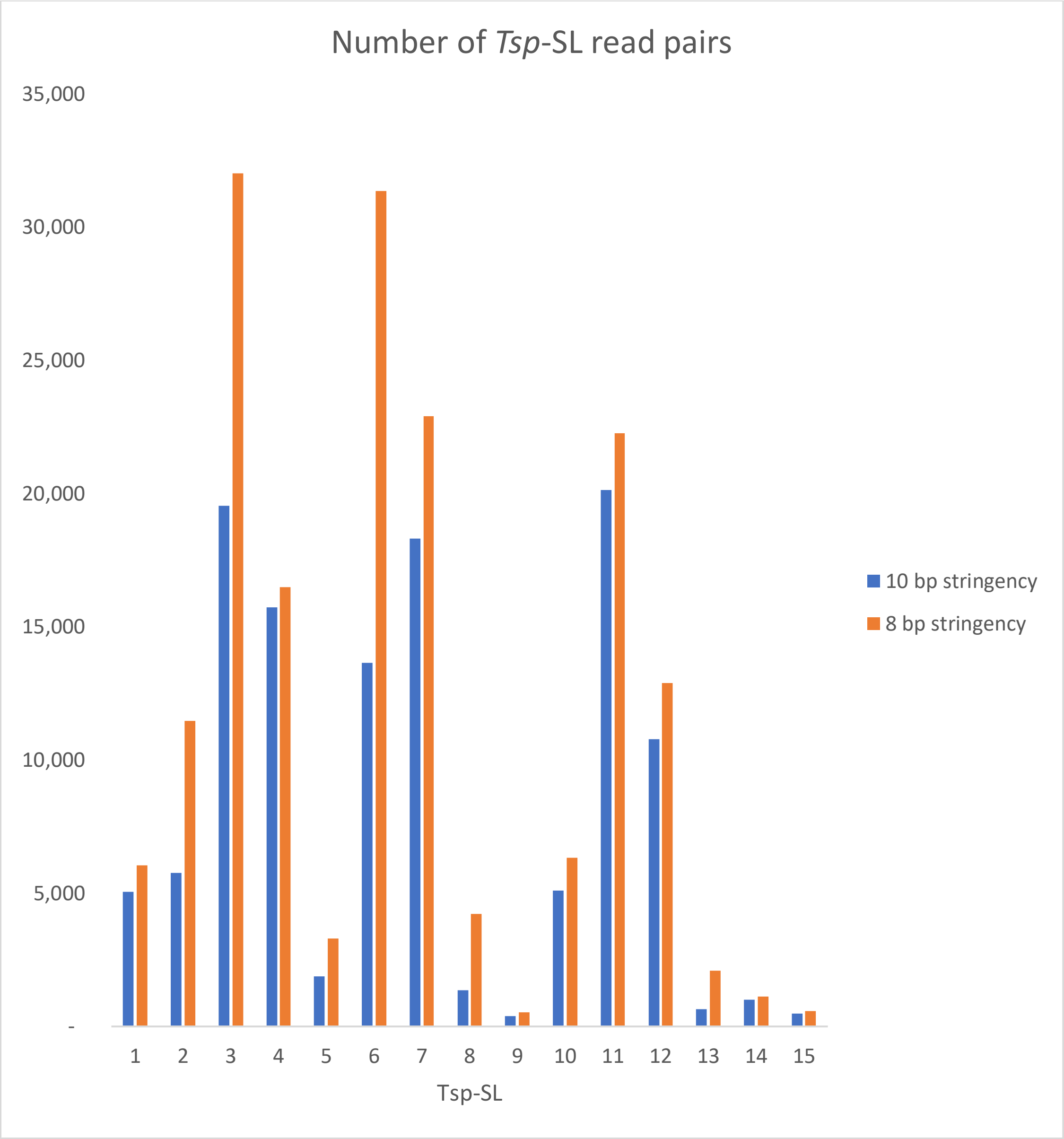

Number of *Tsp*-SL read pairs
35,000
30,000
25,000
20,000
15,000
10,000
5,000
-
Tsp-SL
10 bp stringency
8 bp stringency

**Read mapping**

| | LIB7135 | | | | | LIB7136 | | | | |
|---|---|---|---|---|---|---|---|---|---|---|
| *10 bp stringency* | | | | | | | | | | |
| | **Read pairs** | **Mapped** | | **Properly paired** | | **Read pairs** | **Mapped** | | **Properly paired** | |
| TSP-SL1 | 5,052 | 10,745 | 98.22% | 9,570 | 94.71% | 4,874 | 10,375 | 97.99% | 9,204 | 94.42% |
| TSP-SL2 | 5,762 | 12,584 | 97.90% | 10,874 | 94.36% | 6,281 | 13,602 | 97.69% | 11,782 | 93.79% |
| TSP-SL3 | 19,557 | 41,521 | 98.46% | 37,542 | 95.98% | 18,483 | 39,368 | 98.62% | 35,544 | 96.15% |
| TSP-SL4 | 15,722 | 33,209 | 98.50% | 30,238 | 96.16% | 16,256 | 34,654 | 98.51% | 31,180 | 95.90% |
| TSP-SL5 | 1,889 | 3,893 | 96.84% | 3,502 | 92.69% | 2,435 | 5,043 | 97.58% | 4,592 | 94.29% |
| TSP-SL6 | 13,655 | 28,749 | 98.44% | 26,212 | 95.98% | 17,055 | 36,341 | 98.55% | 32,772 | 96.08% |
| TSP-SL7 | 18,306 | 38,843 | 98.57% | 35,272 | 96.34% | 18,453 | 39,246 | 98.51% | 35,426 | 95.99% |
| TSP-SL8 | 1,365 | 2,905 | 98.01% | 2,588 | 94.80% | 1,053 | 2,221 | 97.97% | 1,978 | 93.92% |
| TSP-SL9 | 397 | 806 | 97.23% | 714 | 89.92% | 487 | 1,042 | 97.38% | 888 | 91.17% |
| TSP-SL10 | 5,100 | 11,597 | 98.00% | 9,646 | 94.57% | 4,902 | 11,093 | 97.98% | 9,260 | 94.45% |
| TSP-SL11 | 20,130 | 42,943 | 98.71% | 38,890 | 96.60% | 20,282 | 43,371 | 98.73% | 39,068 | 96.31% |
| TSP-SL12 | 10,786 | 24,761 | 98.52% | 20,638 | 95.67% | 10,473 | 23,871 | 98.35% | 19,996 | 95.46% |
| TSP-SL13 | 640 | 1,362 | 97.01% | 1,182 | 92.34% | 682 | 1,448 | 97.18% | 1,262 | 92.52% |

| | | | | | | | | | | |
|---|---|---|---|---|---|---|---|---|---|---|
| TSP-SL14 | 1,014 | 2,075 | 95.93% | 1,848 | 91.12% | 958 | 1,999 | 97.09% | 1,776 | 92.69% |
| TSP-SL15 | 486 | 1,009 | 96.28% | 892 | 91.77% | 504 | 1,051 | 96.16% | 916 | 90.87% |
| *8 bp stringency* | | | | | | | | | | |
| TSP-SL1 | 6,059 | 12,862 | 98.18% | 11,480 | 94.74% | 6,007 | 12,744 | 97.94% | 11,348 | 94.46% |
| TSP-SL2 | 11,469 | 24,803 | 98.02% | 21,754 | 94.84% | 12,426 | 26,835 | 98.06% | 23,568 | 94.83% |
| TSP-SL3 | 32,022 | 68,155 | 98.90% | 62,110 | 96.98% | 30,907 | 65,784 | 98.94% | 59,918 | 96.93% |
| TSP-SL4 | 16,480 | 34,796 | 98.51% | 31,712 | 96.21% | 17,099 | 36,410 | 98.47% | 32,766 | 95.81% |
| TSP-SL5 | 3,293 | 6,670 | 94.99% | 5,872 | 89.16% | 3,875 | 7,915 | 96.02% | 7,086 | 91.43% |
| TSP-SL6 | 31,355 | 66,682 | 98.84% | 60,740 | 96.86% | 36,258 | 77,373 | 98.90% | 70,250 | 96.88% |
| TSP-SL7 | 22,904 | 48,686 | 98.54% | 44,094 | 96.26% | 22,865 | 48,721 | 98.59% | 43,984 | 96.18% |
| TSP-SL8 | 4,221 | 8,892 | 97.50% | 7,928 | 93.91% | 3,202 | 6,692 | 97.62% | 6,014 | 93.91% |
| TSP-SL9 | 533 | 1,058 | 92.97% | 864 | 81.05% | 613 | 1,260 | 93.47% | 1,022 | 83.36% |
| TSP-SL10 | 6,335 | 15,994 | 97.69% | 11,832 | 93.39% | 6,121 | 15,463 | 97.68% | 11,394 | 93.07% |
| TSP-SL11 | 22,269 | 47,426 | 98.59% | 42,868 | 96.25% | 22,554 | 48,154 | 98.59% | 43,282 | 95.95% |
| TSP-SL12 | 12,881 | 29,461 | 98.21% | 24,466 | 94.97% | 12,128 | 27,662 | 98.16% | 23,070 | 95.11% |
| TSP-SL13 | 2,100 | 4,319 | 95.96% | 3,824 | 91.05% | 2,105 | 4,322 | 95.26% | 3,764 | 89.41% |

| | | | | | | | | | | |
|---|---|---|---|---|---|---|---|---|---|---|
| TSP-SL14 | 1,111 | 2,235 | 94.14% | 1,946 | 87.58% | 1,047 | 2,143 | 95.12% | 1,854 | 88.54% |
| TSP-SL15 | 588 | 1,170 | 92.56% | 990 | 84.18% | 683 | 1,409 | 93.93% | 1,176 | 86.09% |

TSP-SL13, 14 and 15 are not reliably distinguishable at 8 bp stringency.

| | | | | | | | | | | |
|---|---|---|---|---|---|---|---|---|---|---|
| Min | | | 92.56% | | 81.05% | | | 93.47% | | 83.36% |
| Max | | | 98.90% | | 96.98% | | | 98.94% | | 96.93% |

| 8bp stringency | | Total alignments | Exons Assigned | Exons % | Genes Assigned | Genes % | Unassigned_Unmapped | Unassigned_MultiMapping | Unassigned_NoFeatures | Exons Unassigned_Ambiguity | Genes Unassigned_Ambiguity |
|---|---|---|---|---|---|---|---|---|---|---|---|
| LIB7135 | TSP-SL10 | 8217 | 4232 | 51.50% | 4241 | 51.61% | 214 | 2749 | 1013 | 9 | 0 |
| LIB7135 | TSP-SL11 | 24051 | 16774 | 69.74% | 16781 | 69.77% | 326 | 3080 | 3863 | 8 | 1 |
| LIB7135 | TSP-SL12 | 14986 | 8522 | 56.87% | 8539 | 56.98% | 319 | 3455 | 2673 | 17 | 0 |
| LIB7135 | TSP-SL13 | 2241 | 1406 | 62.74% | 1406 | 62.74% | 142 | 241 | 452 | 0 | 0 |
| LIB7135 | TSP-SL14 | 1184 | 794 | 67.06% | 794 | 67.06% | 106 | 131 | 153 | 0 | 0 |
| LIB7135 | TSP-SL15 | 627 | 403 | 64.27% | 403 | 64.27% | 81 | 67 | 76 | 0 | 0 |
| LIB7135 | TSP-SL1 | 6551 | 4445 | 67.85% | 4446 | 67.87% | 98 | 852 | 1155 | 1 | 0 |
| LIB7135 | TSP-SL2 | 12646 | 8142 | 64.38% | 8152 | 64.46% | 252 | 2014 | 2228 | 10 | 0 |
| LIB7135 | TSP-SL3 | 34455 | 23968 | 69.56% | 23985 | 69.61% | 330 | 4256 | 5884 | 17 | 0 |
| LIB7135 | TSP-SL4 | 17664 | 12475 | 70.62% | 12480 | 70.65% | 271 | 2106 | 2807 | 5 | 0 |
| LIB7135 | TSP-SL5 | 3502 | 2269 | 64.79% | 2269 | 64.79% | 261 | 373 | 599 | 0 | 0 |
| LIB7135 | TSP-SL6 | 33723 | 23863 | 70.76% | 23875 | 70.80% | 429 | 4088 | 5331 | 12 | 0 |
| LIB7135 | TSP-SL7 | 24715 | 17256 | 69.82% | 17262 | 69.84% | 350 | 3151 | 3952 | 6 | 0 |
| LIB7135 | TSP-SL8 | 4559 | 3089 | 67.76% | 3091 | 67.80% | 115 | 583 | 770 | 2 | 0 |
| LIB7135 | TSP-SL9 | 565 | 325 | 57.52% | 325 | 57.52% | 74 | 53 | 113 | 0 | 0 |
| LIB7136 | TSP-SL10 | 7925 | 4005 | 50.54% | 4013 | 50.64% | 207 | 2638 | 1067 | 8 | 0 |
| LIB7136 | TSP-SL11 | 24423 | 16835 | 68.93% | 16841 | 68.96% | 304 | 3218 | 4060 | 6 | 0 |

| | | | | | | | | | | | |
|---|---|---|---|---|---|---|---|---|---|---|---|
| LIB7136 | TSP-SL12 | 14092 | 8006 | 56.81% | 8018 | 56.90% | 272 | 3238 | 2564 | 12 | 0 |
| LIB7136 | TSP-SL13 | 2259 | 1373 | 60.78% | 1373 | 60.78% | 172 | 252 | 462 | 0 | 0 |
| LIB7136 | TSP-SL14 | 1124 | 722 | 64.23% | 723 | 64.32% | 86 | 136 | 179 | 1 | 0 |
| LIB7136 | TSP-SL15 | 746 | 478 | 64.08% | 479 | 64.21% | 74 | 103 | 90 | 1 | 0 |
| LIB7136 | TSP-SL1 | 6510 | 4383 | 67.33% | 4385 | 67.36% | 125 | 872 | 1128 | 2 | 0 |
| LIB7136 | TSP-SL2 | 13687 | 8834 | 64.54% | 8846 | 64.63% | 253 | 2163 | 2425 | 12 | 0 |
| LIB7136 | TSP-SL3 | 33262 | 23169 | 69.66% | 23185 | 69.70% | 314 | 4105 | 5658 | 16 | 0 |
| LIB7136 | TSP-SL4 | 18488 | 12798 | 69.22% | 12801 | 69.24% | 308 | 2442 | 2937 | 3 | 0 |
| LIB7136 | TSP-SL5 | 4111 | 2776 | 67.53% | 2777 | 67.55% | 220 | 421 | 693 | 1 | 0 |
| LIB7136 | TSP-SL6 | 39119 | 27359 | 69.94% | 27384 | 70.00% | 420 | 4928 | 6386 | 26 | 1 |
| LIB7136 | TSP-SL7 | 24711 | 17178 | 69.52% | 17188 | 69.56% | 346 | 3205 | 3971 | 11 | 1 |
| LIB7136 | TSP-SL8 | 3427 | 2349 | 68.54% | 2349 | 68.54% | 84 | 396 | 598 | 0 | 0 |
| LIB7136 | TSP-SL9 | 670 | 359 | 53.58% | 359 | 53.58% | 69 | 98 | 144 | 0 | 0 |
| LIB7137 | TSP-SL10 | 7990 | 3896 | 48.76% | 3904 | 48.86% | 250 | 2719 | 1117 | 8 | 0 |
| LIB7137 | TSP-SL11 | 22174 | 15267 | 68.85% | 15271 | 68.87% | 322 | 2934 | 3647 | 4 | 0 |
| LIB7137 | TSP-SL12 | 11947 | 6631 | 55.50% | 6639 | 55.57% | 315 | 2759 | 2234 | 8 | 0 |
| LIB7137 | TSP-SL13 | 2741 | 1685 | 61.47% | 1685 | 61.47% | 219 | 299 | 538 | 0 | 0 |
| LIB7137 | TSP-SL14 | 1168 | 743 | 63.61% | 744 | 63.70% | 127 | 146 | 151 | 1 | 0 |
| LIB7137 | TSP-SL15 | 828 | 525 | 63.41% | 526 | 63.53% | 105 | 92 | 105 | 1 | 0 |

| | | | | | | | | | | | |
|---|---|---|---|---|---|---|---|---|---|---|---|
| LIB7137 | TSP-SL1 | 4899 | 3367 | 68.73% | 3369 | 68.77% | 105 | 598 | 827 | 2 | 0 |
| LIB7137 | TSP-SL2 | 9417 | 6023 | 63.96% | 6035 | 64.09% | 216 | 1386 | 1780 | 12 | 0 |
| LIB7137 | TSP-SL3 | 31266 | 21801 | 69.73% | 21817 | 69.78% | 300 | 3953 | 5196 | 16 | 0 |
| LIB7137 | TSP-SL4 | 17446 | 12059 | 69.12% | 12064 | 69.15% | 259 | 2315 | 2808 | 5 | 0 |
| LIB7137 | TSP-SL5 | 4391 | 2790 | 63.54% | 2794 | 63.63% | 327 | 562 | 708 | 4 | 0 |
| LIB7137 | TSP-SL6 | 32523 | 22771 | 70.02% | 22798 | 70.10% | 402 | 4173 | 5150 | 27 | 0 |
| LIB7137 | TSP-SL7 | 26352 | 18084 | 68.62% | 18096 | 68.67% | 372 | 3613 | 4268 | 15 | 3 |
| LIB7137 | TSP-SL8 | 2884 | 1897 | 65.78% | 1899 | 65.85% | 87 | 380 | 518 | 2 | 0 |
| LIB7137 | TSP-SL9 | 619 | 334 | 53.96% | 334 | 53.96% | 82 | 73 | 130 | 0 | 0 |
| **10bp stringency** | | | **Assigned** | **%** | **Assigned** | **%** | **Unassigned_Unmapped** | **Unassigned_MultiMapping** | **Unassigned_NoFeatures** | **Unassigned_Ambiguity** | **Unassigned_Ambiguity** |
| LIB7135 | TSP-SL10 | 5921 | 3532 | 59.65% | 3541 | 59.80% | 110 | 1363 | 907 | 9 | 0 |
| LIB7135 | TSP-SL11 | 21760 | 15215 | 69.92% | 15220 | 69.94% | 218 | 2813 | 3509 | 5 | 0 |
| LIB7135 | TSP-SL12 | 12572 | 7225 | 57.47% | 7239 | 57.58% | 183 | 2919 | 2231 | 14 | 0 |
| LIB7135 | TSP-SL13 | 703 | 375 | 53.34% | 375 | 53.34% | 28 | 107 | 193 | 0 | 0 |
| LIB7135 | TSP-SL14 | 1082 | 761 | 70.33% | 761 | 70.33% | 61 | 121 | 139 | 0 | 0 |
| LIB7135 | TSP-SL15 | 524 | 365 | 69.66% | 365 | 69.66% | 29 | 65 | 65 | 0 | 0 |
| LIB7135 | TSP-SL1 | 5473 | 3692 | 67.46% | 3693 | 67.48% | 81 | 729 | 970 | 1 | 0 |
| LIB7135 | TSP-SL2 | 6426 | 4068 | 63.31% | 4072 | 63.37% | 122 | 1117 | 1115 | 4 | 0 |

| | | | | | | | | | | | |
|---|---|---|---|---|---|---|---|---|---|---|---|
| LIB7135 | TSP-SL3 | 21092 | 14546 | 68.96% | 14554 | 69.00 % | 256 | 2715 | 3567 | 8 | 0 |
| LIB7135 | TSP-SL4 | 16863 | 11925 | 70.72% | 11927 | 70.73 % | 247 | 2029 | 2660 | 2 | 0 |
| LIB7135 | TSP-SL5 | 2008 | 1394 | 69.42% | 1394 | 69.42 % | 73 | 213 | 328 | 0 | 0 |
| LIB7135 | TSP-SL6 | 14610 | 10224 | 69.98% | 10227 | 70.00 % | 187 | 1672 | 2524 | 3 | 0 |
| LIB7135 | TSP-SL7 | 19713 | 13771 | 69.86% | 13775 | 69.88 % | 232 | 2453 | 3253 | 4 | 0 |
| LIB7135 | TSP-SL8 | 1483 | 994 | 67.03% | 994 | 67.03 % | 24 | 207 | 258 | 0 | 0 |
| LIB7135 | TSP-SL9 | 415 | 303 | 73.01% | 305 | 73.49 % | 14 | 32 | 64 | 2 | 0 |
| LIB7136 | TSP-SL10 | 5662 | 3331 | 58.83% | 3339 | 58.97 % | 102 | 1273 | 948 | 8 | 0 |
| LIB7136 | TSP-SL11 | 21975 | 15171 | 69.04% | 15176 | 69.06 % | 203 | 2912 | 3684 | 5 | 0 |
| LIB7136 | TSP-SL12 | 12148 | 6969 | 57.37% | 6978 | 57.44 % | 177 | 2764 | 2229 | 9 | 0 |
| LIB7136 | TSP-SL13 | 744 | 415 | 55.78% | 415 | 55.78 % | 30 | 105 | 194 | 0 | 0 |
| LIB7136 | TSP-SL14 | 1029 | 695 | 67.54% | 696 | 67.64 % | 40 | 125 | 168 | 1 | 0 |
| LIB7136 | TSP-SL15 | 545 | 359 | 65.87% | 360 | 66.06 % | 28 | 74 | 83 | 1 | 0 |
| LIB7136 | TSP-SL1 | 5299 | 3550 | 66.99% | 3552 | 67.03 % | 93 | 734 | 920 | 2 | 0 |
| LIB7136 | TSP-SL2 | 6963 | 4432 | 63.65% | 4433 | 63.67 % | 134 | 1173 | 1223 | 1 | 0 |
| LIB7136 | TSP-SL3 | 19969 | 13779 | 69.00% | 13786 | 69.04 % | 232 | 2596 | 3355 | 7 | 0 |
| LIB7136 | TSP-SL4 | 17593 | 12162 | 69.13% | 12165 | 69.15 % | 276 | 2350 | 2802 | 3 | 0 |
| LIB7136 | TSP-SL5 | 2581 | 1813 | 70.24% | 1813 | 70.24 % | 61 | 271 | 436 | 0 | 0 |
| LIB7136 | TSP-SL6 | 18448 | 12655 | 68.60% | 12655 | 68.60 % | 217 | 2413 | 3163 | 0 | 0 |

| | | | | | | | | | | | |
|---|---|---|---|---|---|---|---|---|---|---|---|
| LIB7136 | TSP-SL7 | 19928 | 13788 | 69.19% | 13794 | 69.22% | 267 | 2570 | 3296 | 7 | 1 |
| LIB7136 | TSP-SL8 | 1135 | 765 | 67.40% | 765 | 67.40% | 17 | 148 | 205 | 0 | 0 |
| LIB7136 | TSP-SL9 | 534 | 348 | 65.17% | 348 | 65.17% | 13 | 80 | 93 | 0 | 0 |
| LIB7137 | TSP-SL10 | 5720 | 3285 | 57.43% | 3292 | 57.55% | 89 | 1339 | 1000 | 7 | 0 |
| LIB7137 | TSP-SL11 | 19974 | 13786 | 69.02% | 13789 | 69.03% | 198 | 2672 | 3315 | 3 | 0 |
| LIB7137 | TSP-SL12 | 10158 | 5726 | 56.37% | 5731 | 56.42% | 164 | 2344 | 1919 | 5 | 0 |
| LIB7137 | TSP-SL13 | 770 | 394 | 51.17% | 394 | 51.17% | 52 | 107 | 217 | 0 | 0 |
| LIB7137 | TSP-SL14 | 1081 | 714 | 66.05% | 715 | 66.14% | 83 | 142 | 141 | 1 | 0 |
| LIB7137 | TSP-SL15 | 671 | 473 | 70.49% | 474 | 70.64% | 35 | 73 | 89 | 1 | 0 |
| LIB7137 | TSP-SL1 | 4107 | 2833 | 68.98% | 2835 | 69.03% | 76 | 504 | 692 | 2 | 0 |
| LIB7137 | TSP-SL2 | 4955 | 3078 | 62.12% | 3082 | 62.20% | 94 | 819 | 960 | 4 | 0 |
| LIB7137 | TSP-SL3 | 18218 | 12482 | 68.51% | 12486 | 68.54% | 218 | 2417 | 3097 | 4 | 0 |
| LIB7137 | TSP-SL4 | 16709 | 11552 | 69.14% | 11557 | 69.17% | 234 | 2230 | 2688 | 5 | 0 |
| LIB7137 | TSP-SL5 | 2420 | 1603 | 66.24% | 1604 | 66.28% | 75 | 324 | 417 | 1 | 0 |
| LIB7137 | TSP-SL6 | 12709 | 8611 | 67.76% | 8614 | 67.78% | 176 | 1696 | 2223 | 3 | 0 |
| LIB7137 | TSP-SL7 | 21136 | 14560 | 68.89% | 14566 | 68.92% | 246 | 2840 | 3481 | 9 | 3 |
| LIB7137 | TSP-SL8 | 864 | 567 | 65.63% | 567 | 65.63% | 17 | 117 | 163 | 0 | 0 |
| LIB7137 | TSP-SL9 | 434 | 291 | 67.05% | 291 | 67.05% | 11 | 50 | 82 | 0 | 0 |

| | | |
|---|---|---|
| Min | 48.76% | 48.86 % |
| Mean | 65.05% | 65.10 % |
| Max | 73.01% | 73.49 % |

**Quantification (brakerTRINITY)**

| 8bp stringency | | Total alignments | Exons Assigned | Exons % | Genes Assigned | Genes % | Unassigned_Unmapped | Unassigned_MultiMapping | Unassigned_NoFeatures | Exons Unassigned_Ambiguity | Genes Unassigned_Ambiguity |
|---|---|---|---|---|---|---|---|---|---|---|---|
| LIB7135 | TSL10 | 8217 | 4741 | 57.70% | 4747 | 57.77% | 214 | 2749 | 507 | 6 | 0 |
| LIB7135 | TSL11 | 24051 | 18511 | 76.97% | 18523 | 77.02% | 326 | 3080 | 2119 | 15 | 3 |
| LIB7135 | TSL12 | 14986 | 9940 | 66.33% | 9949 | 66.39% | 319 | 3455 | 1263 | 9 | 0 |
| LIB7135 | TSL13 | 2241 | 1564 | 69.79% | 1566 | 69.88% | 142 | 241 | 292 | 2 | 0 |
| LIB7135 | TSL14 | 1184 | 841 | 71.03% | 841 | 71.03% | 106 | 131 | 106 | 0 | 0 |
| LIB7135 | TSL15 | 627 | 415 | 66.19% | 415 | 66.19% | 81 | 67 | 64 | 0 | 0 |
| LIB7135 | TSL1 | 6551 | 4930 | 75.26% | 4932 | 75.29% | 98 | 852 | 668 | 3 | 1 |
| LIB7135 | TSL2 | 12646 | 9217 | 72.88% | 9243 | 73.09% | 252 | 2014 | 1136 | 27 | 1 |
| LIB7135 | TSL3 | 34455 | 26843 | 77.91% | 26881 | 78.02% | 330 | 4256 | 2985 | 41 | 3 |
| LIB7135 | TSL4 | 17664 | 13771 | 77.96% | 13779 | 78.01% | 271 | 2106 | 1505 | 11 | 3 |
| LIB7135 | TSL5 | 3502 | 2512 | 71.73% | 2513 | 71.76% | 261 | 373 | 352 | 4 | 3 |
| LIB7135 | TSL6 | 33723 | 26354 | 78.15% | 26396 | 78.27% | 429 | 4088 | 2807 | 45 | 3 |
| LIB7135 | TSL7 | 24715 | 19055 | 77.10% | 19066 | 77.14% | 350 | 3151 | 2147 | 12 | 1 |
| LIB7135 | TSL8 | 4559 | 3470 | 76.11% | 3476 | 76.24% | 115 | 583 | 385 | 6 | 0 |
| LIB7135 | TSL9 | 565 | 348 | 61.59% | 348 | 61.59% | 74 | 53 | 90 | 0 | 0 |

| | | | | | | | | | | | |
|---|---|---|---|---|---|---|---|---|---|---|---|
| LIB7136 | TSL10 | 7925 | 4600 | 58.04% | 4605 | 58.11% | 207 | 2638 | 475 | 5 | 0 |
| LIB7136 | TSL11 | 24423 | 18783 | 76.91% | 18807 | 77.01% | 304 | 3218 | 2093 | 25 | 1 |
| LIB7136 | TSL12 | 14092 | 9403 | 66.73% | 9412 | 66.79% | 272 | 3238 | 1170 | 9 | 0 |
| LIB7136 | TSL13 | 2259 | 1532 | 67.82% | 1533 | 67.86% | 172 | 252 | 302 | 1 | 0 |
| LIB7136 | TSL14 | 1124 | 775 | 68.95% | 776 | 69.04% | 86 | 136 | 126 | 1 | 0 |
| LIB7136 | TSL15 | 746 | 490 | 65.68% | 490 | 65.68% | 74 | 103 | 79 | 0 | 0 |
| LIB7136 | TSL1 | 6510 | 4915 | 75.50% | 4921 | 75.59% | 125 | 872 | 592 | 6 | 0 |
| LIB7136 | TSL2 | 13687 | 10018 | 73.19% | 10044 | 73.38% | 253 | 2163 | 1225 | 28 | 2 |
| LIB7136 | TSL3 | 33262 | 25803 | 77.58% | 25852 | 77.72% | 314 | 4105 | 2989 | 51 | 2 |
| LIB7136 | TSL4 | 18488 | 14212 | 76.87% | 14227 | 76.95% | 308 | 2442 | 1506 | 20 | 5 |
| LIB7136 | TSL5 | 4111 | 3083 | 74.99% | 3084 | 75.02% | 220 | 421 | 386 | 1 | 0 |
| LIB7136 | TSL6 | 39119 | 30345 | 77.57% | 30402 | 77.72% | 420 | 4928 | 3368 | 58 | 1 |
| LIB7136 | TSL7 | 24711 | 19064 | 77.15% | 19086 | 77.24% | 346 | 3205 | 2071 | 25 | 3 |
| LIB7136 | TSL8 | 3427 | 2606 | 76.04% | 2617 | 76.36% | 84 | 396 | 330 | 11 | 0 |
| LIB7136 | TSL9 | 670 | 403 | 60.15% | 403 | 60.15% | 69 | 98 | 98 | 2 | 2 |
| LIB7137 | TSL10 | 7990 | 4500 | 56.32% | 4501 | 56.33% | 250 | 2719 | 520 | 1 | 0 |
| LIB7137 | TSL11 | 22174 | 16941 | 76.40% | 16958 | 76.48% | 322 | 2934 | 1960 | 17 | 0 |
| LIB7137 | TSL12 | 11947 | 7754 | 64.90% | 7760 | 64.95% | 315 | 2759 | 1113 | 6 | 0 |
| LIB7137 | TSL13 | 2741 | 1867 | 68.11% | 1869 | 68.19% | 219 | 299 | 354 | 2 | 0 |

| | | | | | | | | | | | |
|---|---|---|---|---|---|---|---|---|---|---|---|
| LIB7137 | TSL14 | 1168 | 795 | 68.07% | 797 | 68.24% | 127 | 146 | 98 | 2 | 0 |
| LIB7137 | TSL15 | 828 | 540 | 65.22% | 541 | 65.34% | 105 | 92 | 90 | 1 | 0 |
| LIB7137 | TSL1 | 4899 | 3739 | 76.32% | 3741 | 76.36% | 105 | 598 | 454 | 3 | 1 |
| LIB7137 | TSL2 | 9417 | 6809 | 72.31% | 6824 | 72.46% | 216 | 1386 | 990 | 16 | 1 |
| LIB7137 | TSL3 | 31266 | 24255 | 77.58% | 24288 | 77.68% | 300 | 3953 | 2721 | 37 | 4 |
| LIB7137 | TSL4 | 17446 | 13420 | 76.92% | 13433 | 77.00% | 259 | 2315 | 1436 | 16 | 3 |
| LIB7137 | TSL5 | 4391 | 3102 | 70.64% | 3105 | 70.71% | 327 | 562 | 397 | 3 | 0 |
| LIB7137 | TSL6 | 32523 | 25171 | 77.39% | 25224 | 77.56% | 402 | 4173 | 2721 | 56 | 3 |
| LIB7137 | TSL7 | 26352 | 20162 | 76.51% | 20188 | 76.61% | 372 | 3613 | 2175 | 30 | 4 |
| LIB7137 | TSL8 | 2884 | 2133 | 73.96% | 2144 | 74.34% | 87 | 380 | 273 | 11 | 0 |
| LIB7137 | TSL9 | 619 | 363 | 58.64% | 364 | 58.80% | 82 | 73 | 100 | 1 | 0 |
| 10bp stringency | | | Assigned | | Assigned | | Unassigned_Unmapped | Unassigned_MultiMapping | Unassigned_NoFeatures | Unassigned_Ambiguity | Unassigned_Ambiguity |
| LIB7135 | TSL10 | 5921 | 4017 | 67.84% | 4020 | 67.89% | 110 | 1363 | 428 | 3 | 0 |
| LIB7135 | TSL11 | 21760 | 16869 | 77.52% | 16880 | 77.57% | 218 | 2813 | 1846 | 14 | 3 |
| LIB7135 | TSL12 | 12572 | 8402 | 66.83% | 8404 | 66.85% | 183 | 2919 | 1066 | 2 | 0 |
| LIB7135 | TSL13 | 703 | 393 | 55.90% | 393 | 55.90% | 28 | 107 | 175 | 0 | 0 |
| LIB7135 | TSL14 | 1082 | 807 | 74.58% | 807 | 74.58% | 61 | 121 | 93 | 0 | 0 |
| LIB7135 | TSL15 | 524 | 386 | 73.66% | 386 | 73.66% | 29 | 65 | 44 | 0 | 0 |

| | | | | | | | | | | | |
|---|---|---|---|---|---|---|---|---|---|---|---|
| **LIB7135** | **TSL1** | **5473** | **4096** | **74.84%** | **4098** | **74.88%** | **81** | **729** | **564** | **3** | **1** |
| **LIB7135** | **TSL2** | **6426** | **4587** | **71.38%** | **4596** | **71.52%** | **122** | **1117** | **590** | **10** | **1** |
| **LIB7135** | **TSL3** | **21092** | **16162** | **76.63%** | **16178** | **76.70%** | **256** | **2715** | **1940** | **19** | **3** |
| **LIB7135** | **TSL4** | **16863** | **13150** | **77.98%** | **13156** | **78.02%** | **247** | **2029** | **1428** | **9** | **3** |
| **LIB7135** | **TSL5** | **2008** | **1532** | **76.29%** | **1532** | **76.29%** | **73** | **213** | **187** | **3** | **3** |
| **LIB7135** | **TSL6** | **14610** | **11352** | **77.70%** | **11368** | **77.81%** | **187** | **1672** | **1381** | **18** | **2** |
| **LIB7135** | **TSL7** | **19713** | **15274** | **77.48%** | **15279** | **77.51%** | **232** | **2453** | **1749** | **5** | **0** |
| **LIB7135** | **TSL8** | **1483** | **1104** | **74.44%** | **1105** | **74.51%** | **24** | **207** | **147** | **1** | **0** |
| **LIB7135** | **TSL9** | **415** | **320** | **77.11%** | **321** | **77.35%** | **14** | **32** | **48** | **1** | **0** |
| **LIB7136** | **TSL10** | **5662** | **3892** | **68.74%** | **3896** | **68.81%** | **102** | **1273** | **391** | **4** | **0** |
| **LIB7136** | **TSL11** | **21975** | **17002** | **77.37%** | **17022** | **77.46%** | **203** | **2912** | **1838** | **20** | **0** |
| **LIB7136** | **TSL12** | **12148** | **8210** | **67.58%** | **8217** | **67.64%** | **177** | **2764** | **990** | **7** | **0** |
| **LIB7136** | **TSL13** | **744** | **435** | **58.47%** | **435** | **58.47%** | **30** | **105** | **174** | **0** | **0** |
| **LIB7136** | **TSL14** | **1029** | **749** | **72.79%** | **750** | **72.89%** | **40** | **125** | **114** | **1** | **0** |
| **LIB7136** | **TSL15** | **545** | **389** | **71.38%** | **389** | **71.38%** | **28** | **74** | **54** | **0** | **0** |
| **LIB7136** | **TSL1** | **5299** | **3973** | **74.98%** | **3977** | **75.05%** | **93** | **734** | **495** | **4** | **0** |
| **LIB7136** | **TSL2** | **6963** | **4994** | **71.72%** | **4996** | **71.75%** | **134** | **1173** | **658** | **4** | **2** |
| **LIB7136** | **TSL3** | **19969** | **15325** | **76.74%** | **15348** | **76.86%** | **232** | **2596** | **1792** | **24** | **1** |
| **LIB7136** | **TSL4** | **17593** | **13512** | **76.80%** | **13527** | **76.89%** | **276** | **2350** | **1436** | **19** | **4** |

| | | | | | | | | | | | |
|---|---|---|---|---|---|---|---|---|---|---|---|
| LIB7136 | TSL5 | 2581 | 2025 | 78.46 % | 2025 | 78.46 % | 61 | 271 | 224 | 0 | 0 |
| LIB7136 | TSL6 | 18448 | 14076 | 76.30 % | 14090 | 76.38 % | 217 | 2413 | 1728 | 14 | 0 |
| LIB7136 | TSL7 | 19928 | 15372 | 77.14 % | 15388 | 77.22 % | 267 | 2570 | 1701 | 18 | 2 |
| LIB7136 | TSL8 | 1135 | 866 | 76.30 % | 867 | 76.39 % | 17 | 148 | 103 | 1 | 0 |
| LIB7136 | TSL9 | 534 | 389 | 72.85 % | 389 | 72.85 % | 13 | 80 | 50 | 2 | 2 |
| LIB7137 | TSL10 | 5720 | 3860 | 67.48 % | 3861 | 67.50 % | 89 | 1339 | 431 | 1 | 0 |
| LIB7137 | TSL11 | 19974 | 15365 | 76.93 % | 15382 | 77.01 % | 198 | 2672 | 1722 | 17 | 0 |
| LIB7137 | TSL12 | 10158 | 6697 | 65.93 % | 6700 | 65.96 % | 164 | 2344 | 950 | 3 | 0 |
| LIB7137 | TSL13 | 770 | 414 | 53.77 % | 414 | 53.77 % | 52 | 107 | 197 | 0 | 0 |
| LIB7137 | TSL14 | 1081 | 768 | 71.05 % | 769 | 71.14 % | 83 | 142 | 87 | 1 | 0 |
| LIB7137 | TSL15 | 671 | 501 | 74.66 % | 502 | 74.81 % | 35 | 73 | 61 | 1 | 0 |
| LIB7137 | TSL1 | 4107 | 3137 | 76.38 % | 3139 | 76.43 % | 76 | 504 | 387 | 3 | 1 |
| LIB7137 | TSL2 | 4955 | 3473 | 70.09 % | 3476 | 70.15 % | 94 | 819 | 566 | 3 | 0 |
| LIB7137 | TSL3 | 18218 | 13883 | 76.20 % | 13896 | 76.28 % | 218 | 2417 | 1684 | 16 | 3 |
| LIB7137 | TSL4 | 16709 | 12842 | 76.86 % | 12855 | 76.93 % | 234 | 2230 | 1387 | 16 | 3 |
| LIB7137 | TSL5 | 2420 | 1794 | 74.13 % | 1795 | 74.17 % | 75 | 324 | 226 | 1 | 0 |
| LIB7137 | TSL6 | 12709 | 9602 | 75.55 % | 9611 | 75.62 % | 176 | 1696 | 1224 | 11 | 2 |
| LIB7137 | TSL7 | 21136 | 16235 | 76.81 % | 16251 | 76.89 % | 246 | 2840 | 1796 | 19 | 3 |
| LIB7137 | TSL8 | 864 | 642 | 74.31 % | 642 | 74.31 % | 17 | 117 | 88 | 0 | 0 |

| | | | | | | | | | | | |
|---|---|---|---|---|---|---|---|---|---|---|---|
| LIB7137 | TSL9 | 434 | 322 | 74.19 % | 323 | 74.42 % | 11 | 50 | 50 | 1 | 0 |
| | | Min | | 53.77 % | | 53.77 % | | | | | |
| | | Mean | | 72.13 % | | 72.20 % | | | | | |
| | | Max | | 78.46 % | | 78.46 % | | | | | |

**Quantification (brakerBAM)**

| 8bp stringency | | Total alignments | Exons Assigned | Exons % | Genes Assigned | Genes % | Unassigned_ Unmapped | Unassigned_ MultiMapping | Unassigned_ NoFeatures | Exons Unassigned_ Ambiguity | Genes Unassigned_ Ambiguity |
|---|---|---|---|---|---|---|---|---|---|---|---|
| LIB7135 | TSL10 | 8217 | 4717 | 57.41% | 4728 | 57.54% | 214 | 2749 | 526 | 11 | 0 |
| LIB7135 | TSL11 | 24051 | 18358 | 76.33% | 18380 | 76.42% | 326 | 3080 | 2263 | 24 | 2 |
| LIB7135 | TSL12 | 14986 | 9952 | 66.41% | 9962 | 66.48% | 319 | 3455 | 1250 | 10 | 0 |
| LIB7135 | TSL13 | 2241 | 1489 | 66.44% | 1491 | 66.53% | 142 | 241 | 367 | 2 | 0 |
| LIB7135 | TSL14 | 1184 | 821 | 69.34% | 822 | 69.43% | 106 | 131 | 125 | 1 | 0 |
| LIB7135 | TSL15 | 627 | 414 | 66.03% | 414 | 66.03% | 81 | 67 | 65 | 0 | 0 |
| LIB7135 | TSL1 | 6551 | 4908 | 74.92% | 4911 | 74.97% | 98 | 852 | 689 | 4 | 1 |
| LIB7135 | TSL2 | 12646 | 9228 | 72.97% | 9256 | 73.19% | 252 | 2014 | 1124 | 28 | 0 |
| LIB7135 | TSL3 | 34455 | 26777 | 77.72% | 26818 | 77.83% | 330 | 4256 | 3049 | 43 | 2 |
| LIB7135 | TSL4 | 17664 | 13630 | 77.16% | 13644 | 77.24% | 271 | 2106 | 1641 | 16 | 2 |
| LIB7135 | TSL5 | 3502 | 2493 | 71.19% | 2495 | 71.25% | 261 | 373 | 373 | 2 | 0 |
| LIB7135 | TSL6 | 33723 | 26203 | 77.70% | 26247 | 77.83% | 429 | 4088 | 2957 | 46 | 2 |
| LIB7135 | TSL7 | 24715 | 18926 | 76.58% | 18945 | 76.65% | 350 | 3151 | 2269 | 19 | 0 |
| LIB7135 | TSL8 | 4559 | 3478 | 76.29% | 3484 | 76.42% | 115 | 583 | 377 | 6 | 0 |
| LIB7135 | TSL9 | 565 | 349 | 61.77% | 350 | 61.95% | 74 | 53 | 88 | 1 | 0 |
| LIB7136 | TSL10 | 7925 | 4568 | 57.64% | 4573 | 57.70% | 207 | 2638 | 507 | 5 | 0 |
| LIB7136 | TSL11 | 24423 | 18654 | 76.38% | 18694 | 76.54% | 304 | 3218 | 2206 | 41 | 1 |
| LIB7136 | TSL12 | 14092 | 9404 | 66.73% | 9416 | 66.82% | 272 | 3238 | 1166 | 12 | 0 |
| LIB7136 | TSL13 | 2259 | 1477 | 65.38% | 1479 | 65.47% | 172 | 252 | 356 | 2 | 0 |
| LIB7136 | TSL14 | 1124 | 768 | 68.33% | 769 | 68.42% | 86 | 136 | 133 | 1 | 0 |
| LIB7136 | TSL15 | 746 | 479 | 64.21% | 479 | 64.21% | 74 | 103 | 90 | 0 | 0 |
| LIB7136 | TSL1 | 6510 | 4874 | 74.87% | 4885 | 75.04% | 125 | 872 | 628 | 11 | 0 |
| LIB7136 | TSL2 | 13687 | 10012 | 73.15% | 10036 | 73.33% | 253 | 2163 | 1233 | 26 | 2 |
| LIB7136 | TSL3 | 33262 | 25753 | 77.42% | 25802 | 77.57% | 314 | 4105 | 3039 | 51 | 2 |
| LIB7136 | TSL4 | 18488 | 14142 | 76.49% | 14159 | 76.58% | 308 | 2442 | 1578 | 18 | 1 |

| | | | | | | | | | | | |
|---|---|---|---|---|---|---|---|---|---|---|---|
| LIB7136 | TSL5 | 4111 | 3041 | 73.97% | 3044 | 74.05% | 220 | 421 | 426 | 3 | 0 |
| LIB7136 | TSL6 | 39119 | 30195 | 77.19% | 30269 | 77.38% | 420 | 4928 | 3502 | 74 | 0 |
| LIB7136 | TSL7 | 24711 | 18964 | 76.74% | 18991 | 76.85% | 346 | 3205 | 2167 | 29 | 2 |
| LIB7136 | TSL8 | 3427 | 2592 | 75.63% | 2601 | 75.90% | 84 | 396 | 346 | 9 | 0 |
| LIB7136 | TSL9 | 670 | 408 | 60.90% | 408 | 60.90% | 69 | 98 | 95 | 0 | 0 |
| LIB7137 | TSL10 | 7990 | 4483 | 56.11% | 4484 | 56.12% | 250 | 2719 | 537 | 1 | 0 |
| LIB7137 | TSL11 | 22174 | 16746 | 75.52% | 16773 | 75.64% | 322 | 2934 | 2145 | 27 | 0 |
| LIB7137 | TSL12 | 11947 | 7755 | 64.91% | 7763 | 64.98% | 315 | 2759 | 1110 | 8 | 0 |
| LIB7137 | TSL13 | 2741 | 1823 | 66.51% | 1825 | 66.58% | 219 | 299 | 398 | 2 | 0 |
| LIB7137 | TSL14 | 1168 | 773 | 66.18% | 775 | 66.35% | 127 | 146 | 120 | 2 | 0 |
| LIB7137 | TSL15 | 828 | 528 | 63.77% | 529 | 63.89% | 105 | 92 | 102 | 1 | 0 |
| LIB7137 | TSL1 | 4899 | 3706 | 75.65% | 3708 | 75.69% | 105 | 598 | 488 | 2 | 0 |
| LIB7137 | TSL2 | 9417 | 6810 | 72.32% | 6829 | 72.52% | 216 | 1386 | 985 | 20 | 1 |
| LIB7137 | TSL3 | 31266 | 24106 | 77.10% | 24143 | 77.22% | 300 | 3953 | 2869 | 38 | 1 |
| LIB7137 | TSL4 | 17446 | 13310 | 76.29% | 13328 | 76.40% | 259 | 2315 | 1544 | 18 | 0 |
| LIB7137 | TSL5 | 4391 | 3051 | 69.48% | 3054 | 69.55% | 327 | 562 | 448 | 3 | 0 |
| LIB7137 | TSL6 | 32523 | 24964 | 76.76% | 25021 | 76.93% | 402 | 4173 | 2926 | 58 | 1 |
| LIB7137 | TSL7 | 26352 | 19944 | 75.68% | 19968 | 75.77% | 372 | 3613 | 2396 | 27 | 3 |
| LIB7137 | TSL8 | 2884 | 2136 | 74.06% | 2147 | 74.45% | 87 | 380 | 270 | 11 | 0 |
| LIB7137 | TSL9 | 619 | 373 | 60.26% | 374 | 60.42% | 82 | 73 | 90 | 1 | 0 |
| 10bp stringency | | | Assigned | | Assigned | | Unassigned_ Unmapped | Unassigned_ MultiMapping | Unassigned_ NoFeatures | Unassigned_ Ambiguity | Unassigned_ Ambiguity |
| LIB7135 | TSL10 | 5921 | 3994 | 67.45% | 4002 | 67.59% | 110 | 1363 | 446 | 8 | 0 |
| LIB7135 | TSL11 | 21760 | 16722 | 76.85% | 16742 | 76.94% | 218 | 2813 | 1985 | 22 | 2 |
| LIB7135 | TSL12 | 12572 | 8409 | 66.89% | 8411 | 66.90% | 183 | 2919 | 1059 | 2 | 0 |
| LIB7135 | TSL13 | 703 | 379 | 53.91% | 379 | 53.91% | 28 | 107 | 189 | 0 | 0 |
| LIB7135 | TSL14 | 1082 | 787 | 72.74% | 788 | 72.83% | 61 | 121 | 112 | 1 | 0 |
| LIB7135 | TSL15 | 524 | 383 | 73.09% | 383 | 73.09% | 29 | 65 | 47 | 0 | 0 |
| LIB7135 | TSL1 | 5473 | 4071 | 74.38% | 4074 | 74.44% | 81 | 729 | 588 | 4 | 1 |

| | | | | | | | | | | | |
|---|---|---|---|---|---|---|---|---|---|---|---|
| LIB7135 | TSL2 | 6426 | 4597 | 71.54% | 4606 | 71.68% | 122 | 1117 | 581 | 9 | 0 |
| LIB7135 | TSL3 | 21092 | 16071 | 76.19% | 16088 | 76.28% | 256 | 2715 | 2031 | 19 | 2 |
| LIB7135 | TSL4 | 16863 | 13003 | 77.11% | 13016 | 77.19% | 247 | 2029 | 1569 | 15 | 2 |
| LIB7135 | TSL5 | 2008 | 1527 | 76.05% | 1528 | 76.10% | 73 | 213 | 194 | 1 | 0 |
| LIB7135 | TSL6 | 14610 | 11318 | 77.47% | 11334 | 77.58% | 187 | 1672 | 1416 | 17 | 1 |
| LIB7135 | TSL7 | 19713 | 15146 | 76.83% | 15158 | 76.89% | 232 | 2453 | 1870 | 12 | 0 |
| LIB7135 | TSL8 | 1483 | 1110 | 74.85% | 1111 | 74.92% | 24 | 207 | 141 | 1 | 0 |
| LIB7135 | TSL9 | 415 | 324 | 78.07% | 324 | 78.07% | 14 | 32 | 45 | 0 | 0 |
| LIB7136 | TSL10 | 5662 | 3854 | 68.07% | 3858 | 68.14% | 102 | 1273 | 429 | 4 | 0 |
| LIB7136 | TSL11 | 21975 | 16864 | 76.74% | 16897 | 76.89% | 203 | 2912 | 1963 | 33 | 0 |
| LIB7136 | TSL12 | 12148 | 8209 | 67.57% | 8217 | 67.64% | 177 | 2764 | 990 | 8 | 0 |
| LIB7136 | TSL13 | 744 | 429 | 57.66% | 429 | 57.66% | 30 | 105 | 180 | 0 | 0 |
| LIB7136 | TSL14 | 1029 | 742 | 72.11% | 743 | 72.21% | 40 | 125 | 121 | 1 | 0 |
| LIB7136 | TSL15 | 545 | 378 | 69.36% | 378 | 69.36% | 28 | 74 | 65 | 0 | 0 |
| LIB7136 | TSL1 | 5299 | 3928 | 74.13% | 3934 | 74.24% | 93 | 734 | 538 | 6 | 0 |
| LIB7136 | TSL2 | 6963 | 4988 | 71.64% | 4990 | 71.66% | 134 | 1173 | 664 | 4 | 2 |
| LIB7136 | TSL3 | 19969 | 15173 | 75.98% | 15200 | 76.12% | 232 | 2596 | 1941 | 27 | 0 |
| LIB7136 | TSL4 | 17593 | 13448 | 76.44% | 13465 | 76.54% | 276 | 2350 | 1502 | 17 | 0 |
| LIB7136 | TSL5 | 2581 | 1982 | 76.79% | 1983 | 76.83% | 61 | 271 | 266 | 1 | 0 |
| LIB7136 | TSL6 | 18448 | 13990 | 75.83% | 14015 | 75.97% | 217 | 2413 | 1803 | 25 | 0 |
| LIB7136 | TSL7 | 19928 | 15292 | 76.74% | 15311 | 76.83% | 267 | 2570 | 1779 | 20 | 1 |
| LIB7136 | TSL8 | 1135 | 844 | 74.36% | 845 | 74.45% | 17 | 148 | 125 | 1 | 0 |
| LIB7136 | TSL9 | 534 | 394 | 73.78% | 394 | 73.78% | 13 | 80 | 47 | 0 | 0 |
| LIB7137 | TSL10 | 5720 | 3843 | 67.19% | 3844 | 67.20% | 89 | 1339 | 448 | 1 | 0 |
| LIB7137 | TSL11 | 19974 | 15181 | 76.00% | 15208 | 76.14% | 198 | 2672 | 1896 | 27 | 0 |
| LIB7137 | TSL12 | 10158 | 6700 | 65.96% | 6703 | 65.99% | 164 | 2344 | 947 | 3 | 0 |
| LIB7137 | TSL13 | 770 | 414 | 53.77% | 414 | 53.77% | 52 | 107 | 197 | 0 | 0 |
| LIB7137 | TSL14 | 1081 | 747 | 69.10% | 748 | 69.20% | 83 | 142 | 108 | 1 | 0 |
| LIB7137 | TSL15 | 671 | 489 | 72.88% | 490 | 73.03% | 35 | 73 | 73 | 1 | 0 |
| LIB7137 | TSL1 | 4107 | 3099 | 75.46% | 3101 | 75.51% | 76 | 504 | 426 | 2 | 0 |

| | | | | | | | | | | | |
|---|---|---|---|---|---|---|---|---|---|---|---|
| LIB7137 | TSL2 | 4955 | 3473 | 70.09% | 3477 | 70.17% | 94 | 819 | 565 | 4 | 0 |
| LIB7137 | TSL3 | 18218 | 13698 | 75.19% | 13710 | 75.26% | 218 | 2417 | 1872 | 13 | 1 |
| LIB7137 | TSL4 | 16709 | 12732 | 76.20% | 12750 | 76.31% | 234 | 2230 | 1495 | 18 | 0 |
| LIB7137 | TSL5 | 2420 | 1761 | 72.77% | 1762 | 72.81% | 75 | 324 | 259 | 1 | 0 |
| LIB7137 | TSL6 | 12709 | 9522 | 74.92% | 9531 | 74.99% | 176 | 1696 | 1306 | 9 | 0 |
| LIB7137 | TSL7 | 21136 | 16056 | 75.97% | 16072 | 76.04% | 246 | 2840 | 1975 | 19 | 3 |
| LIB7137 | TSL8 | 864 | 639 | 73.96% | 639 | 73.96% | 17 | 117 | 91 | 0 | 0 |
| LIB7137 | TSL9 | 434 | 333 | 76.73% | 334 | 76.96% | 11 | 50 | 39 | 1 | 0 |
| | | Min | | 53.77% | | 53.77% | | | | | |
| | | Mean | | 71.56% | | 71.66% | | | | | |
| | | Max | | 78.07% | | 78.07% | | | | | |

**Quantification (stringtie)**

| | | | Exons | | Genes | | | | | Exons | Genes |
|---|---|---|---|---|---|---|---|---|---|---|---|
| 8bp stringency | | Total alignments | Assigned | % | Assigned | % | Unassigned_ Unmapped | Unassigned _MultiMap ping | Unassigned_ NoFeatures | Unassigned _Ambiguity | Unassigned _Ambiguity |
| LIB7135 | TSL10 | 8217 | 4902 | 59.66% | 4916 | 59.83% | 214 | 2749 | 338 | 14 | 0 |
| LIB7135 | TSL11 | 24051 | 19581 | 81.41% | 19589 | 81.45% | 326 | 3080 | 1056 | 8 | 0 |
| LIB7135 | TSL12 | 14986 | 10357 | 69.11% | 10373 | 69.22% | 319 | 3455 | 839 | 16 | 0 |
| LIB7135 | TSL13 | 2241 | 1722 | 76.84% | 1724 | 76.93% | 142 | 241 | 134 | 2 | 0 |
| LIB7135 | TSL14 | 1184 | 898 | 75.84% | 898 | 75.84% | 106 | 131 | 49 | 0 | 0 |
| LIB7135 | TSL15 | 627 | 452 | 72.09% | 452 | 72.09% | 81 | 67 | 27 | 0 | 0 |
| LIB7135 | TSL1 | 6551 | 5318 | 81.18% | 5335 | 81.44% | 98 | 852 | 266 | 17 | 0 |
| LIB7135 | TSL2 | 12646 | 9830 | 77.73% | 9835 | 77.77% | 252 | 2014 | 545 | 5 | 0 |
| LIB7135 | TSL3 | 34455 | 28367 | 82.33% | 28407 | 82.45% | 330 | 4256 | 1462 | 40 | 0 |
| LIB7135 | TSL4 | 17664 | 14581 | 82.55% | 14614 | 82.73% | 271 | 2106 | 673 | 33 | 0 |
| LIB7135 | TSL5 | 3502 | 2684 | 76.64% | 2689 | 76.78% | 261 | 373 | 179 | 5 | 0 |
| LIB7135 | TSL6 | 33723 | 27782 | 82.38% | 27808 | 82.46% | 429 | 4088 | 1398 | 26 | 0 |
| LIB7135 | TSL7 | 24715 | 20246 | 81.92% | 20254 | 81.95% | 350 | 3151 | 960 | 8 | 0 |
| LIB7135 | TSL8 | 4559 | 3676 | 80.63% | 3680 | 80.72% | 115 | 583 | 181 | 4 | 0 |
| LIB7135 | TSL9 | 565 | 401 | 70.97% | 401 | 70.97% | 74 | 53 | 37 | 0 | 0 |
| LIB7136 | TSL10 | 7925 | 4741 | 59.82% | 4751 | 59.95% | 207 | 2638 | 329 | 10 | 0 |
| LIB7136 | TSL11 | 24423 | 19837 | 81.22% | 19858 | 81.31% | 304 | 3218 | 1043 | 21 | 0 |
| LIB7136 | TSL12 | 14092 | 9826 | 69.73% | 9844 | 69.86% | 272 | 3238 | 738 | 18 | 0 |
| LIB7136 | TSL13 | 2259 | 1677 | 74.24% | 1677 | 74.24% | 172 | 252 | 158 | 0 | 0 |
| LIB7136 | TSL14 | 1124 | 848 | 75.44% | 848 | 75.44% | 86 | 136 | 54 | 0 | 0 |
| LIB7136 | TSL15 | 746 | 535 | 71.72% | 535 | 71.72% | 74 | 103 | 34 | 0 | 0 |
| LIB7136 | TSL1 | 6510 | 5225 | 80.26% | 5236 | 80.43% | 125 | 872 | 277 | 11 | 0 |
| LIB7136 | TSL2 | 13687 | 10650 | 77.81% | 10660 | 77.88% | 253 | 2163 | 611 | 10 | 0 |
| LIB7136 | TSL3 | 33262 | 27493 | 82.66% | 27536 | 82.79% | 314 | 4105 | 1307 | 43 | 0 |
| LIB7136 | TSL4 | 18488 | 14930 | 80.76% | 14971 | 80.98% | 308 | 2442 | 767 | 41 | 0 |

| | | | | | | | | | | | |
|---|---|---|---|---|---|---|---|---|---|---|---|
| **LIB7136** | **TSL5** | **4111** | **3267** | **79.47%** | **3277** | **79.71%** | **220** | **421** | **193** | **10** | **0** |
| **LIB7136** | **TSL6** | **39119** | **32126** | **82.12%** | **32150** | **82.19%** | **420** | **4928** | **1621** | **24** | **0** |
| **LIB7136** | **TSL7** | **24711** | **20166** | **81.61%** | **20177** | **81.65%** | **346** | **3205** | **983** | **11** | **0** |
| **LIB7136** | **TSL8** | **3427** | **2785** | **81.27%** | **2786** | **81.30%** | **84** | **396** | **161** | **1** | **0** |
| **LIB7136** | **TSL9** | **670** | **463** | **69.10%** | **463** | **69.10%** | **69** | **98** | **40** | **0** | **0** |
| **LIB7137** | **TSL10** | **7990** | **4660** | **58.32%** | **4667** | **58.41%** | **250** | **2719** | **354** | **7** | **0** |
| **LIB7137** | **TSL11** | **22174** | **17903** | **80.74%** | **17914** | **80.79%** | **322** | **2934** | **1004** | **11** | **0** |
| **LIB7137** | **TSL12** | **11947** | **8157** | **68.28%** | **8175** | **68.43%** | **315** | **2759** | **698** | **18** | **0** |
| **LIB7137** | **TSL13** | **2741** | **2060** | **75.16%** | **2061** | **75.19%** | **219** | **299** | **162** | **1** | **0** |
| **LIB7137** | **TSL14** | **1168** | **840** | **71.92%** | **841** | **72.00%** | **127** | **146** | **54** | **1** | **0** |
| **LIB7137** | **TSL15** | **828** | **587** | **70.89%** | **587** | **70.89%** | **105** | **92** | **44** | **0** | **0** |
| **LIB7137** | **TSL1** | **4899** | **3989** | **81.42%** | **3999** | **81.63%** | **105** | **598** | **197** | **10** | **0** |
| **LIB7137** | **TSL2** | **9417** | **7356** | **78.11%** | **7366** | **78.22%** | **216** | **1386** | **449** | **10** | **0** |
| **LIB7137** | **TSL3** | **31266** | **25726** | **82.28%** | **25765** | **82.41%** | **300** | **3953** | **1248** | **39** | **0** |
| **LIB7137** | **TSL4** | **17446** | **14111** | **80.88%** | **14144** | **81.07%** | **259** | **2315** | **728** | **33** | **0** |
| **LIB7137** | **TSL5** | **4391** | **3308** | **75.34%** | **3311** | **75.40%** | **327** | **562** | **191** | **3** | **0** |
| **LIB7137** | **TSL6** | **32523** | **26634** | **81.89%** | **26643** | **81.92%** | **402** | **4173** | **1305** | **9** | **0** |
| **LIB7137** | **TSL7** | **26352** | **21301** | **80.83%** | **21312** | **80.87%** | **372** | **3613** | **1055** | **11** | **0** |
| **LIB7137** | **TSL8** | **2884** | **2287** | **79.30%** | **2287** | **79.30%** | **87** | **380** | **130** | **0** | **0** |
| **LIB7137** | **TSL9** | **619** | **428** | **69.14%** | **428** | **69.14%** | **82** | **73** | **36** | **0** | **0** |
| **10bp stringency** | | | **Assigned** | | **Assigned** | | **Unassigned_ Unmapped** | **Unassigned _MultiMap ping** | **Unassigned_ NoFeatures** | **Unassigned _Ambiguity** | **Unassigned _Ambiguity** |
| **LIB7135** | **TSL10** | **5921** | **4160** | **70.26%** | **4173** | **70.48%** | **110** | **1363** | **275** | **13** | **0** |
| **LIB7135** | **TSL11** | **21760** | **17807** | **81.83%** | **17814** | **81.87%** | **218** | **2813** | **915** | **7** | **0** |
| **LIB7135** | **TSL12** | **12572** | **8783** | **69.86%** | **8791** | **69.93%** | **183** | **2919** | **679** | **8** | **0** |
| **LIB7135** | **TSL13** | **703** | **501** | **71.27%** | **502** | **71.41%** | **28** | **107** | **66** | **1** | **0** |
| **LIB7135** | **TSL14** | **1082** | **858** | **79.30%** | **858** | **79.30%** | **61** | **121** | **42** | **0** | **0** |
| **LIB7135** | **TSL15** | **524** | **412** | **78.63%** | **412** | **78.63%** | **29** | **65** | **18** | **0** | **0** |

| | | | | | | | | | | | |
|---|---|---|---|---|---|---|---|---|---|---|---|
| LIB7135 | TSL1 | 5473 | 4421 | 80.78% | 4438 | 81.09% | 81 | 729 | 225 | 17 | 0 |
| LIB7135 | TSL2 | 6426 | 4911 | 76.42% | 4914 | 76.47% | 122 | 1117 | 273 | 3 | 0 |
| LIB7135 | TSL3 | 21092 | 17137 | 81.25% | 17177 | 81.44% | 256 | 2715 | 944 | 40 | 0 |
| LIB7135 | TSL4 | 16863 | 13901 | 82.43% | 13933 | 82.62% | 247 | 2029 | 654 | 32 | 0 |
| LIB7135 | TSL5 | 2008 | 1624 | 80.88% | 1629 | 81.13% | 73 | 213 | 93 | 5 | 0 |
| LIB7135 | TSL6 | 14610 | 12151 | 83.17% | 12176 | 83.34% | 187 | 1672 | 575 | 25 | 0 |
| LIB7135 | TSL7 | 19713 | 16275 | 82.56% | 16283 | 82.60% | 232 | 2453 | 745 | 8 | 0 |
| LIB7135 | TSL8 | 1483 | 1188 | 80.11% | 1192 | 80.38% | 24 | 207 | 60 | 4 | 0 |
| LIB7135 | TSL9 | 415 | 351 | 84.58% | 352 | 84.82% | 14 | 32 | 17 | 1 | 0 |
| LIB7136 | TSL10 | 5662 | 4000 | 70.65% | 4010 | 70.82% | 102 | 1273 | 277 | 10 | 0 |
| LIB7136 | TSL11 | 21975 | 17931 | 81.60% | 17952 | 81.69% | 203 | 2912 | 908 | 21 | 0 |
| LIB7136 | TSL12 | 12148 | 8557 | 70.44% | 8570 | 70.55% | 177 | 2764 | 637 | 13 | 0 |
| LIB7136 | TSL13 | 744 | 524 | 70.43% | 524 | 70.43% | 30 | 105 | 85 | 0 | 0 |
| LIB7136 | TSL14 | 1029 | 815 | 79.20% | 815 | 79.20% | 40 | 125 | 49 | 0 | 0 |
| LIB7136 | TSL15 | 545 | 417 | 76.51% | 417 | 76.51% | 28 | 74 | 26 | 0 | 0 |
| LIB7136 | TSL1 | 5299 | 4234 | 79.90% | 4245 | 80.11% | 93 | 734 | 227 | 11 | 0 |
| LIB7136 | TSL2 | 6963 | 5315 | 76.33% | 5320 | 76.40% | 134 | 1173 | 336 | 5 | 0 |
| LIB7136 | TSL3 | 19969 | 16296 | 81.61% | 16336 | 81.81% | 232 | 2596 | 805 | 40 | 0 |
| LIB7136 | TSL4 | 17593 | 14203 | 80.73% | 14244 | 80.96% | 276 | 2350 | 723 | 41 | 0 |
| LIB7136 | TSL5 | 2581 | 2128 | 82.45% | 2138 | 82.84% | 61 | 271 | 111 | 10 | 0 |
| LIB7136 | TSL6 | 18448 | 15087 | 81.78% | 15110 | 81.91% | 217 | 2413 | 708 | 23 | 0 |
| LIB7136 | TSL7 | 19928 | 16305 | 81.82% | 16316 | 81.87% | 267 | 2570 | 775 | 11 | 0 |
| LIB7136 | TSL8 | 1135 | 907 | 79.91% | 907 | 79.91% | 17 | 148 | 63 | 0 | 0 |
| LIB7136 | TSL9 | 534 | 412 | 77.15% | 412 | 77.15% | 13 | 80 | 29 | 0 | 0 |
| LIB7137 | TSL10 | 5720 | 3984 | 69.65% | 3991 | 69.77% | 89 | 1339 | 301 | 7 | 0 |
| LIB7137 | TSL11 | 19974 | 16227 | 81.24% | 16238 | 81.30% | 198 | 2672 | 866 | 11 | 0 |
| LIB7137 | TSL12 | 10158 | 7055 | 69.45% | 7061 | 69.51% | 164 | 2344 | 589 | 6 | 0 |
| LIB7137 | TSL13 | 770 | 518 | 67.27% | 519 | 67.40% | 52 | 107 | 92 | 1 | 0 |
| LIB7137 | TSL14 | 1081 | 814 | 75.30% | 814 | 75.30% | 83 | 142 | 42 | 0 | 0 |
| LIB7137 | TSL15 | 671 | 534 | 79.58% | 534 | 79.58% | 35 | 73 | 29 | 0 | 0 |

| | | | | | | | | | | | |
|---|---|---|---|---|---|---|---|---|---|---|---|
| LIB7137 | TSL1 | 4107 | 3349 | 81.54% | 3359 | 81.79% | 76 | 504 | 168 | 10 | 0 |
| LIB7137 | TSL2 | 4955 | 3797 | 76.63% | 3802 | 76.73% | 94 | 819 | 240 | 5 | 0 |
| LIB7137 | TSL3 | 18218 | 14775 | 81.10% | 14813 | 81.31% | 218 | 2417 | 770 | 38 | 0 |
| LIB7137 | TSL4 | 16709 | 13511 | 80.86% | 13544 | 81.06% | 234 | 2230 | 701 | 33 | 0 |
| LIB7137 | TSL5 | 2420 | 1922 | 79.42% | 1925 | 79.55% | 75 | 324 | 96 | 3 | 0 |
| LIB7137 | TSL6 | 12709 | 10363 | 81.54% | 10371 | 81.60% | 176 | 1696 | 466 | 8 | 0 |
| LIB7137 | TSL7 | 21136 | 17187 | 81.32% | 17198 | 81.37% | 246 | 2840 | 852 | 11 | 0 |
| LIB7137 | TSL8 | 864 | 699 | 80.90% | 699 | 80.90% | 17 | 117 | 31 | 0 | 0 |
| LIB7137 | TSL9 | 434 | 354 | 81.57% | 354 | 81.57% | 11 | 50 | 19 | 0 | 0 |
| | | Min | | 58.32% | | 58.41% | | | | | |
| | | Mean | | 77.27% | | 77.37% | | | | | |
| | | Max | | 84.58% | | 84.82% | | | | | |

**Quantification (denovo_all)**

| 8bp stringency | | Total alignments | Exons Assigned | Exons % | Genes Assigned | Genes % | Unassigned_ Unmapped | Unassigned _MultiMap ping | Unassigned_ NoFeatures | Exons Unassigned_ Ambiguity | Genes Unassigned_ Ambiguity |
|---|---|---|---|---|---|---|---|---|---|---|---|
| LIB7135 | TSL10 | 8217 | 5145 | 62.61% | 5159 | 62.78% | 214 | 2749 | 95 | 14 | 0 |
| LIB7135 | TSL11 | 24051 | 20343 | 84.58% | 20352 | 84.62% | 326 | 3080 | 293 | 9 | 0 |
| LIB7135 | TSL12 | 14986 | 11000 | 73.40% | 11017 | 73.52% | 319 | 3455 | 195 | 17 | 0 |
| LIB7135 | TSL13 | 2241 | 1781 | 79.47% | 1784 | 79.61% | 142 | 241 | 74 | 3 | 0 |
| LIB7135 | TSL14 | 1184 | 929 | 78.46% | 929 | 78.46% | 106 | 131 | 18 | 0 | 0 |
| LIB7135 | TSL15 | 627 | 461 | 73.52% | 461 | 73.52% | 81 | 67 | 18 | 0 | 0 |
| LIB7135 | TSL1 | 6551 | 5510 | 84.11% | 5527 | 84.37% | 98 | 852 | 74 | 17 | 0 |
| LIB7135 | TSL2 | 12646 | 10228 | 80.88% | 10234 | 80.93% | 252 | 2014 | 146 | 6 | 0 |
| LIB7135 | TSL3 | 34455 | 29501 | 85.62% | 29542 | 85.74% | 330 | 4256 | 327 | 41 | 0 |
| LIB7135 | TSL4 | 17664 | 15070 | 85.31% | 15103 | 85.50% | 271 | 2106 | 184 | 33 | 0 |
| LIB7135 | TSL5 | 3502 | 2788 | 79.61% | 2793 | 79.75% | 261 | 373 | 75 | 5 | 0 |
| LIB7135 | TSL6 | 33723 | 28880 | 85.64% | 28912 | 85.73% | 429 | 4088 | 294 | 32 | 0 |
| LIB7135 | TSL7 | 24715 | 20970 | 84.85% | 20978 | 84.88% | 350 | 3151 | 236 | 8 | 0 |
| LIB7135 | TSL8 | 4559 | 3799 | 83.33% | 3804 | 83.44% | 115 | 583 | 57 | 5 | 0 |
| LIB7135 | TSL9 | 565 | 420 | 74.34% | 420 | 74.34% | 74 | 53 | 18 | 0 | 0 |
| LIB7136 | TSL10 | 7925 | 4984 | 62.89% | 4996 | 63.04% | 207 | 2638 | 84 | 12 | 0 |
| LIB7136 | TSL11 | 24423 | 20626 | 84.45% | 20649 | 84.55% | 304 | 3218 | 252 | 23 | 0 |
| LIB7136 | TSL12 | 14092 | 10379 | 73.65% | 10399 | 73.79% | 272 | 3238 | 183 | 20 | 0 |
| LIB7136 | TSL13 | 2259 | 1746 | 77.29% | 1746 | 77.29% | 172 | 252 | 89 | 0 | 0 |
| LIB7136 | TSL14 | 1124 | 879 | 78.20% | 879 | 78.20% | 86 | 136 | 23 | 0 | 0 |
| LIB7136 | TSL15 | 746 | 550 | 73.73% | 550 | 73.73% | 74 | 103 | 19 | 0 | 0 |
| LIB7136 | TSL1 | 6510 | 5417 | 83.21% | 5429 | 83.39% | 125 | 872 | 84 | 12 | 0 |
| LIB7136 | TSL2 | 13687 | 11098 | 81.08% | 11109 | 81.16% | 253 | 2163 | 162 | 11 | 0 |
| LIB7136 | TSL3 | 33262 | 28438 | 85.50% | 28486 | 85.64% | 314 | 4105 | 357 | 48 | 0 |
| LIB7136 | TSL4 | 18488 | 15500 | 83.84% | 15541 | 84.06% | 308 | 2442 | 197 | 41 | 0 |

| | | | | | | | | | | | |
|---|---|---|---|---|---|---|---|---|---|---|---|
| LIB7136 | TSL5 | 4111 | 3392 | 82.51% | 3402 | 82.75% | 220 | 421 | 68 | 10 | 0 |
| LIB7136 | TSL6 | 39119 | 33335 | 85.21% | 33369 | 85.30% | 420 | 4928 | 402 | 34 | 0 |
| LIB7136 | TSL7 | 24711 | 20936 | 84.72% | 20949 | 84.78% | 346 | 3205 | 211 | 13 | 0 |
| LIB7136 | TSL8 | 3427 | 2896 | 84.51% | 2898 | 84.56% | 84 | 396 | 49 | 2 | 0 |
| LIB7136 | TSL9 | 670 | 484 | 72.24% | 484 | 72.24% | 69 | 98 | 19 | 0 | 0 |
| LIB7137 | TSL10 | 7990 | 4920 | 61.58% | 4927 | 61.66% | 250 | 2719 | 94 | 7 | 0 |
| LIB7137 | TSL11 | 22174 | 18652 | 84.12% | 18663 | 84.17% | 322 | 2934 | 255 | 11 | 0 |
| LIB7137 | TSL12 | 11947 | 8677 | 72.63% | 8696 | 72.79% | 315 | 2759 | 177 | 19 | 0 |
| LIB7137 | TSL13 | 2741 | 2118 | 77.27% | 2120 | 77.34% | 219 | 299 | 103 | 2 | 0 |
| LIB7137 | TSL14 | 1168 | 876 | 75.00% | 877 | 75.09% | 127 | 146 | 18 | 1 | 0 |
| LIB7137 | TSL15 | 828 | 606 | 73.19% | 606 | 73.19% | 105 | 92 | 25 | 0 | 0 |
| LIB7137 | TSL1 | 4899 | 4137 | 84.45% | 4147 | 84.65% | 105 | 598 | 49 | 10 | 0 |
| LIB7137 | TSL2 | 9417 | 7661 | 81.35% | 7671 | 81.46% | 216 | 1386 | 144 | 10 | 0 |
| LIB7137 | TSL3 | 31266 | 26633 | 85.18% | 26677 | 85.32% | 300 | 3953 | 336 | 44 | 0 |
| LIB7137 | TSL4 | 17446 | 14658 | 84.02% | 14691 | 84.21% | 259 | 2315 | 181 | 33 | 0 |
| LIB7137 | TSL5 | 4391 | 3429 | 78.09% | 3432 | 78.16% | 327 | 562 | 70 | 3 | 0 |
| LIB7137 | TSL6 | 32523 | 27654 | 85.03% | 27671 | 85.08% | 402 | 4173 | 277 | 17 | 0 |
| LIB7137 | TSL7 | 26352 | 22141 | 84.02% | 22158 | 84.08% | 372 | 3613 | 209 | 17 | 0 |
| LIB7137 | TSL8 | 2884 | 2363 | 81.93% | 2367 | 82.07% | 87 | 380 | 50 | 4 | 0 |
| LIB7137 | TSL9 | 619 | 443 | 71.57% | 443 | 71.57% | 82 | 73 | 21 | 0 | 0 |
| 10bp stringency | | | Assigned | % | Assigned | % | Unassigned_ Unmapped | Unassigned _MultiMap ping | Unassigned_ NoFeatures | Unassigned_ Ambiguity | Unassigned_ Ambiguity |
| LIB7135 | TSL10 | 5921 | 4364 | 73.70% | 4377 | 73.92% | 110 | 1363 | 71 | 13 | 0 |
| LIB7135 | TSL11 | 21760 | 18499 | 85.01% | 18507 | 85.05% | 218 | 2813 | 222 | 8 | 0 |
| LIB7135 | TSL12 | 12572 | 9301 | 73.98% | 9310 | 74.05% | 183 | 2919 | 160 | 9 | 0 |
| LIB7135 | TSL13 | 703 | 512 | 72.83% | 513 | 72.97% | 28 | 107 | 55 | 1 | 0 |
| LIB7135 | TSL14 | 1082 | 889 | 82.16% | 889 | 82.16% | 61 | 121 | 11 | 0 | 0 |
| LIB7135 | TSL15 | 524 | 421 | 80.34% | 421 | 80.34% | 29 | 65 | 9 | 0 | 0 |

| | | | | | | | | | | | |
|---|---|---|---|---|---|---|---|---|---|---|---|
| LIB7135 | TSL1 | 5473 | 4585 | 83.77% | 4602 | 84.09% | 81 | 729 | 61 | 17 | 0 |
| LIB7135 | TSL2 | 6426 | 5111 | 79.54% | 5115 | 79.60% | 122 | 1117 | 72 | 4 | 0 |
| LIB7135 | TSL3 | 21092 | 17867 | 84.71% | 17907 | 84.90% | 256 | 2715 | 214 | 40 | 0 |
| LIB7135 | TSL4 | 16863 | 14378 | 85.26% | 14410 | 85.45% | 247 | 2029 | 177 | 32 | 0 |
| LIB7135 | TSL5 | 2008 | 1690 | 84.16% | 1695 | 84.41% | 73 | 213 | 27 | 5 | 0 |
| LIB7135 | TSL6 | 14610 | 12574 | 86.06% | 12599 | 86.24% | 187 | 1672 | 152 | 25 | 0 |
| LIB7135 | TSL7 | 19713 | 16845 | 85.45% | 16853 | 85.49% | 232 | 2453 | 175 | 8 | 0 |
| LIB7135 | TSL8 | 1483 | 1230 | 82.94% | 1235 | 83.28% | 24 | 207 | 17 | 5 | 0 |
| LIB7135 | TSL9 | 415 | 364 | 87.71% | 365 | 87.95% | 14 | 32 | 4 | 1 | 0 |
| LIB7136 | TSL10 | 5662 | 4205 | 74.27% | 4217 | 74.48% | 102 | 1273 | 70 | 12 | 0 |
| LIB7136 | TSL11 | 21975 | 18641 | 84.83% | 18664 | 84.93% | 203 | 2912 | 196 | 23 | 0 |
| LIB7136 | TSL12 | 12148 | 9044 | 74.45% | 9059 | 74.57% | 177 | 2764 | 148 | 15 | 0 |
| LIB7136 | TSL13 | 744 | 549 | 73.79% | 549 | 73.79% | 30 | 105 | 60 | 0 | 0 |
| LIB7136 | TSL14 | 1029 | 847 | 82.31% | 847 | 82.31% | 40 | 125 | 17 | 0 | 0 |
| LIB7136 | TSL15 | 545 | 432 | 79.27% | 432 | 79.27% | 28 | 74 | 11 | 0 | 0 |
| LIB7136 | TSL1 | 5299 | 4393 | 82.90% | 4404 | 83.11% | 93 | 734 | 68 | 11 | 0 |
| LIB7136 | TSL2 | 6963 | 5551 | 79.72% | 5556 | 79.79% | 134 | 1173 | 100 | 5 | 0 |
| LIB7136 | TSL3 | 19969 | 16887 | 84.57% | 16928 | 84.77% | 232 | 2596 | 213 | 41 | 0 |
| LIB7136 | TSL4 | 17593 | 14740 | 83.78% | 14781 | 84.02% | 276 | 2350 | 186 | 41 | 0 |
| LIB7136 | TSL5 | 2581 | 2206 | 85.47% | 2216 | 85.86% | 61 | 271 | 33 | 10 | 0 |
| LIB7136 | TSL6 | 18448 | 15592 | 84.52% | 15615 | 84.64% | 217 | 2413 | 203 | 23 | 0 |
| LIB7136 | TSL7 | 19928 | 16917 | 84.89% | 16929 | 84.95% | 267 | 2570 | 162 | 12 | 0 |
| LIB7136 | TSL8 | 1135 | 958 | 84.41% | 958 | 84.41% | 17 | 148 | 12 | 0 | 0 |
| LIB7136 | TSL9 | 534 | 434 | 81.27% | 434 | 81.27% | 13 | 80 | 7 | 0 | 0 |
| LIB7137 | TSL10 | 5720 | 4209 | 73.58% | 4216 | 73.71% | 89 | 1339 | 76 | 7 | 0 |
| LIB7137 | TSL11 | 19974 | 16903 | 84.63% | 16914 | 84.68% | 198 | 2672 | 190 | 11 | 0 |
| LIB7137 | TSL12 | 10158 | 7500 | 73.83% | 7507 | 73.90% | 164 | 2344 | 143 | 7 | 0 |
| LIB7137 | TSL13 | 770 | 532 | 69.09% | 533 | 69.22% | 52 | 107 | 78 | 1 | 0 |
| LIB7137 | TSL14 | 1081 | 844 | 78.08% | 844 | 78.08% | 83 | 142 | 12 | 0 | 0 |
| LIB7137 | TSL15 | 671 | 552 | 82.27% | 552 | 82.27% | 35 | 73 | 11 | 0 | 0 |

| | | | | | | | | | | | |
|---|---|---|---|---|---|---|---|---|---|---|---|
| LIB7137 | TSL1 | 4107 | 3476 | 84.64% | 3486 | 84.88% | 76 | 504 | 41 | 10 | 0 |
| LIB7137 | TSL2 | 4955 | 3958 | 79.88% | 3963 | 79.98% | 94 | 819 | 79 | 5 | 0 |
| LIB7137 | TSL3 | 18218 | 15326 | 84.13% | 15364 | 84.33% | 218 | 2417 | 219 | 38 | 0 |
| LIB7137 | TSL4 | 16709 | 14038 | 84.01% | 14071 | 84.21% | 234 | 2230 | 174 | 33 | 0 |
| LIB7137 | TSL5 | 2420 | 1997 | 82.52% | 2000 | 82.64% | 75 | 324 | 21 | 3 | 0 |
| LIB7137 | TSL6 | 12709 | 10713 | 84.29% | 10721 | 84.36% | 176 | 1696 | 116 | 8 | 0 |
| LIB7137 | TSL7 | 21136 | 17858 | 84.49% | 17873 | 84.56% | 246 | 2840 | 177 | 15 | 0 |
| LIB7137 | TSL8 | 864 | 716 | 82.87% | 716 | 82.87% | 17 | 117 | 14 | 0 | 0 |
| LIB7137 | TSL9 | 434 | 365 | 84.10% | 365 | 84.10% | 11 | 50 | 8 | 0 | 0 |
| | | Min | | 61.58% | | 61.66% | | | | | |
| | | Mean | | 80.32% | | 80.43% | | | | | |
| | | Max | | 87.71% | | 87.95% | | | | | |
