## Supplemental Table S2 for "Deep evolutionary origin of nematode SL2 *trans*-splicing revealed by genome-wide analysis of the *Trichinella spiralis* transcriptome"

**Supplementary Table S2. Transcripts derived from *T. spiralis* predicted downstream operonic genes show strict association with SL2-type *trans*-splicing.**

The table shows the association between putative *T.* spiralis (*Tsp*) operonic genes and the two categories of spliced leader: SL1-type (*Tsp*-SL1, 3-9, 11, 13-15) or SL2-type (*Tsp*-SL2, 10, 12). 'None' indicates transcripts that are not *trans*-spliced. $ *Tsp-uxt-1* transcripts receive equal numbers of SL1- and SL2-type spliced leaders, indicating that is gene likely contains an internal promoter as has been reported for some *C. elegans* operons (Williams et al., 1999). Operons were predicted based on syntenic gene pairs conserved between *T. spiralis* and *C. elegans* (blue text). Shading indicates genes belonging to the same operon; unshaded genes are not part of the same operon or are monocistronic. *T. spiralis* operons (TSPOP) are assigned an arbitrary number based on the BRAKER-Trinity gene set, using 8 bp *Tsp-SL* screening stringency and exon-based gene prediction (see Methods). TSPOP operons indicated by an asterisk were defined by manual annotation alone, since they were not computationally predicted. Note that TSPOP53 and 110 were identified experimentally (Pettitt et al., 2014). *T. spiralis* genes were named based on either their *C. elegans* or human orthologues, where applicable. NDO – no detectable orthologue; NA – not applicable.

| TSPOP | *Tsp* Gene | SL-type | *Cel* Gene | CEOP |
|---|---|---|---|---|
| 1 | *ulp-3* | SL1 | *ulp-3* | 4052 |
| | *shq-1* | None | *Y48A5A.1* | |
| | *Tsp-07337* | SL2 | NDO | NA |
| 5 | *arhgap-27* | SL1 | *tag-325* | 3220 |
| | *mettl-5* | SL2 | *C38D4.9* | |
| 10 | *snu-23* | SL1 | *snu-23* | 3452 |
| | *magt-1* | SL2 | *ZK686.3* | |
| | *csl-4* | SL2 | NDO | NA |
| 11 | *ulp-1* | SL1 | *ulp-1* | 3260 |
| | *slc-25.A15* | SL2 | *T10F2.2* | |
| 21 | *rcl-1* | SL1 | *ZK1127.5* | 2280 |
| | *rnaseh-2C* | SL2 | *ZK1127.13* | |
| 38 | *lpd-5* | SL1 | *lpd-5* | 1176 |
| | *ZK973.9* | SL2 | *ZK973.9* | |
| 47 | *tmem-167B* | None | *Y62E10A.2* | 4540 |
| | *fdxr-1* | None | *Y62E10A.6* | |
| | *cpped-1* | SL2 | NDO | NA |
| 53 | *cpt-2* | None | *cpt-2* | 4424 |
| | *nuaf-3* | SL2 | *nuaf-3* | |
| 61 | *sel-1* | SL1 | *sel-1* | 5356 |
| | *mrps-5* | SL2 | *mrps-5* | |
| 88 | *rom-4* | None | *rom-4* | |
| | *rwdd-1* | SL2 | *T26E3.4* | 1672 |
| | *par-6* | SL2 | *par-6* | |
| 94 | *serinc-2* | SL1 | *Y57E12AL.1* | 5534 |
| | *tmem-258* | SL2 | *Y57E12AM.1* | |
| 103 | *rpl-20* | SL1 | *rpl-20* | 4148 |
| | *cyc-2.1* | SL2 | *cyc-2.1* | |
| 110 | *zgpa-1* | None | *zgpa-1* | 4270 |
| | *dif-1* | SL2 | *dif-1* | 4228 |

| | | | | |
|---|---|---|---|---|
| 117 | *nprl-1* | SL1 | *nprl-1* | 4280 |
| | *uxt-1* | SL1/SL2[$] | *F35H10.6* | |
| | *emc-3* | SL2 | *emc-3* | 4544 |
| | *nme-5* | SL2 | NDO | NA |
| 123 | *fam-210A* | SL1 | *Y56A3A.22* | 3744 |
| | *nft-1* | SL2 | *nft-1* | |
| 124 | *ubxn-2* | SL1 | *ubxn-2* | 4068 |
| | *nfda-12* | SL2 | *Y94H6A.8* | |
| 139 | *lis-1* | SL1 | *lis-1* | - |
| | *mrps-17* | SL2 | *mrps-17* | 3372 |
| | *pmrp-1* | SL2 | *C05D11.9* | |
| 176 | *cyy-1* | None | *cyy-1* | 3522 |
| | *ptihd-1* | SL2 | *ZK353.9* | |
| 182 | *snx-13* | SL1 | *snx-13* | 4632 |
| | *fn3k-1* | SL2 | *Y116A8C.25* | |
| 204 | *rps-29* | SL1 | *rps-29* | 3036 |
| | *trpp-11* | SL2 | *trpp-11* | |
| 206 | *clec-16A* | SL1 | *gop-1* | 3272 |
| | *gpn-1* | SL2 | *gop-2* | |
| | *esyt-2* | SL2 | *esyt-2* | - |
| | *tfbm-1* | SL2 | *tfbm-1* | 1120 |
| 214 | *isy-1* | SL1 | *isy-1* | - |
| | *rab-51.F* | SL2 | *F44E7.9* | 5084 |
| | *b3glct-1* | SL2 | *ZC250.2* | |
| 229 | *let-92* | none | *let-92* | 4480 |
| | *mob-1* | SL2 | *mob-1* | |
| 230 | *eogt-1* | SL1 | *H12D21.10* | - |
| | *ndufaf-2* | SL2 | *Y116A8C.30* | 4596 |
| | *atpaf-2* | SL2 | *Y116A8C.27* | |
| 263 | *exos-4.1* | SL1 | *exos-4.1* | 4532 |
| | *vars-1* | SL2 | *vars-1* | 5176 |
| | *algn-12* | SL2 | *algn-12* | |
| 272 | *uqcr-11* | SL1 | NDO | NA |
| | *spg-20* | SL2 | *spg-20* | 1332 |
| | *abt-1* | SL2 | *F57B10.8* | |
| 274 | *snrk-1* | SL1 | *ZK524.4* | 1396 |
| | *lars-2* | SL2 | *lars-2* | |
| 275 | *mafr-1* | SL1 | *mafr-1* | 1396 |
| | *arch-1* | SL2 | *arch-1* | |
| 279 | *inx-12* | SL1 | *inx-12* | 1108 |
| | *inx-13* | SL2 | *inx-13* | |
| 282 | *sgn-1* | None | *sgn-1* | - |
| | *pigm-1* | SL2 | *pigm-1* | 2532 |
| | *dph-5* | SL2 | *B0491.7* | |
| 283 | *trpp-8* | None | *trpp-8* | 1264 |
| | *vha-10* | SL2 | *vha-10* | |

| | | | | |
|---|---|---|---|---|
| 284 | *zipt-1*<br>*kiaa-100* | SL1<br>SL2 | *zipt-1*<br>*F31C3.3* | 1776 |
| 305 | *zipt-11*<br>*mrpl-24* | SL1<br>SL2 | *zipt-11*<br>*mrpl-24* | 1256 |
| 308 | *apg-1*<br>*hpo-13* | None<br>SL2 | *apg-1*<br>*hpo-13* | 1904 |
| 337 | *mrps-27*<br>*blos-9* | SL1<br>SL2 | *mrps-27*<br>*blos-9* | 1764 |
| 338 | *mrps-22*<br>*algn-9* | SL1<br>SL2 | *mrps-22*<br>*algn-9* | 2496 |
| 342 | *lst-6*<br>*sqv-7* | SL1<br>SL2 | *lst-6*<br>*sqv-7* | 2276 |
| 345 | *ncap-1*<br>*ubc-26*<br>*hpo-20* | SL1<br>SL2<br>SL2 | *ncap-1*<br>*ubc-26*<br>*hpo-20* | 2076<br>2072 |
| 347 | *tmem-222*<br>*mog-2* | SL1<br>SL2 | *H20J04.6*<br>*mog-2* | 2124 |
| 348 | *rtraf-1*<br>*adk-6* | SL1<br>SL2 | *E02H1.5*<br>*E02H1.6* | 2436 |
| 379* | *ribo-1*<br>*impdh-2* | SL1<br>SL2 | *ribo-1*<br>*T22D1.3* | 4204 |
| 381* | *unc-32*<br>*tpk-1* | SL1<br>SL2 | *unc-32*<br>*tpk-1* | 3556 |
| 382* | *cul-1*<br>*colg-1*<br>*stk-16* | SL1<br>SL2<br>SL2 | *cul-1*<br>*D2045.9*<br>*D2045.7* | 3676 |
| 383* | *slc-25.A3*<br>*pigq-1* | SL1<br>SL2 | *F01G4.6*<br>*pigq-1* | 4416 |
| 384* | *csn-6*<br>*mipep-1* | SL1<br>SL2 | *csn-6*<br>*Y67H2A.7* | 4538 |
