## Supplemental Figure S1 for "Deep evolutionary origin of nematode SL2 *trans*-splicing revealed by genome-wide analysis of the *Trichinella spiralis* transcriptome"

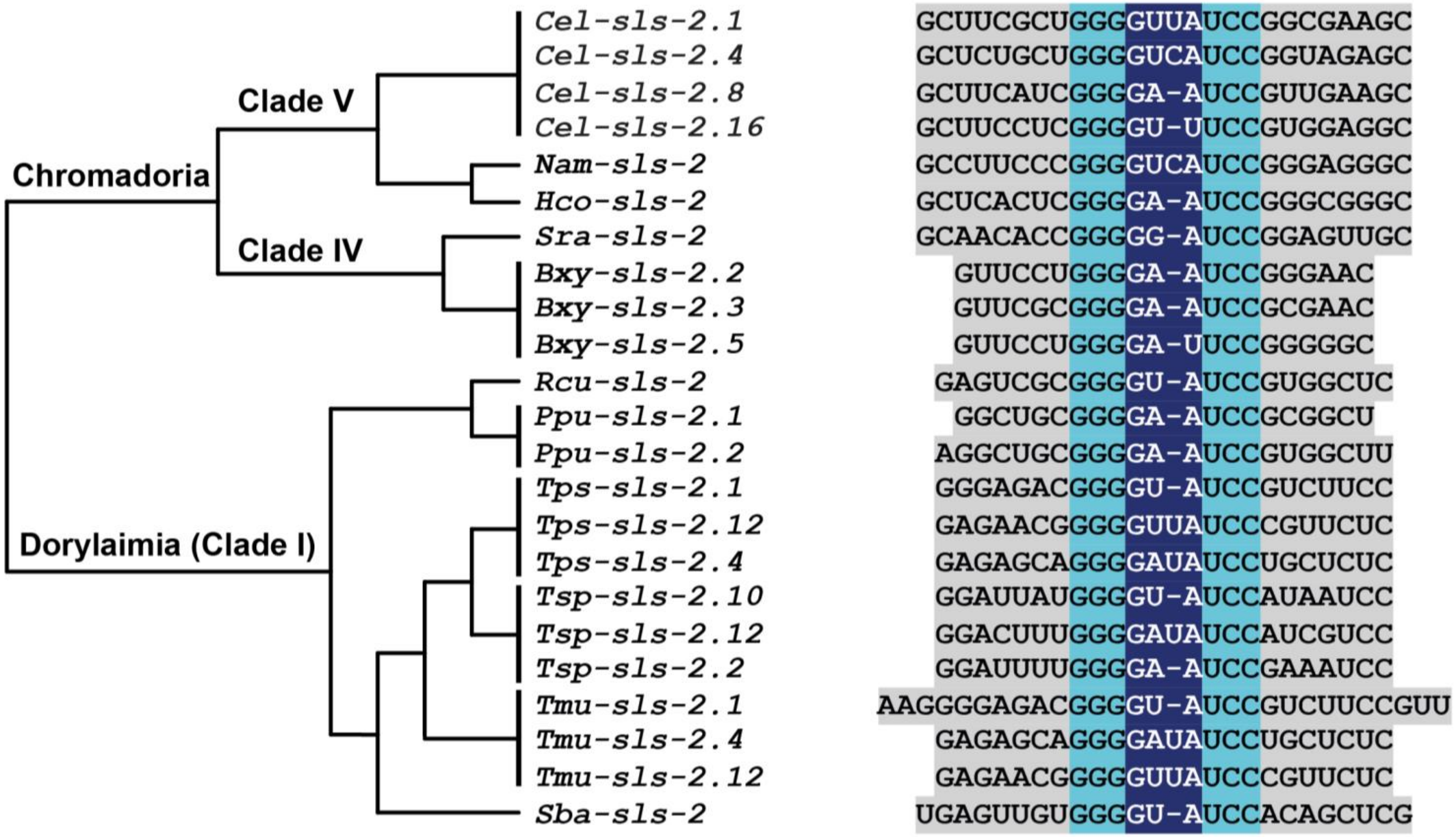


**Supplementary Figure 1. SL2-type stem-loop III motif is conserved across multiple nematode clades.** Representative primary sequences of the stem-loop III motif are shown. Grey countershading indicates non-conserved residues that form part of the stem, whereas light blue indicates the conserved residues of the stem. Residues predicted to form the loop, are counter shaded dark-blue. The genes encoding each SL2-type RNA are indicated, named on the basis of *C. elegans* nomenclature, thus *Tsp*-SL2, SL10 and SL12 are encoded by *Tsp-sls-2.2*, *Tsp-sls-2.10* and *Tsp-sls-2.12*, respectively. The phylogram shows the evolutionary relationship between the species from which the genes were derived. *Cel* - *C. elegans*; *Nam* - *Necator americanus*; *Hco* - *Haemonchus contortus*; *Sra* - *Strongyloides ratti*; *Bxy* - *Bursaphelenchus xylophilus*; *Rcu* - *Romanomermis culicovorax*; *Tps* - *Trichinella pseudospiralis*; *Tsp* - *T. spiralis*; *Tmu* - *Trichuris muris*; *Sba* - *Soboliphyme baturini*.
