## Supplemental Figure S2 for "Deep evolutionary origin of nematode SL2 *trans*-splicing revealed by genome-wide analysis of the *Trichinella spiralis* transcriptome"

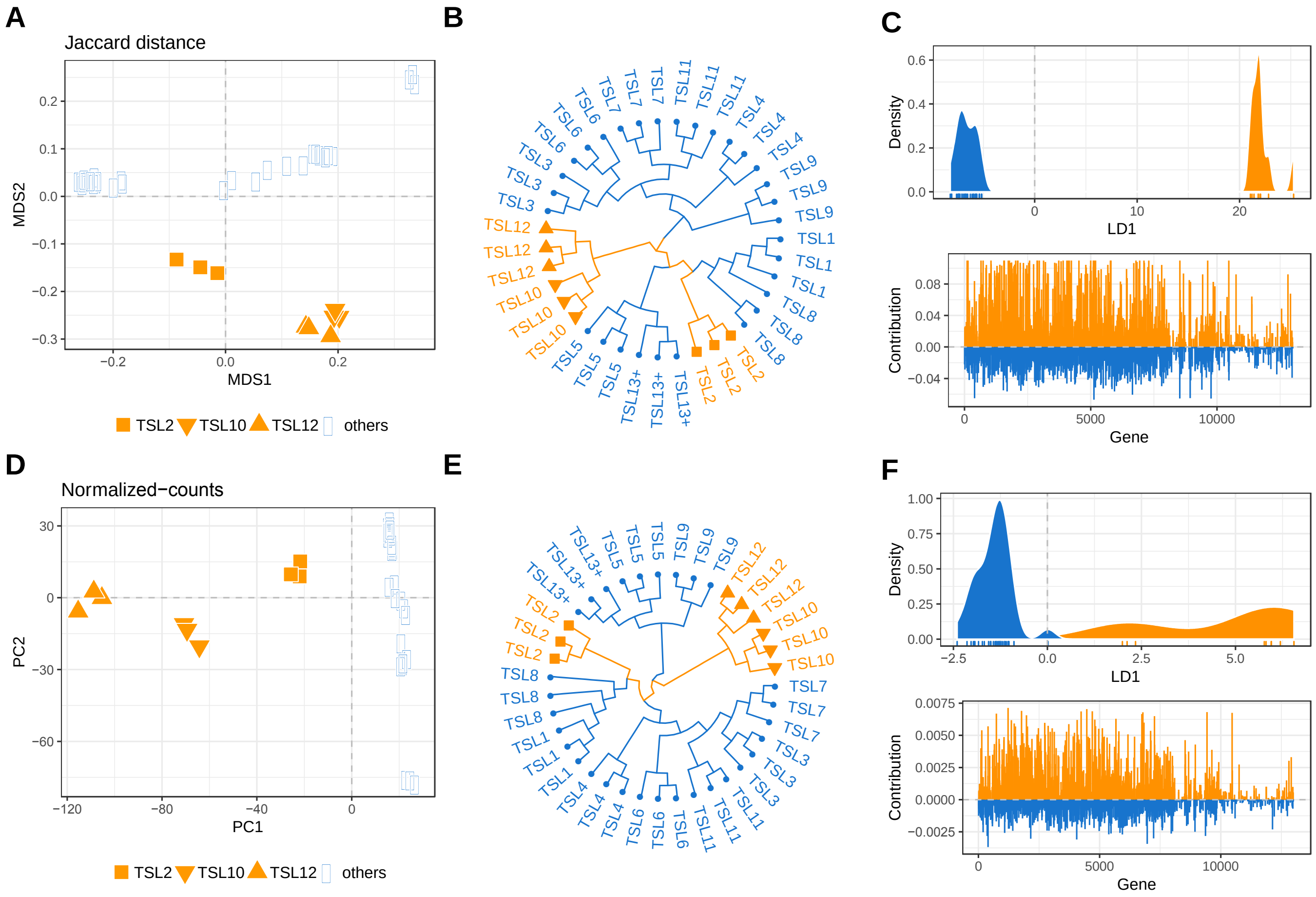


**Supplementary Figure 2. Multivariate analysis of *trans*-splicing events highlighting possible functional sub-division among SL2-type trans-spliced leaders in *T. spiralis*.**

(A, D) Ordinations (MDS or PCA) of *Tsp*-SL (TSL) libraries based on gene-based Jaccard (A) or normalized-counts distances (D). (B, E) Hierarchical clustering (Ward's criterion) of these distances. (C, F) The corresponding densities of the first linear discriminant function among SL1-type and SL2-type *Tsp*-SL libraries and the contribution of each gene to the group discrimination (DAPC). Data are based on uncorrected *de novo* gene annotations (STRINGTIE) and 8-bp *Tsp*-SL read classification stringency. *Tsp*-SL is abbreviated to 'TSL' for clarity.
